# Machine learning enables accurate and rapid prediction of active molecules against breast cancer cells

**DOI:** 10.1101/2021.09.06.459060

**Authors:** Shuyun He, Duancheng Zhao, Yanle Ling, Hanxuan Cai, Yike Cai, Jiquan Zhang, Ling Wang

## Abstract

**Summary:** Breast cancer (BC) has surpassed lung cancer as the most frequently occurring cancer, and it is the leading cause of cancer-related death in women. Therefore, there is an urgent need to discover or design new drug candidates for BC treatment. In this study, we first collected a series of structurally diverse datasets consisting of 33,757 active and 21,152 inactive compounds for 13 breast cancer cell lines and one normal breast cell line commonly used in *in vitro* antiproliferative assays. Predictive models were then developed using five conventional machine learning algorithms, including naïve Bayesian, support vector machine, k-Nearest Neighbors, random forest, and extreme gradient boosting, as well as five deep learning algorithms, including deep neural networks, graph convolutional networks, graph attention network, message passing neural networks, and Attentive FP. A total of 476 single models and 112 fusion models were constructed based on three types of molecular representations including molecular descriptors, fingerprints, and graphs. The evaluation results demonstrate that the best model for each BC cell subtype can achieve high predictive accuracy for the test sets with AUC values of 0.689–0.993. Moreover, important structural fragments related to BC cell inhibition were identified and interpreted. To facilitate the use of the model, an online webserver called ChemBC and its local version software were developed to predict potential anti-BC agents.

**Availability:** ChemBC webserver is available at http://chembc.idruglab.cn/ and its local version Python software is maintained at a GitHub repository (https://github.com/idruglab/ChemBC).

**Supplementary information:** Supplementary data are available at Bioinformatics online.

## 1 Introduction

According to the latest data on the global cancer burden for 2020 released by the International Agency for Research on Cancer of the World Health Organization, breast cancer (BC) surpassed lung cancer in 2020 to become the most common cancer worldwide. BC is the leading cause of cancer-related death among women worldwide (Sung, et al., 2021). BC consists of the uncontrolled proliferation of mammary epithelial cells under the action of many carcinogenic factors (Escala-Garcia, et al., 2020), including alcohol consumption, smoking, overweight, and mammographic density. BC is classified according to the expression of the estrogen receptor (ER), progesterone receptor (PR), human epidermal growth factor receptor 2 (HER2), and Ki-67 into five subtypes: Luminal A, Luminal B (HER2-positive or HER2-negative), HER2-positive, and triple-negative breast cancer (TNBC) (Harbeck, et al., 2013). Among these BC subtypes, TNBC is associated with poor survival mediated by treatment resistance, and it is the most difficult to treat with curative intent (Liao, et al., 2021). Several drugs (e.g., anthracyclines and trastuzumab) have been approved by the U.S. Food and Drug Administration (FDA) for the treatment of BC; however, issues such as poor efficacy, toxicity, adverse drug reactions, and the emergence of drug resistance have limited their clinical use (Brower, 2013; Cameron, et al., 2017; Daniyal, et al., 2021; Li and Li, 2021; Shah and Gradishar, 2018). Therefore, there is an urgent need to discover and develop new drugs for the treatment of BC, especially for TNBC.

Increased amounts of phenotypical pharmacological data on cancer, Alzheimer’s disease, and infectious diseases have been accumulated in the past three decades. Inspired by the available phenotypic screening data, several efficient and cost-saving computational models have been developed to accelerate the drug design and discovery process (Buckner, et al., 2020; Chandrasekaran, et al., 2020; Malandraki-Miller and Riley, 2021; Zoffmann, et al., 2019). For example, in 2020, Stokes et al. first reported directed message passing neural network models using a collection of 2,335 compounds for those that inhibited the growth of *Escherichia coli* (phenotype screening data) and then identified the lead compound halicin with broad-spectrum antibacterial activity (Stokes, et al., 2020). Other machine learning-based models have been established to identify new agents against Methicillin-Resistant *Staphylococcus aureus* (Wang, et al., 2014), *Mycobacterium tuberculosis* (Ye, et al., 2021), *Pseudomonas aeruginosa* (Fields, et al., 2020), *Plasmodium falciparum* (Ashdown, et al., 2020), and *Schistosoma* (Zheng, et al., 2021). In the field of anticancer drug design and discovery, phenotypical whole cell-based screening methods have substantially advanced our ability to identify new anticancer drugs. In previous studies, we reported the development of computational models using integrated NCI-60 cell-based phenotype screening data to identify new anticancer agents (e.g., **G03** and **I2**) with significant inhibitory activity against various cancer cell lines (Guo, et al., 2019; Luo, et al., 2019). Although the reported integrated computational anticancer models provided valuable data for discovering anticancer agents, these models cannot distinguish or selectively predict specific cancer cell subtypes (such as BC and its subtypes). In addition, these prediction models have not been developed into easy-to-use tools (e.g., local software packages or online prediction platforms), which limits the use of these models by practitioners in the field.

In the present study, we expanded our earlier efforts aimed at developing reliable computational cell-based models to predict cell inhibitory activity in BC and subtypes and provided a free platform to share our models. A total of 588 cell-based models for BC and subtypes were developed using five conventional machine learning (ML) and five deep learning (DL) algorithms based on three major types of molecular descriptors, fingerprints, and graphs. We used the local outlier factor (LOF) (Breunig, et al., 2000) algorithm to evaluate the applicability domain of the best model for each BC cell line and applied the SHapley Additive exPlanations (SHAP) (Lundberg and Lee, 2017; Lundberg, et al., 2020) algorithm to highlight important structural fragments. Finally, an online platform (ChemBC) and local software were constructed based on reliable models to contribute to future research.

## 2 Methods

### 2.1 Dataset collection and preparation

All quantitative compound-cell associations (cell-based assays, assay type: F) for available BC cell lines and normal BC cell lines were collected from ChEMBL (Mendez, et al., 2019) (downloaded in March 2021) after the exclusion of metastatic cell lines. Each BC cell dataset was then processed using the following steps: (i) compounds with biological activity reported as IC_50_, EC_50_, or GI_50_ were kept, whereas molecules that had no bioactivity record were removed; (ii) the units of bioactivity (i.e., g/mL, M, nM) were converted into the standard unit in μM; (iii) for a molecule with multiple bioactivity values, the final bioactivity value was obtained by averaging the available bioactivity records; (iv) according to previous studies (Fields, et al., 2020; Ye, et al., 2021), compounds with bioactivity values (e.g., IC_50_, EC_50_, GI_50_) ≤10 μM were considered as active and vice versa; molecules whose label could not be unequivocally assigned (e.g., activity <100 μM or activity >1 μM) were excluded from the dataset; (v) all molecules were processed by removing salt and optimized based on the MMFF94X force field using MOE software (version 2018) with the default parameters. Finally, 14 cell lines with the number of active molecules (actives) and inactive molecules (inactives) >50 were retained. Each cell-compound dataset was randomly split into three sub-datasets: training (80%), validation (10%), and test (10%).

### 2.2 Molecular representations calculation

Choosing suitable molecular representations is essential for developing acceptable and robust QSAR models. To a certain extent, the molecular representation determines the upper limit of the accuracy of the model. To fully characterize the chemical information of these molecules, three distinct types of features were calculated and used, including molecular descriptors-, fingerprints-, and graph-based representations. RDKit descriptors (RDKitDes), a set of 208 descriptors, were used. Four fingerprint-based features including Morgan fingerprints (ECFP-like, 1024-bits) (Rogers, et al., 2010), MACCS keys (166-bits) (Durant, et al., 2002), AtomParis fingerprints (1024-bits) (Carhart, et al., 1985), and 2D Pharmacophore Fingerprints (PharmacoPFP, 38-bits) (Gobbi, et al., 1998) were implemented. The molecular descriptor- and fingerprint-based representations were calculated using RDKit (Landrum, 2016) (version: 2020.03.1). Molecular graph-based representations were generated using Deepchem (version: 2.5.0).

### 2.3 Machine learning algorithms

Five conventional ML algorithms (i.e., RF, SVM, XGBoost, KNN, and NB) and five DL algorithms (i.e., DNN, GCN, GAT, MPNN, and Attentive FP) were used to develop classification models for discriminating actives from inactives against breast cell lines.

The RF, SVM, KNN, and NB models were constructed using the Scikit-learn (Pedregosa, et al., 2011) python package; the XGBoost (Chen and Guestrin, 2016) models were developed using the XGBoost python package; and other graph-based models were established using the DeepChem python package (https://deepchem.io/). The detailed machine learning methods can be found in Part I of the Supplementary Material.

### 2.4 Performance evaluation of models

The following classification evaluation metrics were used to evaluate the performance of the classification models: specificity (SP/TNR), sensitivity (SE/TPR/Recall), accuracy (ACC), F1-measure (F1 score), Matthews correlation coefficient (MCC), the area under the receiver operating characteristic (AUC), and Balanced accuracy (BA).

These evaluation metrics are defined as follows:

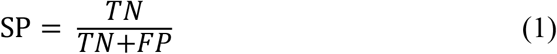

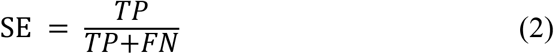

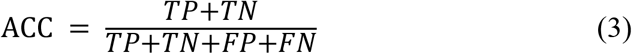

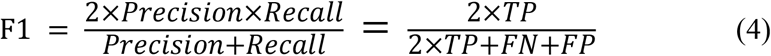

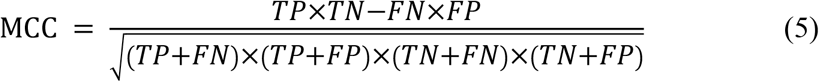

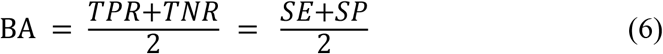

where TP, TN, FP, and FN represent the number of true positives, true negatives, false positives, and false negatives, respectively.

### 2.5 Model interpretation

We used a recently-developed model-agnostic interpretation framework termed SHapley Additive exPlanation (SHAP) to interpret the established ML models presented in this study. Inspired by the idea of cooperative game theory, the SHAP method constructs an additive explanatory model. In this model, all features are considered contributors. For each prediction sample, the model generates a predicted value, and the SHAP value is the value assigned to each feature in the sample. The greater the SHAP value, the greater the contribution of the corresponding feature to the ML model.

### 2.6 Model applicability domain

We used the LOF algorithm (Breunig, et al., 2000) to detect super-applicability domain compounds for the best model for each BC or normal breast cell line. LOF is based on the concept of local density, where the local area is given by k-nearest neighbors, whose distance is used to estimate the density. Regions of similar density can be identified by comparing the local density of an object with that of its neighbors, and points that are much lower in density than their neighbors are considered outliers.

## 3 Results and discussions

### 3.1 Dataset analysis and model construction

According to the above-predefined criteria, 14 breast-associated cell lines were obtained and distributed as follows: (1) two Luminal A subtypes including MCF-7 and T-47D; (2) two Luminal B subtypes including BT-474 and MDA-MB-361; (3) three HER-2+ subtypes including MDA-MB-435, MDA-MB-453, and SK-BR-3; (4) six TNBC subtypes including Bcap37, BT-20, BT-549, HS-578T, MDA-MB-231, and MDA-MB-468; and (5) one normal breast cell line, HBL-100. Accordingly, we selected these cell-based phenotypical datasets for subsequent modeling. Details on the 14 cell lines and their corresponding cell-associated compound datasets are summarized in Table 1. The compiled cell-based phenotype datasets included 34,801 unique compounds and 54,909 cell–compound associations. Among them, in 14 cell line datasets, 33,757 compounds were labeled as actives and 21,152 compounds were labeled as inactives (Fig. S1A). Fig. S1B shows the proportions of actives and inactives in the 14 cell datasets (due to the natural, although it may not be the best, we did not add theoretical decoys to deliberately balance the data), with active compounds accounting for approximately 40%–78%.

**Table 1.** Breast cell line datasets used in this study.

| Cell lines | Classification | No. of compounds | No. of scaffolds | scaffolds/compounds (%) |
| --- | --- | --- | --- | --- |
| MDA-MB-435 | HER-2+<sup>a</sup> | 3030 | 870 | 28.71% |
| MDA-MB-453 | HER-2+ | 440 | 215 | 48.86% |
| SK-BR-3 | HER-2+ | 2026 | 571 | 28.18% |
| MCF-7 | Luminal A<sup>b</sup> | 29378 | 5787 | 19.70% |
| T-47D | Luminal A | 3135 | 926 | 29.54% |
| BT-474 | Luminal B<sup>c</sup> | 811 | 308 | 37.98% |
| MDA-MB-361 | Luminal B | 367 | 196 | 53.41% |
| HBL-100 | Normal cell line | 316 | 110 | 34.81% |
| Bcap37 | TNBC<sup>d</sup> | 275 | 73 | 26.55% |
| BT-20 | TNBC | 292 | 146 | 50.00% |
| BT-549 | TNBC | 1182 | 497 | 42.05% |
| HS-578T | TNBC | 469 | 215 | 45.84% |
| MDA-MB-231 | TNBC | 11202 | 2672 | 23.85% |
| MDA-MB-468 | TNBC | 1986 | 685 | 34.49% |

<sup>a</sup> HER-2+: HER2-positive breast cancers. <sup>b</sup> Luminal A: Luminal A breast cancer is hormone-receptor positive (estrogen-receptor and/or progesterone-receptor positive), HER2-negative, and has low levels of the protein Ki-67, which helps control how fast cancer cells grow. <sup>c</sup> Luminal B: Luminal B breast cancer is hormone-receptor positive (estrogen-receptor and/or progesterone-receptor positive), and either HER2 positive or HER2 negative with high levels of Ki-67. <sup>d</sup> TNBC: triple-negative breast cancer.

The structural diversity and chemical space of compounds in datasets play a key role in the predictive ability of the ML models. Bemis–Murcko scaffold analysis (Bemis and Murcko, 1996) showed that the proportion of the scaffolds for each BC cell line dataset was between 19.70% and 53.41% (Table 1), suggesting that the anti-BC compounds of each cell line were structurally more diverse. In addition, the chemical space of the compounds in each dataset can be depicted in a two-dimensional space using molecular weight (MW) and AlogP. As shown in Fig. S2, the training, validation, and test set compounds were distributed over a wide range of MW (108.10–5714.45) and AlogP (−55.54–42.62), demonstrating that the compounds in the modeling datasets have a broad chemical space. Based on the three different types of molecular features (i.e., molecular descriptors-, fingerprints-, and graph-based features) and the selected ten types of ML algorithms, 476 single models and 112 fusion models were developed.

All models were optimized based on the validation sets and selected based on the F1 score (Kc, et al., 2021). The best models were selected for the evaluation of external test datasets. The performance of the established models is discussed in the following sections.

### 3.2 Performance of descriptor-based prediction models for breast-associated cells

Firstly, 84 predictive models were constructed based on the RDKit-descriptors using five traditional types of ML algorithms (KNN, NB, RF, SVM, and XGBoost) and one deep learning DNN method. For these traditional ML methods, the optimized RDKit-descriptors were obtained using the SelectPercentile module (Percentile = 30) implemented in the scikit-learn package and then used as input features to construct models. Each model is denoted as a combination of a given molecular representation and ML algorithm (e.g., RF::RDKitDes). For each cell dataset and the corresponding ML methods, hyperparameters were optimized based on the validation sets (detailed in the Methods section), and the best set of hyperparameters are shown in Table S1. The detailed performance results for descriptor-based models are listed in Table S2. The performance of the models (F1 score, BA, and AUC) for the test sets is summarized in Fig. S3. Overall, most descriptor-based models performed well in BC cell inhibitory prediction tasks, achieving a mean F1 score and BA value >0.5. The RF model performed the best in all cell lines, with higher average F1 scores (0.840 ± 0.073), BA (0.725 ± 0.073), and AUC (0.835 ± 0.067). Meanwhile, the XGBoost model also achieved good and/or comparable performance results (Fig. S3). The detailed best-performing RF::RdkitDes models results were achieved in five breast cancer cell lines (BT-20, HS-578T, MCF-7, MDA-MB-231, and T-47D), while the XGBoost::RDKit models also showed superior performance in five breast-associated cell lines (BT-474, HBL-100, MDA-MB-453, MDA-MB-468, and SK-BR-3). The KNN::RDKitDes models exhibited the best performance in the Bcap37, MDA-MB-361, and MDA-MB-435 cell lines. The SVM::RDKit models performed well in BT-549.

### 3.3 Performance of fingerprint-based prediction models for breast-associated cells

There were 336 models developed based on four types of fingerprints (Morgan, MACCS, Atompairs, and PharmacoPFP) using six types of ML algorithms (KNN, NB, RF, SVM, XGBoost, and DNN). The detailed performance results for fingerprint-based models are listed in Table S3-S6. The F1, AUC, and BA values of the test sets are shown in Fig. 2 and Fig. S4, S5. Taking the average F1 score as a point metric into consideration, the numbers of cell lines for which each model was identified as the best-performing are shown in Fig. 3. No model, fingerprint, or ML algorithm could be identified as the best-performing for the 14 cell line datasets, demonstrating that it is necessary to screen different fingerprints and different ML algorithms for the current breast cell-associated modeling datasets (Fig. 3B–F). Although the characteristics of the four molecular fingerprints are different, the RF models performed better than the other five ML models against most of the 14 cell lines (Fig. 2, 3A, and S4). Meanwhile, the Morgan fingerprint represents the best molecular feature representation because the ML models based on Morgan fingerprints achieved the best results for these modeling datasets (Table 2). Global analysis of four fingerprint-based models also demonstrated that RF methods can achieve a better performance than other ML methods, with the highest average F1 score (0.848 ± 0.006), BA (0.750 ± 0.013), and AUC (0.853 ± 0.009).

**Fig. 1.**
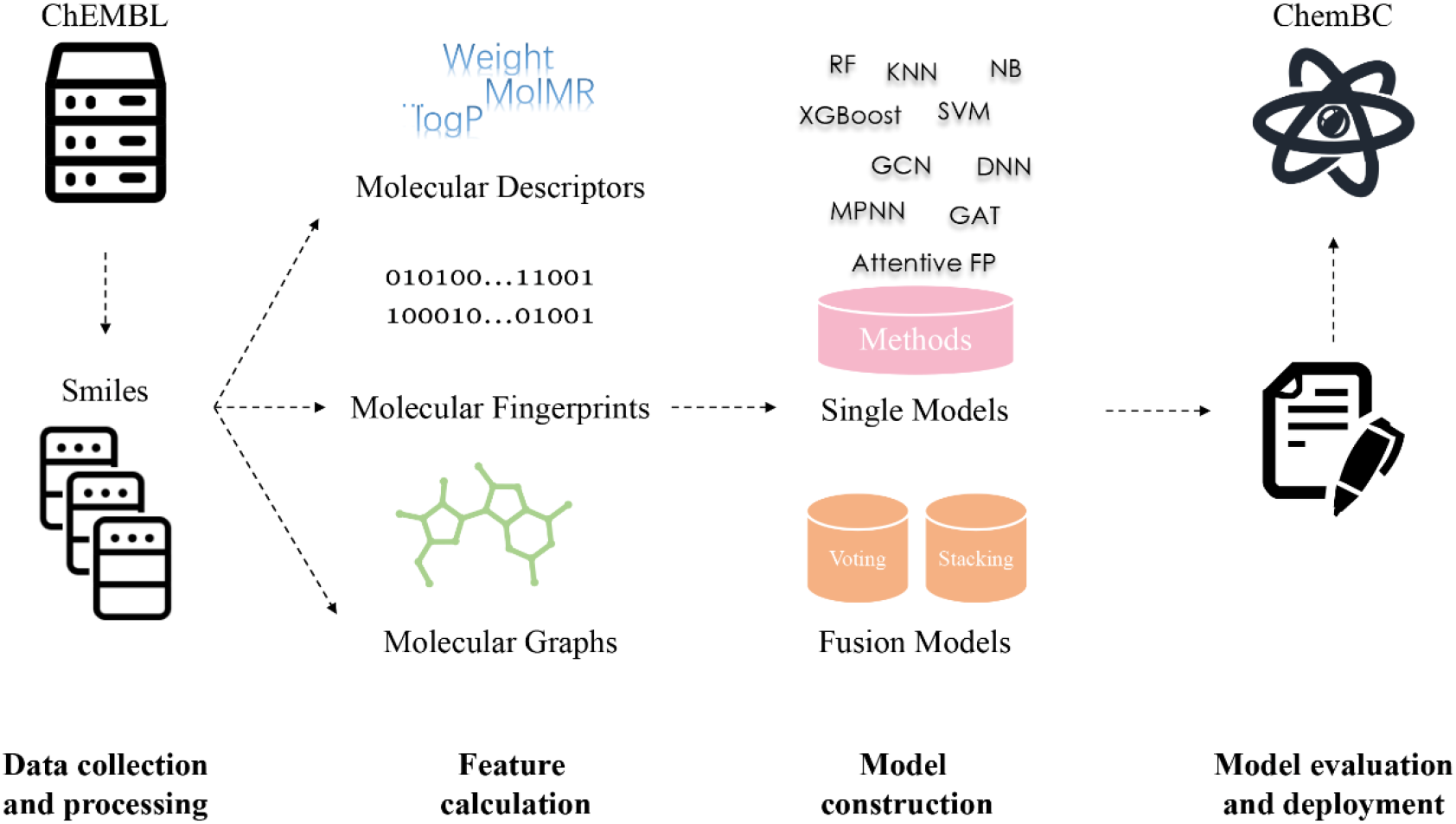
Model construction pipeline.

**Fig. 2.**
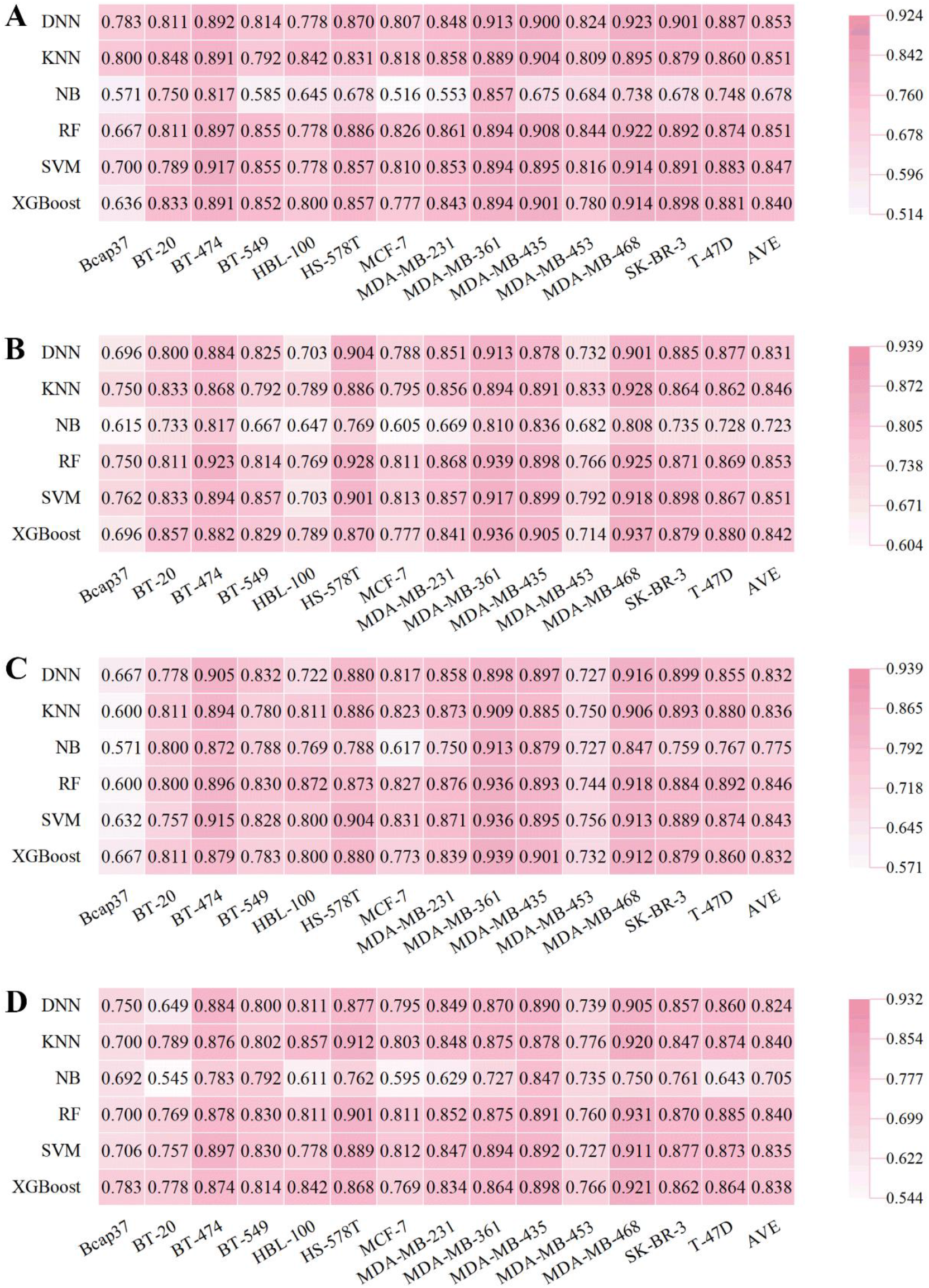
Performance of fingerprint-based BC prediction models. (A) F1 scores of the AtomPairs-based models. (B) F1 scores of the MACCS-based models. (C) F1 scores of the Morgan-based models. (D) F1 scores of the PharmacoPFP-based models.

**Fig. 3.**
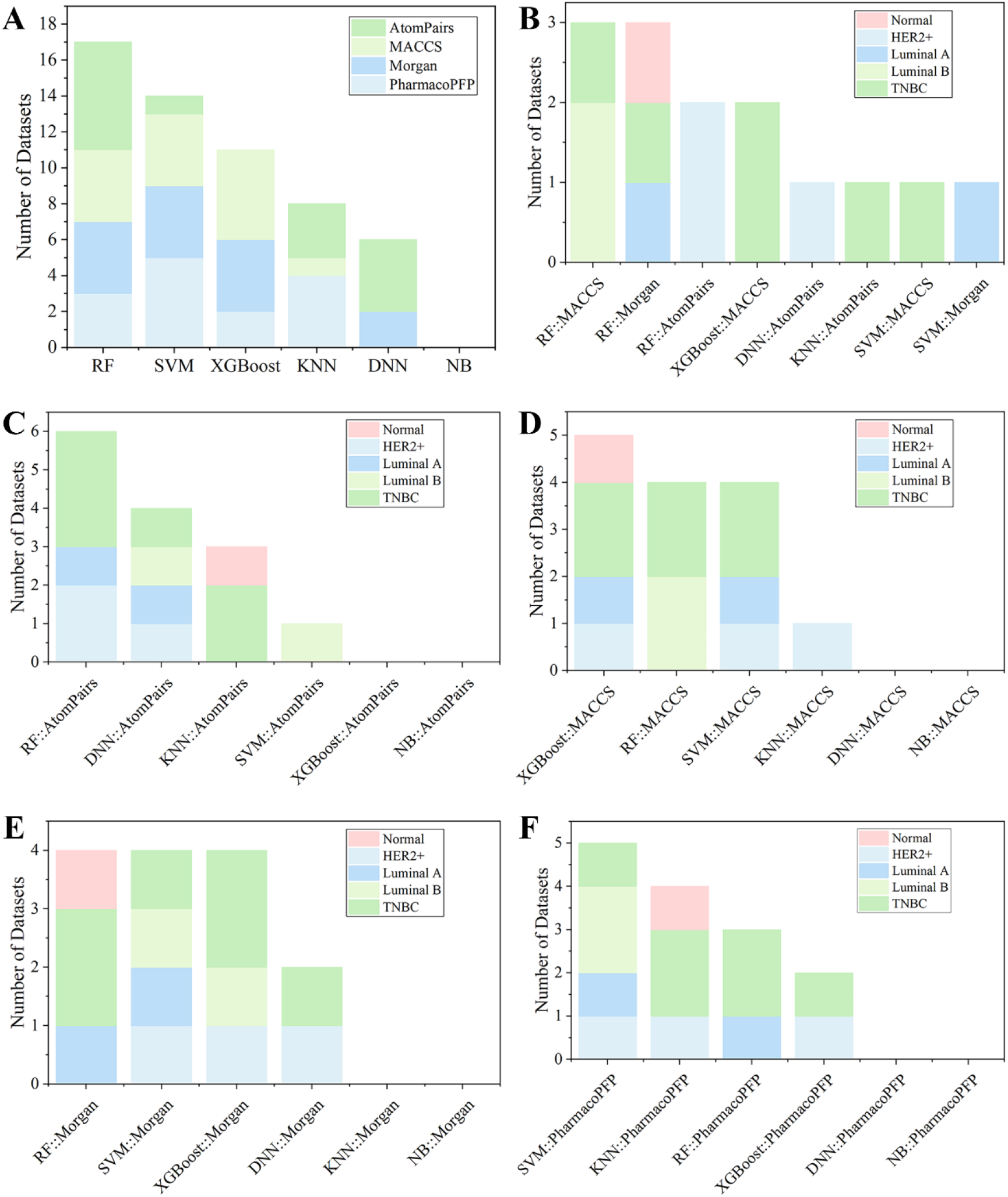
(A) Summary of the optimal models for each fingerprint-based feature. (B) The best models among various fingerprint-based models for different kinds of breast cell lines. The optimal models based on (C) AtomPairs, (D) MACCS, (E) Morgan, and (F) PharmacoPFP for different subtypes of breast cell lines.

**Table 2.** Optimal models in different datasets and the evaluation of test datasets.

| Molecular features | Algorithms | F1 scores<sup>j</sup> | BA<sup>k</sup> | AUC<sup>l</sup> |
| --- | --- | --- | --- | --- |
| Morgan | DNN<sup>a</sup> | 0.832 ± 0.080 | 0.735 ± 0.058 | 0.822 ± 0.078 |
|  | KNN<sup>b</sup> | 0.836 ± 0.084 | 0.771 ± 0.063 | 0.821 ± 0.069 |
|  | NB<sup>c</sup> | 0.775 ± 0.094 | 0.720 ± 0.079 | 0.782 ± 0.078 |
|  | RF<sup>d</sup> | 0.846 ± 0.087 | 0.754 ± 0.068 | 0.852 ± 0.072 |
|  | SVM<sup>e</sup> | 0.843 ± 0.084 | 0.747 ± 0.067 | 0.838 ± 0.072 |
|  | XGBoost<sup>f</sup> | 0.832 ± 0.076 | 0.728 ± 0.062 | 0.813 ± 0.079 |
|  | **Mean** | **0.827 ± 0.026** | **0.743 ± 0.019** | **0.821 ± 0.024** |
| MACCS | DNN | 0.831 ± 0.076 | 0.737 ± 0.060 | 0.822 ± 0.067 |
|  | KNN | 0.846 ± 0.050 | 0.759 ± 0.056 | 0.798 ± 0.067 |
|  | NB | 0.723 ± 0.077 | 0.637 ± 0.073 | 0.722 ± 0.103 |
|  | RF | 0.853 ± 0.066 | 0.761 ± 0.064 | 0.860 ± 0.067 |
|  | SVM | 0.851 ± 0.064 | 0.755 ± 0.059 | 0.830 ± 0.068 |
|  | XGBoost | 0.842 ± 0.074 | 0.760 ± 0.056 | 0.842 ± 0.068 |
|  | **Mean** | **0.824 ± 0.050** | **0.735 ± 0.049** | **0.812 ± 0.049** |
| AtomPairs | DNN | 0.853 ± 0.050 | 0.759 ± 0.057 | 0.842 ± 0.063 |
|  | KNN | 0.851 ± 0.037 | 0.781 ± 0.051 | 0.828 ± 0.064 |
|  | NB | 0.678 ± 0.099 | 0.668 ± 0.083 | 0.732 ± 0.085 |
| | RF | $0.851 \pm 0.066$ | $0.753 \pm 0.054$ | $0.858 \pm 0.059$ |
| | SVM | $0.847 \pm 0.062$ | $0.737 \pm 0.069$ | $0.829 \pm 0.066$ |
| | XGBoost | $0.840 \pm 0.074$ | $0.755 \pm 0.041$ | $0.837 \pm 0.075$ |
|  | **Mean** | **$0.820 \pm 0.070$** | **$0.742 \pm 0.039$** | **$0.821 \pm 0.045$** |
| Molecular Graph | Attentive FP | $0.831 \pm 0.070$ | $0.721 \pm 0.086$ | $0.809 \pm 0.087$ |
| | GAT<sup>g</sup> | $0.810 \pm 0.086$ | $0.695 \pm 0.088$ | $0.774 \pm 0.075$ |
| | GCN<sup>h</sup> | $0.818 \pm 0.076$ | $0.710 \pm 0.091$ | $0.798 \pm 0.100$ |
| | MPNN<sup>i</sup> | $0.821 \pm 0.080$ | $0.696 \pm 0.109$ | $0.781 \pm 0.090$ |
|  | **Mean** | **$0.820 \pm 0.009$** | **$0.708 \pm 0.011$** | **$0.793 \pm 0.015$** |
| PharmacPFP | DNN | $0.824 \pm 0.072$ | $0.705 \pm 0.091$ | $0.803 \pm 0.105$ |
| | KNN | $0.840 \pm 0.060$ | $0.755 \pm 0.075$ | $0.782 \pm 0.070$ |
| | NB | $0.705 \pm 0.088$ | $0.619 \pm 0.075$ | $0.680 \pm 0.080$ |
| | RF | $0.840 \pm 0.064$ | $0.731 \pm 0.070$ | $0.840 \pm 0.060$ |
| | SVM | $0.835 \pm 0.068$ | $0.722 \pm 0.064$ | $0.823 \pm 0.059$ |
| | XGBoost | $0.838 \pm 0.049$ | $0.727 \pm 0.072$ | $0.825 \pm 0.058$ |
|  | **Mean** | **$0.814 \pm 0.054$** | **$0.710 \pm 0.047$** | **$0.792 \pm 0.059$** |
| RDKit | DNN | $0.817 \pm 0.063$ | $0.671 \pm 0.089$ | $0.782 \pm 0.070$ |
| | KNN | $0.831 \pm 0.053$ | $0.736 \pm 0.065$ | $0.778 \pm 0.068$ |
| | NB | $0.753 \pm 0.068$ | $0.605 \pm 0.083$ | $0.672 \pm 0.108$ |
| | RF | $0.840 \pm 0.073$ | $0.725 \pm 0.073$ | $0.835 \pm 0.067$ |
| | SVM | $0.805 \pm 0.091$ | $0.656 \pm 0.086$ | $0.761 \pm 0.077$ |
| | XGBoost | $0.836 \pm 0.084$ | $0.740 \pm 0.071$ | $0.839 \pm 0.060$ |
|  | **Mean** | **$0.814 \pm 0.032$** | **$0.689 \pm 0.054$** | **$0.778 \pm 0.061$** |

<sup>a</sup> DNN: Deep neural networks. <sup>b</sup> KNN: K-Nearest Neighbor. <sup>c</sup> NB: Naïve Bayesian. <sup>d</sup> RF: Random forest. <sup>e</sup> SVM: Support vector machine. <sup>f</sup> XGBoost: Extreme gradient boosting. <sup>g</sup> GCN: Graph convolutional networks. <sup>h</sup> GAT: Graph attention network; <sup>i</sup> MPNN: Message passing neural networks. <sup>j</sup> F1 scores: F1-measure. <sup>k</sup> BA: Balanced accuracy. <sup>l</sup> AUC: Area under the receiver operating characteristics curve.

### 3.4 Performance of graph-based prediction models for breast-associated cells

Compared with the traditional pre-tailored molecular descriptors and/or fingerprints, the key feature of GNN is its capacity to automatically learn task-specific molecular representations using graph convolutions. The SOAT accuracies of GNN models and their variants (e.g., GCN, MPNN, GAT, and Attentive FP) have been reported in various molecular property prediction tasks (Wu, et al., 2017; Xiong, et al., 2020; Yang, et al., 2019). Therefore, 56 molecular graph-based models were established using four types of DL algorithms, including GCN, MPNN, GAT, and Attentive FP. The detailed performance results of molecular graph-based models are listed in Table S7. As shown in Fig. S6, the Attentive FP models exhibited the overall best performance compared with other GNN methods, with a relatively higher average F1 score (0.831 ± 0.070) and AUC (0.809 ± 0.086). The BA results are shown in Fig. S7. Fig. S6C shows that the Attentive FP models performed the best in six breast cancer cell lines including Bcap37, MCF-7, MDA-MB-453, MDA-MB-468, SK-BR-3, and T-47D, making it the most frequent choice. The GCN models showed the best performance in four breast cell lines (BT-549, HBL-100, MDA-MB-231, and MDA-MB-361), the MPNN models performed the best in BT-20 and BT-474 cell lines, and the GAT models performed the best in HS-578T and MDA-MB-435 cell lines.

One advantage of the DL model is its capacity for multi-task model building for attribute-related datasets to improve the accuracy of the single-task model (Li, et al., 2018). Therefore, the multi-task models were trained by the entire 13 breast cancer cell-compound datasets based on the features of the Morgan fingerprints using DNN and molecular graphs using GCN, Attentive FP. Table S8 shows that the AUC of the multi-task models was not better than that of the single-task models. Further data point distribution analysis found that the number of common compounds shared by 13 cell line datasets was small (only 12 molecules, Fig. S8), which explains the poor performance results (Table S8) of the multi-task models.

### 3.5 The optimal model for each breast cell line and further validation

Comparison of the established molecular descriptor-, fingerprint-, and graph-based models showed that (1) the RF algorithm had a better performance capability than the other five ML methods, with higher average metric values of F1 score, BA, and AUC (Table 2) in both descriptor- and fingerprint-based models, while XGBoost also achieved comparable results for these 14 modeling datasets (Table 2 and Fig. 3A); (2) among the established 56 graph-based models, Attentive FP architecture outperformed the other three deep graph learning approaches (i.e., GCN, MPNN, and GAT) on average across all 14 datasets (Table 2); and (3) the performance of molecular fingerprint-based models is generally better than that of both descriptor- and graph-based models at least in these 14 datasets (Table 2), implying that graph DL methods do not achieve better results than the traditional ML learning methods (especially for the two most efficient algorithms, XGBoost and RF), which is consistent with a recent systematic comparison study (Jiang, et al., 2021).

According to the metrics of F1 score, BA, and AUC from the test sets, the optimal *in silico* predictive model for each breast cell line is listed in Table S9. Fingerprint-based RF models performed the best because they ranked first in eight of 14 cell lines. Fingerprint-based XGBoost and SVM models are tied for second place and performed best in two of 14 breast cell lines each. For example, the RF::Morgan model achieved higher prediction results against MDA-MB-231 and T-47D breast cancer cell lines, with ACC values of 83.7% and 84.0%, respectively, and AUC values of 0.904 and 0.885, respectively. The lack of selectivity for cancer cells rather than normal cells is one of the main factors that limit the development of anticancer drugs for clinical use (Dy and Adjei, 2013; Guo, et al., 2020). For one normal breast cell line (HBL-100), the RF::Morgan model also showed good prediction results, with ACC and AUC values of 83.9%, and 0.823, respectively, suggesting that this model can be used to detect whether a given molecule selectively inhibits breast cancer cells over normal human breast cells.

Model fusion may improve the classification prediction performance of a single model by combining the classification prediction results from the corresponding multiple models. Both voting and stacking methods were used in this study for model fusion. As shown in Table 2, Morgan fingerprint-based models performed the best in different kinds of fingerprint-based models with an average F1 score of 0.827 ± 0.026, and RF, XGBoost, and SVM algorithms performed best in most of the datasets (Fig. 3A and E). Therefore, RF, SVM, and XGBoost models for model fusion were applied based on Morgan fingerprints. A total of 112 fusion models were established, and detailed performance results for these voting and stacking models are listed in Tables S10 and S11. As shown in Fig. S9, the average F1 scores of voting or stacking models was similar in each dataset. In all the datasets of breast cell lines, the RF + XGBoost voting model showed the best average performance among fusion models, with average F1, BA, and AUC of 0.849 ± 0.066, 0.749 ± 0.075, and 0.845 ± 0.075, respectively. The fusion models based on Morgan fingerprints were slightly but not significantly better than the single models.

To validate the stability and reliability of the models presented, 10-fold cross-validation and 10 different random seeds of data were used to retrain the models based on the combination of Morgan fingerprints and two ML algorithms (RF and XGBoost). The performance of 10-fold cross-validation classification models is summarized in Table S12 and Fig. 4. Overall, all RF::Morgan models performed well, showing high F1 scores of 0.582–0.914, AUC values of 0.704–0.960, and ACC values of 0.685–0.878. XGBoost::Morgan models showed a similar trend in the 10-fold cross-validation experiment. In 14 cell line datasets, both RF::Morgan and XGBoost::Morgan models consistently exhibited better performance with different seeds (Fig. S10), and the performance showed comparable or smaller variation compared with the previous models based on a specific random seed. Taken together, these results demonstrate that the models presented in this study show stability and reliability. Y-scrambling testing was used to demonstrate that the results are not attributed to chance correlation. As illustrated in Fig. S11 and S12, the F1 scores, BA, and AUC values of the RF::Morgan and XGBoost::Morgan models were significantly higher than those of any of the Y-scrambled models, which confirmed that the results were not chance correlations.

**Fig. 4.**
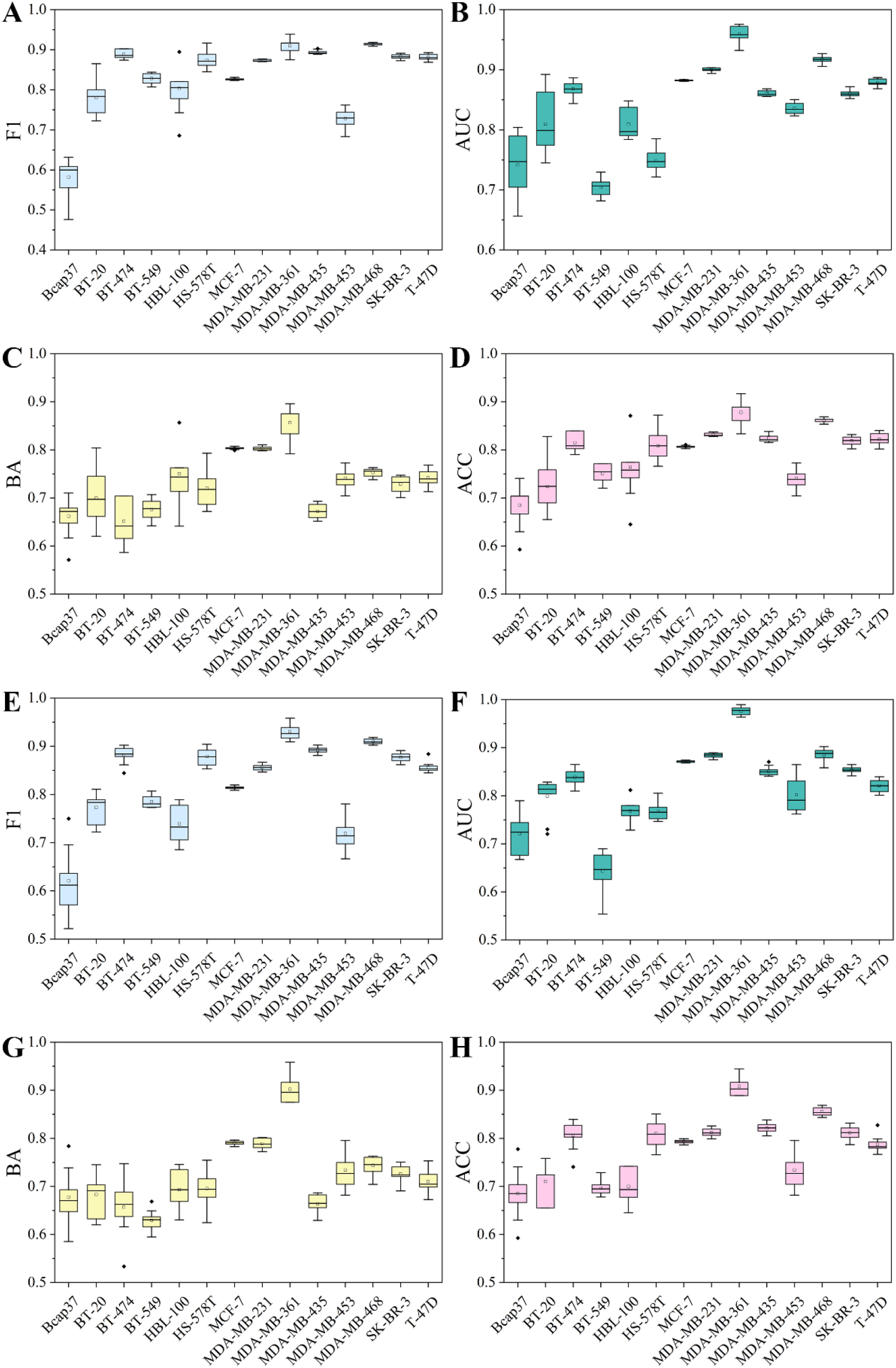
The performance of 10-fold cross-validation results in RF::Morgan and XGBoost::Morgan models. (A-D) F1 scores, AUC, BA, and ACC results in RF::Morgan models. (E-H) F1 scores, AUC, BA, and ACC results in XGBoost::Morgan models.

### 3.6 Interpretation of the optimal model for each breast cell line

To gain a deeper understanding of the established models, we used the SHAP method to calculate the contribution of important structural fragments. Because models based on the combination of the RF and Morgan fingerprints had relatively high predictive performance, we used TreeExplainer, a tree explanation method in SHAP, to calculate the optimal local explanation for these RF::Morgan models. In the MDA-MB-231 cell line as an example, the top 20 favorable and unfavorable structural fragments for MDA-MB-231 inhibition were determined based on the SHAP value and are displayed in Fig. 5 and S33. Morgan 128, Morgan 926, and Morgan 694 (Fig. 5A) had a higher contribution to the RF::Morgan model, indicating a higher probability that the predicted molecules with these fragments will have anti-BC activity. The top 20 important structural fragments for other breast cell lines are shown in Supplementary Fig. S13-S39, which may facilitate anti-BC lead compound selection and optimization.

**Fig. 5.**
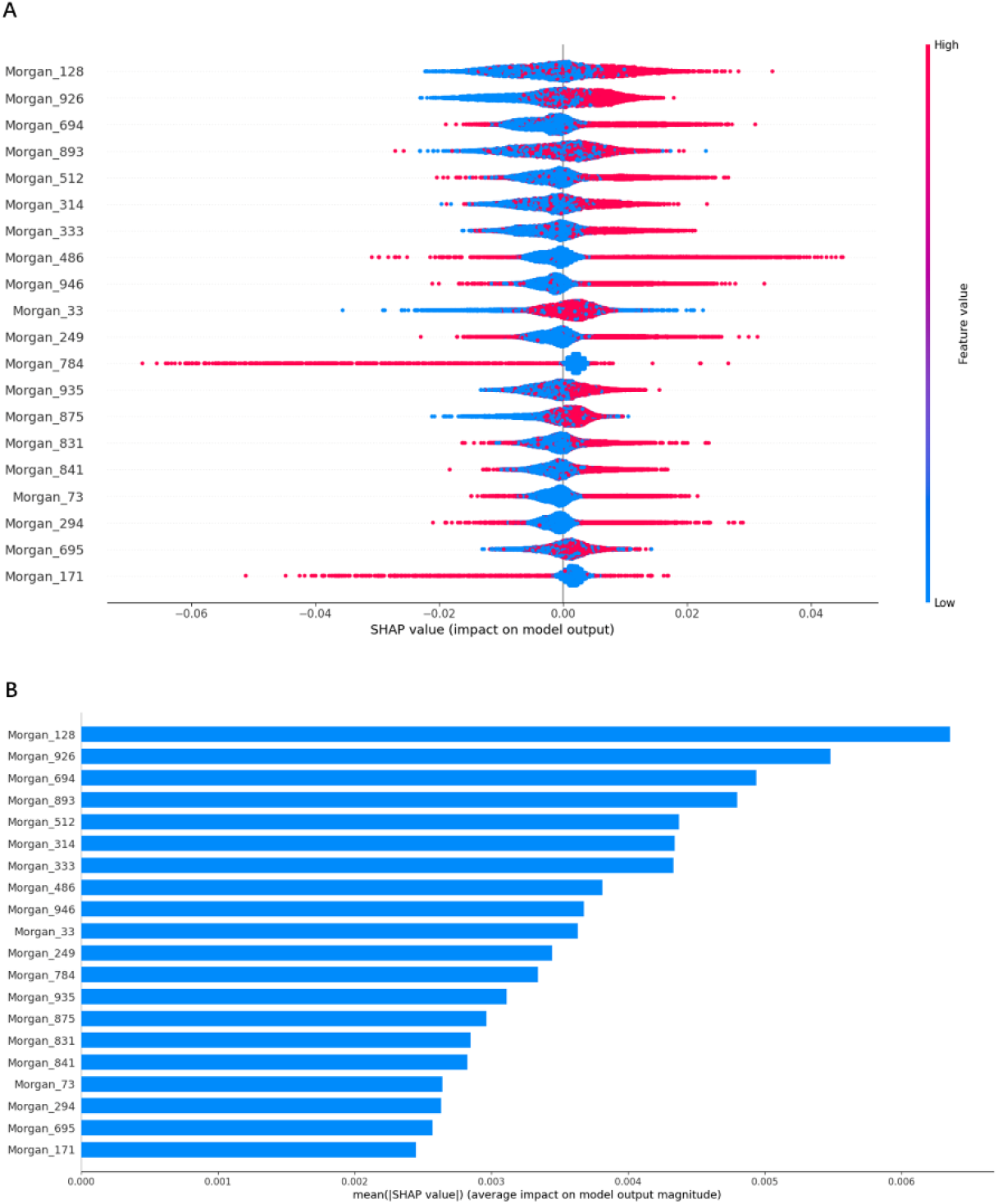
Based on the top 20 most important features of the RF::Morgan model in MDA-MB-231, (A) the SHAP values for each molecular substructure, and (B) the mean of the absolute value of the SHAP value for each molecular substructure.

### 3.7 Model AD

To further evaluate the generalization capability of our models, the LOF algorithm was applied to detect super-applicability domain compounds in the datasets. We first reduced the Morgan fingerprints of 1024 bits to two dimensions by Principal Component Analysis in Scikit-learn and then used the LOF module for calculation. As shown in Fig. S40, there are fewer red points, which indicates that each dataset has fewer super-applicability domain compounds. Therefore, selecting compounds that are similar to those in the datasets of this study may result in higher prediction accuracy when using the present model. The molecular (feature) spaces can be used to define the applicability domain, thus, a simpler way to determine whether a molecule fits the models of this study is to directly calculate the molecular weight of the molecule. Since the molecular weight range of the molecules in this study is 108.10-5714.45, we recommend using molecules in this range for prediction, which can make the prediction more accurate.

### 3.8 Webserver and local version software for the prediction of anti-BC agents

To facilitate the use of these models by experts and non-experts in the field, we built a web-based online forecasting system called ChemBC (http://chembc.idruglab.cn/). To expand the AD threshold of the established model, we retained models for each breast cell line according to the combination of Morgan fingerprint and RF using the entire dataset, and then implemented these retained models into ChemBC and its local version. According to the 10-fold cross-validation (AUC = 0.780-0.928, ACC = 0.714-0.880), the retrained models for 14 breast cell line datasets showed excellent predictive performance. ChemBC was developed based on the Django framework using the Python package. The main functional module of ChemBC is prediction (Fig. 6) in which users can upload and/or online draw a structure to easily and quickly predict the inhibitory activity against 13 breast cancer cell lines and one normal breast cell line. In addition, a local version executable software (https://github.com/idruglab/ChemBC) was developed to perform large-scale VS screening.

**Fig. 6.**
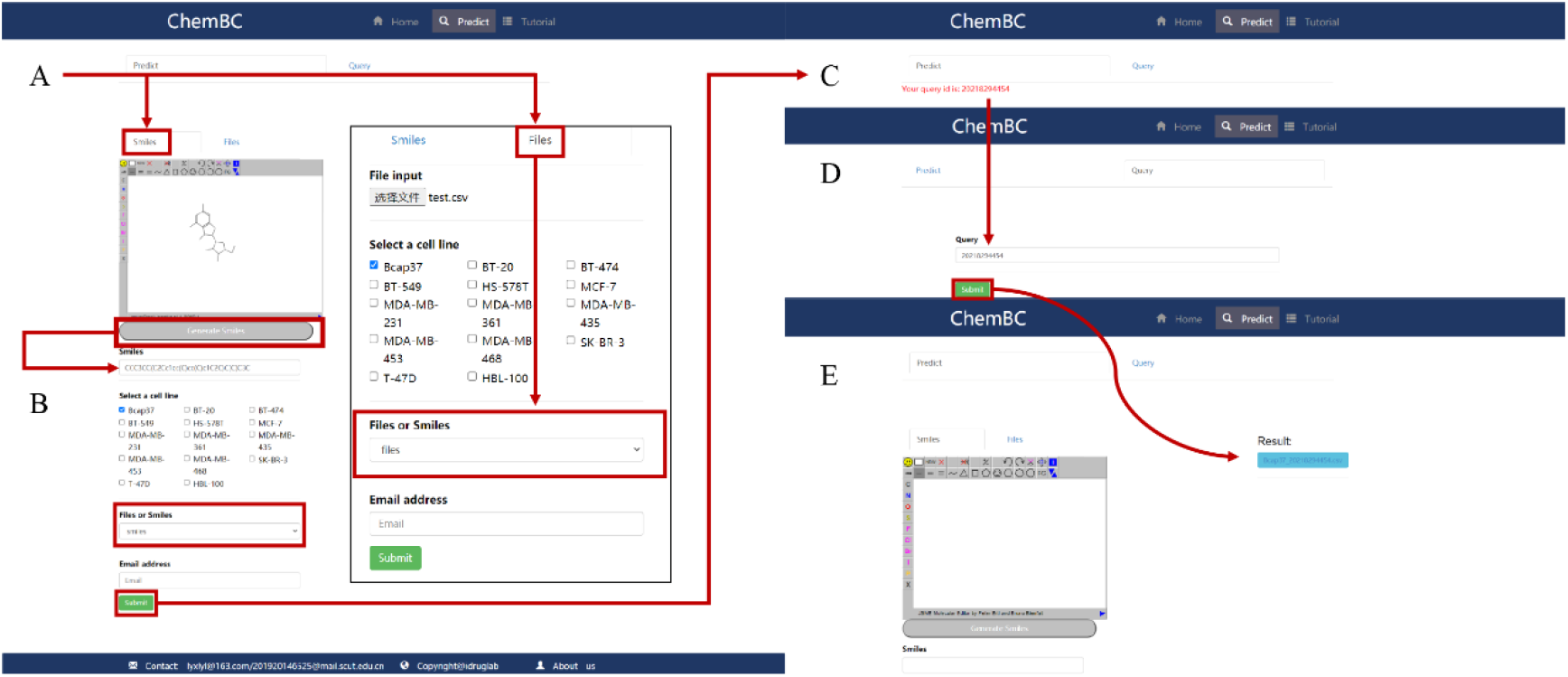
Website schematic diagram of bioactivity prediction. (A) Draw a molecule or upload a file as input. (B) Select a breast cancer cell line for prediction. (C) Get a query id to view the result. (D) Enter the query id. (E) Download the results.

## 4 Conclusions

In this study, we collected datasets of phenotypic compound-cell association bioactivity toward 13 breast cancer cell lines and one normal breast cell line and constructed 588 models based on three molecular representatives, including molecular descriptors, fingerprints, and graphs using five conventional ML and five DL algorithms. Compared with these established models, the performance of RF::Morgan models was superior to that of the other models based on molecular descriptors and graphs. Based on RF::Morgan models, the important favorable and unfavorable fragments for each breast cell line generated using SHAP algorithms will be helpful for lead optimization or the design of new agents with better anti-BC activity. Although some fusion models based on voting and stacking methods showed better performance than single models, the observed improvement was minor. Finally, the online platform ChemBC and its local version software were developed based on well-established models, which could contribute to research aimed at designing and discovering new anti-BC agents.

## Acknowledgements

We acknowledge the use of computational resources from the SCUT supercomputing platform.

## Funding

This work was supported in part by the National Natural Science Foundation of China (Nos. 81973241 and 82060625), the Natural Science Foundation of Guangdong Province (2020A1515010548), and the Guizhou Provincial Natural Science Foundation ([2020]1Z073).

---

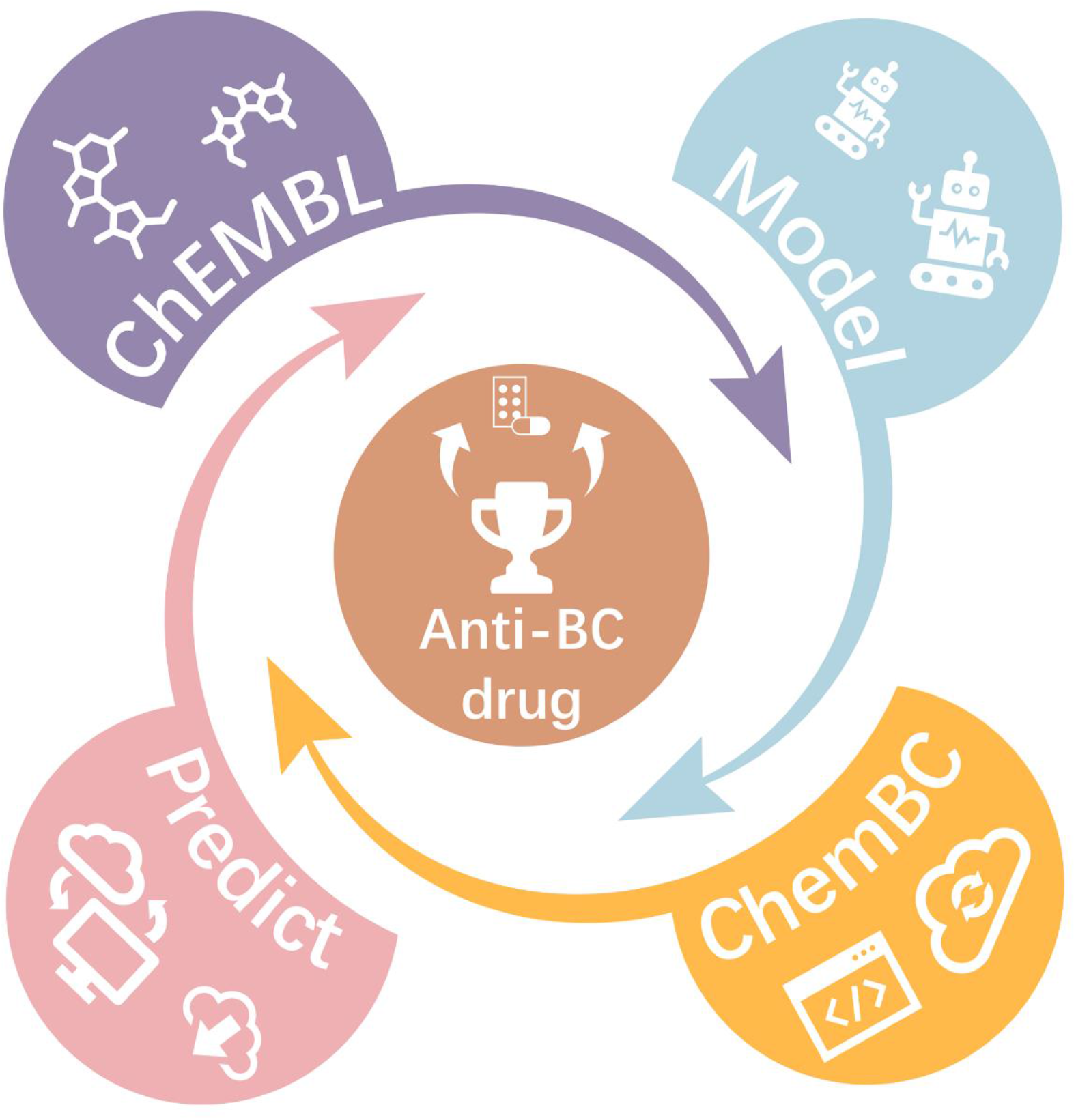

## Notes

http://chembc.idruglab.cn/

https://github.com/idruglab/ChemBC

### Competing Interest Statement

The authors have declared no competing interest.

## References

Ashdown, G.W., et al. (2020) A machine learning approach to define antimalarial drug action from heterogeneous cell-based screens. Sci Adv., 6, eaba9338.

Bemis, G.W. and Murcko, M.A. (1996) The Properties of Known Drugs. 1. Molecular Frameworks. J Med Chem., 39, 2887–2893.

Breunig, M.M., et al. (2000) LOF: identifying density-based local outliers. In Proceedings of the 2000 ACM SIGMOD international conference on Management of data, pp. 93–104.

Brower, V. (2013) Cardiotoxicity debated for anthracyclines and trastuzumab in breast cancer. J Natl Cancer Inst., 105, 835–836.

Buckner, F.S., et al. (2020) Phenotypic drug discovery for human African trypanosomiasis: A powerful approach. Trop Med Infect Dis., 5, 23.

Cameron, D., et al. (2017) 11 years’ follow-up of trastuzumab after adjuvant chemotherapy in HER2-positive early breast cancer: final analysis of the HERceptin Adjuvant (HERA) trial. Lancet., 389, 1195–1205.

Carhart, R.E., et al. (1985) Atom pairs as molecular features in structure-activity studies: definition and applications. J. Chem. Inf. Comput. Sci., 25, 64–73.

Chandrasekaran, S.N., et al. (2021) Image-based profiling for drug discovery: due for a machine-learning upgrade? Nat Rev Drug Discov. 20, 145–159.

Chen, T. and Guestrin, C. (2016) Xgboost: A scalable tree boosting system. In, Proceedings of the 22nd acm sigkdd international conference on knowledge discovery and data mining. pp. 785–794.

Daniyal, A., et al. (2021) Genetic Influences in Breast Cancer Drug Resistance. Breast Cancer (Dove Med Press) 13, 59–85.

Durant, J.L., et al. (2002) Reoptimization of MDL keys for use in drug discovery. J Chem Inf Comput Sci., 42, 1273–1280.

Escala-Garcia, M., et al. (2020) Breast cancer risk factors and their effects on survival: a Mendelian randomisation study. BMC Med., 18, 1–10.

Fields, F.R., et al. (2020) Novel antimicrobial peptide discovery using machine learning and biophysical selection of minimal bacteriocin domains. Drug Dev Res., 81, 43–51.

Gobbi, A., Poppinger, D. (1998) Genetic optimization of combinatorial libraries. Biotechnol Bioeng. 61, 47–54.

Guo, Q., et al. (2019) Discovery, biological evaluation, structure–activity relationships and mechanism of action of pyrazolo [3, 4-b] pyridin-6-one derivatives as a new class of anticancer agents. Org Biomol Chem. 17, 6201–6214.

Harbeck, N., Thomssen, C. and Gnant, M. (2013) St. Gallen 2013: brief preliminary summary of the consensus discussion. Breast Care (Basel), 8, 102–109.

Jiang, D., et al. (2021) Could graph neural networks learn better molecular representation for drug discovery? A comparison study of descriptor-based and graph-based models. J Cheminform, 13, 12.

Kc, G.B., et al. (2021) A machine learning platform to estimate anti-SARS-CoV-2 activities. Nature Machine Intelligence, 3, 527–535.

Li, X., et al. (2018) Prediction of Human Cytochrome P450 Inhibition Using a Multitask Deep Autoencoder Neural Network. Mol Pharm., 15, 4336–4345.

Li, Y. and Li, Z. (2021) Potential Mechanism Underlying the Role of Mitochondria in Breast Cancer Drug Resistance and Its Related Treatment Prospects. Front Oncol., 11, 629614.

Liao, M., et al. (2021) Small-molecule drug discovery in triple negative breast cancer: current situation and future directions. J Med Chem., 64, 2382–2418.

Lundberg, S. and Lee, SI. (2017) A unified approach to interpreting model predictions. arXiv, Preprint arXiv:1705.07874

Lundberg, S.M., et al. (2020) From local explanations to global understanding with explainable AI for trees. Nat Mach Intell., 2, 56–67.

Luo, Y., et al. (2019) Identifying a novel anticancer agent with microtubule-stabilizing effects through computational cell-based bioactivity prediction models and bioassays. Org Biomol Chem., 17, 1519–1530.

Malandraki-Miller S, Riley PR. (2021) Use of artificial intelligence to enhance phenotypic drug discovery. Drug Discov Today., 26, 887–901.

Mendez, D., et al. (2019) ChEMBL: towards direct deposition of bioassay data. Nucleic Acids Res., 47, D930–D940.

Pedregosa, F., et al. (2011) Scikit-learn: Machine learning in Python. J. Mach. Learn. Res. 12, 2825–2830.

Rogers, D., Hahn, M. (2010) Extended-connectivity fingerprints. J Chem Inf Model., 50, 742–754.

Shah, A.N. and Gradishar, W.J. (2018) Adjuvant Anthracyclines in Breast Cancer: What Is Their Role? Oncologist, 23, 1153–1161.

Stokes, J.M., et al. (2020) A deep learning approach to antibiotic discovery. Cell, 180, 688–702.

Sung, H., et al. (2021) Global cancer statistics 2020: GLOBOCAN estimates of incidence and mortality worldwide for 36 cancers in 185 countries. CA Cancer J Clin., 71, 209–249.

Wang, L., et al. (2014) Discovering new agents active against methicillin-resistant Staphylococcus aureus with ligand-based approaches. J Chem Inf Model., 54, 3186–3197.

Wu, Z., et al. (2017) MoleculeNet: a benchmark for molecular machine learning. Chem Sci, 9, 513–530.

Xiong, Z., et al. (2020) Pushing the Boundaries of Molecular Representation for Drug Discovery with the Graph Attention Mechanism. J Med Chem., 63, 8749–8760.

Yang, K., et al. (2019) Analyzing Learned Molecular Representations for Property Prediction. J Chem Inf Model., 59, 3370–3388.

Ye, Q., et al. (2021) Identification of active molecules against Mycobacterium tuberculosis through machine learning. Brief Bioinform., bbab068.

Zheng, J.-X., et al. (2021) Infestation risk of the intermediate snail host of Schistosoma japonicum in the Yangtze River Basin: improved results by spatial reassessment and a random forest approach. Infect Dis Poverty., 10, 74.

Zoffmann, S., et al. (2019) Machine learning-powered antibiotics phenotypic drug discovery. Sci Rep., 9, 1–14.

