## Supplemental data for "Machine learning enables accurate and rapid prediction of active molecules against breast cancer cells"

**Contents**

### Part I: Machine learning algorithms

#### Random forest (RF)

RF is a representative ensemble learning approach. It establishes a classifier or regressor by an ensemble of individual decision trees and makes predictions as final output by vote or by averaging multiple decision trees (Svetnik V, 2003). Compared with a decision tree, RF has high prediction accuracy, good tolerance to outliers and noise, and is not easy to overfit. To obtain the best RF model, the following five hyperparameters were optimized: n_estimators (10–500), criterion (‘*gini’* and ‘*entropy’*), max_depth (0–15), min_samples_leaf (1–10), and max_features (‘*log2’*, ‘*auto’* and ‘*sqrt’*).

**Support vector machine (SVM)**

SVM is a supervised ML algorithm that can be used for both classification and regression tasks (Zernov, et al., 2003). The basic idea underlying SVM is to find the optimal hyperplane in the feature space that can be obtained by maximizing the boundary between classes in N-dimensional space, which distinguishes objects with different class labels. SVM has been widely used in drug discovery-relevant applications such as compound activity and property prediction (Heikamp and Bajorath, 2014). In the training of SVM models, two hyperparameters, Kernel coefficient (gamma, ‘auto’, 0.1–0.2) and penalty parameter C of the error term (C, from 1 to 100), were optimized.

**Extreme gradient boosting (XGBoost)**

XGBoost is one of the so-called ensemble learning algorithms under the Gradient Boosting framework and has achieved state-of-the-art ranking results in many ML competitions. It has been widely used in molecular property/activity prediction tasks (Jiang, et al., 2021; Li, et al., 2021; Ye, et al., 2021). Seven hyperparameters were optimized in the training of XGBoost models: learning_rate (0.01–0.1), gamma (0–0.1), min_child_weight (1–3), max_depth (3–5), n_estimators (50–100), subsample (0.8–1.0), and colsample bytree (0.8–1.0).

**K-Nearest Neighbor (KNN)**

The basic idea of the KNN ML algorithm (Cover and Hart, 1967) is to identify the *k* training samples closest to the test samples in the training set based on distance measures (e.g., Euclidean, Manhattan, and Jaccard distance), and to make a prediction based on the information of the *k* samples. The default distance measure Euclidean was used in this study. The following three hyperparameters were optimized: n_neighbors (1–5), p (1–2), and weight function (‘uniform’, ‘distance’).

**Naïve Bayesian (NB)**

NB is a classic classification ML method based on Bayes’ theorem (Duda and Hart, 1973) and independent assumption of characteristic conditions. For a given dataset, the joint probability distribution of input and output is first learned based on the independent hypothesis of characteristic conditions. NB is also widely used in drug discovery practices (Guo, et al., 2020; Wang, et al., 2014; Wang, et al., 2016; Wang, et al., 2016). Two hyperparameters were optimized: alpha (0.01–1) and binarize (0, 0.5, 0.8).

**Deep neural networks (DNN)**

DNN is a typical DL algorithm and is essentially an artificial neural network (McCulloch and Pitts, 1943) with multiple hidden layers. It consists of many independent neurons, each of which collects information from its connected neurons, and the aggregated information is then activated through a nonlinear activation function. The following key hyperparameters were optimized: dropouts (0.1, 0.2, 0.5), layer_sizes (64, 128, 256, 512) and weight_decay_penalty (0.01, 0.001, 0.0001).

**Graph convolutional network (GCN)**

GCN is a classic neural network that can use graph-structured data as input (Kipf and Welling, 2016). It is composed of graph convolution layers, a readout layer, fully connected layers, and an output layer. The core idea of graph convolution is to use edge information for aggregating node information, thereby generating a new node representation. Various GCN frameworks have been proposed. Duvenaud et al. (Duvenaud, et al., 2015) introduced a convolutional neural network that allows end-to-end learning of prediction pipelines. In this study, we used Duvenaud’s GCN method, and the following hyperparameters were optimized: weight_decay (0, 10e-8, 10e-6, 10e-4), graph_conv_layers ([64, 64], [128, 128], [256, 256]), learning rate (0.01, 0.001, 0.0001) and dense_layer_size (64, 128, 256).

**Graph attention network (GAT)**

Attention mechanism (AM) is one component of a neural network architecture, which can be embedded in the DL models to automatically learn and calculate the contribution of input data to output data. GCN cannot complete the inductive task, namely, dynamic graph problems, and it is not easy for GCN to assign different learning weights to different neighbors. GAT (Veličković, et al., 2017) introduces an AM to address the disadvantages of previous approaches based on GCN or its approximation. The weight of the features of adjacent nodes depends entirely on the features of the nodes and is independent of the graph structure. In the training of the GAT model, the following hyperparameters were optimized: weight_decay (0, 10e-8, 10e-6, 10e-4), learning rate (0.01, 0.001, 0.0001), n_attention_heads (8, 16, 32), and dropouts (0, 0.1, 0.3, 0.5).

**Message passing neural network (MPNN)**

MPNN, proposed by Gilmer et al. in 2017 (Gilmer, et al., 2017), is a common graph neural network (GNN) framework for chemical prediction tasks. It can directly learn the molecular characteristics from the molecular diagram and is not affected by the graph isomorphism. In the training of the MPNN model, six hyperparameters were optimized: weight_decay (10e-8, 10e-6, 10e-4), learning rate (0.01, 0.001, 0.0001), graph_conv_layers ([64, 64], [128, 128], [256, 256]), num_layer_set2set (2, 3, 4), node_out_feats (16, 32, 64), and edge_hidden_feats (16, 32, 64).

**Attentive FP**

Attentive FP, which was proposed by Xiong et al. (Xiong, et al., 2019), is currently a state-of-the-art GNN model for molecular property prediction, and what is learned from the established model is interpretable. It allows the model to focus on the most relevant parts of the input by applying a graph AM. Herein, the main hyperparameters were optimized as follows: dropout (0, 0.1, 0.5), graph_feat_size (50, 100, 200), num_timesteps (1, 2, 3), num_layers (2, 3, 4), learning rate (0.0001, 0.001, 0.01), and weight_decay (0, 0.01, 0.0001).

### References


Cover, T. and Hart, P. (1967) Nearest neighbor pattern classification. *IEEE Transactions on Information Theory.* vol. 13, pp. 21-27.

Duda, R.O. and Hart, P.E. (1973) Pattern classification and scene analysis. Wiley New York.

Duvenaud, D*., et al.* (2015) Convolutional networks on graphs for learning molecular fingerprints. arXiv, Preprint arXiv: 1509.09292.

.Gilmer, J., *et al.* (2017) Neural message passing for quantum chemistry. *In Proceedings of the 34th International Conference on Machine Learning*. vol. 70, pp. 1263-1272.

Guo, Q., *et al.* (2020) Ligand- and structural-based discovery of potential small molecules that target the colchicine site of tubulin for cancer treatment. *European Journal of Medicinal Chemistry*, **196**, 112328.

Jiang, Z., *et al.* (2021) A comprehensive comparative assessment of 3D molecular similarity tools in ligand-based virtual screening. *Brief Bioinform*. bbab231.

Kipf, T.N. and Welling, M. (2016) Semi-supervised classification with graph convolutional networks. arXiv, Preprint arXiv: 1609.02907.

Li, S., *et al.* (2021) HDAC3i-Finder: A Machine Learning-based Computational Tool to Screen for HDAC3 Inhibitors. **40**, e2000105.

McCulloch WS, Pitts W. (1943) A logical calculus of the ideas immanent in nervous activity. *Bull Math Biol.*, **5**, 115-133.

Svetnik V., *et al.* (2003) A Classification and Regression Tool for Compound Classification and QSAR Modeling. *J Chem Inf Comput Sci.* **43**, 1947-1958.

Veličković, P., *et al.* (2017) Graph attention networks. arXiv, Preprint arXiv: 1710.10903

Wang, L., *et al.* (2014) Predicting mTOR inhibitors with a classifier using recursive partitioning and Naive Bayesian approaches. *PLoS One.*, **9**, e95221.

Wang, L., *et al.* (2016) Discovering new mTOR inhibitors for cancer treatment through virtual screening methods and in vitro assays. *Sci Rep.*, **6**, 1-13.

Wang, L., *et al.* (2016) Chemical fragment-based CDK4/6 inhibitors prediction and web server. *RSC Adv.,* **6**, 16972-16981.

Xiong, Z., *et al.* (2019) Pushing the Boundaries of Molecular Representation for Drug Discovery with the Graph Attention Mechanism. *J Med Chem.,* **63**, 8749-8760.

Ye, Q., *et al.* (2021) Identification of active molecules against Mycobacterium tuberculosis through machine learning. *Brief Bioinform.*, bbab068.

Zernov, V.V., *et al.* (2003) Drug discovery using support vector machines. The case studies of drug-likeness, agrochemical-likeness, and enzyme inhibition predictions. *J Chem Inf Comput Sci.*, **43**, 2048-2056.

### Part II: Supplementary figures and tables

**Fig. S1.** Analysis of 14 datasets. (A) Number of compounds in all datasets. (B) Percentage of active and inactive compounds in all breast cell lines.

**Fig. S2.** The chemical space of the compounds in (A) Bcap37, (B) BT-20, (C) BT-474, (D) BT-549, (E) HS-578T, (F) MCF-7, (G) MDA-MB-231, (H) MDA-MB-361, (I) MDA-MB-435, (J) MDA-MB-453, (K) MDA-MB-468, (L) SK-BR-3, (M) T-47D, (N) HBL-100 datasets.

**Fig. S3.** Performance of descriptor-based BC prediction models. (A) F1 scores of descriptor-based models. (B) AUC results of descriptor-based models. (C) BC results of descriptor-based models.

**Fig. S4.** Performance of fingerprint-based BC prediction models. (A) AUC results of the AtomPairs-based models. (B) AUC results of the MACCS-based models. (C) AUC results of the Morgan-based models. (D) AUC results of the PharmacoPFP-based models.

**Fig. S5.** Performance of fingerprint-based BC prediction models. (A) BA results of the AtomPairs-based models. (B) BA results of the MACCS-based models. (C) BA results of the Morgan-based models. (D) BA results of the PharmacoPFP-based models.

**Fig. S6.** Performance of graph-based BC prediction models. (A) F1 scores of graph-based models. (B) AUC results of graph-based models. (C) The optimal models based on molecular graph for different subtypes of breast cell lines.

**Fig. S7.** BA results of graph-based BC prediction models.

**Fig. S8.** Common molecules in 13 breast cancer cell lines.

**Fig. S9.** F1 scores of fusion models.

**Fig. S10.** The performance of 10 random seeds in RF and XGBoost::Morgan models. (A-D) F1scores, AUC, BA, and ACC results in RF::Morgan models; (E-H) F1scores, AUC, BA, and ACC results in XGBoost::Morgan models.

**Fig. S11.** Y-scrambling results for the RF::Morgan models. Both the training sets and testing sets were unscrambled (gold). The training sets were scrambled, whereas the test sets were unscrambled (green). The training sets were unscrambled, whereas the test sets were scrambled (purple). F1: F1-measure. BA: Balanced accuracy. AUC: Area under the receiver operating characteristics curve.

**Fig. S12.** Y-scrambling results for the XGB::Morgan models. Both the training sets and testing sets were unscrambled (gold). The training sets were scrambled, whereas the test sets were unscrambled (green). The training sets were unscrambled, whereas the test sets were scrambled (purple).F1: F1-measure. BA: Balanced accuracy. AUC: Area under the receiver operating characteristics curve.

**Fig. S13.** Based on the top 20 most important features of the RF::Morgan model in Bcap37, (A) the SHAP values for each molecular substructure, and (B) the mean of the absolute value of the SHAP value for each molecular substructure.

**Fig. S14.** Based on the top 20 most important features of the RF::Morgan model in BT-20, (A) the SHAP values for each molecular substructure, and (B) the mean of the absolute value of the SHAP value for each molecular substructure.

**Fig. S15.** Based on the top 20 most important features of the RF::Morgan model in BT-474, (A) the SHAP values for each molecular substructure, and (B) the mean of the absolute value of the SHAP value for each molecular substructure.

**Fig. S16.** Based on the top 20 most important features of the RF::Morgan model in BT-549, (A) the SHAP values for each molecular substructure, and (B) the mean of the absolute value of the SHAP value for each molecular substructure.

**Fig. S17.** Based on the top 20 most important features of the RF::Morgan model in HBL-100, (A) the SHAP values for each molecular substructure, and (B) the mean of the absolute value of the SHAP value for each molecular substructure.

**Fig. S18.** Based on the top 20 most important features of the RF::Morgan model in HS-578T (A) the SHAP values for each molecular substructure, and (B) the mean of the absolute value of the SHAP value for each molecular substructure.

**Fig. S19.** Based on the top 20 most important features of the RF::Morgan model in MCF-7 (A) the SHAP values for each molecular substructure, and (B) the mean of the absolute value of the SHAP value for each molecular substructure.

**Fig. S20.** Based on the top 20 most important features of the RF::Morgan model in MDA-MB-361 (A) the SHAP values for each molecular substructure, and (B) the mean of the absolute value of the SHAP value for each molecular substructure.

**Fig. S21.** Based on the top 20 most important features of the RF::Morgan model in MDA-MB-435 (A) the SHAP values for each molecular substructure, and (B) the mean of the absolute value of the SHAP value for each molecular substructure.

**Fig. S22.** Based on the top 20 most important features of the RF::Morgan model in MDA-MB-453 (A) the SHAP values for each molecular substructure, and (B) the mean of the absolute value of the SHAP value for each molecular substructure.

**Fig. S23.** Based on the top 20 most important features of the RF::Morgan model in MDA-MB-468 (A) the SHAP values for each molecular substructure, and (B) the mean of the absolute value of the SHAP value for each molecular substructure.

**Fig. S24.** Based on the top 20 most important features of the RF::Morgan model in SK-BR-3 (A) the SHAP values for each molecular substructure, and (B) the mean of the absolute value of the SHAP value for each molecular substructure.

**Fig. S25.** Based on the top 20 most important features of the RF::Morgan model in T-47D (A) the SHAP values for each molecular substructure, and (B) the mean of the absolute value of the SHAP value for each molecular substructure.

**Fig. S26.** Important molecular substructures of the RF::Morgan model in Bcap37.

**Fig. S27.** Important molecular substructures of the RF::Morgan model in BT-20.

**Fig. S28.** Important molecular substructures of the RF::Morgan model in BT-474.

**Fig. S29.** Important molecular substructures of the RF::Morgan model in BT-549.

**Fig. S30.** Important molecular substructures of the RF::Morgan model in HBL-100.

**Fig. S31.** Important molecular substructures of the RF::Morgan model in HS-578T.

**Fig. S32.** Important molecular substructures of the RF::Morgan model in MCF-7.

**Fig. S33.** Important molecular substructures of the RF::Morgan model in MDA-MB-231.

**Fig. S34.** Important molecular substructures of the RF::Morgan model in MDA-MB-361.

**Fig. S35.** Important molecular substructures of the RF::Morgan model in MDA-MB-435.

**Fig. S36.** Important molecular substructures of the RF::Morgan model in MDA-MB-453.

**Fig. S37.** Important molecular substructures of the RF::Morgan model in MDA-MB-468.

**Fig. S38.** Important molecular substructures of the RF::Morgan model in SK-BR-3.

**Fig. S39.** Important molecular substructures of the RF::Morgan model in T-47D.

**Fig. S40.** Model AD in training sets and test sets in all breast cell lines. K was set to 5. By comparing the density of each point p and its five neighborhood points, whether this point is abnormal is judged. The lower the density of point p is, the more likely it is to be identified as an abnormal point. Exceptions are shown in red.

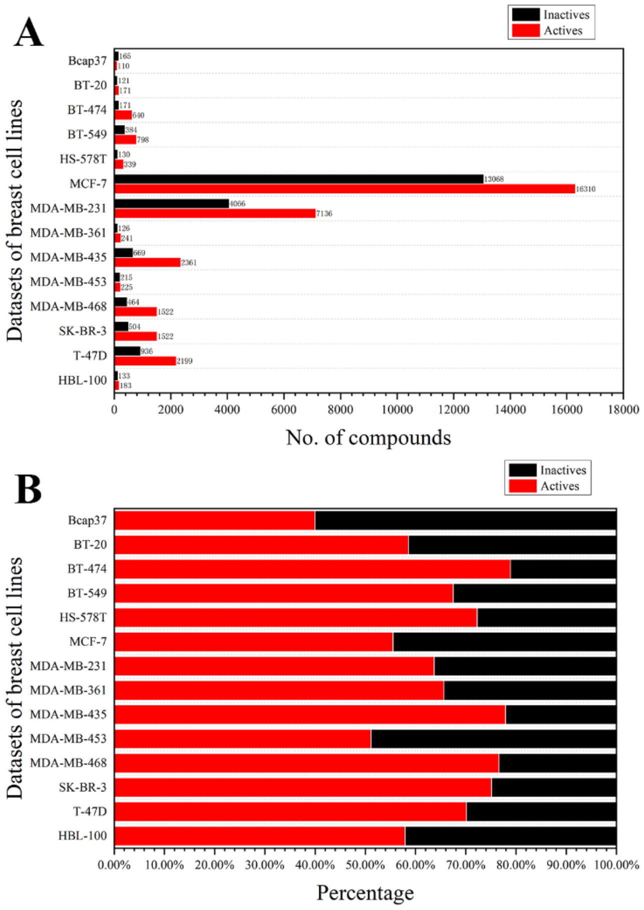


**Fig. S1.** Analysis of 14 datasets. (A) Number of compounds in all datasets. (B) Percentage of active and inactive compounds in all breast cell lines.

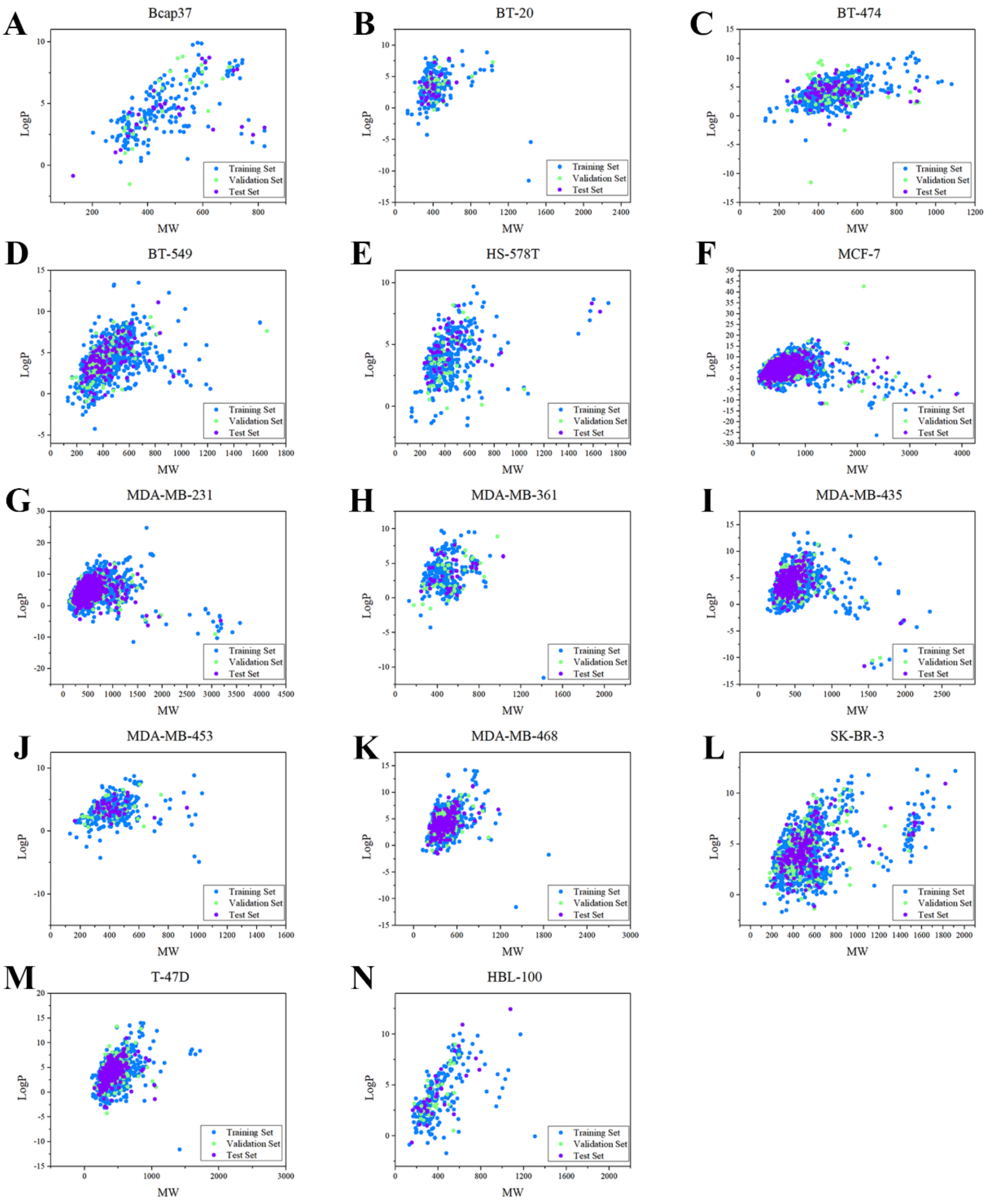


**Fig. S2.** The chemical space of the compounds in (A) Bcap37, (B) BT-20, (C) BT-474, (D) BT-549, (E) HS-578T, (F) MCF-7, (G) MDA-MB-231, (H) MDA-MB-361, (I) MDA-MB-435, (J) MDA-MB-453, (K) MDA-MB-468, (L) SK-BR-3, (M) T-47D, (N) HBL-100 datasets.

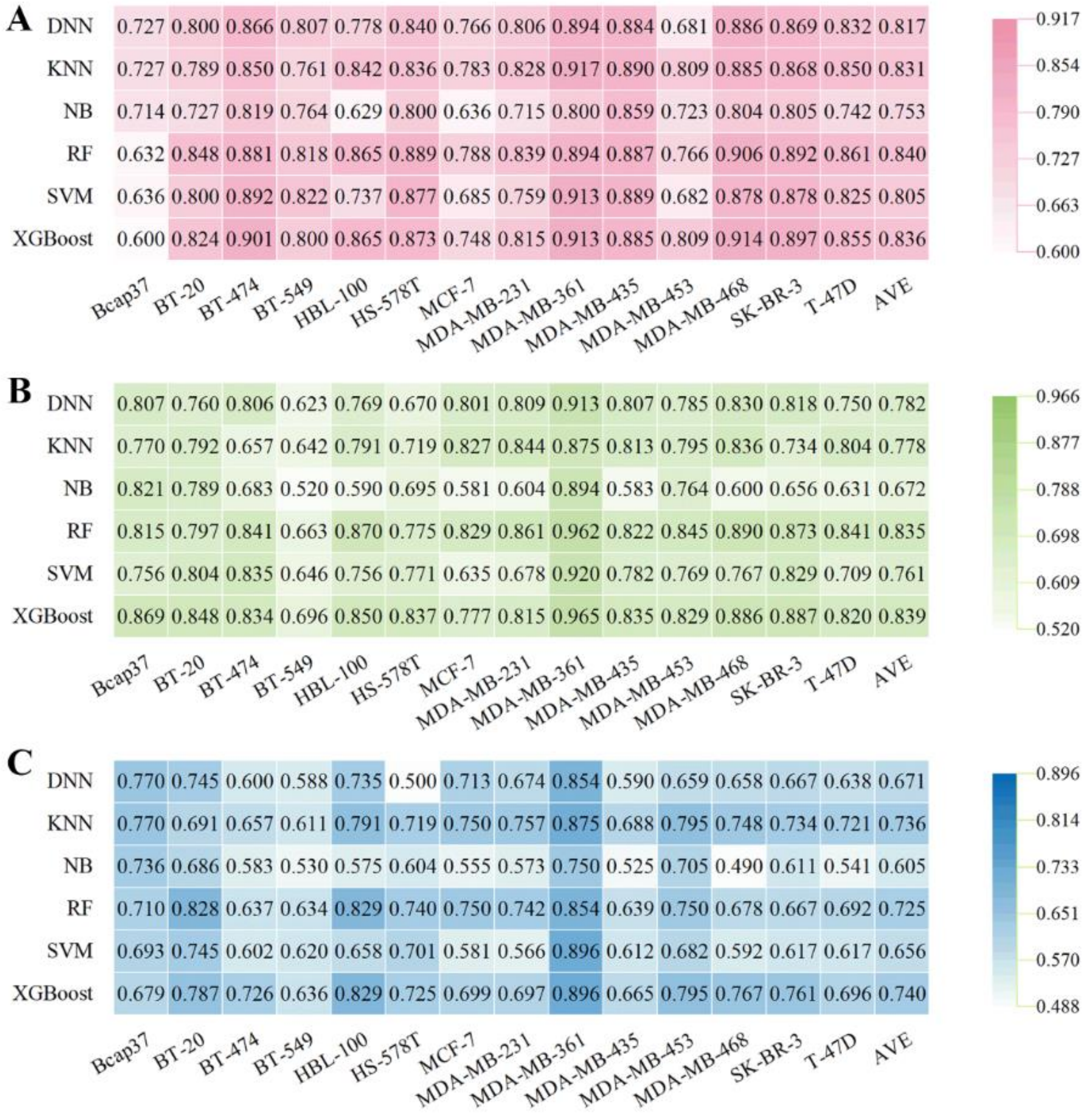


**Fig. S3.** Performance of descriptor-based BC prediction models. (A) F1 scores of descriptor-based models. (B) AUC results of descriptor-based models. (C) BC results of descriptor-based models.

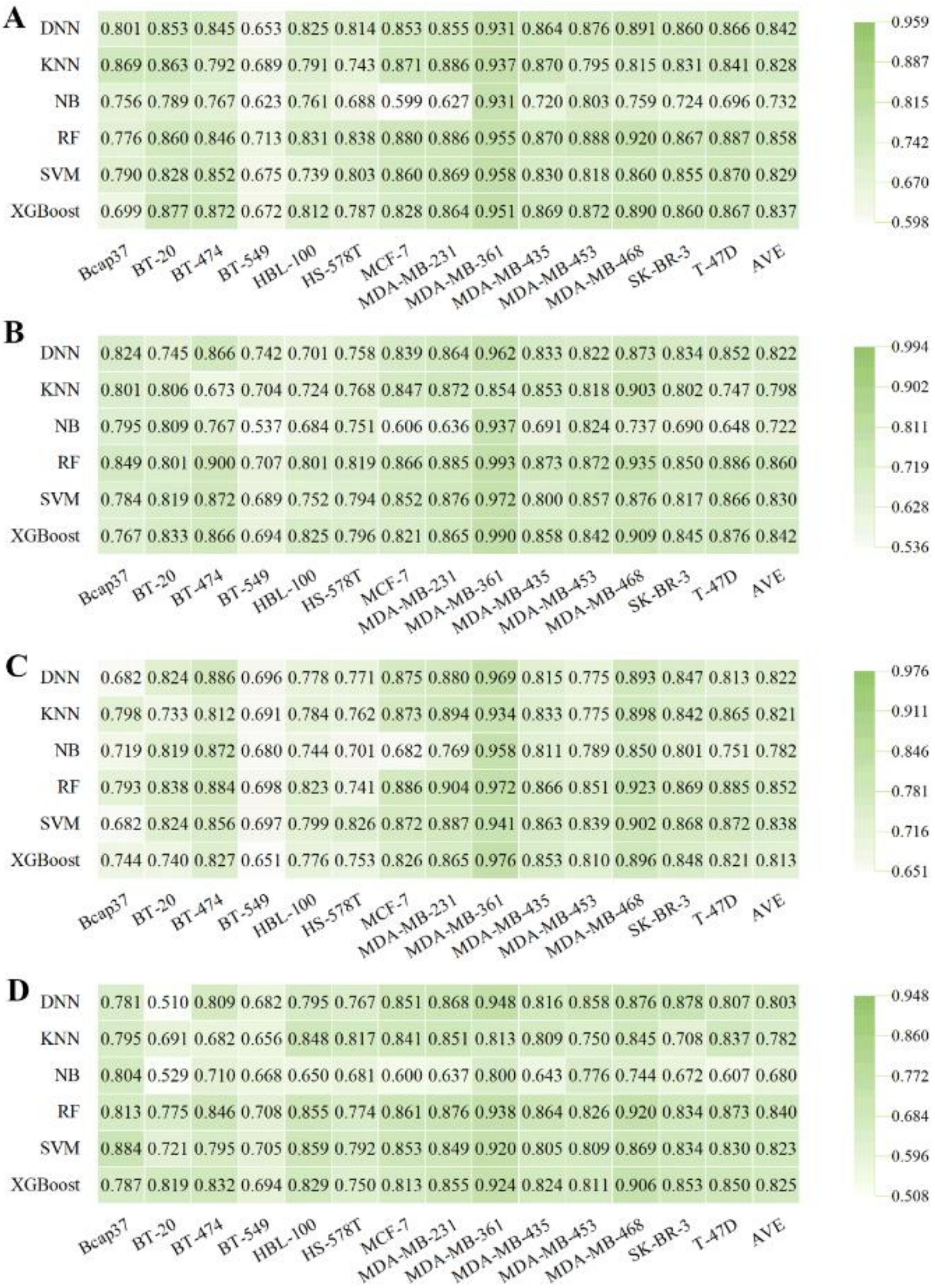


**Fig. S4.** Performance of fingerprint-based BC prediction models. (A) AUC results of the AtomPairs-based models. (B) AUC results of the MACCS-based models. (C) AUC results of the Morgan-based models. (D) AUC results of the PharmacoPFP-based models.

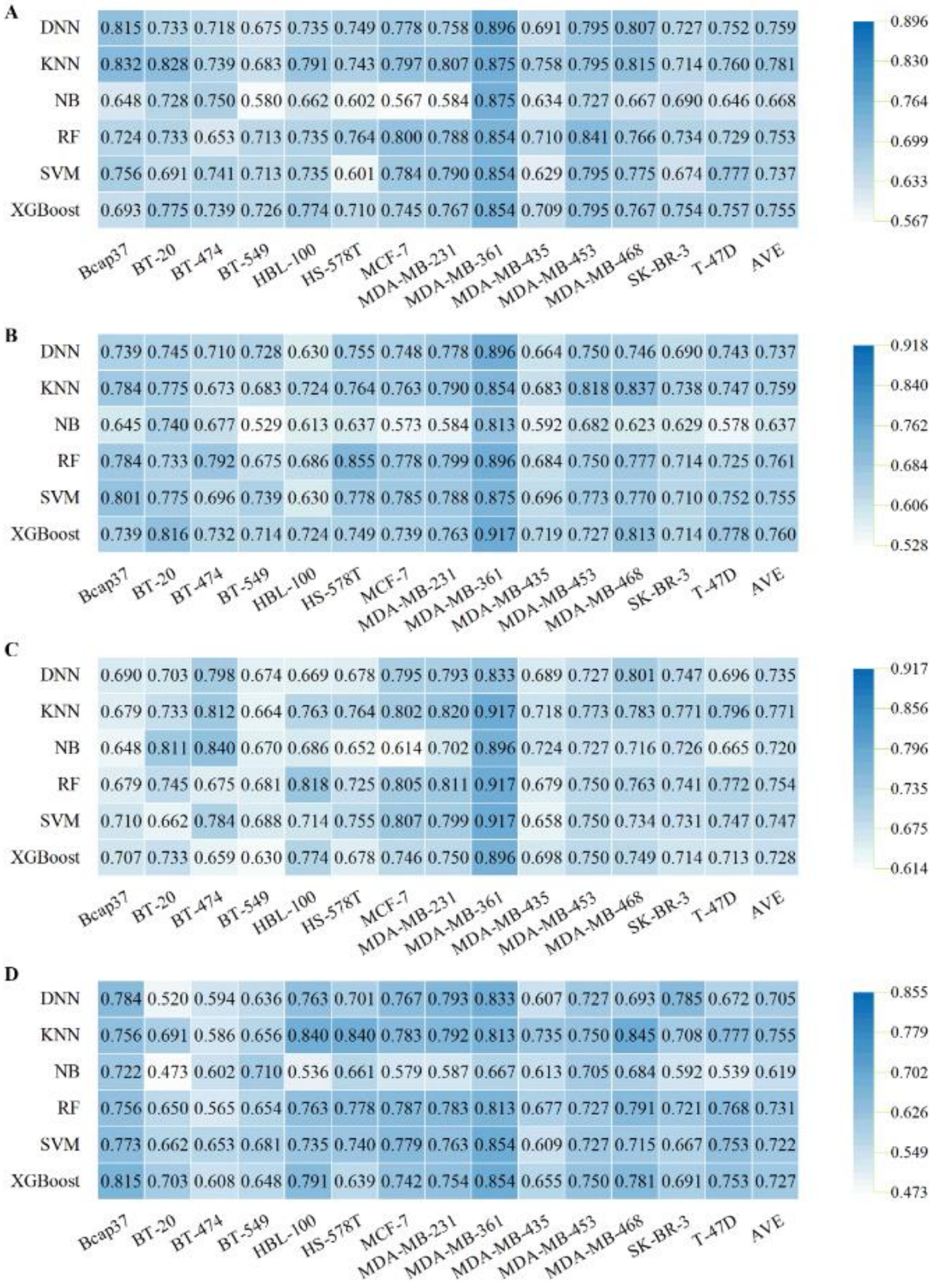


**Fig. S5.** Performance of fingerprint-based BC prediction models. (A) BA results of the AtomPairs-based models. (B) BA results of the MACCS-based models. (C) BA results of the Morgan-based models. (D) BA results of the PharmacoPFP-based models.

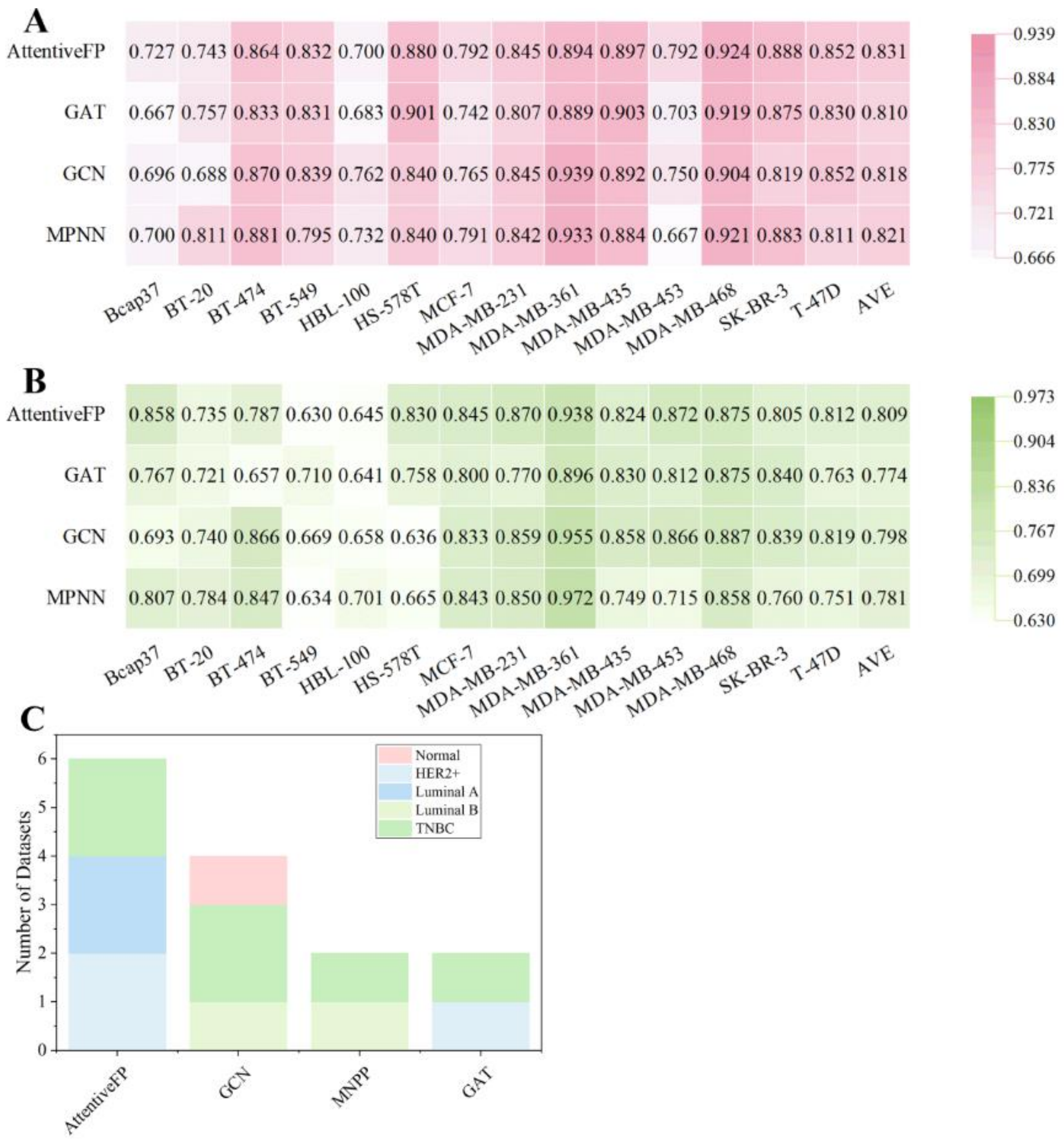


**Fig. S6.** Performance of graph-based BC prediction models. (A) F1 scores of graph-based models. (B) AUC results of graph-based models. (C) The optimal models based on molecular graph for different subtypes of breast cell lines.

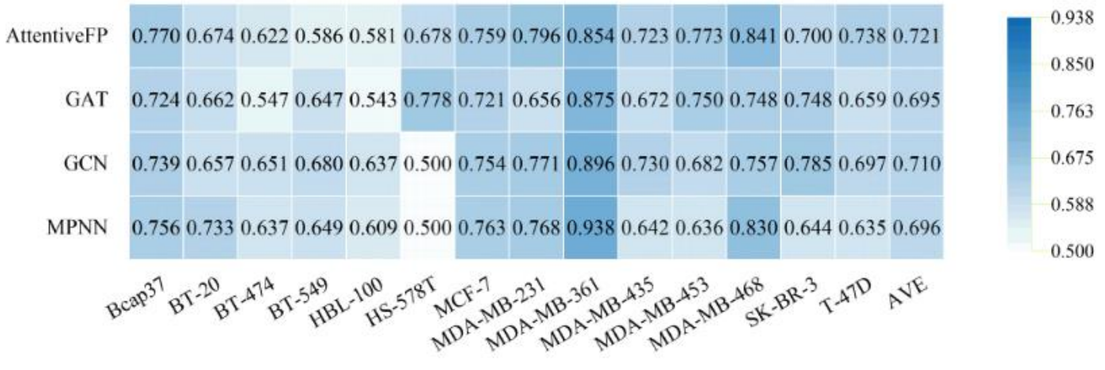


**Fig. S7.** BA results of graph-based BC prediction models.

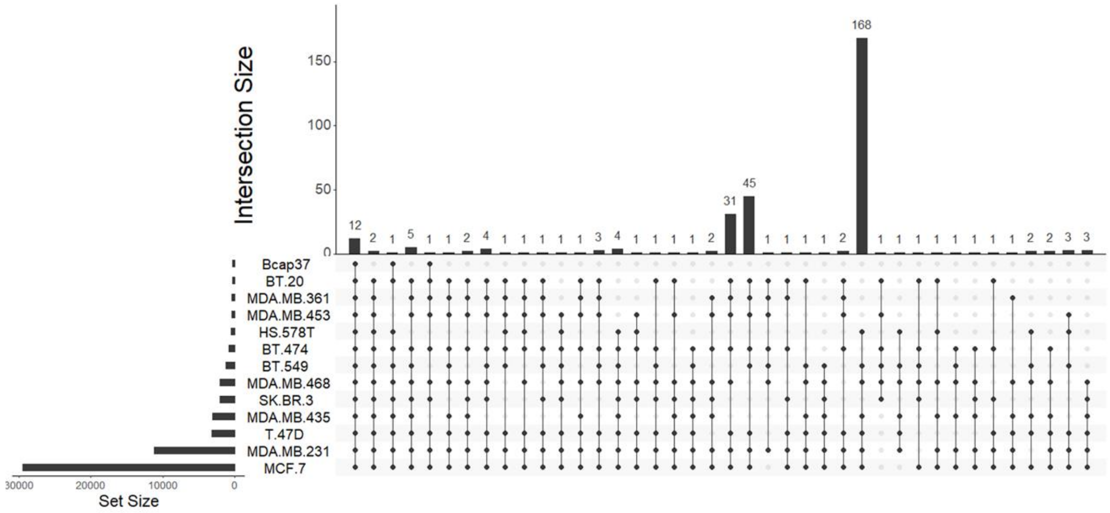


**Fig. S8.** Common molecules in 13 breast cancer cell lines.

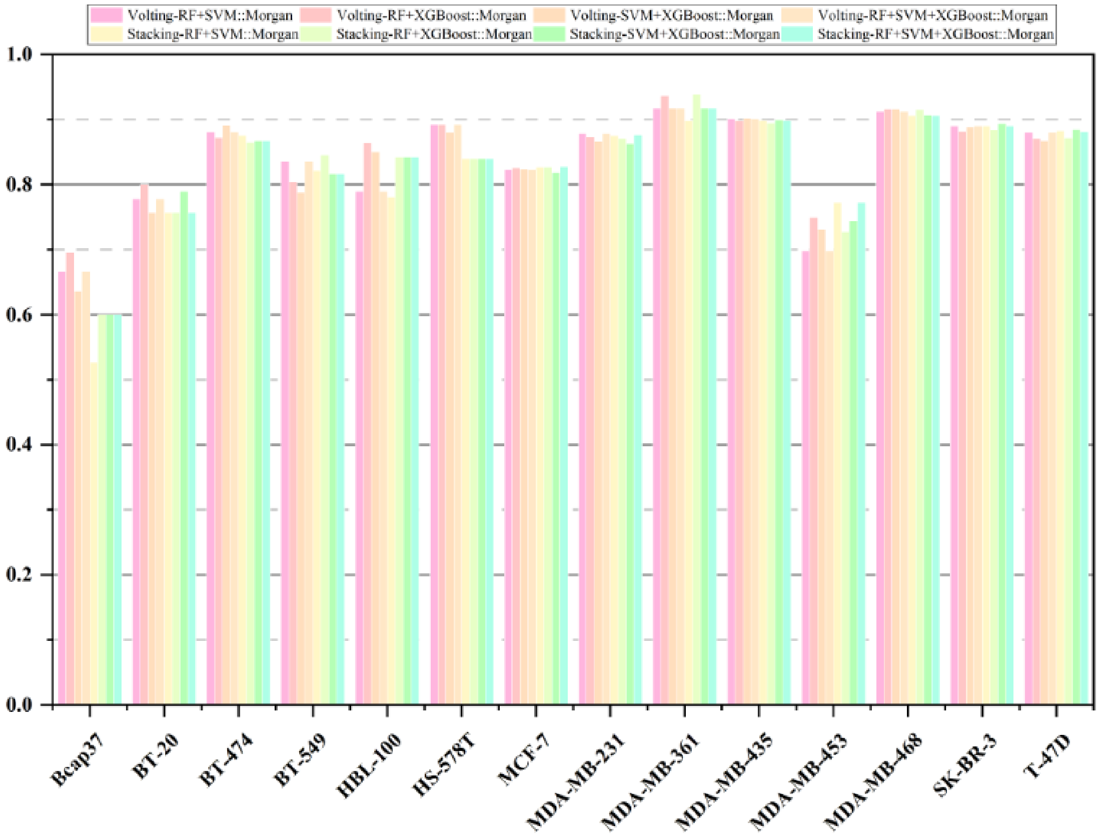


**Fig. S9.** F1 scores of fusion models.

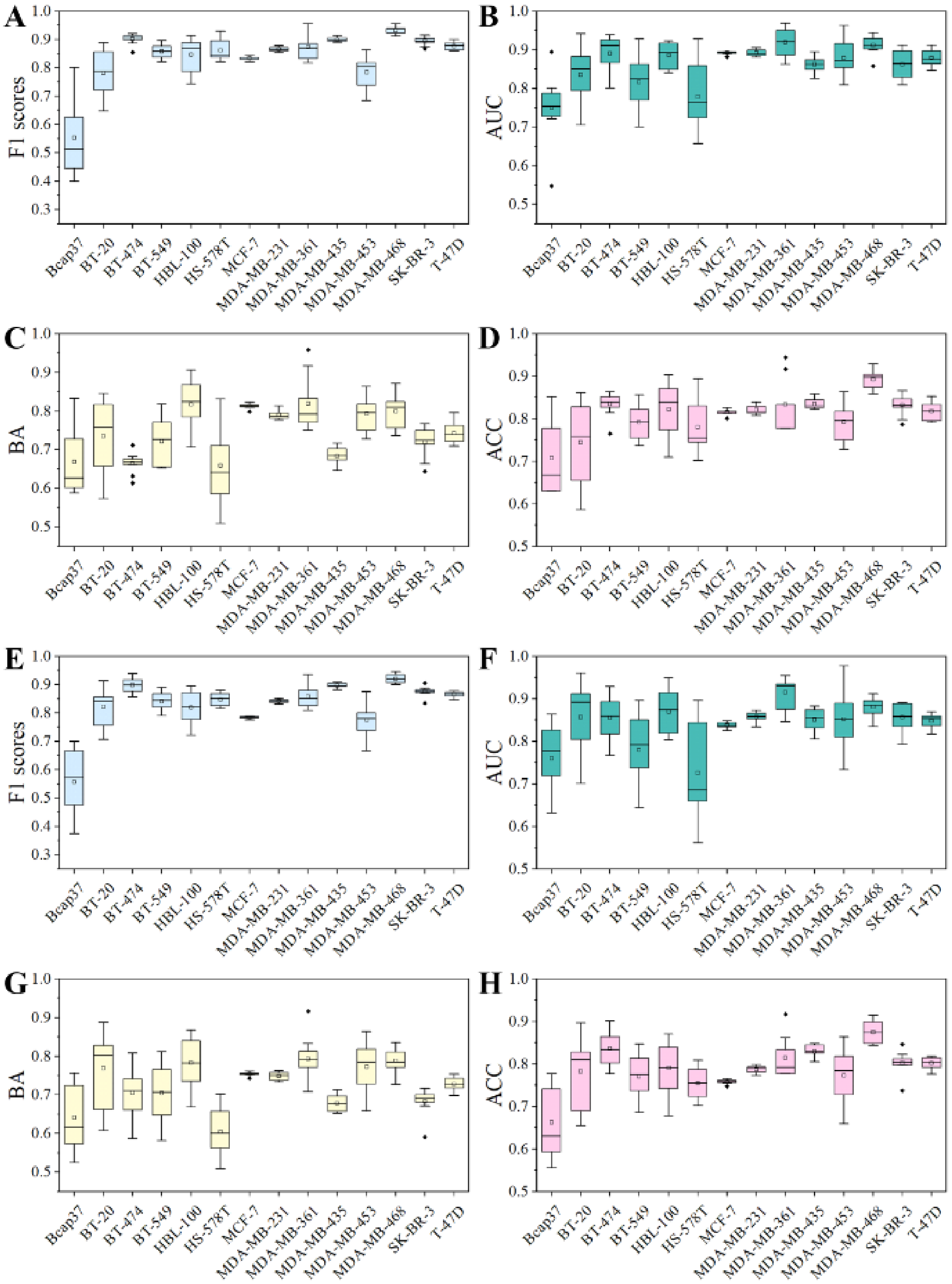


**Fig. S10.** The performance of 10 random seeds in RF and XGBoost::Morgan models. (A-D) F1scores, AUC, BA, and ACC results in RF::Morgan models; (E-H) F1scores, AUC, BA, and ACC results in XGBoost::Morgan models.

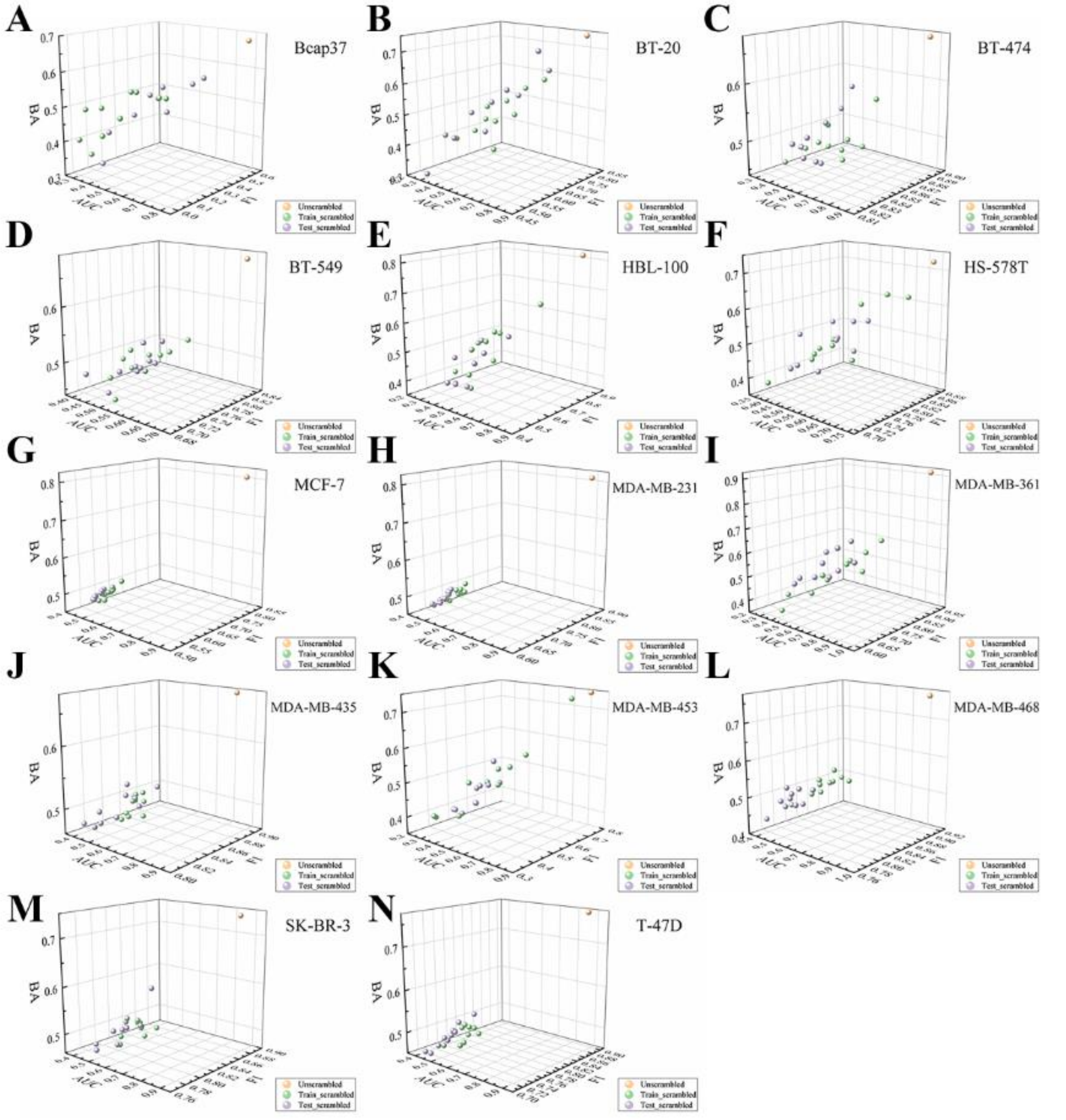


**Fig. S11.** Y-scrambling results for the RF::Morgan models. Both the training sets and testing sets were unscrambled (gold). The training sets were scrambled, whereas the test sets were unscrambled (green). The training sets were unscrambled, whereas the test sets were scrambled (purple). F1: F1-measure. BA: Balanced accuracy. AUC: Area under the receiver operating characteristics curve.

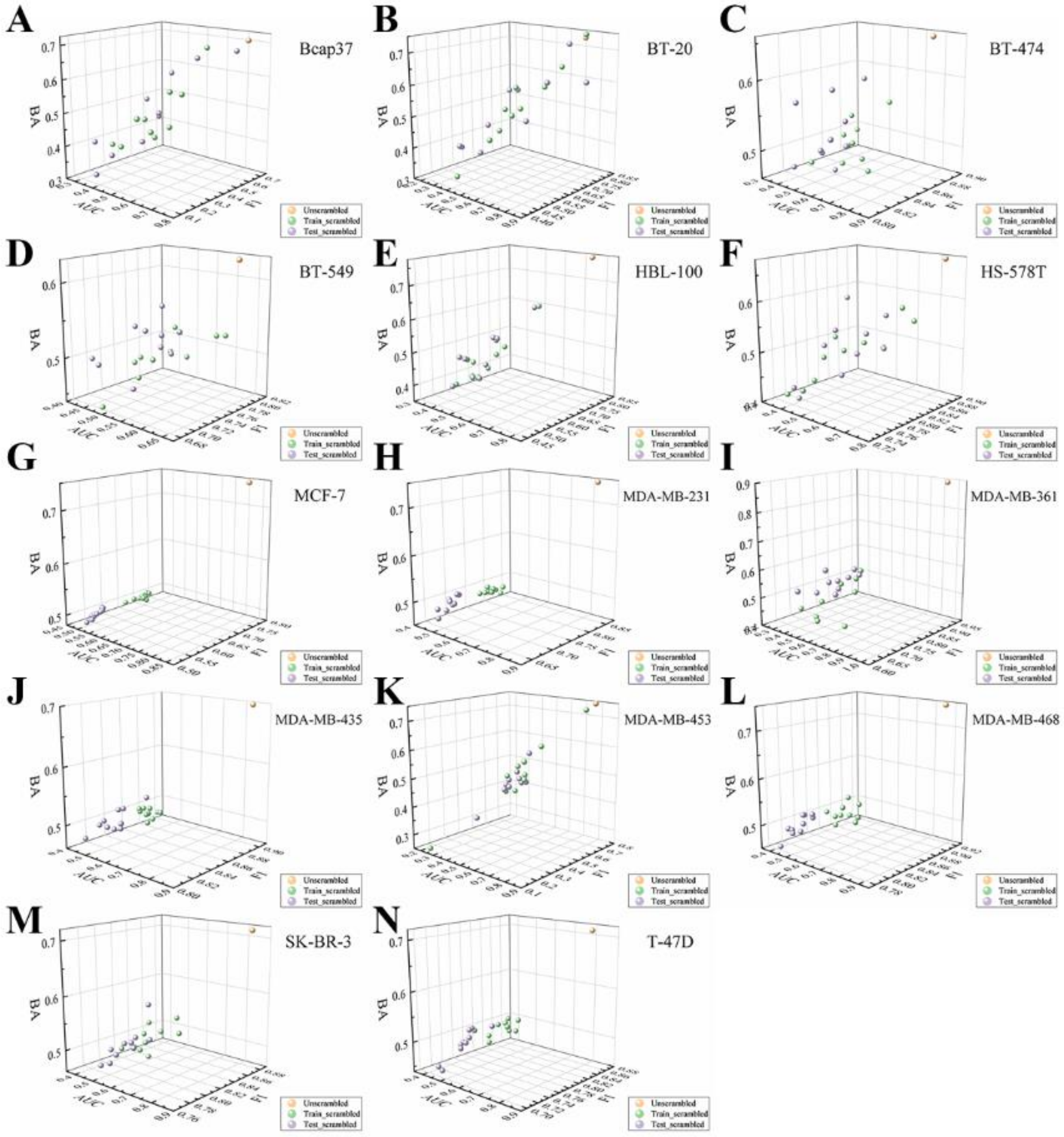


**Fig. S12.** Y-scrambling results for the XGB::Morgan models. Both the training sets and testing sets were unscrambled (gold). The training sets were scrambled, whereas the test sets were unscrambled (green). The training sets were unscrambled, whereas the test sets were scrambled (purple).F1: F1-measure. BA: Balanced accuracy. AUC: Area under the receiver operating characteristics curve.

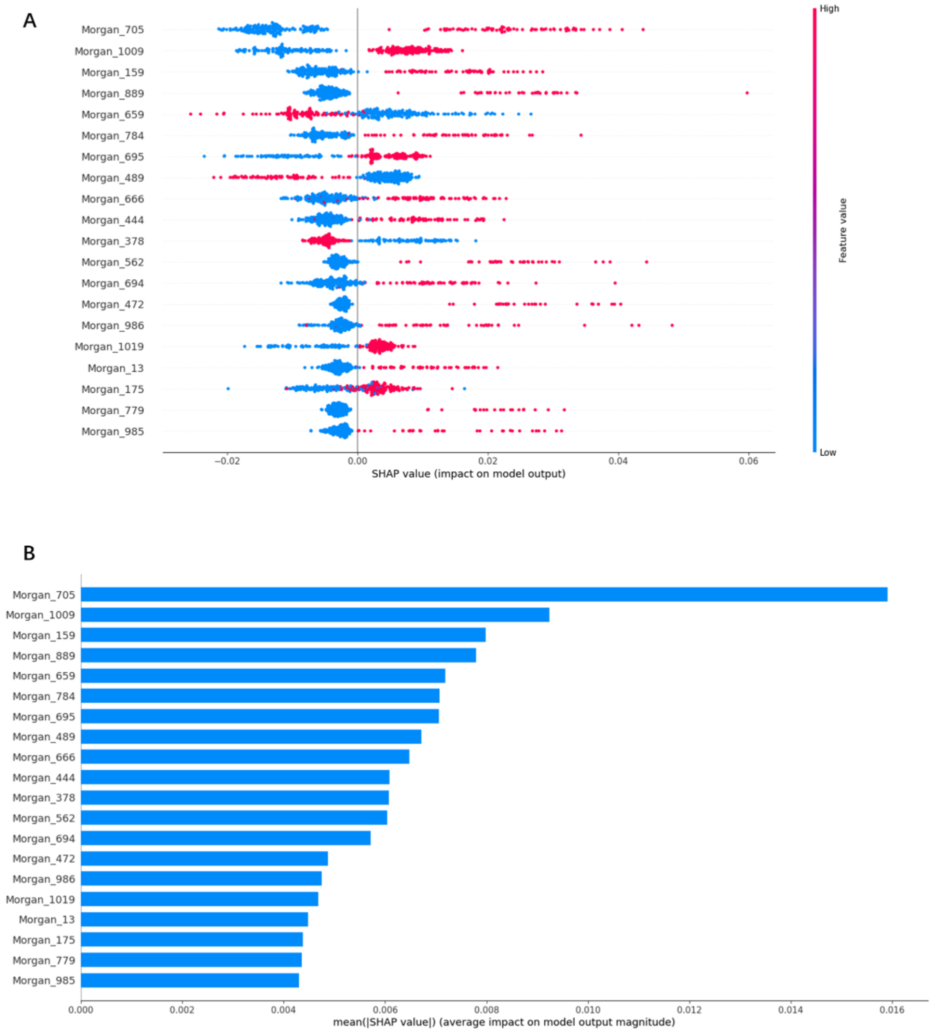


**Fig. S13.** Based on the top 20 most important features of the RF::Morgan model in Bcap37, (A) the SHAP values for each molecular substructure, and (B) the mean of the absolute value of the SHAP value for each molecular substructure.

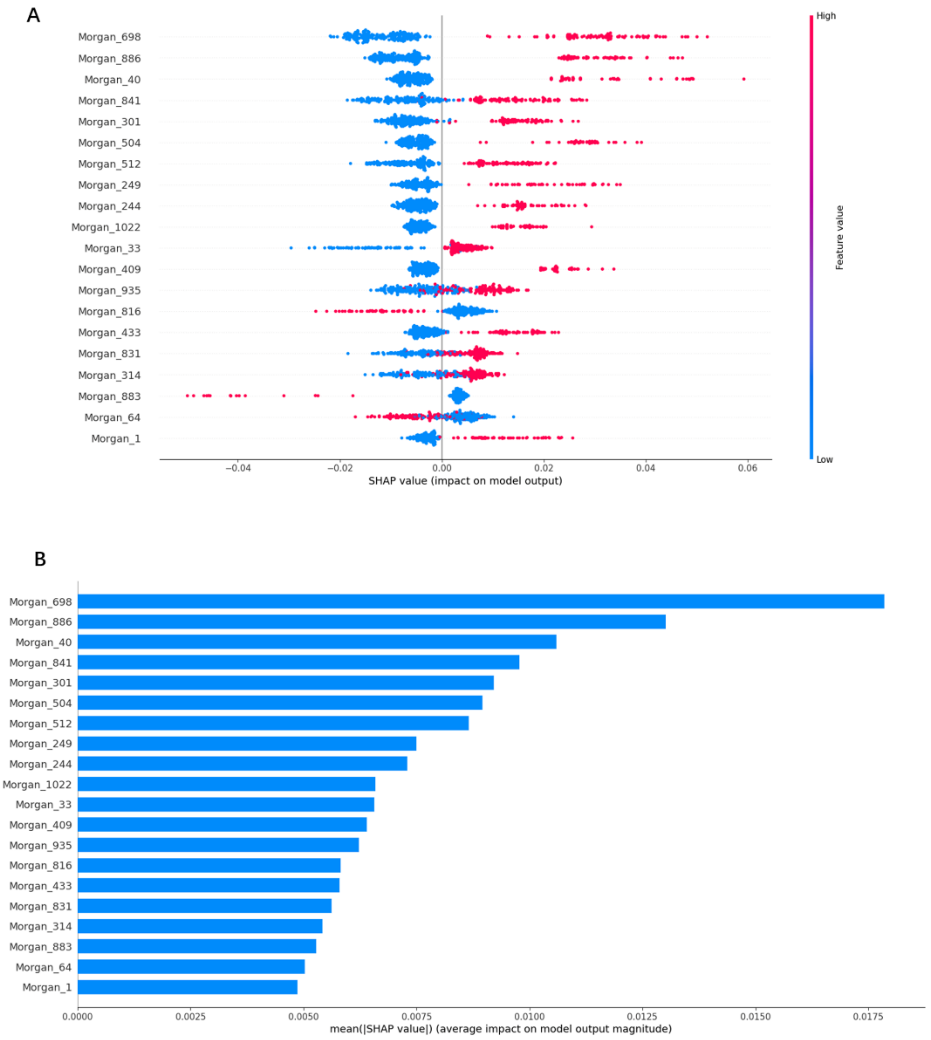


**Fig. S14.** Based on the top 20 most important features of the RF::Morgan model in BT-20, (A) the SHAP values for each molecular substructure, and (B) the mean of the absolute value of the SHAP value for each molecular substructure.

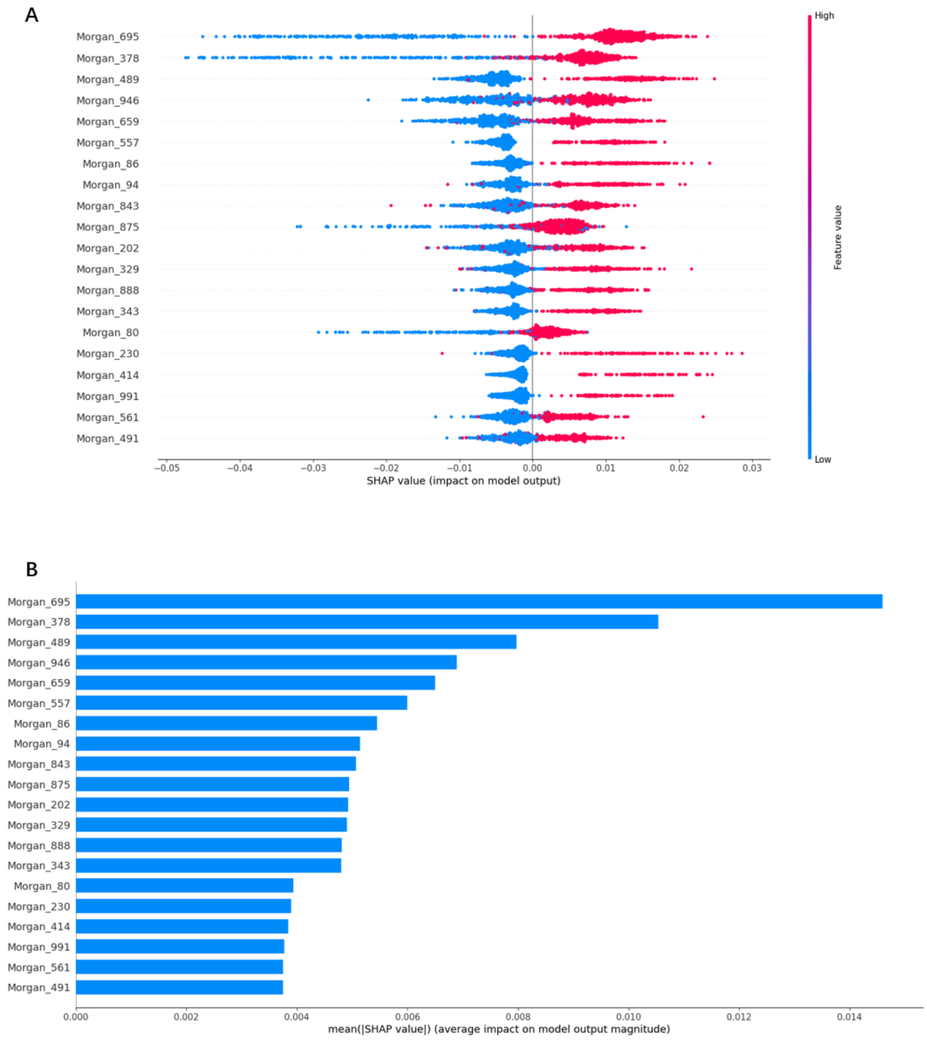


**Fig. S15.** Based on the top 20 most important features of the RF::Morgan model in BT-474, (A) the SHAP values for each molecular substructure, and (B) the mean of the absolute value of the SHAP value for each molecular substructure.

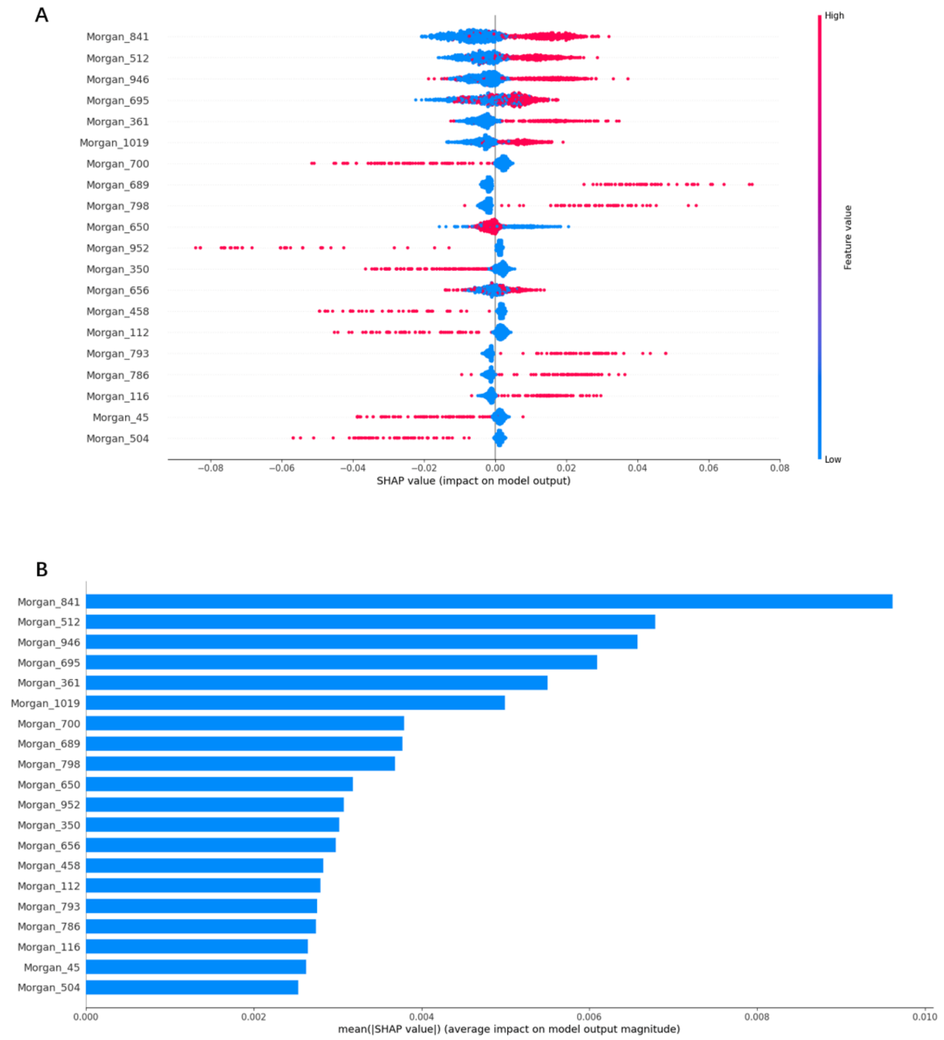


**Fig. S16.** Based on the top 20 most important features of the RF::Morgan model in BT-549, (A) the SHAP values for each molecular substructure, and (B) the mean of the absolute value of the SHAP value for each molecular substructure.

A

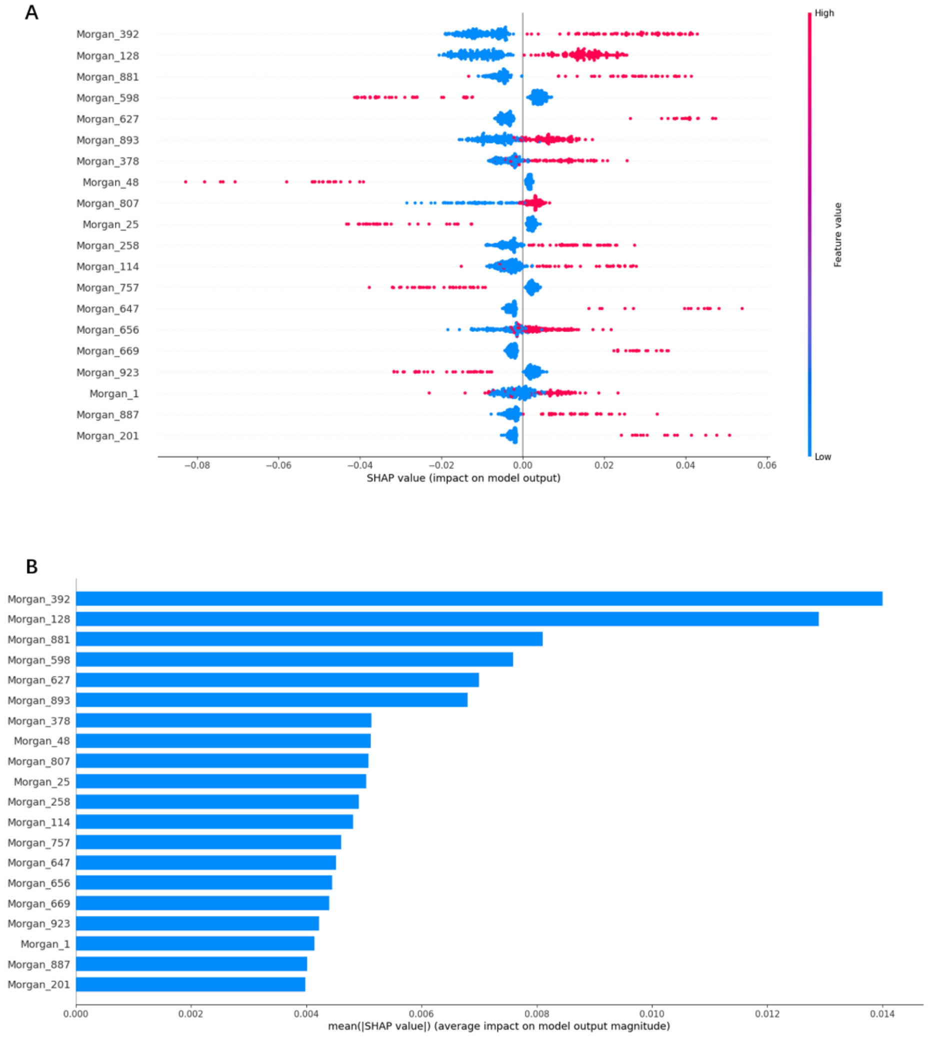


B



**Fig. S17.** Based on the top 20 most important features of the RF::Morgan model in HBL-100, (A) the SHAP values for each molecular substructure, and (B) the mean of the absolute value of the SHAP value for each molecular substructure.

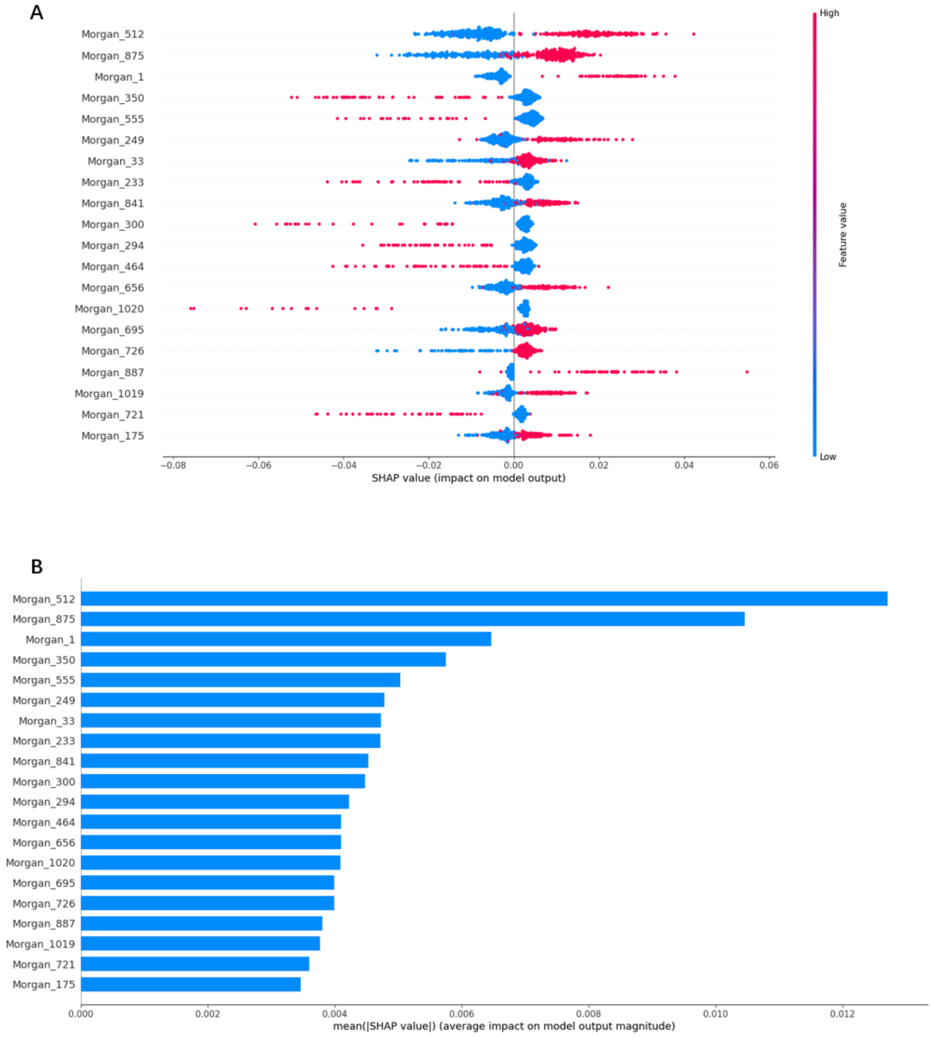


**Fig. S18.** Based on the top 20 most important features of the RF::Morgan model in HS-578T (A) the SHAP values for each molecular substructure, and (B) the mean of the absolute value of the SHAP value for each molecular substructure.

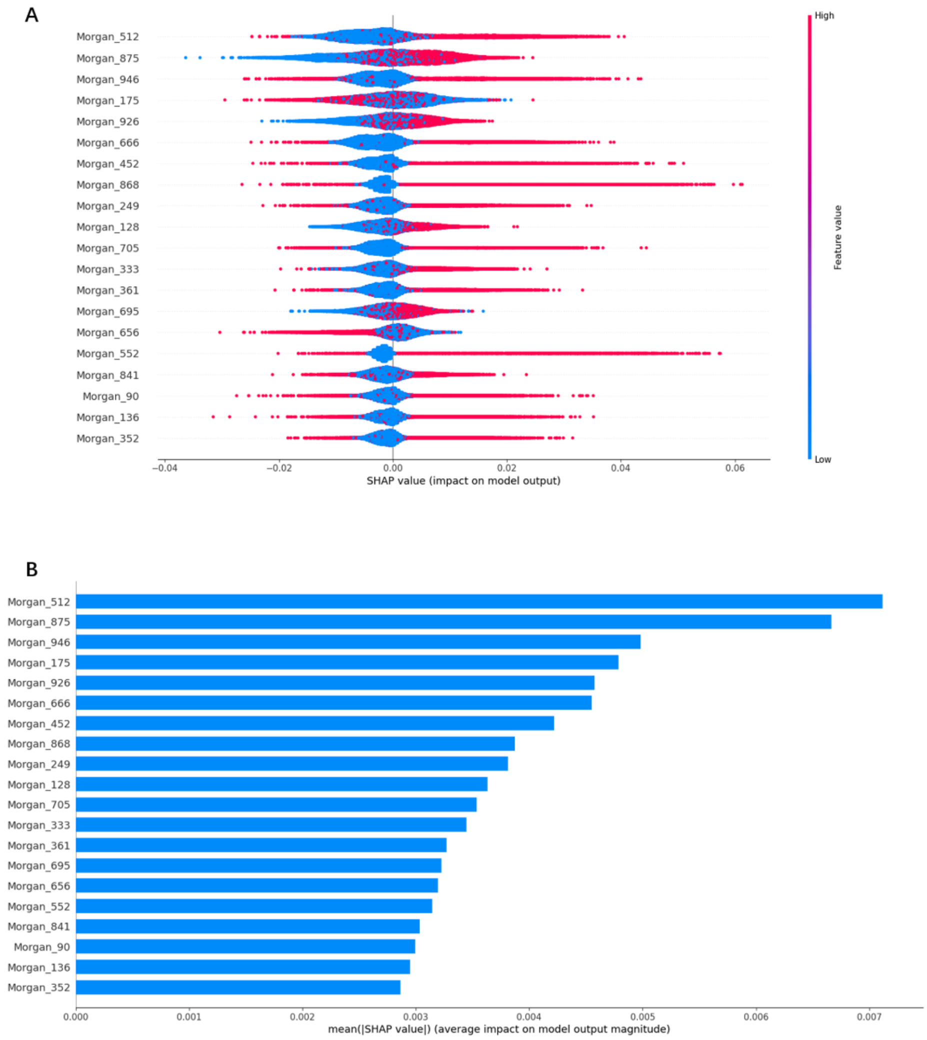


**Fig. S19.** Based on the top 20 most important features of the RF::Morgan model in MCF-7 (A) the SHAP values for each molecular substructure, and (B) the mean of the absolute value of the SHAP value for each molecular substructure.

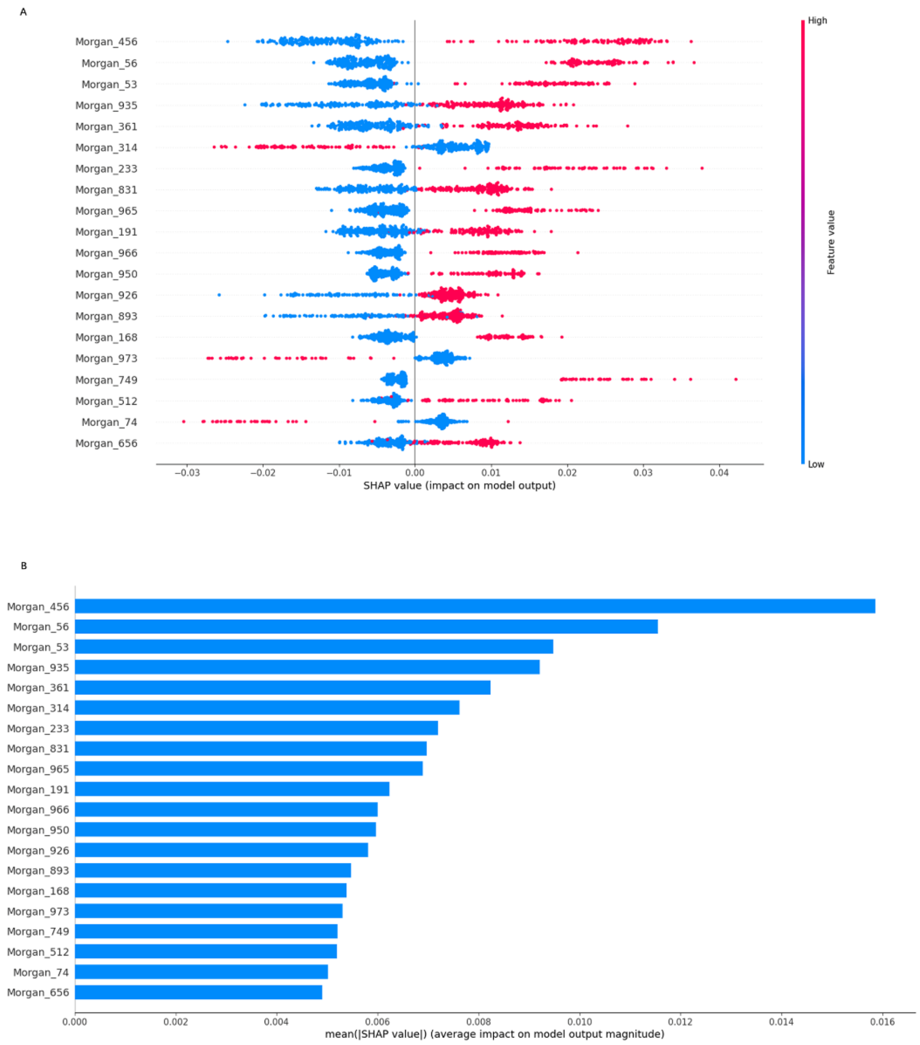


**Fig. S20.** Based on the top 20 most important features of the RF::Morgan model in MDA-MB-361 (A) the SHAP values for each molecular substructure, and (B) the mean of the absolute value of the SHAP value for each molecular substructure.

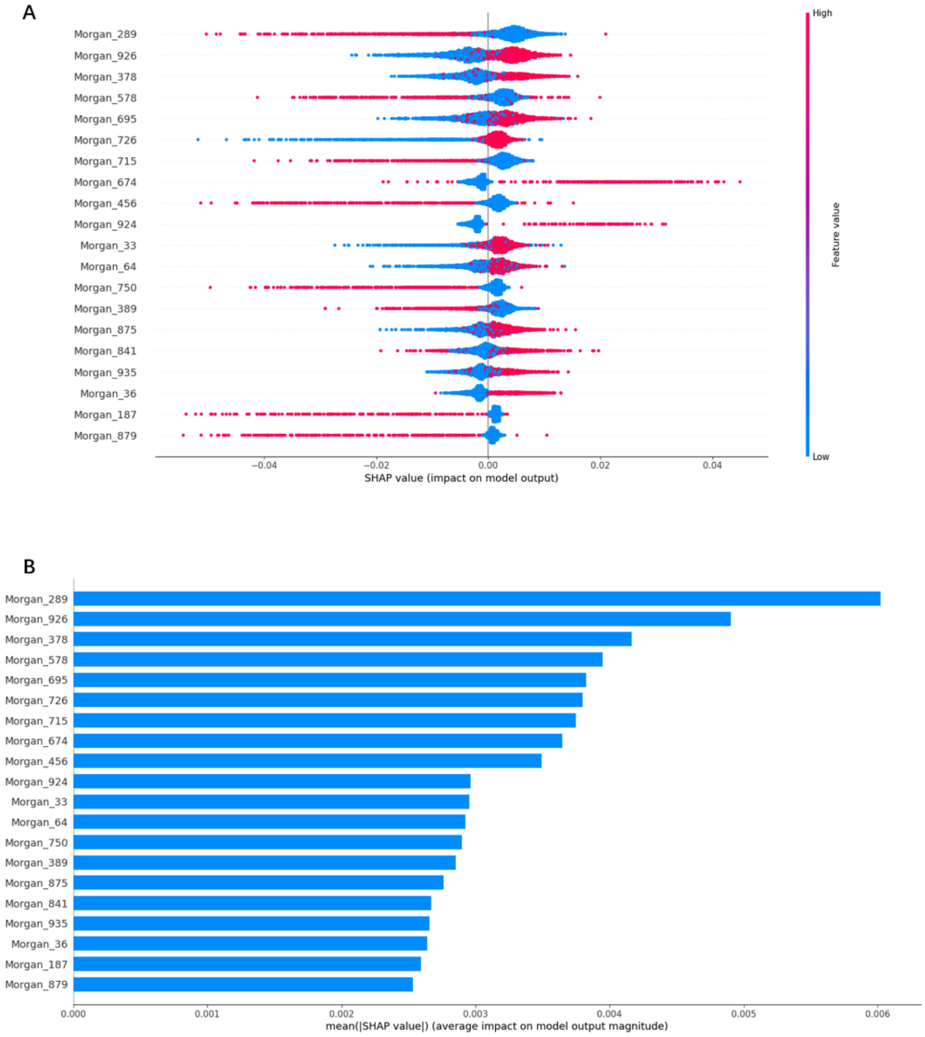


**Fig. S21.** Based on the top 20 most important features of the RF::Morgan model in MDA-MB-435 (A) the SHAP values for each molecular substructure, and (B) the mean of the absolute value of the SHAP value for each molecular substructure.

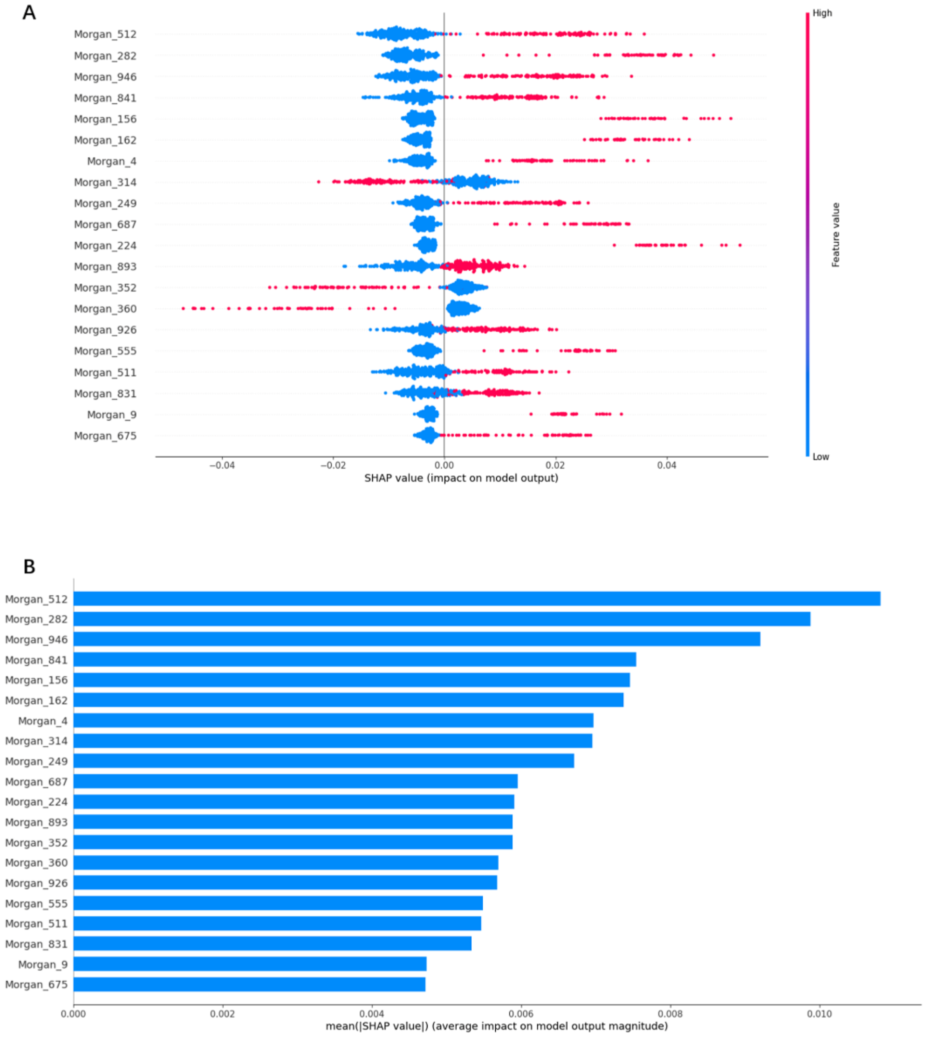


**Fig. S22.** Based on the top 20 most important features of the RF::Morgan model in MDA-MB-453 (A) the SHAP values for each molecular substructure, and (B) the mean of the absolute value of the SHAP value for each molecular substructure.

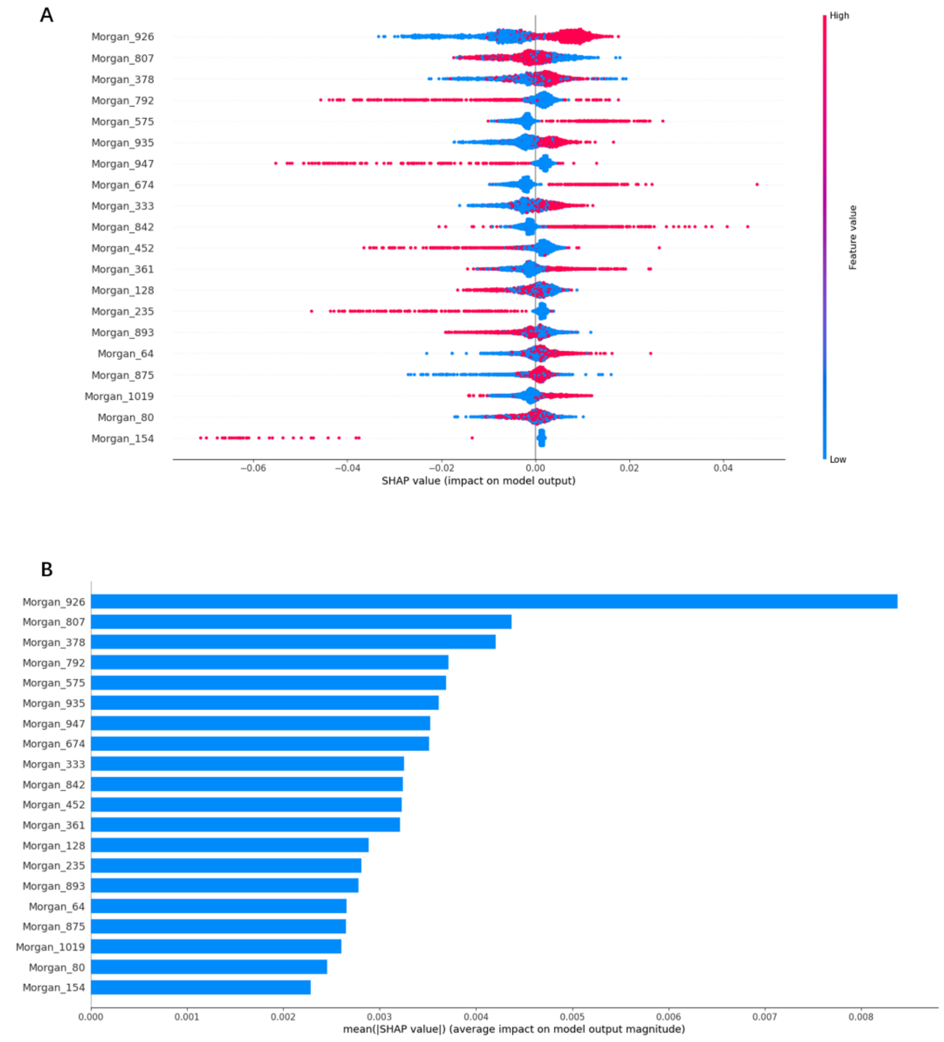


**Fig. S23.** Based on the top 20 most important features of the RF::Morgan model in MDA-MB-468 (A) the SHAP values for each molecular substructure, and (B) the mean of the absolute value of the SHAP value for each molecular substructure.

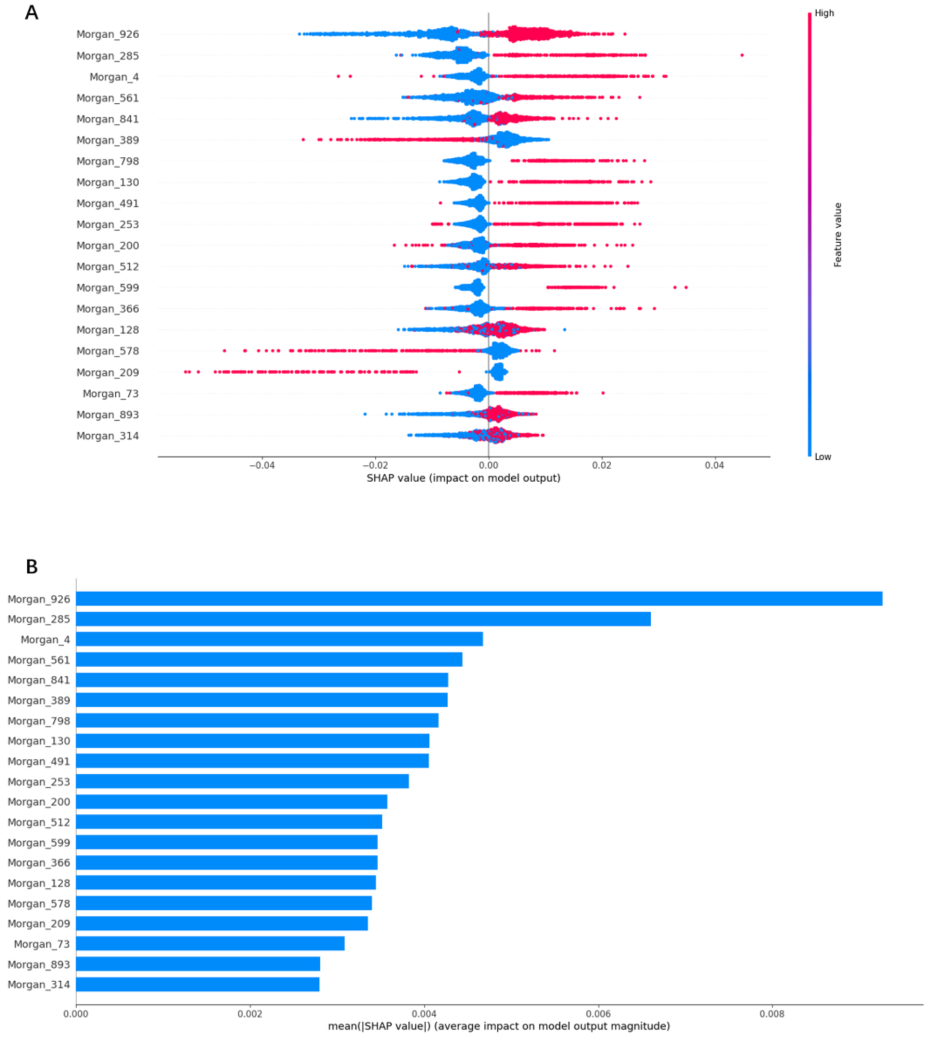


**Fig. S24.** Based on the top 20 most important features of the RF::Morgan model in SK-BR-3 (A) the SHAP values for each molecular substructure, and (B) the mean of the absolute value of the SHAP value for each molecular substructure.

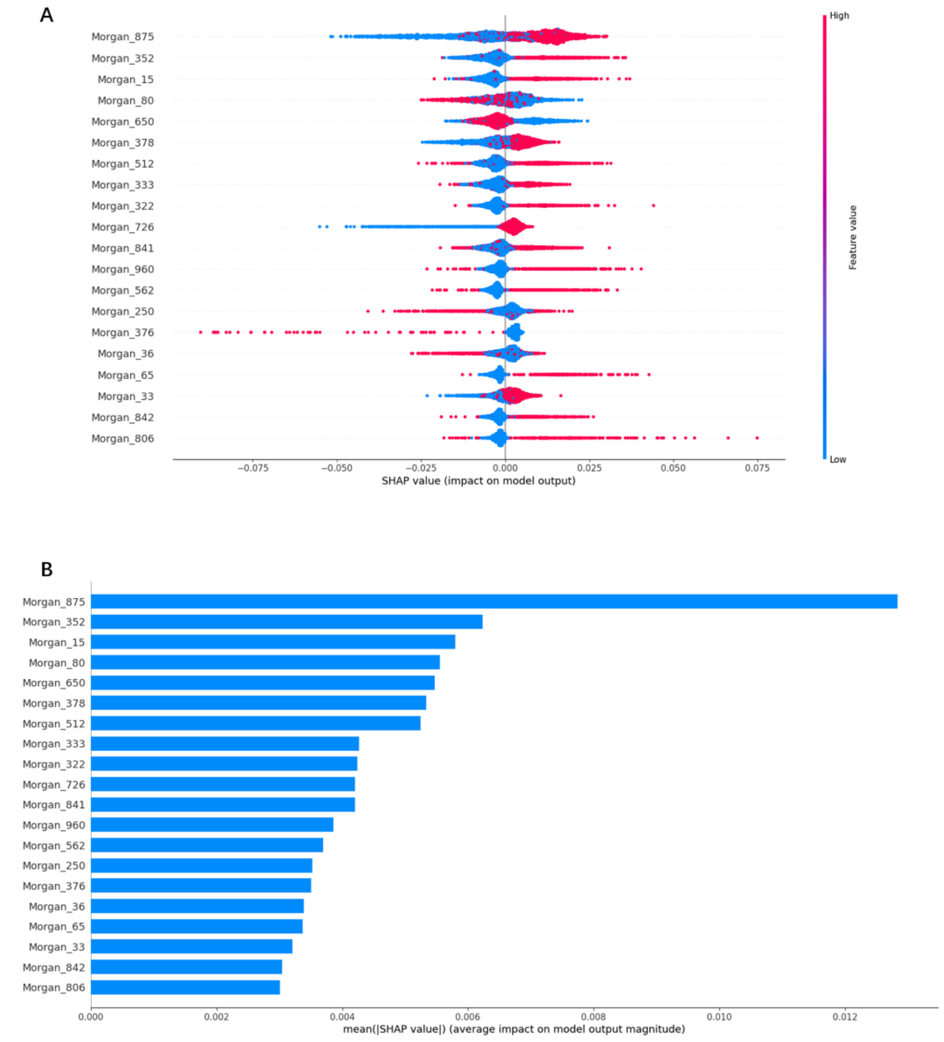


**Fig. S25.** Based on the top 20 most important features of the RF::Morgan model in T-47D (A) the SHAP values for each molecular substructure, and (B) the mean of the absolute value of the SHAP value for each molecular substructure.

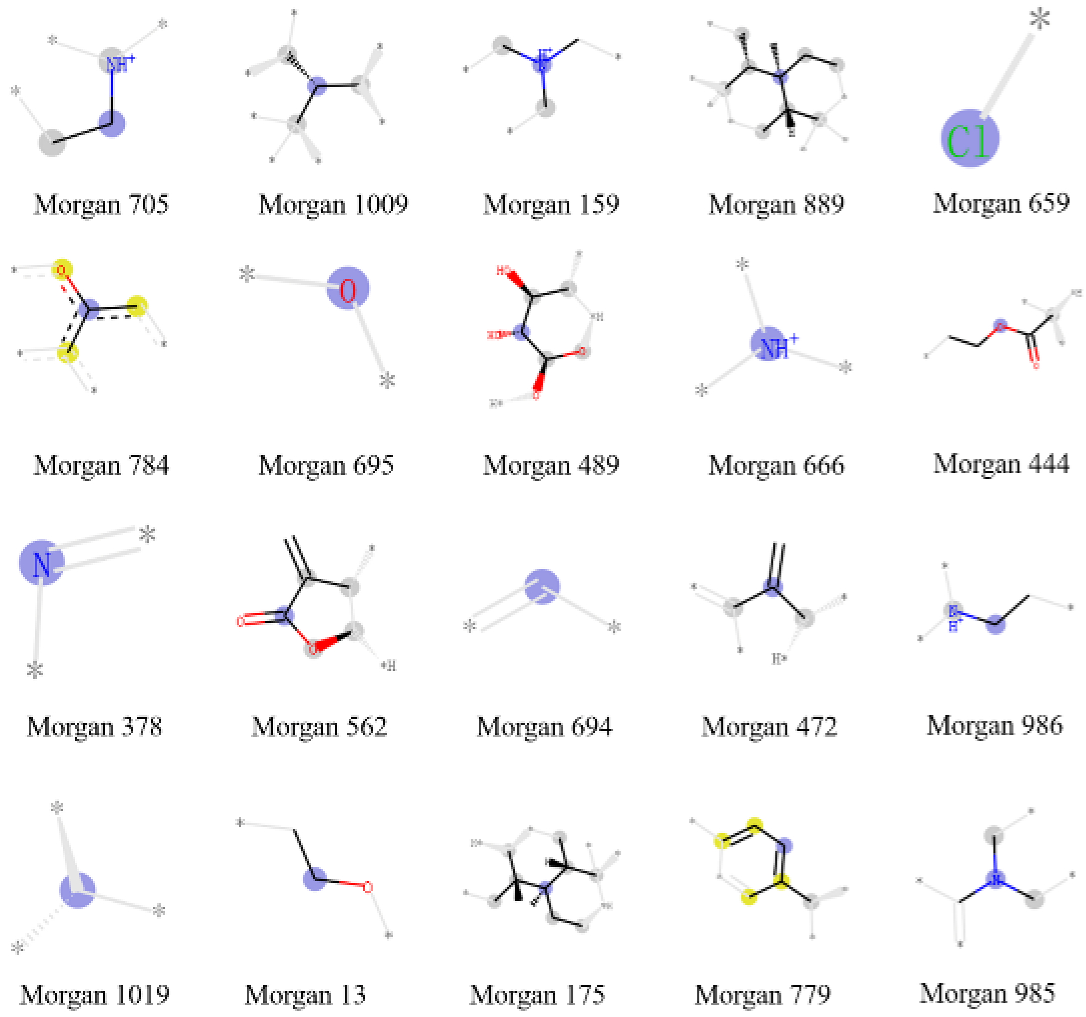


**Fig. S26.** Important molecular substructures of the RF::Morgan model in Bcap37.

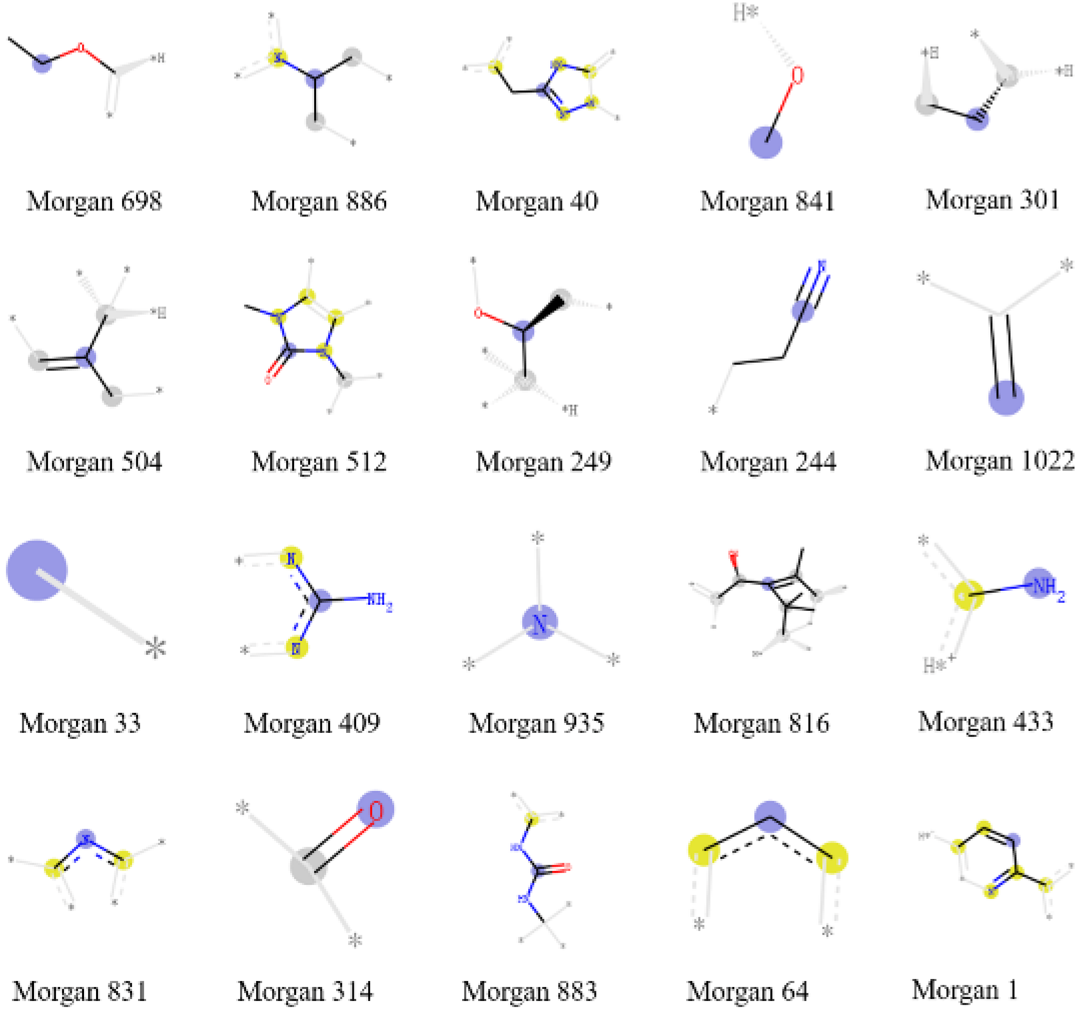


**Fig. S27.** Important molecular substructures of the RF::Morgan model in BT-20.

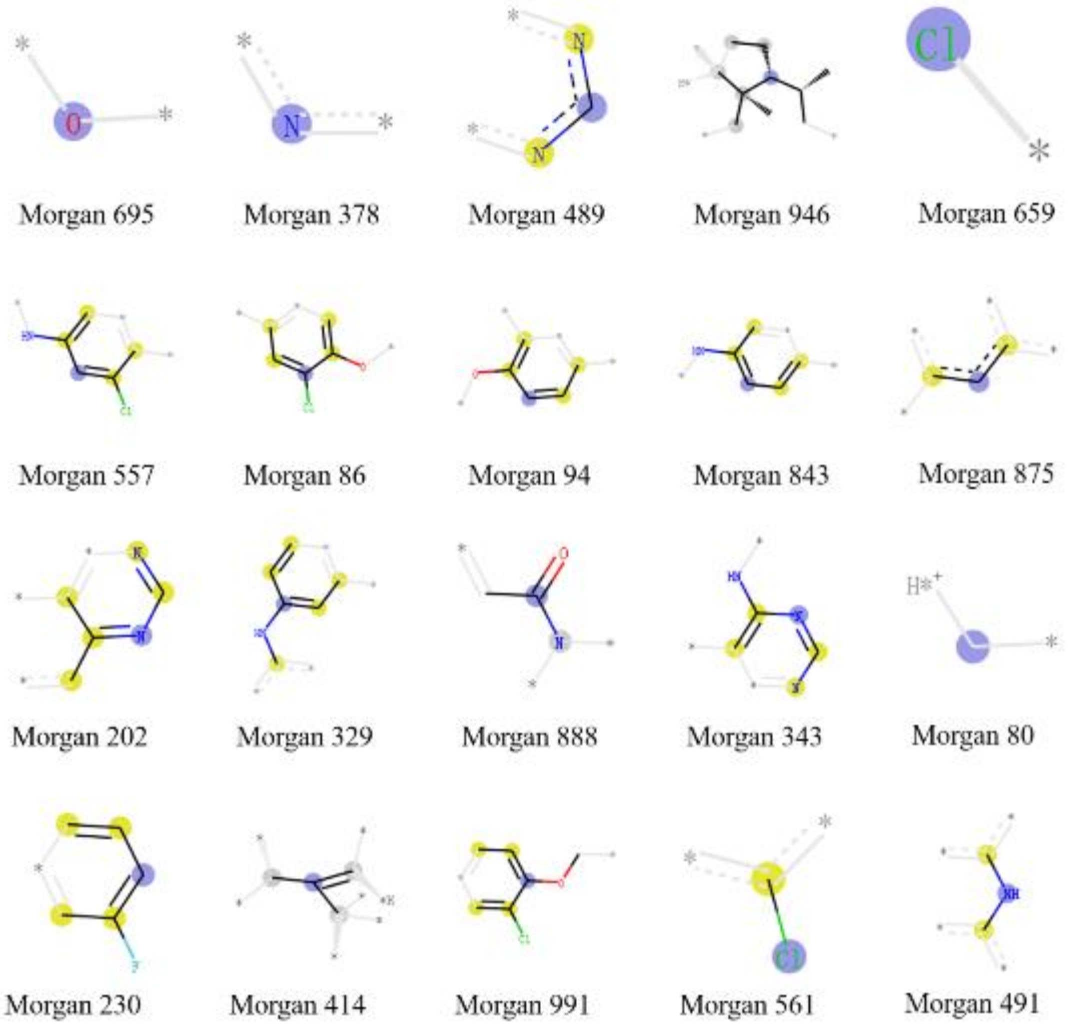


**Fig. S28.** Important molecular substructures of the RF::Morgan model in BT-474.

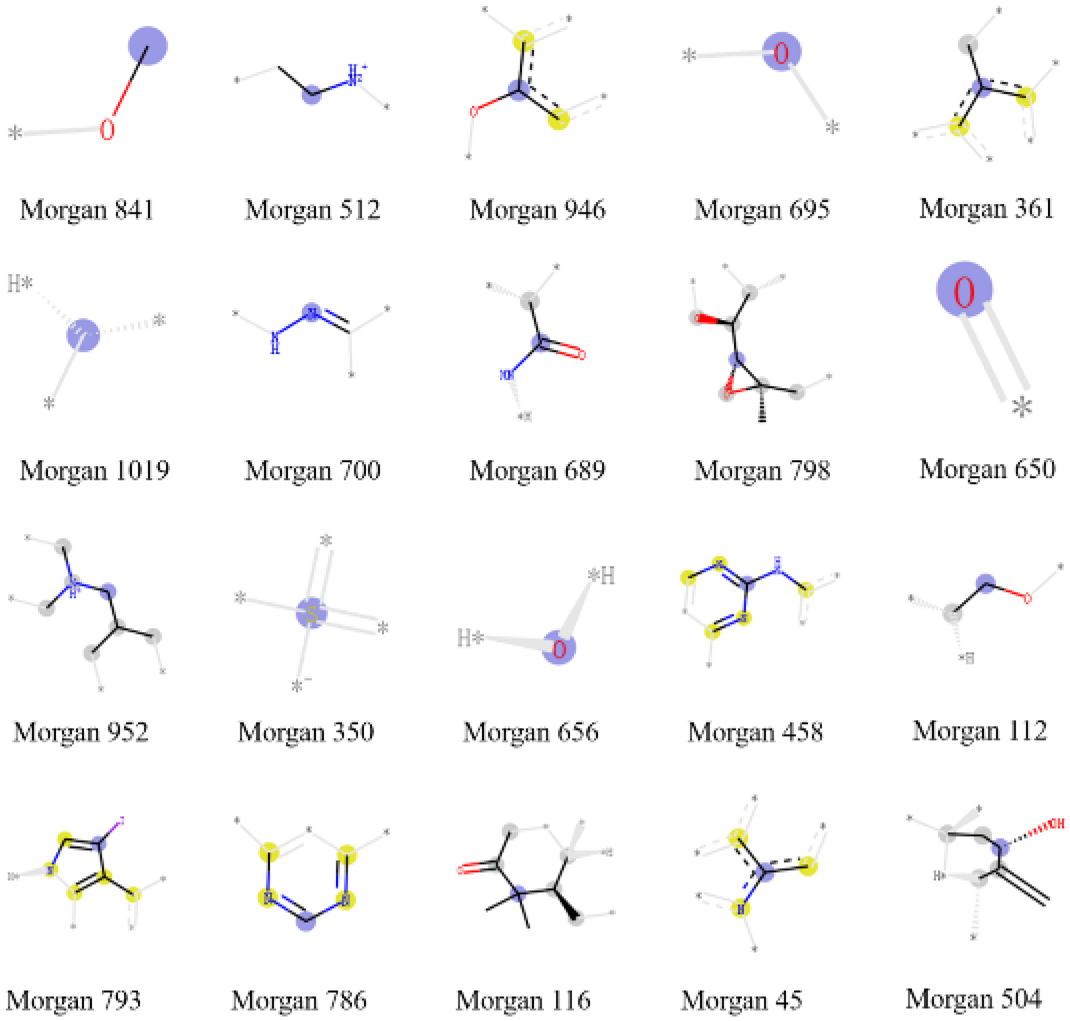


**Fig. S29.** Important molecular substructures of the RF::Morgan model in BT-549.

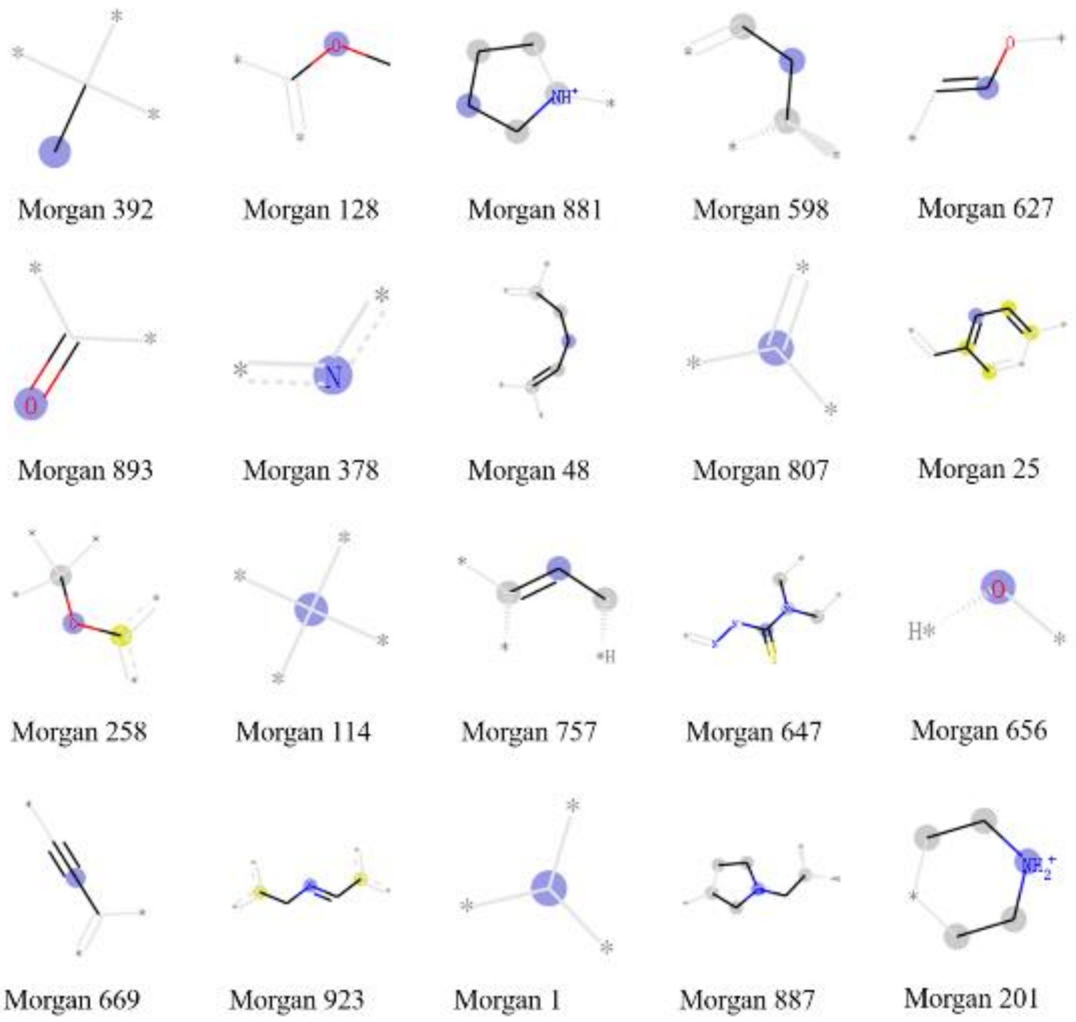


**Fig. S30.** Important molecular substructures of the RF::Morgan model in HBL-100.

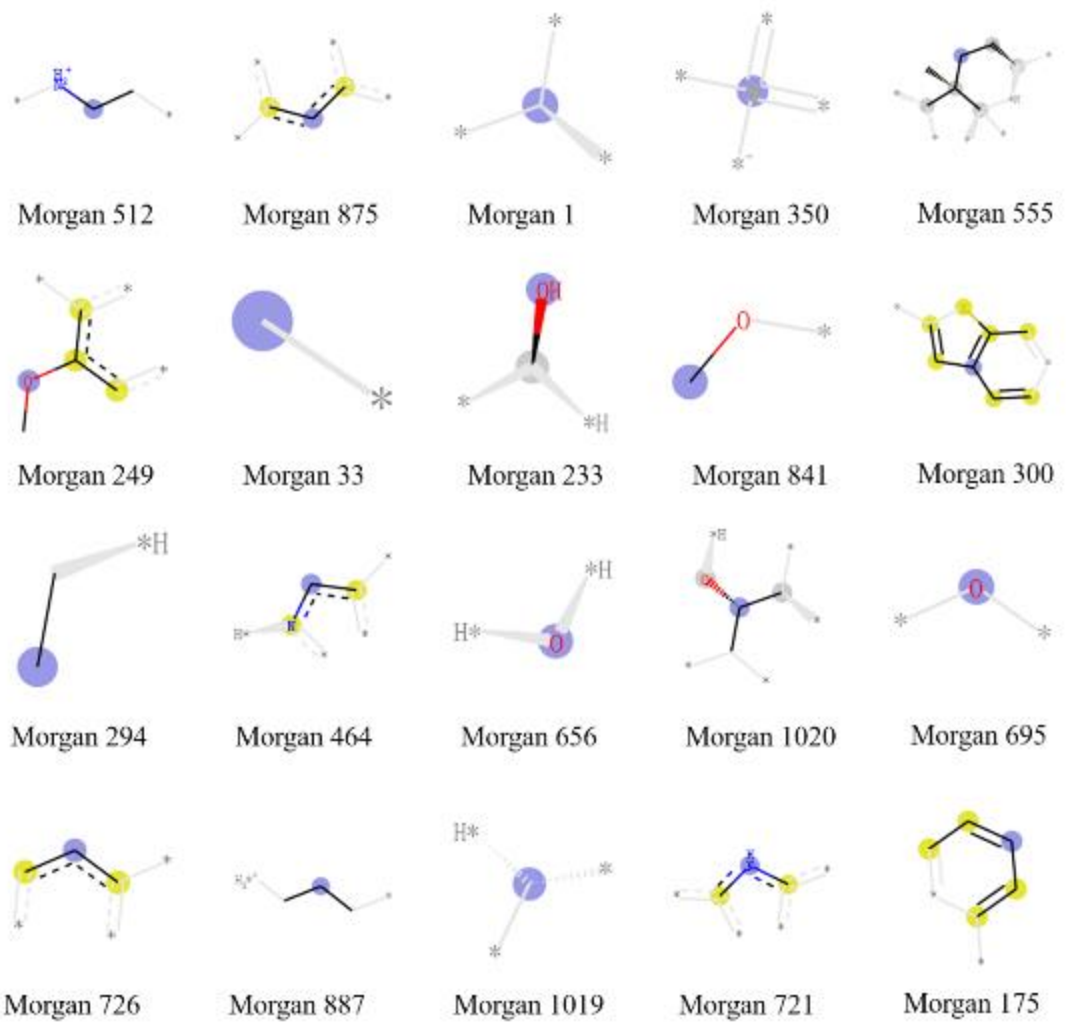


**Fig. S31.** Important molecular substructures of the RF::Morgan model in HS-578T.

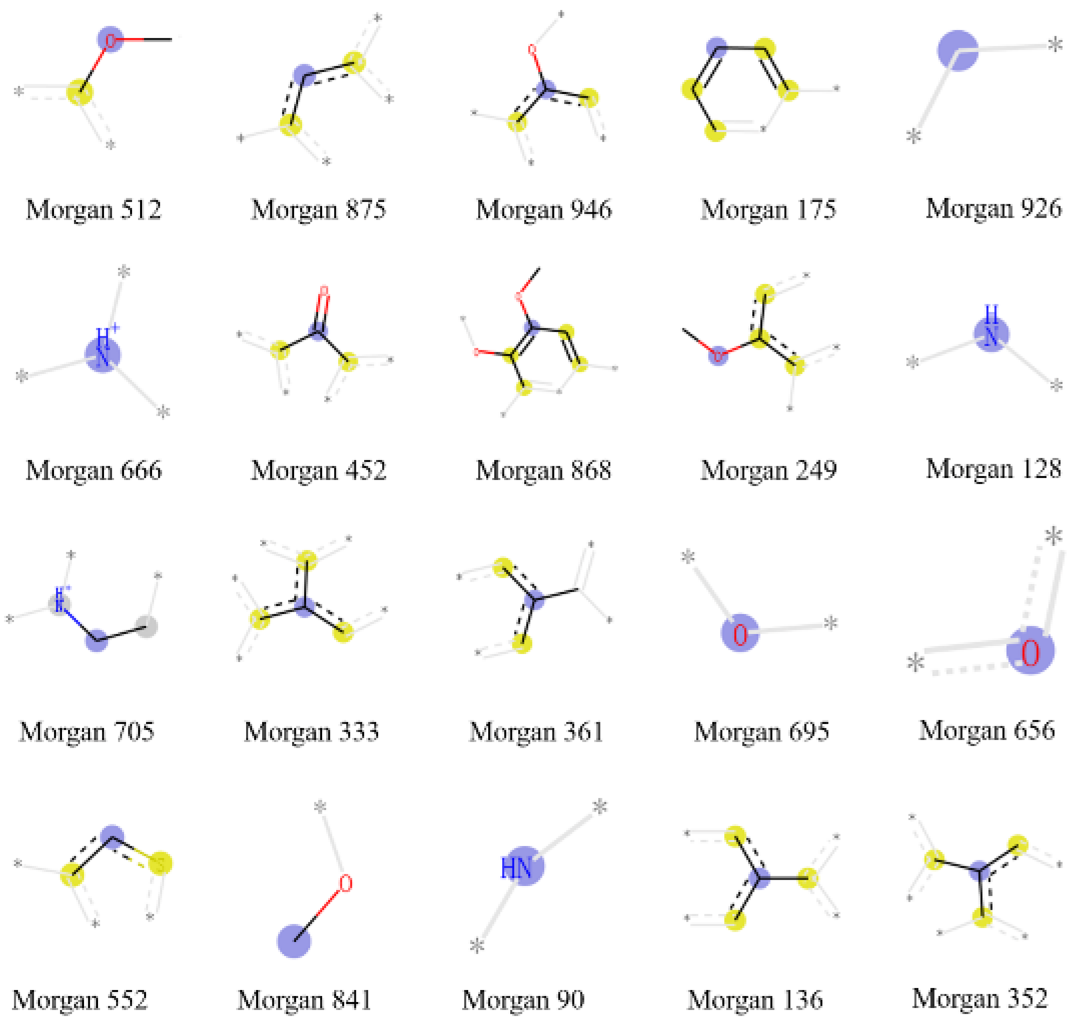


**Fig. S32.** Important molecular substructures of the RF::Morgan model in MCF-7.

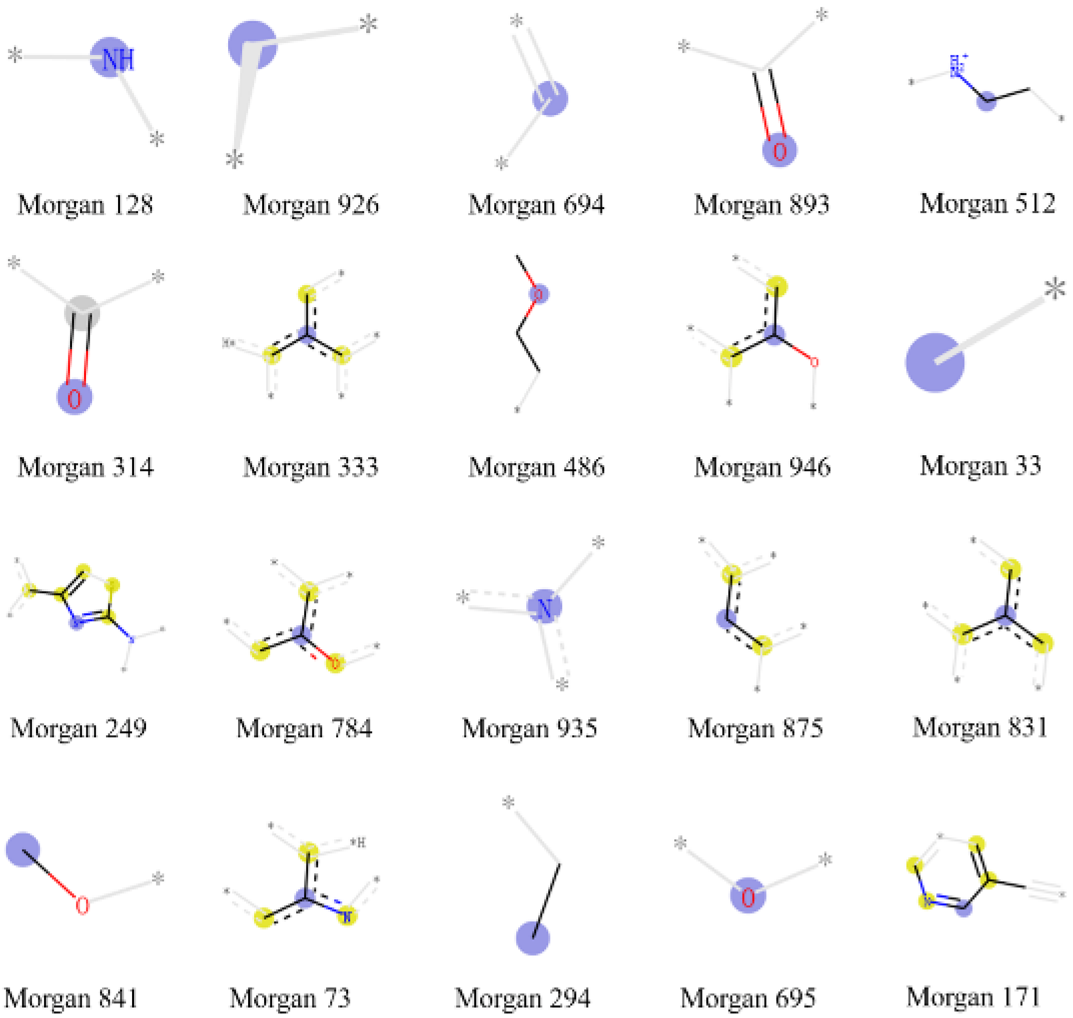


**Fig. S33.** Important molecular substructures of the RF::Morgan model in MDA-MB-231.

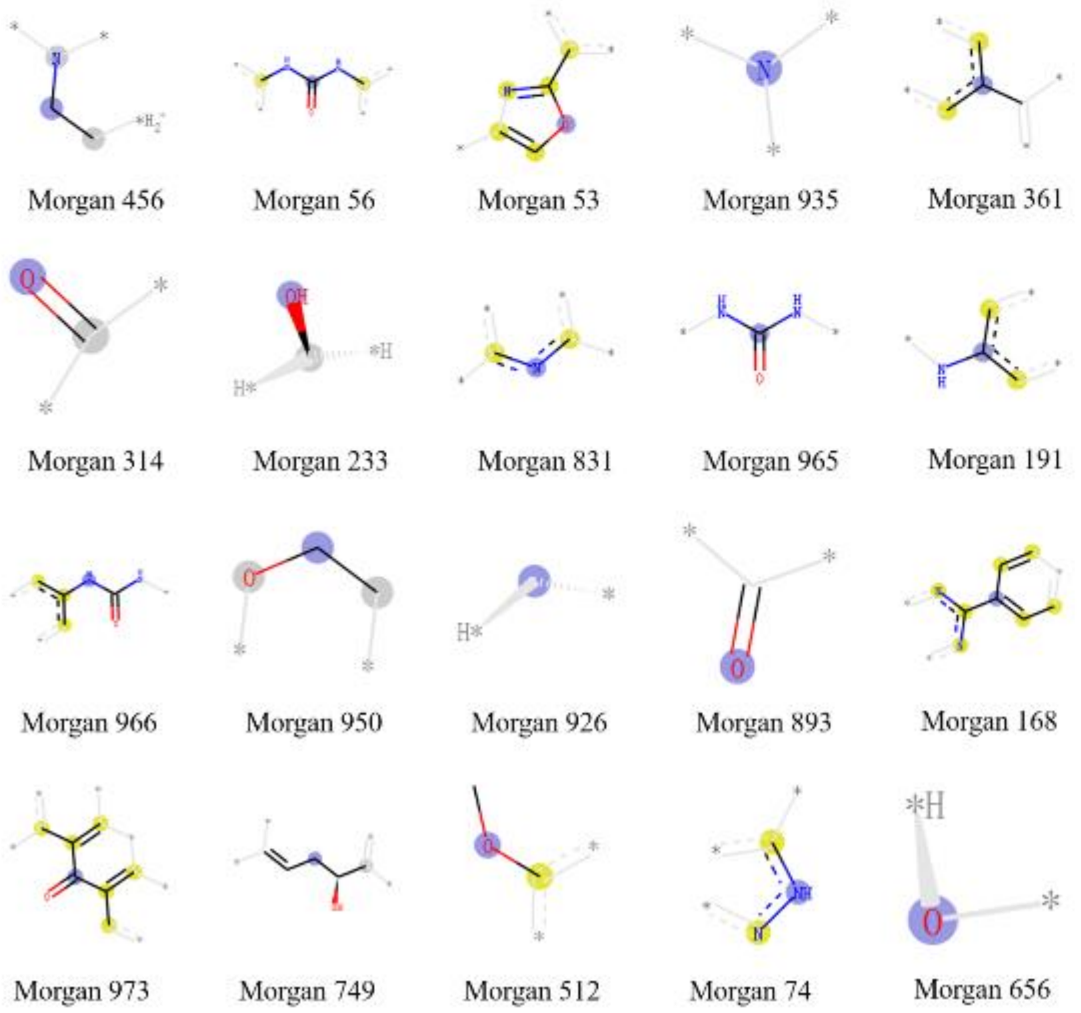


**Fig. S34.** Important molecular substructures of the RF::Morgan model in MDA-MB-361.

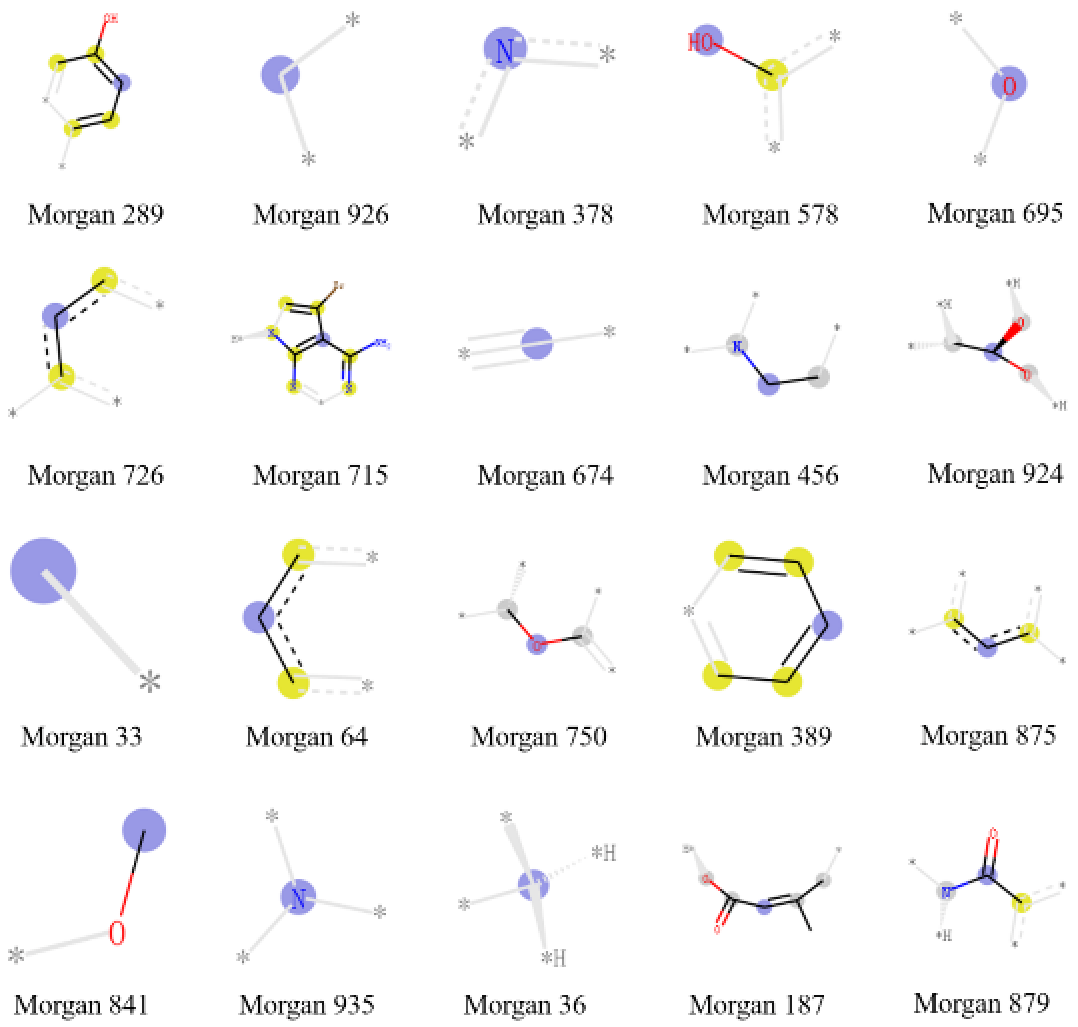


**Fig. S35.** Important molecular substructures of the RF::Morgan model in MDA-MB-435.

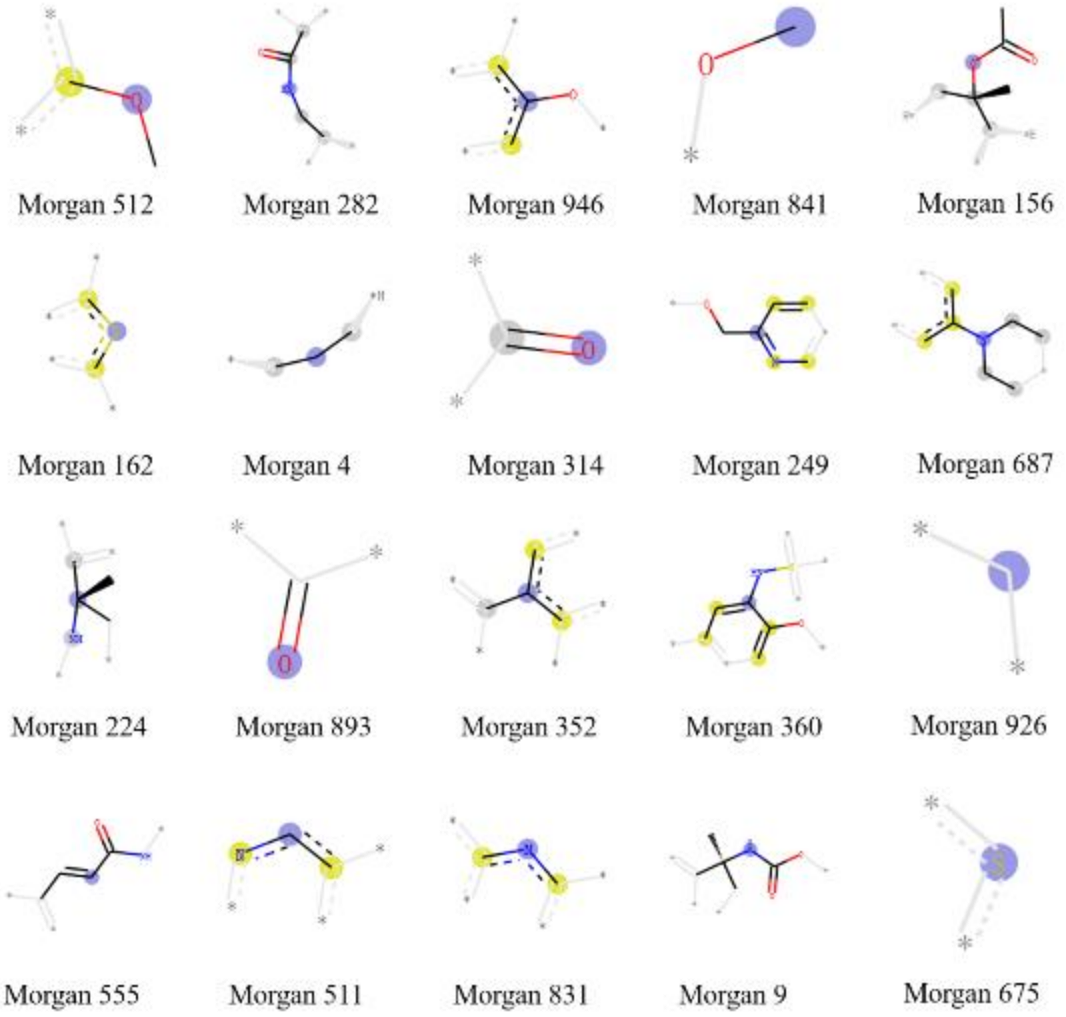


**Fig. S36.** Important molecular substructures of the RF::Morgan model in MDA-MB-453.

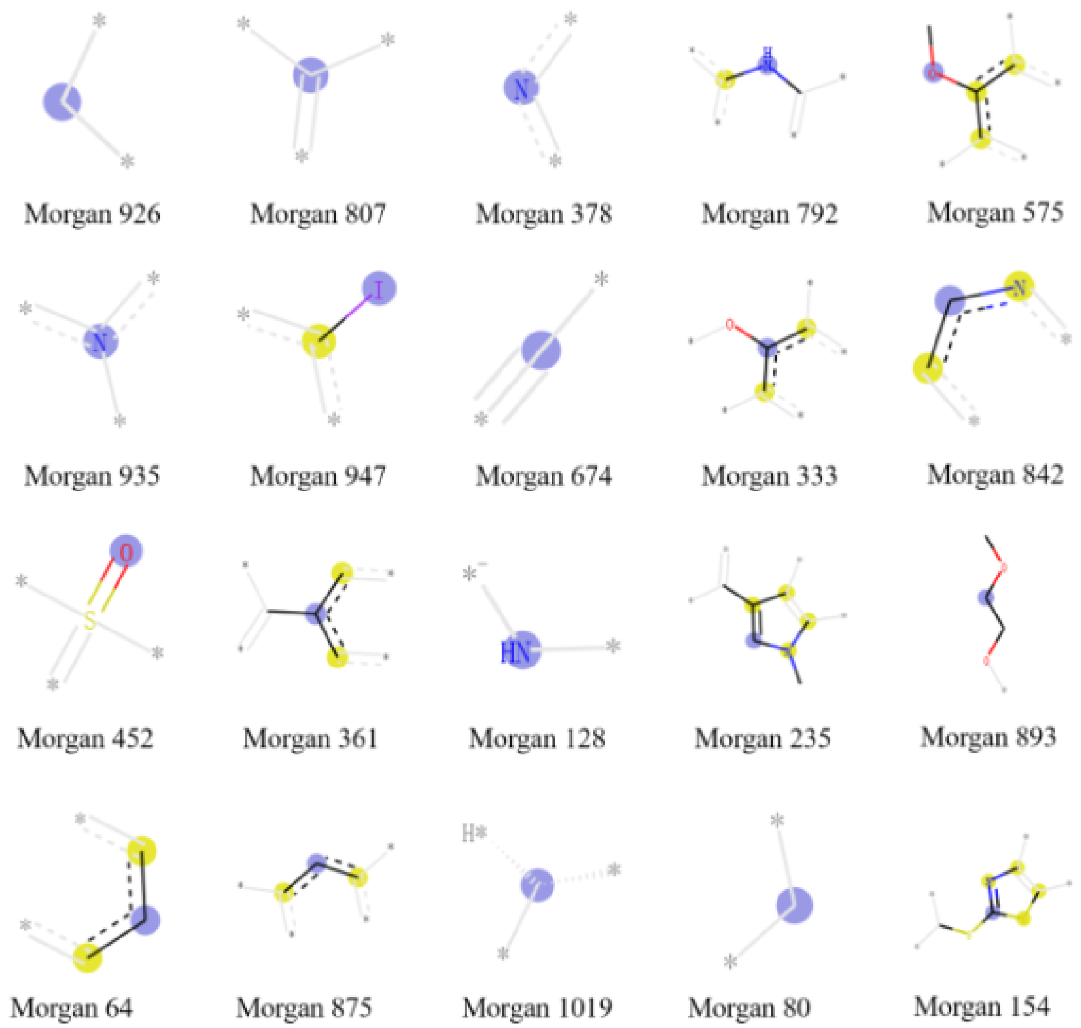


**Fig. S37.** Important molecular substructures of the RF::Morgan model in MDA-MB-468.

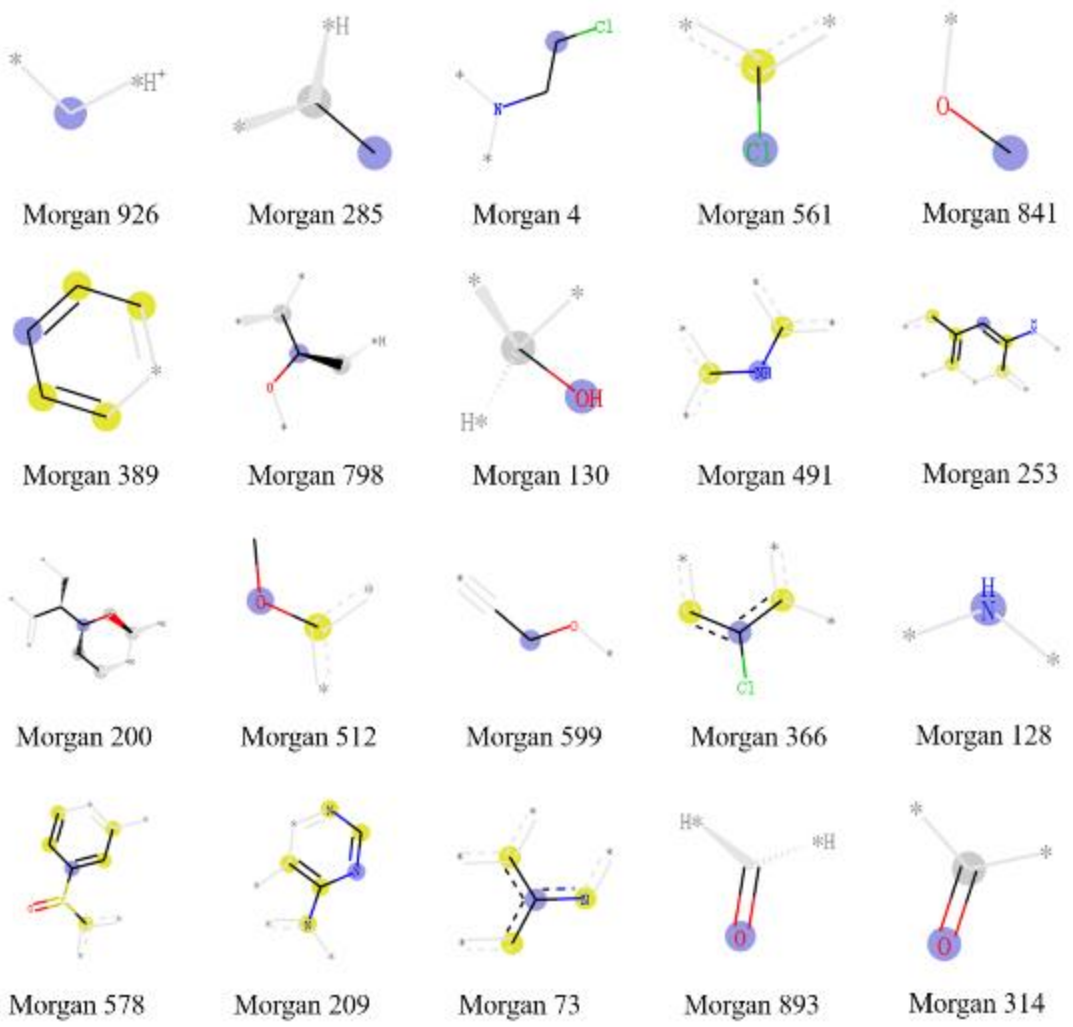


**Fig. S38.** Important molecular substructures of the RF::Morgan model in SK-BR-3.

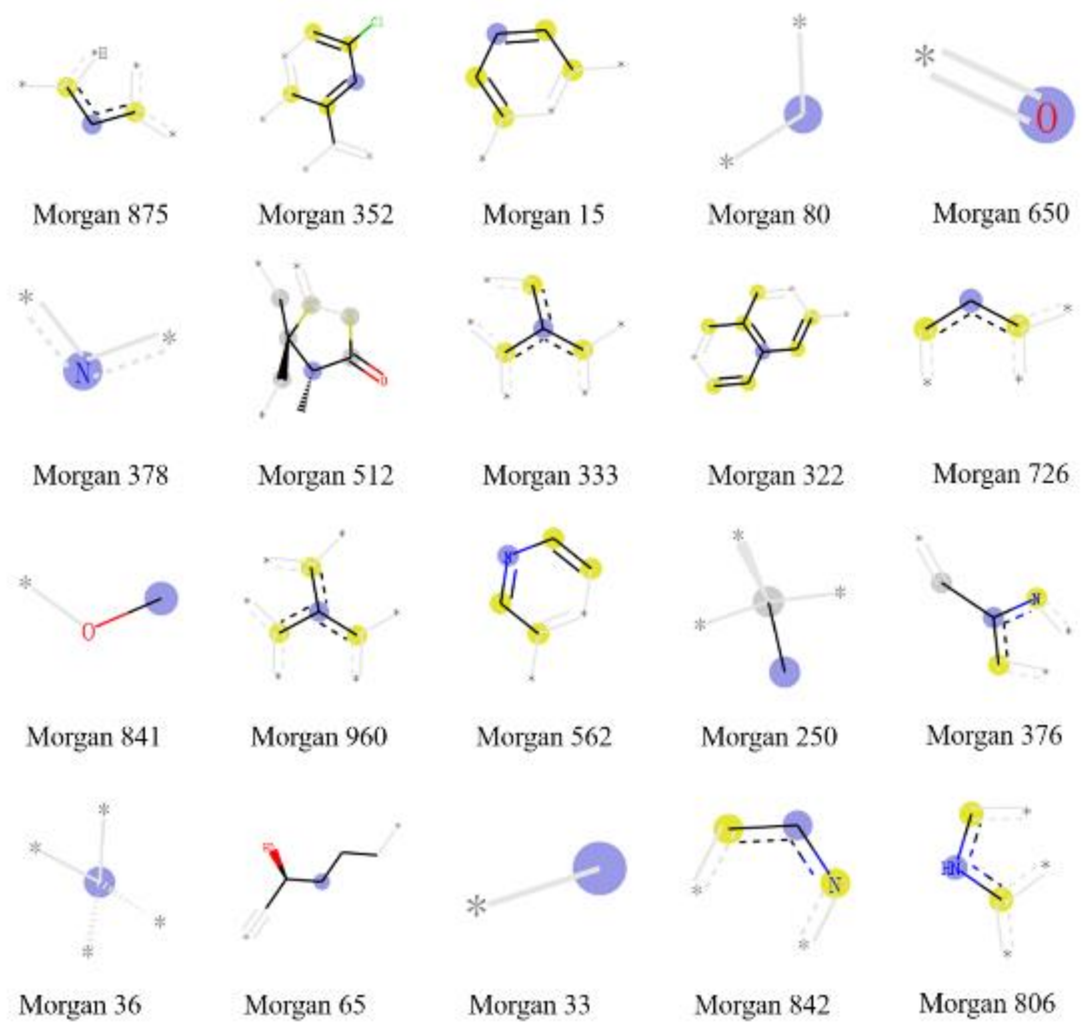


**Fig. S39.** Important molecular substructures of the RF::Morgan model in T-47D.

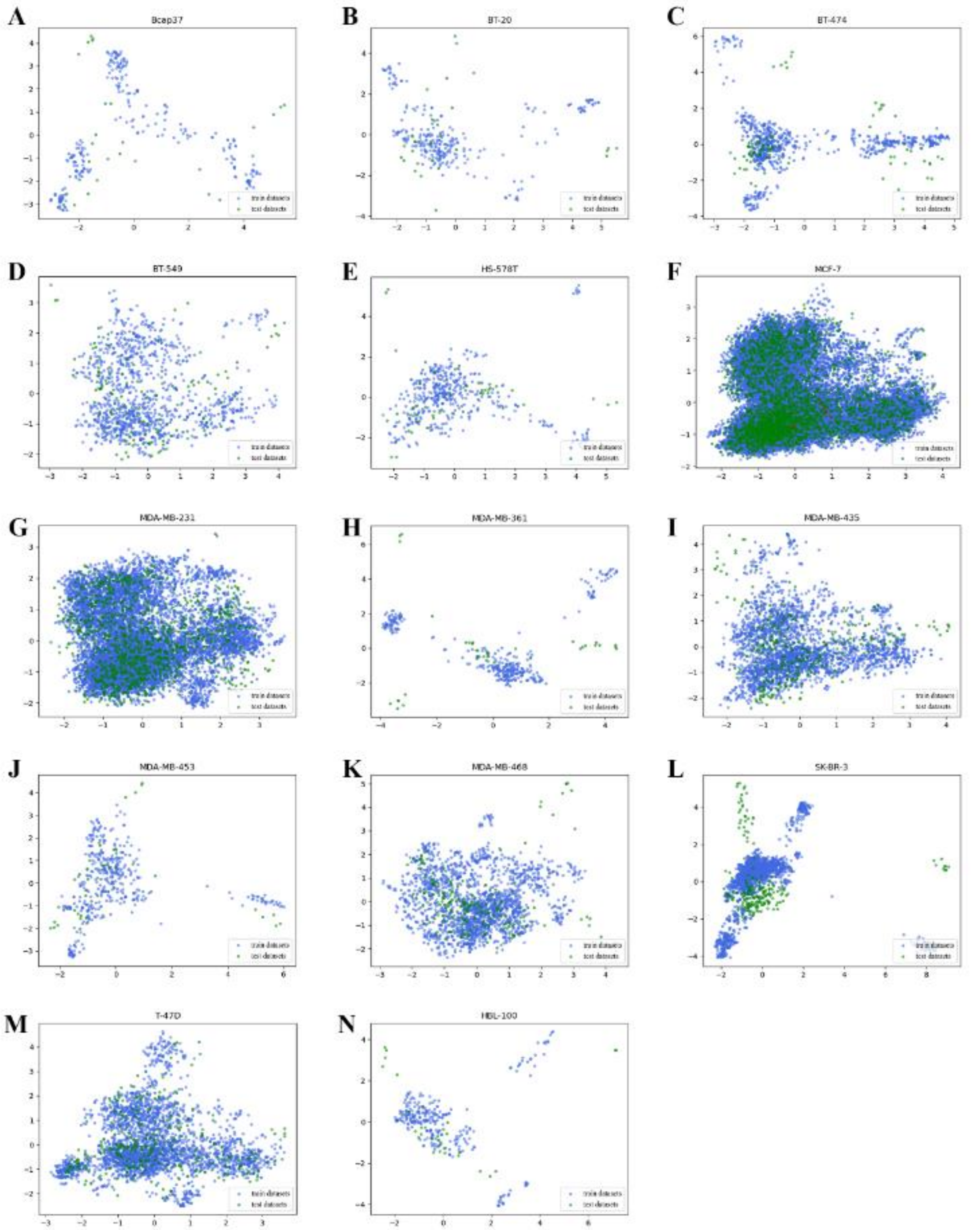


**Fig. S40.** Model AD in training sets and test sets in all breast cell lines. K was set to 5. By comparing the density of each point p and its five neighborhood points, whether this point is abnormal is judged. The lower the density of point p is, the more likely it is to be identified as an abnormal point. Exceptions are shown in red.
