## Supplemental Table for "Machine learning enables accurate and rapid prediction of active molecules against breast cancer cells"

**Table S1.** Summary of hyperparameter setting in different models.

| Methods | Hyperparameters |
|---|---|
| DNN | 'dropouts':[0.1,0.2,0.5],<br>'layer_sizes':[[64],[128],[256],[512]],<br>'weight_decay_penalty':[0.01,0.001,0.0001] |
| KNN | 'n_neighbors':[1,2,3,4,5],<br>'p':[1,2],<br>'weights':['uniform','distance'] |
| NB | 'alpha':[1,0.1,0.01],<br>'binarize':[0,0.5,0.8] |
| RF | 'n_estimators':[10, 50, 100, 200, 300, 400, 500],<br>'criterion':["gini", "entropy"],<br>'max_depth': [None,5,10,15],<br>'min_samples_leaf': [1,3,5,10],<br>'max_features':["log2","auto","sqrt"] |
| SVM | 'gamma':['auto',0.1,0.2],<br>'C': [1, 10, 100] |
| XGBoost | 'learning_rate':[0.01,0.1],<br>'gamma':[0,0.1],<br>'min_child_weight':[1,3],<br>'max_depth':[3,5],<br>'n_estimators':[50,100],<br>'subsample':[0.8,1.0],<br>'colsample_bytree':[0.8,1.0] |
| GCN | 'weight_decay':[0, 10e-8, 10e-6, 10e-4],<br>'graph_conv_layers': [[64, 64],[128, 128],[256,256]],<br>'learning_rate': [0.01,0.001, 0.0001],<br>'dense_layer_size': [64, 128, 256] |
| GAT | 'weight_decay':[0, 10e-8, 10e-6, 10e-4],<br>'learning_rate': [0.01,0.001, 0.0001],<br>'n_attention_heads':[8,16,32],<br>'dropout': [0,0.1,0.3,0.5] |
| MPNN | 'weight_decay': [10e-8, 10e-6, 10e-4],<br>'learning_rate': [0.01, 0.001, 0.0001],<br>'graph_conv_layers': [[64, 64], [128, 128],[256,256]],<br>'num_layer_set2set':[2,3,4],<br>'node_out_feats':[16,32,64],<br>'edge_hidden_feats':[16,32,64] |
| Attentive FP | 'dropout': [0,0.1,0.5],<br>'graph_feat_size': [50,100,200],<br>'num_timesteps':[1,2,3],<br>'num_layers':[2,3,4],<br>'learning_rate': [0.01,0.001, 0.0001],<br>'weight_decay':[0, 10e-4, 10e-2] |

| DNN | | |
|---|---|---|
| Methods | Datasets | Hyperparameters |
| DNN::AtomPairs | Bcap37 | default[a] |
| | BT-20 | 0.5, [128], 0.0001 |
| | BT-474 | 0.5, [64], 0.0001 |
| | BT-549 | default |
| | HBL-100 | 0.1, [512], 0.01 |
| | HS-578T | default |
| | MCF-7 | 0.2,[256],0.0001 |
| | MDA-MB-231 | 0.2,[64],0.001 |
| | MDA-MB-361 | 0.5, [128], 0.01 |
| | MDA-MB-435 | 0.2, [256], 0.001 |
| | MDA-MB-453 | default |
| | MDA-MB-468 | default |
| | SK-BR-3 | default |
| | T-47D | default |
| DNN::MACCS | Bcap37 | default |
| | BT-20 | default |
| | BT-474 | 0.2, [512], 0.001 |
| | BT-549 | 0.1, [512], 0.0001 |
| | HBL-100 | default |
| | HS-578T | default |
| | MCF-7 | 0.2, [256], 0.0001 |
| | MDA-MB-231 | 0.5,[256],0.0001 |
| | MDA-MB-361 | 0.2, [512], 0.001 |
| | MDA-MB-435 | 0.1, [128], 0.0001 |
| | MDA-MB-453 | 0.2, [512], 0.001 |
| | MDA-MB-468 | 0.5, [128], 0.001 |
| | SK-BR-3 | 0.2, [256], 0.001 |
| | T-47D | default |
| DNN::Morgan | Bcap37 | 0.5, [512], 0.001 |
| | BT-20 | 0.1, [512], 0.0001 |
| | BT-474 | default |
| | BT-549 | 0.5, [512], 0.01 |
| | HBL-100 | 0.5, [256], 0.001 |
| | HS-578T | 0.5, [128], 0.0001 |
| | MCF-7 | 0.5, [512], 0.0001 |
| | MDA-MB-231 | 0.5,[64],0.001 |
| | MDA-MB-361 | 0.1, [64], 0.0001 |
| | MDA-MB-435 | 0.5, [256], 0.0001 |
| | MDA-MB-453 | default |
| | MDA-MB-468 | default |
| | SK-BR-3 | 0.1, [512], 0.0001 |
| | T-47D | 0.5,[128],0.01 |
| DNN::PharmacoPFP | Bcap37 | default |
| | BT-20 | default |
| | BT-474 | 0.1, [512], 0.0001 |
| | BT-549 | 0.5, [64], 0.0001 |
| | HBL-100 | default |
| | HS-578T | default |
| | MCF-7 | 0.2,[128],0.0001 |
| | MDA-MB-231 | 0.2,[128],0.0001 |
| | MDA-MB-361 | default |
| | MDA-MB-435 | default |
| | MDA-MB-453 | 0.5, [64], 0.001 |
| | MDA-MB-468 | 0.5, [64], 0.01 |
| | SK-BR-3 | default |
| | T-47D | 0.5,[256],0.01 |
| DNN::rdkit | Bcap37 | 0.5, [64], 0.01 |
| | BT-20 | 0.5, [128], 0.001 |
| | BT-474 | default |
| | BT-549 | default |
| | HBL-100 | default |
| | HS-578T | default |
| | MCF-7 | 0.5, [128], 0.0001 |
| | MDA-MB-231 | 0.5,[64],0.0001 |
| | MDA-MB-361 | default |
| | MDA-MB-435 | default |
| | MDA-MB-453 | default |
| | MDA-MB-468 | default |
| | SK-BR-3 | 0.1, [128], 0.001 |
| | T-47D | default |

[*a]DNN default: weight_decay_penalty=0.001

| KNN | | |
|---|---|---|
| Methods | Datasets | Hyperparameters |
| KNN::AtomPairs | Bcap37 | 4, 1, 'distance' |
| | BT-20 | 5, 2, 'distance' |
| | BT-474 | 4, 1, 'distance' |
| | BT-549 | 3, 2, 'distance' |
| | HBL-100 | default[b] |
| | HS-578T | default |
| | MCF-7 | 5, 1, 'distance' |
| | MDA-MB-231 | 5, 1, 'distance' |
| | MDA-MB-361 | 4, 1, 'uniform' |
| | MDA-MB-435 | 5, 1, 'distance' |
| | MDA-MB-453 | default |
| | MDA-MB-468 | default |
| | SK-BR-3 | 5, 2, 'uniform' |
| | T-47D | 5, 1, 'distance' |
| KNN::MACCS | Bcap37 | 5, 1, 'distance' |
| | BT-20 | 4, 1, 'distance' |
| | BT-474 | default |
| | BT-549 | 4, 1, 'distance' |
| | HBL-100 | default |
| | HS-578T | 5, 1, 'distance' |
| | MCF-7 | 5, 2, 'uniform' |
| | MDA-MB-231 | 5, 2, 'uniform' |
| | MDA-MB-361 | default |
| | MDA-MB-435 | 5, 2, 'uniform' |
| | MDA-MB-453 | default |
| | MDA-MB-468 | 5, 1, 'distance' |
| | SK-BR-3 | 4, 2, 'distance' |
| | T-47D | default |
| KNN::Morgan | Bcap37 | 2, 2, 'distance' |
| | BT-20 | default |
| | BT-474 | default |
| | BT-549 | 5, 1, 'distance' |
| | HBL-100 | 4, 2, 'distance' |
| | HS-578T | 5, 1, 'distance' |
| | MCF-7 | 5, 1, 'distance' |
| | MDA-MB-231 | 5, 1, 'distance' |
| | MDA-MB-361 | 5, 2, 'distance' |
| | MDA-MB-435 | 5, 1, 'distance' |
| | MDA-MB-453 | 5, 2, 'uniform' |
| | MDA-MB-468 | 5, 2, 'distance' |
| | SK-BR-3 | 5, 2, 'distance' |
| | T-47D | 4, 1, 'distance' |
| KNN::PharmacoPFP | Bcap37 | 2, 2, 'distance' |
| | BT-20 | default |
| | BT-474 | 5, 2, 'distance' |
| | BT-549 | default |
| | HBL-100 | 3, 2, 'uniform' |
| | HS-578T | 4, 2, 'distance' |
| | MCF-7 | 5, 1, 'distance' |
| | MDA-MB-231 | 5, 1, 'distance' |
| | MDA-MB-361 | default |
| | MDA-MB-435 | 4, 2, 'uniform' |
| | MDA-MB-453 | default |
| | MDA-MB-468 | default |
| | SK-BR-3 | default |
| | T-47D | 5, 1, 'distance' |
| KNN::rdkit | Bcap37 | default |
| | BT-20 | 5, 2, 'uniform' |
| | BT-474 | default |
| | BT-549 | 3, 2, 'distance' |
| | HBL-100 | default |
| | HS-578T | default |
| | MCF-7 | 5, 1, 'distance' |
| | MDA-MB-231 | 5, 1, 'distance' |
| | MDA-MB-361 | default |
| | MDA-MB-435 | 5, 2, 'distance' |
| | MDA-MB-453 | default |
| | MDA-MB-468 | 4, 1, 'distance' |
| | SK-BR-3 | default |
| | T-47D | 5, 1, 'distance' |

[b]KNN default: n_neighbors=1

| NB | | |
|---|---|---|
| Methods | Datasets | Hyperparameters |
| NB::AtomPairs | Bcap37 | default[c] |
| | BT-20 | default |
| | BT-474 | default |
| | BT-549 | 0.01, 0.8 |
| | HBL-100 | 0.01, 0.8 |
| | HS-578T | default |
| | MCF-7 | 0.01, 0.8 |
| | MDA-MB-231 | 0.1, 0.8 |
| | MDA-MB-361 | default |
| | MDA-MB-435 | default |
| | MDA-MB-453 | default |
| | MDA-MB-468 | default |
| | SK-BR-3 | 0.01, 0.8 |
| | T-47D | 0.01, 0.8 |
| NB::MACCS | Bcap37 | 0.01, 0.8 |
| | BT-20 | 0.01, 0.8 |
| | BT-474 | default |
| | BT-549 | default |
| | HBL-100 | 0.01, 0.8 |
| | HS-578T | 0.01, 0.8 |
| | MCF-7 | 0.01, 0.8 |
| | MDA-MB-231 | 0.01, 0.8 |
| | MDA-MB-361 | default |
| | MDA-MB-435 | default |
| | MDA-MB-453 | 0.01, 0.8 |
| | MDA-MB-468 | default |
| | SK-BR-3 | 0.01, 0.8 |
| | T-47D | 0.1, 0.8 |
| NB::Morgan | Bcap37 | default |
| | BT-20 | default |
| | BT-474 | 0.1, 0.8 |
| | BT-549 | 0.01, 0.8 |
| | HBL-100 | 0.01, 0.8 |
| | HS-578T | 0.1, 0.8 |
| | MCF-7 | 0.01, 0.8 |
| | MDA-MB-231 | 0.1, 0.8 |
| | MDA-MB-361 | default |
| | MDA-MB-435 | default |
| | MDA-MB-453 | default |
| | MDA-MB-468 | default |
| | SK-BR-3 | 0.1, 0.8 |
| | T-47D | 0.1, 0.8 |
| NB::PharmacoPFP | Bcap37 | default |
| | BT-20 | default |
| | BT-474 | default |
| | BT-549 | default |
| | HBL-100 | default |
| | HS-578T | default |
| | MCF-7 | default |
| | MDA-MB-231 | default |
| | MDA-MB-361 | default |
| | MDA-MB-435 | default |
| | MDA-MB-453 | 0.01, 0.8 |
| | MDA-MB-468 | default |
| | SK-BR-3 | default |
| | T-47D | default |
| NB::rdkit | Bcap37 | default |
| | BT-20 | 0.01, 0 |
| | BT-474 | default |
| | BT-549 | 0.01, 0 |
| | HBL-100 | default |
| | HS-578T | default |
| | MCF-7 | default |
| | MDA-MB-231 | default |
| | MDA-MB-361 | default |
| | MDA-MB-435 | default |
| | MDA-MB-453 | default |
| | MDA-MB-468 | 0.01, 0 |
| | SK-BR-3 | 0.1, 0 |
| | T-47D | default |

[c]NB default: alpha=1

| RF | | |
|---|---|---|
| Methods | Datasets | Hyperparameter |
| RF::AtomPairs | Bcap37 | default[d] |
| | BT-20 | default |
| | BT-474 | default |
| | BT-549 | 200, 'gini', None, 1, 'sqrt' |
| | HBL-100 | 10, 'gini', None, 5, 'auto' |
| | HS-578T | default |
| | MCF-7 | 500, 'entropy', None, 1, 'sqrt' |
| | MDA-MB-231 | default |
| | MDA-MB-361 | 10, 'gini', None, 5, 'log2' |
| | MDA-MB-435 | default |
| | MDA-MB-453 | default |
| | MDA-MB-468 | 400, 'gini', None, 1, 'sqrt' |
| | SK-BR-3 | default |
| | T-47D | 50, 'entropy', None, 1, 'log2' |
| RF::MACCS | Bcap37 | 10, 'gini', 10, 1, 'log2' |
| | BT-20 | default |
| | BT-474 | 100, 'entropy', 15, 1, 'auto' |
| | BT-549 | default |
| | HBL-100 | default |
| | HS-578T | 10, 'gini', 15, 1, 'sqrt' |
| | MCF-7 | 500, 'entropy', None, 3, 'auto' |
| | MDA-MB-231 | default |
| | MDA-MB-361 | default |
| | MDA-MB-435 | default |
| | MDA-MB-453 | 100, 'gini', None, 1, 'auto' |
| | MDA-MB-468 | default |
| | SK-BR-3 | default |
| | T-47D | default |
| RF::Morgan | Bcap37 | 10, 'entropy', 5, 1, 'auto' |
| | BT-20 | 10, 'entropy', 10, 1, 'log2' |
| | BT-474 | default |
| | BT-549 | default |
| | HBL-100 | default |
| | HS-578T | default |
| | MCF-7 | 400, 'entropy', None, 1, 'sqrt' |
| | MDA-MB-231 | default |
| | MDA-MB-361 | default |
| | MDA-MB-435 | default |
| | MDA-MB-453 | default |
| | MDA-MB-468 | default |
| | SK-BR-3 | 100, 'gini', None, 1, 'log2' |
| | T-47D | default |
| RF::PharmacoPFP | Bcap37 | default |
| | BT-20 | default |
| | BT-474 | 100, 'entropy', 10, 1, 'log2' |
| | BT-549 | default |
| | HBL-100 | default |
| | HS-578T | 100, 'gini', 15, 1, 'auto' |
| | MCF-7 | default |
| | MDA-MB-231 | default |
| | MDA-MB-361 | default |
| | MDA-MB-435 | default |
| | MDA-MB-453 | default |
| | MDA-MB-468 | 100, 'gini', None, 1, 'auto' |
| | SK-BR-3 | default |
| | T-47D | default |
| RF::rdkit | Bcap37 | 100, 'entropy', 15, 1, 'sqrt' |
| | BT-20 | 10, 'gini', 10, 1, 'sqrt' |
| | BT-474 | default |
| | BT-549 | 100, 'entropy', 15, 1, 'log2' |
| | HBL-100 | default |
| | HS-578T | 10, 'entropy', 10, 5, 'sqrt' |
| | MCF-7 | 300, 'entropy', None, 3, 'auto' |
| | MDA-MB-231 | 500, 'entropy', None, 3, 'auto' |
| | MDA-MB-361 | default |
| | MDA-MB-435 | 200, 'entropy', None, 1, 'log2' |
| | MDA-MB-453 | default |
| | MDA-MB-468 | default |
| | SK-BR-3 | default |
| | T-47D | 100, 'entropy', None, 3, 'sqrt' |

[d]RF default: n_estimators=100;max_features='log2'

| SVM | | |
|---|---|---|
| Methods | Datasets | Hyperparameters |
| SVM::AtomPairs | Bcap37 | 'auto', 100 |
| | BT-20 | default[e] |
| | BT-474 | default |
| | BT-549 | default |
| | HBL-100 | 'auto', 100 |
| | HS-578T | 'auto', 100 |
| | MCF-7 | 'auto', 10 |
| | MDA-MB-231 | 'auto', 10 |
| | MDA-MB-361 | default |
| | MDA-MB-435 | default |
| | MDA-MB-453 | default |
| | MDA-MB-468 | 'auto', 100 |
| | SK-BR-3 | default |
| | T-47D | 'auto', 10 |
| SVM::MACCS | Bcap37 | 0.2, 100 |
| | BT-20 | default |
| | BT-474 | 'auto', 10 |
| | BT-549 | default |
| | HBL-100 | default |
| | HS-578T | default |
| | MCF-7 | 0.2, 1 |
| | MDA-MB-231 | 0.1, 1 |
| | MDA-MB-361 | default |
| | MDA-MB-435 | 0.1, 100 |
| | MDA-MB-453 | default |
| | MDA-MB-468 | 'auto', 100 |
| | SK-BR-3 | default |
| | T-47D | 'auto', 100 |
| SVM::Morgan | Bcap37 | 0.1, 100 |
| | BT-20 | default |
| | BT-474 | 'auto', 100 |
| | BT-549 | default |
| | HBL-100 | 0.1, 100 |
| | HS-578T | default |
| | MCF-7 | 0.1, 1 |
| | MDA-MB-231 | 0.1, 100 |
| | MDA-MB-361 | 'auto', 10 |
| | MDA-MB-435 | default |
| | MDA-MB-453 | default |
| | MDA-MB-468 | default |
| | SK-BR-3 | default |
| | T-47D | default |
| SVM::PharmacoPFP | Bcap37 | 0.2, 100 |
| | BT-20 | default |
| | BT-474 | default |
| | BT-549 | default |
| | HBL-100 | default |
| | HS-578T | default |
| | MCF-7 | 0.1, 1 |
| | MDA-MB-231 | 0.1, 10 |
| | MDA-MB-361 | default |
| | MDA-MB-435 | default |
| | MDA-MB-453 | 0.1, 100 |
| | MDA-MB-468 | default |
| | SK-BR-3 | default |
| | T-47D | default |
| SVM::rdkit | Bcap37 | 0.1, 100 |
| | BT-20 | default |
| | BT-474 | default |
| | BT-549 | 0.2, 100 |
| | HBL-100 | default |
| | HS-578T | 0.2, 100 |
| | MCF-7 | default |
| | MDA-MB-231 | default |
| | MDA-MB-361 | 0.2, 10 |
| | MDA-MB-435 | default |
| | MDA-MB-453 | default |
| | MDA-MB-468 | default |
| | SK-BR-3 | 0.2, 100 |
| | T-47D | 0.2, 100 |

[e]SVM default: C=1

| XGBoost | | |
|---|---|---|
| Methods | Datasets | Hyperparameters |
| XGBoost::AtomPairs | Bcap37 | default[f] |
| | BT-20 | default |
| | BT-474 | 0.1, 0.1, 3, 3, 100, 0.8, 1.0 |
| | BT-549 | default |
| | HBL-100 | default |
| | HS-578T | default |
| | MCF-7 | default |
| | MDA-MB-231 | default |
| | MDA-MB-361 | default |
| | MDA-MB-435 | default |
| | MDA-MB-453 | default |
| | MDA-MB-468 | default |
| | SK-BR-3 | 0.1, 0.1, 1, 5, 100, 1.0, 0.8 |
| | T-47D | 0.1, 0, 1, 5, 100, 0.8, 1.0 |
| XGBoost::MACCS | Bcap37 | default |
| | BT-20 | default |
| | BT-474 | default |
| | BT-549 | 0.1, 0.1, 1, 5, 100, 0.8, 1.0 |
| | HBL-100 | default |
| | HS-578T | 0.1, 0, 1, 5, 50, 0.8, 0.8 |
| | MCF-7 | default |
| | MDA-MB-231 | default |
| | MDA-MB-361 | default |
| | MDA-MB-435 | default |
| | MDA-MB-453 | 0.1, 0, 1, 3, 100, 1.0, 0.8 |
| | MDA-MB-468 | default |
| | SK-BR-3 | 0.1, 0, 1, 5, 100, 0.8, 1.0 |
| | T-47D | default |
| XGBoost::Morgan | Bcap37 | default |
| | BT-20 | default |
| | BT-474 | default |
| | BT-549 | default |
| | HBL-100 | default |
| | HS-578T | 0.1, 0.1, 1, 5, 50, 0.8, 0.8 |
| | MCF-7 | default |
| | MDA-MB-231 | default |
| | MDA-MB-361 | default |
| | MDA-MB-435 | default |
| | MDA-MB-453 | default |
| | MDA-MB-468 | default |
| | SK-BR-3 | default |
| | T-47D | default |
| XGBoost::PharmacoPFP | Bcap37 | default |
| | BT-20 | default |
| | BT-474 | default |
| | BT-549 | default |
| | HBL-100 | default |
| | HS-578T | 0.1, 0, 1, 3, 100, 0.8, 0.8 |
| | MCF-7 | default |
| | MDA-MB-231 | default |
| | MDA-MB-361 | default |
| | MDA-MB-435 | 0.1, 0, 1, 5, 100, 0.8, 1.0 |
| | MDA-MB-453 | default |
| | MDA-MB-468 | default |
| | SK-BR-3 | 0.1, 0, 1, 5, 100, 0.8, 1.0 |
| | T-47D | default |
| XGBoost::rdkit | Bcap37 | 0.1, 0.1, 1, 3, 50, 0.8, 1.0 |
| | BT-20 | 0.1, 0, 1, 5, 100, 0.8, 1.0 |
| | BT-474 | default |
| | BT-549 | default |
| | HBL-100 | default |
| | HS-578T | default |
| | MCF-7 | default |
| | MDA-MB-231 | 0.1, 0, 1, 5, 100, 0.8, 1.0 |
| | MDA-MB-361 | 0.1, 0.1, 3, 5, 50, 1.0, 1.0 |
| | MDA-MB-435 | default |
| | MDA-MB-453 | default |
| | MDA-MB-468 | 0.1, 0.1, 1, 5, 100, 0.8, 1.0 |
| | SK-BR-3 | default |
| | T-47D | default |

[f]XGBoost default: n_estimators=50

| Graph | | |
|---|---|---|
| Methods | Datasets | Hyperparameters |
| Graph::AttentiveFP | Bcap37 | default[g] |
| | BT-20 | default |
| | BT-474 | 0.1, 100, 1, 2, 0.01, 0.1 |
| | BT-549 | 0, 100, 3, 2, 0.01, 0.1 |
| | HBL-100 | 0.1, 50, 2, 4, 0.01, 0 |
| | HS-578T | 0.1, 100, 3, 2, 0.01, 0.1 |
| | MCF-7 | default |
| | MDA-MB-231 | default |
| | MDA-MB-361 | default |
| | MDA-MB-435 | 0, 200, 3, 2, 0.001, 0.1 |
| | MDA-MB-453 | default |
| | MDA-MB-468 | 0.1, 200, 2, 3, 0.001, 0.001 |
| | SK-BR-3 | default |
| | T-47D | default |
| Graph::GAT | Bcap37 | 0, 0.001, 32, 0.5 |
| | BT-20 | default[h] |
| | BT-474 | 1e-07, 0.01, 32, 0 |
| | BT-549 | default |
| | HBL-100 | default |
| | HS-578T | default |
| | MCF-7 | default |
| | MDA-MB-231 | 1e-07, 0.001, 16, 0 |
| | MDA-MB-361 | 1e-07, 0.01, 8, 0 |
| | MDA-MB-435 | 1e-05, 0.01, 8, 0 |
| | MDA-MB-453 | default |
| | MDA-MB-468 | default |
| | SK-BR-3 | 1e-07, 0.01, 8, 0 |
| | T-47D | 0, 0.01, 32, 0 |
| Graph::GCN | Bcap37 | 0.001, [256, 256], 0.0001, 128 |
| | BT-20 | 1e-07, [64, 64], 0.001, 256 |
| | BT-474 | default[i] |
| | BT-549 | default |
| | HBL-100 | 1e-07, [128, 128], 0.01, 256 |
| | HS-578T | 1e-07, [64, 64], 0.0001, 256 |
| | MCF-7 | 0, [128, 128], 0.0001, 256 |
| | MDA-MB-231 | 0, [256, 256], 0.0001, 256 |
| | MDA-MB-361 | 1e-05, [128, 128], 0.001, 64 |
| | MDA-MB-435 | default |
| | MDA-MB-453 | 0.001, [64, 64], 0.001, 256 |
| | MDA-MB-468 | default |
| | SK-BR-3 | default |
| | T-47D | default |
| Graph::MPNN | Bcap37 | default[j] |
| | BT-20 | 0.001, 0.001, [64, 64], 2, 32, 16 |
| | BT-474 | default |
| | BT-549 | default |
| | HBL-100 | 1e-05, 0.0001, [256, 256], 4, 32, 32 |
| | HS-578T | 1e-07, 0.0001, [256, 256], 3, 32, 16 |
| | MCF-7 | default |
| | MDA-MB-231 | default |
| | MDA-MB-361 | default |
| | MDA-MB-435 | 1e-07, 0.001, [256, 256], 2, 64, 32 |
| | MDA-MB-453 | 1e-05, 0.01, [64, 64], 2, 32, 32 |
| | MDA-MB-468 | 0.001, 0.001, [64, 64], 4, 64, 16 |
| | SK-BR-3 | 1e-07, 0.01, [64, 64], 2, 32, 64 |
| | T-47D | 1e-07, 0.001, [256, 256], 3, 64, 64 |

[g]AttentiveFP default: mode='classification';batch_size=32;learning_rate=0.001

[h]GAT default: mode='classification';batch_size=32;learning_rate=0.001

[i]GCN default: batch_size=64;mode='classification';learning_rate=0.001;dense_layer_size=128

[j]MPNN default: mode='classification';batch_size=32;learning_rate=0.001

**Table S2.** The performance results of models based on RDKit descriptors.

| Methods | Cell lines | Training set | | | | | | | Validation set | | | | | | | Test set | | | | | | |
|---|---|---|---|---|---|---|---|---|---|---|---|---|---|---|---|---|---|---|---|---|---|---|
| | | ACC[a] | F1[b] | BA[c] | SE[d] | SP[e] | MCC[f] | AUC[g] | ACC | F1 | BA | SE | SP | MCC | AUC | ACC | F1 | BA | SE | SP | MCC | AUC |
| DNN | Bcap37 | 0.786 | 0.719 | 0.769 | 0.682 | 0.856 | 0.549 | 0.848 | 0.714 | 0.600 | 0.684 | 0.545 | 0.824 | 0.386 | 0.695 | 0.778 | 0.727 | 0.770 | 0.727 | 0.813 | 0.540 | 0.807 |
| | BT-20 | 0.859 | 0.880 | 0.854 | 0.883 | 0.825 | 0.709 | 0.931 | 0.655 | 0.722 | 0.632 | 0.765 | 0.500 | 0.274 | 0.750 | 0.759 | 0.800 | 0.745 | 0.824 | 0.667 | 0.498 | 0.760 |
| | BT-474 | 0.878 | 0.925 | 0.779 | 0.951 | 0.606 | 0.610 | 0.924 | 0.827 | 0.891 | 0.739 | 0.891 | 0.588 | 0.479 | 0.892 | 0.778 | 0.866 | 0.600 | 0.906 | 0.294 | 0.238 | 0.806 |
| | BT-549 | 0.759 | 0.841 | 0.661 | 0.942 | 0.380 | 0.409 | 0.851 | 0.712 | 0.809 | 0.608 | 0.900 | 0.316 | 0.269 | 0.760 | 0.703 | 0.807 | 0.588 | 0.913 | 0.263 | 0.234 | 0.623 |
| | HBL-100 | 0.842 | 0.861 | 0.842 | 0.844 | 0.840 | 0.679 | 0.925 | 0.656 | 0.732 | 0.631 | 0.833 | 0.429 | 0.289 | 0.718 | 0.742 | 0.778 | 0.735 | 0.778 | 0.692 | 0.470 | 0.769 |
| | HS-578T | 0.723 | 0.839 | 0.500 | 1.000 | 0.000 | nan | 0.734 | 0.723 | 0.840 | 0.500 | 1.000 | 0.000 | nan | 0.794 | 0.723 | 0.840 | 0.500 | 1.000 | 0.000 | nan | 0.670 |
| | MCF-7 | 0.762 | 0.799 | 0.751 | 0.853 | 0.649 | 0.517 | 0.849 | 0.720 | 0.766 | 0.708 | 0.823 | 0.592 | 0.429 | 0.803 | 0.724 | 0.766 | 0.713 | 0.813 | 0.613 | 0.437 | 0.801 |
| | MDA-MB-231 | 0.780 | 0.843 | 0.724 | 0.930 | 0.518 | 0.510 | 0.867 | 0.748 | 0.820 | 0.690 | 0.902 | 0.478 | 0.430 | 0.818 | 0.730 | 0.806 | 0.674 | 0.880 | 0.468 | 0.388 | 0.809 |
| | MDA-MB-361 | 0.820 | 0.860 | 0.808 | 0.845 | 0.772 | 0.608 | 0.917 | 0.838 | 0.880 | 0.804 | 0.917 | 0.692 | 0.636 | 0.881 | 0.861 | 0.894 | 0.854 | 0.875 | 0.833 | 0.695 | 0.913 |
| | MDA-MB-435 | 0.809 | 0.887 | 0.608 | 0.967 | 0.249 | 0.329 | 0.827 | 0.782 | 0.874 | 0.545 | 0.970 | 0.119 | 0.172 | 0.735 | 0.802 | 0.884 | 0.590 | 0.970 | 0.209 | 0.293 | 0.807 |
| | MDA-MB-453 | 0.861 | 0.865 | 0.861 | 0.867 | 0.854 | 0.721 | 0.925 | 0.818 | 0.810 | 0.818 | 0.773 | 0.864 | 0.639 | 0.882 | 0.659 | 0.681 | 0.659 | 0.727 | 0.591 | 0.321 | 0.785 |
| | MDA-MB-468 | 0.822 | 0.893 | 0.651 | 0.971 | 0.332 | 0.428 | 0.862 | 0.844 | 0.905 | 0.707 | 0.967 | 0.447 | 0.522 | 0.858 | 0.813 | 0.886 | 0.658 | 0.947 | 0.370 | 0.403 | 0.830 |
| | SK-BR-3 | 0.821 | 0.888 | 0.694 | 0.947 | 0.442 | 0.470 | 0.849 | 0.793 | 0.872 | 0.647 | 0.941 | 0.353 | 0.375 | 0.827 | 0.792 | 0.869 | 0.667 | 0.914 | 0.420 | 0.386 | 0.818 |
| | T-47D | 0.776 | 0.855 | 0.664 | 0.942 | 0.386 | 0.414 | 0.826 | 0.745 | 0.836 | 0.623 | 0.927 | 0.319 | 0.319 | 0.767 | 0.744 | 0.832 | 0.638 | 0.900 | 0.376 | 0.327 | 0.750 |
| KNN | Bcap37 | 0.995 | 0.994 | 0.994 | 0.989 | 1.000 | 0.991 | 0.994 | 0.893 | 0.842 | 0.864 | 0.727 | 1.000 | 0.786 | 0.864 | 0.778 | 0.727 | 0.770 | 0.727 | 0.813 | 0.540 | 0.770 |
| | BT-20 | 0.855 | 0.875 | 0.852 | 0.869 | 0.835 | 0.702 | 0.919 | 0.897 | 0.909 | 0.900 | 0.882 | 0.917 | 0.791 | 0.946 | 0.724 | 0.789 | 0.691 | 0.882 | 0.500 | 0.421 | 0.792 |
| | BT-474 | 1.000 | 1.000 | 1.000 | 1.000 | 1.000 | 1.000 | 1.000 | 0.864 | 0.919 | 0.720 | 0.969 | 0.471 | 0.544 | 0.720 | 0.765 | 0.850 | 0.657 | 0.844 | 0.471 | 0.308 | 0.657 |
| | BT-549 | 0.998 | 0.998 | 0.998 | 0.997 | 1.000 | 0.995 | 1.000 | 0.797 | 0.855 | 0.746 | 0.888 | 0.605 | 0.518 | 0.830 | 0.669 | 0.761 | 0.611 | 0.775 | 0.447 | 0.227 | 0.642 |
| | HBL-100 | 0.996 | 0.997 | 0.995 | 1.000 | 0.991 | 0.992 | 0.995 | 0.781 | 0.811 | 0.774 | 0.833 | 0.714 | 0.553 | 0.774 | 0.806 | 0.842 | 0.791 | 0.889 | 0.692 | 0.599 | 0.791 |
| | HS-578T | 1.000 | 1.000 | 1.000 | 1.000 | 1.000 | 1.000 | 1.000 | 0.809 | 0.866 | 0.773 | 0.853 | 0.692 | 0.533 | 0.773 | 0.766 | 0.836 | 0.719 | 0.824 | 0.615 | 0.429 | 0.719 |
| | MCF-7 | 0.996 | 0.997 | 0.997 | 0.994 | 0.999 | 0.992 | 1.000 | 0.751 | 0.780 | 0.745 | 0.795 | 0.695 | 0.493 | 0.832 | 0.755 | 0.783 | 0.750 | 0.795 | 0.706 | 0.503 | 0.827 |
| | MDA-MB-231 | 0.998 | 0.999 | 0.998 | 0.998 | 0.999 | 0.996 | 1.000 | 0.779 | 0.832 | 0.751 | 0.854 | 0.648 | 0.514 | 0.837 | 0.779 | 0.828 | 0.757 | 0.836 | 0.677 | 0.518 | 0.844 |
| | MDA-MB-361 | 0.997 | 0.997 | 0.995 | 1.000 | 0.990 | 0.992 | 0.995 | 0.757 | 0.816 | 0.724 | 0.833 | 0.615 | 0.458 | 0.724 | 0.889 | 0.917 | 0.875 | 0.917 | 0.833 | 0.750 | 0.875 |
| | MDA-MB-435 | 0.996 | 0.998 | 0.997 | 0.996 | 0.998 | 0.989 | 1.000 | 0.835 | 0.899 | 0.702 | 0.941 | 0.463 | 0.471 | 0.862 | 0.822 | 0.890 | 0.688 | 0.928 | 0.448 | 0.431 | 0.813 |
| | MDA-MB-453 | 0.994 | 0.995 | 0.994 | 1.000 | 0.988 | 0.989 | 0.994 | 0.750 | 0.744 | 0.750 | 0.727 | 0.773 | 0.501 | 0.750 | 0.795 | 0.809 | 0.795 | 0.864 | 0.727 | 0.596 | 0.795 |
| | MDA-MB-468 | 0.999 | 1.000 | 1.000 | 0.999 | 1.000 | 0.998 | 1.000 | 0.869 | 0.918 | 0.775 | 0.954 | 0.596 | 0.613 | 0.907 | 0.823 | 0.885 | 0.748 | 0.888 | 0.609 | 0.501 | 0.836 |
| | SK-BR-3 | 0.999 | 0.999 | 0.998 | 0.999 | 0.998 | 0.997 | 0.998 | 0.823 | 0.884 | 0.745 | 0.901 | 0.588 | 0.511 | 0.745 | 0.802 | 0.868 | 0.734 | 0.868 | 0.600 | 0.468 | 0.734 |
| | T-47D | 0.998 | 0.999 | 0.999 | 0.997 | 1.000 | 0.995 | 1.000 | 0.764 | 0.838 | 0.695 | 0.868 | 0.521 | 0.413 | 0.808 | 0.783 | 0.850 | 0.721 | 0.873 | 0.570 | 0.462 | 0.804 |
| NB | Bcap37 | 0.645 | 0.602 | 0.650 | 0.670 | 0.629 | 0.293 | 0.734 | 0.679 | 0.609 | 0.671 | 0.636 | 0.706 | 0.338 | 0.711 | 0.704 | 0.714 | 0.736 | 0.909 | 0.563 | 0.480 | 0.821 |
| | BT-20 | 0.709 | 0.728 | 0.719 | 0.664 | 0.773 | 0.431 | 0.770 | 0.655 | 0.688 | 0.657 | 0.647 | 0.667 | 0.309 | 0.784 | 0.690 | 0.727 | 0.686 | 0.706 | 0.667 | 0.369 | 0.789 |
| | BT-474 | 0.775 | 0.855 | 0.689 | 0.838 | 0.540 | 0.360 | 0.781 | 0.790 | 0.862 | 0.738 | 0.828 | 0.647 | 0.435 | 0.835 | 0.716 | 0.819 | 0.583 | 0.813 | 0.353 | 0.162 | 0.683 |
| | BT-549 | 0.666 | 0.777 | 0.562 | 0.861 | 0.263 | 0.151 | 0.650 | 0.678 | 0.791 | 0.555 | 0.900 | 0.211 | 0.151 | 0.612 | 0.644 | 0.764 | 0.530 | 0.850 | 0.211 | 0.075 | 0.520 |
| | HBL-100 | 0.640 | 0.702 | 0.623 | 0.728 | 0.519 | 0.251 | 0.674 | 0.719 | 0.780 | 0.694 | 0.889 | 0.500 | 0.429 | 0.798 | 0.581 | 0.629 | 0.575 | 0.611 | 0.538 | 0.148 | 0.590 |
| | HS-578T | 0.683 | 0.789 | 0.573 | 0.819 | 0.327 | 0.158 | 0.695 | 0.745 | 0.833 | 0.633 | 0.882 | 0.385 | 0.303 | 0.686 | 0.702 | 0.800 | 0.604 | 0.824 | 0.385 | 0.220 | 0.695 |
| | MCF-7 | 0.569 | 0.631 | 0.557 | 0.663 | 0.450 | 0.116 | 0.584 | 0.584 | 0.642 | 0.574 | 0.671 | 0.477 | 0.150 | 0.600 | 0.569 | 0.636 | 0.555 | 0.679 | 0.432 | 0.113 | 0.581 |
| | MDA-MB-231 | 0.605 | 0.704 | 0.554 | 0.740 | 0.368 | 0.113 | 0.592 | 0.591 | 0.693 | 0.541 | 0.723 | 0.360 | 0.086 | 0.579 | 0.621 | 0.715 | 0.573 | 0.745 | 0.401 | 0.153 | 0.604 |
| | MDA-MB-361 | 0.759 | 0.819 | 0.724 | 0.834 | 0.614 | 0.456 | 0.827 | 0.676 | 0.778 | 0.591 | 0.875 | 0.308 | 0.223 | 0.768 | 0.750 | 0.800 | 0.750 | 0.750 | 0.750 | 0.478 | 0.894 |
| | MDA-MB-435 | 0.758 | 0.857 | 0.545 | 0.927 | 0.163 | 0.128 | 0.611 | 0.772 | 0.865 | 0.560 | 0.941 | 0.179 | 0.177 | 0.607 | 0.759 | 0.859 | 0.525 | 0.945 | 0.104 | 0.083 | 0.583 |
| | MDA-MB-453 | 0.688 | 0.718 | 0.685 | 0.773 | 0.596 | 0.376 | 0.796 | 0.705 | 0.698 | 0.705 | 0.682 | 0.727 | 0.410 | 0.769 | 0.705 | 0.723 | 0.705 | 0.773 | 0.636 | 0.413 | 0.764 |
| | MDA-MB-468 | 0.745 | 0.842 | 0.584 | 0.887 | 0.280 | 0.197 | 0.682 | 0.754 | 0.846 | 0.604 | 0.888 | 0.319 | 0.240 | 0.691 | 0.682 | 0.804 | 0.490 | 0.849 | 0.130 | -0.025 | 0.600 |
| | SK-BR-3 | 0.724 | 0.822 | 0.599 | 0.848 | 0.350 | 0.213 | 0.667 | 0.690 | 0.803 | 0.539 | 0.842 | 0.235 | 0.088 | 0.685 | 0.708 | 0.805 | 0.611 | 0.803 | 0.420 | 0.221 | 0.656 |
| | T-47D | 0.682 | 0.778 | 0.607 | 0.794 | 0.419 | 0.220 | 0.682 | 0.694 | 0.789 | 0.611 | 0.818 | 0.404 | 0.236 | 0.699 | 0.629 | 0.742 | 0.541 | 0.759 | 0.323 | 0.085 | 0.631 |
| RF | Bcap37 | 0.995 | 0.994 | 0.994 | 0.989 | 1.000 | 0.991 | 1.000 | 0.750 | 0.696 | 0.746 | 0.727 | 0.765 | 0.486 | 0.882 | 0.741 | 0.632 | 0.710 | 0.545 | 0.875 | 0.452 | 0.815 |
| | BT-20 | 0.987 | 0.989 | 0.988 | 0.985 | 0.990 | 0.974 | 0.999 | 0.897 | 0.909 | 0.900 | 0.882 | 0.917 | 0.791 | 0.975 | 0.828 | 0.848 | 0.828 | 0.824 | 0.833 | 0.651 | 0.797 |
| | BT-474 | 0.968 | 0.980 | 0.926 | 0.998 | 0.854 | 0.902 | 0.998 | 0.840 | 0.905 | 0.661 | 0.969 | 0.353 | 0.439 | 0.906 | 0.802 | 0.881 | 0.637 | 0.922 | 0.353 | 0.327 | 0.841 |
| | BT-549 | 0.998 | 0.998 | 0.997 | 1.000 | 0.994 | 0.995 | 1.000 | 0.763 | 0.839 | 0.680 | 0.913 | 0.447 | 0.418 | 0.801 | 0.729 | 0.818 | 0.634 | 0.900 | 0.368 | 0.322 | 0.663 |
| | HBL-100 | 0.992 | 0.993 | 0.992 | 0.993 | 0.991 | 0.984 | 1.000 | 0.688 | 0.737 | 0.675 | 0.778 | 0.571 | 0.358 | 0.817 | 0.839 | 0.865 | 0.829 | 0.889 | 0.769 | 0.667 | 0.870 |
| | HS-578T | 0.896 | 0.932 | 0.821 | 0.989 | 0.654 | 0.735 | 0.971 | 0.787 | 0.868 | 0.639 | 0.971 | 0.308 | 0.404 | 0.882 | 0.830 | 0.889 | 0.740 | 0.941 | 0.538 | 0.545 | 0.775 |
| | MCF-7 | 0.983 | 0.985 | 0.982 | 0.987 | 0.977 | 0.965 | 0.999 | 0.757 | 0.788 | 0.749 | 0.814 | 0.685 | 0.505 | 0.835 | 0.757 | 0.788 | 0.750 | 0.815 | 0.685 | 0.505 | 0.829 |
| | MDA-MB-231 | 0.982 | 0.986 | 0.978 | 0.991 | 0.966 | 0.961 | 0.999 | 0.781 | 0.841 | 0.732 | 0.910 | 0.554 | 0.510 | 0.852 | 0.782 | 0.839 | 0.742 | 0.888 | 0.596 | 0.514 | 0.861 |
| | MDA-MB-361 | 0.990 | 0.992 | 0.992 | 0.984 | 1.000 | 0.978 | 1.000 | 0.892 | 0.917 | 0.881 | 0.917 | 0.846 | 0.763 | 0.933 | 0.861 | 0.894 | 0.854 | 0.875 | 0.833 | 0.695 | 0.962 |
| | MDA-MB-435 | 0.995 | 0.997 | 0.994 | 0.996 | 0.993 | 0.987 | 1.000 | 0.825 | 0.895 | 0.658 | 0.958 | 0.358 | 0.415 | 0.841 | 0.812 | 0.887 | 0.639 | 0.949 | 0.328 | 0.365 | 0.822 |
| | MDA-MB-453 | 0.991 | 0.992 | 0.991 | 0.994 | 0.988 | 0.983 | 0.999 | 0.818 | 0.818 | 0.818 | 0.818 | 0.818 | 0.636 | 0.909 | 0.750 | 0.766 | 0.750 | 0.818 | 0.682 | 0.505 | 0.845 |
| | MDA-MB-468 | 0.923 | 0.952 | 0.837 | 0.998 | 0.677 | 0.780 | 0.993 | 0.854 | 0.911 | 0.714 | 0.980 | 0.447 | 0.557 | 0.890 | 0.843 | 0.906 | 0.678 | 0.987 | 0.370 | 0.511 | 0.890 |
| | SK-BR-3 | 0.921 | 0.950 | 0.842 | 0.999 | 0.685 | 0.785 | 0.987 | 0.818 | 0.890 | 0.650 | 0.987 | 0.314 | 0.459 | 0.827 | 0.822 | 0.892 | 0.667 | 0.974 | 0.360 | 0.462 | 0.873 |
| | T-47D | 0.970 | 0.979 | 0.952 | 0.995 | 0.909 | 0.928 | 0.998 | 0.783 | 0.858 | 0.684 | 0.932 | 0.436 | 0.440 | 0.854 | 0.789 | 0.861 | 0.692 | 0.932 | 0.452 | 0.454 | 0.841 |
| SVM | Bcap37 | 0.868 | 0.830 | 0.858 | 0.807 | 0.909 | 0.724 | 0.937 | 0.786 | 0.700 | 0.759 | 0.636 | 0.882 | 0.542 | 0.770 | 0.704 | 0.636 | 0.693 | 0.636 | 0.750 | 0.386 | 0.756 |
| | BT-20 | 0.838 | 0.859 | 0.836 | 0.847 | 0.825 | 0.668 | 0.900 | 0.759 | 0.759 | 0.782 | 0.647 | 0.917 | 0.564 | 0.833 | 0.759 | 0.800 | 0.745 | 0.824 | 0.667 | 0.498 | 0.804 |
| | BT-474 | 0.858 | 0.915 | 0.712 | 0.965 | 0.460 | 0.524 | 0.921 | 0.852 | 0.909 | 0.733 | 0.938 | 0.529 | 0.518 | 0.839 | 0.815 | 0.892 | 0.602 | 0.969 | 0.235 | 0.317 | 0.835 |
| | BT-549 | 0.756 | 0.840 | 0.650 | 0.953 | 0.347 | 0.400 | 0.835 | 0.746 | 0.833 | 0.640 | 0.938 | 0.342 | 0.363 | 0.703 | 0.729 | 0.822 | 0.620 | 0.925 | 0.316 | 0.313 | 0.646 |
| | HBL-100 | 0.755 | 0.793 | 0.744 | 0.810 | 0.679 | 0.493 | 0.875 | 0.750 | 0.778 | 0.746 | 0.778 | 0.714 | 0.492 | 0.815 | 0.677 | 0.737 | 0.658 | 0.778 | 0.538 | 0.326 | 0.756 |
| | HS-578T | 0.861 | 0.912 | 0.759 | 0.989 | 0.529 | 0.641 | 0.931 | 0.787 | 0.868 | 0.639 | 0.971 | 0.308 | 0.404 | 0.794 | 0.809 | 0.877 | 0.701 | 0.941 | 0.462 | 0.479 | 0.771 |
| | MCF-7 | 0.607 | 0.688 | 0.586 | 0.778 | 0.395 | 0.187 | 0.651 | 0.608 | 0.688 | 0.587 | 0.780 | 0.394 | 0.189 | 0.669 | 0.603 | 0.685 | 0.581 | 0.778 | 0.384 | 0.177 | 0.635 |
| | MDA-MB-231 | 0.678 | 0.784 | 0.587 | 0.920 | 0.254 | 0.238 | 0.695 | 0.692 | 0.792 | 0.605 | 0.922 | 0.288 | 0.279 | 0.679 | 0.649 | 0.759 | 0.566 | 0.868 | 0.264 | 0.165 | 0.678 |
| | MDA-MB-361 | 0.878 | 0.907 | 0.862 | 0.912 | 0.812 | 0.727 | 0.936 | 0.757 | 0.809 | 0.742 | 0.792 | 0.692 | 0.476 | 0.788 | 0.889 | 0.913 | 0.896 | 0.875 | 0.917 | 0.766 | 0.920 |
| | MDA-MB-435 | 0.810 | 0.889 | 0.598 | 0.978 | 0.219 | 0.329 | 0.838 | 0.772 | 0.868 | 0.533 | 0.962 | 0.104 | 0.123 | 0.724 | 0.812 | 0.889 | 0.612 | 0.970 | 0.254 | 0.344 | 0.782 |
| | MDA-MB-453 | 0.835 | 0.831 | 0.837 | 0.790 | 0.883 | 0.675 | 0.921 | 0.795 | 0.780 | 0.795 | 0.727 | 0.864 | 0.596 | 0.888 | 0.682 | 0.682 | 0.682 | 0.682 | 0.682 | 0.364 | 0.769 |
| | MDA-MB-468 | 0.824 | 0.894 | 0.660 | 0.969 | 0.350 | 0.439 | 0.833 | 0.849 | 0.909 | 0.696 | 0.987 | 0.404 | 0.541 | 0.834 | 0.793 | 0.878 | 0.592 | 0.967 | 0.217 | 0.294 | 0.767 |
| | SK-BR-3 | 0.809 | 0.885 | 0.640 | 0.977 | 0.303 | 0.417 | 0.862 | 0.837 | 0.902 | 0.676 | 1.000 | 0.353 | 0.539 | 0.847 | 0.797 | 0.878 | 0.617 | 0.974 | 0.260 | 0.363 | 0.829 |
| | T-47D | 0.778 | 0.857 | 0.664 | 0.947 | 0.382 | 0.420 | 0.824 | 0.768 | 0.851 | 0.648 | 0.945 | 0.351 | 0.388 | 0.760 | 0.732 | 0.825 | 0.617 | 0.900 | 0.333 | 0.284 | 0.709 |
| XGBoost | Bcap37 | 0.950 | 0.936 | 0.945 | 0.920 | 0.970 | 0.896 | 0.990 | 0.643 | 0.545 | 0.626 | 0.545 | 0.706 | 0.251 | 0.733 | 0.704 | 0.600 | 0.679 | 0.545 | 0.813 | 0.373 | 0.869 |
| | BT-20 | 0.987 | 0.989 | 0.986 | 0.993 | 0.979 | 0.974 | 1.000 | 0.897 | 0.909 | 0.900 | 0.882 | 0.917 | 0.791 | 0.946 | 0.793 | 0.824 | 0.787 | 0.824 | 0.750 | 0.574 | 0.848 |
| | BT-474 | 0.991 | 0.994 | 0.983 | 0.996 | 0.971 | 0.972 | 1.000 | 0.840 | 0.902 | 0.704 | 0.938 | 0.471 | 0.468 | 0.878 | 0.840 | 0.901 | 0.726 | 0.922 | 0.529 | 0.486 | 0.834 |
| | BT-549 | 0.995 | 0.996 | 0.993 | 0.998 | 0.987 | 0.988 | 1.000 | 0.754 | 0.834 | 0.667 | 0.913 | 0.421 | 0.393 | 0.812 | 0.712 | 0.800 | 0.636 | 0.850 | 0.421 | 0.298 | 0.696 |
| | HBL-100 | 0.996 | 0.997 | 0.995 | 1.000 | 0.991 | 0.992 | 1.000 | 0.688 | 0.750 | 0.667 | 0.833 | 0.500 | 0.357 | 0.734 | 0.839 | 0.865 | 0.829 | 0.889 | 0.769 | 0.667 | 0.850 |
| | HS-578T | 0.979 | 0.985 | 0.965 | 0.996 | 0.933 | 0.947 | 0.999 | 0.851 | 0.901 | 0.778 | 0.941 | 0.615 | 0.608 | 0.914 | 0.809 | 0.873 | 0.725 | 0.912 | 0.538 | 0.492 | 0.837 |
| | MCF-7 | 0.832 | 0.854 | 0.826 | 0.880 | 0.772 | 0.659 | 0.914 | 0.724 | 0.761 | 0.716 | 0.791 | 0.641 | 0.438 | 0.786 | 0.708 | 0.748 | 0.699 | 0.780 | 0.619 | 0.405 | 0.777 |
| | MDA-MB-231 | 0.837 | 0.881 | 0.793 | 0.953 | 0.632 | 0.642 | 0.922 | 0.750 | 0.823 | 0.689 | 0.912 | 0.466 | 0.434 | 0.813 | 0.746 | 0.815 | 0.697 | 0.877 | 0.517 | 0.429 | 0.815 |
| | MDA-MB-361 | 0.973 | 0.979 | 0.975 | 0.969 | 0.980 | 0.941 | 0.995 | 0.838 | 0.880 | 0.804 | 0.917 | 0.692 | 0.636 | 0.936 | 0.889 | 0.913 | 0.896 | 0.875 | 0.917 | 0.766 | 0.965 |
| | MDA-MB-435 | 0.970 | 0.981 | 0.943 | 0.992 | 0.893 | 0.912 | 0.997 | 0.802 | 0.879 | 0.648 | 0.924 | 0.373 | 0.353 | 0.801 | 0.812 | 0.885 | 0.665 | 0.928 | 0.403 | 0.390 | 0.835 |
| | MDA-MB-453 | 0.977 | 0.978 | 0.977 | 0.972 | 0.982 | 0.955 | 0.998 | 0.773 | 0.783 | 0.773 | 0.818 | 0.727 | 0.548 | 0.903 | 0.795 | 0.809 | 0.795 | 0.864 | 0.727 | 0.596 | 0.829 |
| | MDA-MB-468 | 0.976 | 0.985 | 0.954 | 0.995 | 0.914 | 0.933 | 0.998 | 0.884 | 0.928 | 0.785 | 0.974 | 0.596 | 0.658 | 0.911 | 0.864 | 0.914 | 0.767 | 0.947 | 0.587 | 0.592 | 0.886 |
| | SK-BR-3 | 0.995 | 0.997 | 0.993 | 0.998 | 0.988 | 0.987 | 1.000 | 0.833 | 0.893 | 0.732 | 0.934 | 0.529 | 0.521 | 0.856 | 0.842 | 0.897 | 0.761 | 0.921 | 0.600 | 0.554 | 0.887 |
| | T-47D | 0.932 | 0.953 | 0.897 | 0.983 | 0.812 | 0.835 | 0.981 | 0.803 | 0.870 | 0.710 | 0.941 | 0.479 | 0.495 | 0.819 | 0.783 | 0.855 | 0.696 | 0.909 | 0.484 | 0.443 | 0.820 |

[a] ACC: Accuracy. [b] F1: F1-measure. [c] BA: Balanced accuracy. [d] SE:Sensitivity. [e] SP:Specificity. [f] MCC:Matthews correlation coefficient. [g] AUC:The area under receiver operating characteristic.

**Table S3.** The performance results of models based on AtomPairs fingerprints.

| Methods | Cell lines | Training set | | | | | | | Validation set | | | | | | | Test set | | | | | | |
|---|---|---|---|---|---|---|---|---|---|---|---|---|---|---|---|---|---|---|---|---|---|---|
| | | ACC[a] | F1[b] | BA[c] | SE[d] | SP[e] | MCC[f] | AUC[g] | ACC | F1 | BA | SE | SP | MCC | AUC | ACC | F1 | BA | SE | SP | MCC | AUC |
| DNN | Bcap37 | 0.905 | 0.890 | 0.915 | 0.966 | 0.864 | 0.814 | 0.970 | 0.786 | 0.750 | 0.791 | 0.818 | 0.765 | 0.571 | 0.893 | 0.815 | 0.783 | 0.815 | 0.818 | 0.813 | 0.624 | 0.801 |
| | BT-20 | 0.996 | 0.996 | 0.996 | 0.993 | 1.000 | 0.991 | 1.000 | 0.724 | 0.750 | 0.728 | 0.706 | 0.750 | 0.449 | 0.917 | 0.759 | 0.811 | 0.733 | 0.882 | 0.583 | 0.496 | 0.853 |
| | BT-474 | 0.972 | 0.983 | 0.950 | 0.988 | 0.912 | 0.916 | 0.991 | 0.877 | 0.924 | 0.771 | 0.953 | 0.588 | 0.601 | 0.901 | 0.827 | 0.892 | 0.718 | 0.906 | 0.529 | 0.457 | 0.845 |
| | BT-549 | 0.994 | 0.995 | 0.991 | 0.998 | 0.984 | 0.986 | 1.000 | 0.797 | 0.859 | 0.733 | 0.913 | 0.553 | 0.511 | 0.826 | 0.737 | 0.814 | 0.675 | 0.850 | 0.500 | 0.372 | 0.653 |
| | HBL-100 | 0.988 | 0.990 | 0.987 | 0.993 | 0.981 | 0.976 | 0.999 | 0.719 | 0.757 | 0.710 | 0.778 | 0.643 | 0.425 | 0.869 | 0.742 | 0.778 | 0.735 | 0.778 | 0.692 | 0.470 | 0.825 |
| | HS-578T | 1.000 | 1.000 | 1.000 | 1.000 | 1.000 | 1.000 | 1.000 | 0.894 | 0.930 | 0.831 | 0.971 | 0.692 | 0.725 | 0.889 | 0.809 | 0.870 | 0.749 | 0.882 | 0.615 | 0.511 | 0.814 |
| | MCF-7 | 0.966 | 0.970 | 0.965 | 0.980 | 0.950 | 0.932 | 0.995 | 0.780 | 0.806 | 0.776 | 0.820 | 0.731 | 0.554 | 0.859 | 0.782 | 0.807 | 0.778 | 0.818 | 0.738 | 0.558 | 0.853 |
| | MDA-MB-231 | 0.903 | 0.927 | 0.879 | 0.967 | 0.792 | 0.790 | 0.972 | 0.789 | 0.843 | 0.752 | 0.888 | 0.616 | 0.531 | 0.864 | 0.796 | 0.848 | 0.758 | 0.894 | 0.623 | 0.545 | 0.855 |
| | MDA-MB-361 | 0.986 | 0.990 | 0.987 | 0.984 | 0.990 | 0.970 | 1.000 | 0.784 | 0.840 | 0.745 | 0.875 | 0.615 | 0.512 | 0.926 | 0.889 | 0.913 | 0.896 | 0.875 | 0.917 | 0.766 | 0.931 |
| | MDA-MB-435 | 0.922 | 0.951 | 0.858 | 0.972 | 0.744 | 0.763 | 0.972 | 0.828 | 0.895 | 0.692 | 0.936 | 0.448 | 0.448 | 0.855 | 0.835 | 0.900 | 0.691 | 0.949 | 0.433 | 0.463 | 0.864 |
| | MDA-MB-453 | 0.989 | 0.989 | 0.988 | 0.994 | 0.982 | 0.977 | 1.000 | 0.727 | 0.769 | 0.727 | 0.909 | 0.545 | 0.488 | 0.889 | 0.795 | 0.824 | 0.795 | 0.955 | 0.636 | 0.623 | 0.876 |
| | MDA-MB-468 | 0.985 | 0.990 | 0.978 | 0.991 | 0.965 | 0.958 | 0.999 | 0.879 | 0.924 | 0.789 | 0.961 | 0.617 | 0.644 | 0.903 | 0.879 | 0.923 | 0.807 | 0.941 | 0.674 | 0.647 | 0.891 |
| | SK-BR-3 | 0.948 | 0.967 | 0.896 | 1.000 | 0.792 | 0.861 | 0.994 | 0.852 | 0.909 | 0.725 | 0.980 | 0.471 | 0.576 | 0.896 | 0.842 | 0.901 | 0.727 | 0.954 | 0.500 | 0.537 | 0.860 |
| | T-47D | 0.940 | 0.958 | 0.909 | 0.986 | 0.832 | 0.855 | 0.988 | 0.761 | 0.840 | 0.671 | 0.895 | 0.447 | 0.387 | 0.817 | 0.831 | 0.887 | 0.752 | 0.945 | 0.559 | 0.572 | 0.866 |
| KNN | Bcap37 | 0.995 | 0.994 | 0.994 | 0.989 | 1.000 | 0.991 | 1.000 | 0.750 | 0.696 | 0.746 | 0.727 | 0.765 | 0.486 | 0.837 | 0.852 | 0.800 | 0.832 | 0.727 | 0.938 | 0.693 | 0.869 |
| | BT-20 | 1.000 | 1.000 | 1.000 | 1.000 | 1.000 | 1.000 | 1.000 | 0.897 | 0.909 | 0.900 | 0.882 | 0.917 | 0.791 | 0.971 | 0.828 | 0.848 | 0.828 | 0.824 | 0.833 | 0.651 | 0.863 |
| | BT-474 | 1.000 | 1.000 | 1.000 | 1.000 | 1.000 | 1.000 | 1.000 | 0.864 | 0.915 | 0.784 | 0.922 | 0.647 | 0.582 | 0.870 | 0.827 | 0.891 | 0.739 | 0.891 | 0.588 | 0.479 | 0.792 |
| | BT-549 | 0.998 | 0.998 | 0.998 | 0.997 | 1.000 | 0.995 | 1.000 | 0.856 | 0.897 | 0.818 | 0.925 | 0.711 | 0.662 | 0.850 | 0.720 | 0.792 | 0.683 | 0.788 | 0.579 | 0.364 | 0.689 |
| | HBL-100 | 0.996 | 0.997 | 0.995 | 1.000 | 0.991 | 0.992 | 0.995 | 0.813 | 0.842 | 0.802 | 0.889 | 0.714 | 0.618 | 0.802 | 0.806 | 0.842 | 0.791 | 0.889 | 0.692 | 0.599 | 0.791 |
| | HS-578T | 1.000 | 1.000 | 1.000 | 1.000 | 1.000 | 1.000 | 1.000 | 0.851 | 0.899 | 0.802 | 0.912 | 0.692 | 0.620 | 0.802 | 0.766 | 0.831 | 0.743 | 0.794 | 0.692 | 0.459 | 0.743 |
| | MCF-7 | 0.995 | 0.995 | 0.995 | 0.991 | 0.999 | 0.989 | 1.000 | 0.794 | 0.814 | 0.791 | 0.813 | 0.770 | 0.583 | 0.875 | 0.799 | 0.818 | 0.797 | 0.813 | 0.782 | 0.594 | 0.871 |
| | MDA-MB-231 | 0.997 | 0.998 | 0.998 | 0.996 | 0.999 | 0.994 | 1.000 | 0.833 | 0.869 | 0.819 | 0.871 | 0.766 | 0.638 | 0.879 | 0.820 | 0.858 | 0.807 | 0.853 | 0.761 | 0.611 | 0.886 |
| | MDA-MB-361 | 0.850 | 0.876 | 0.869 | 0.808 | 0.931 | 0.706 | 0.940 | 0.946 | 0.957 | 0.958 | 0.917 | 1.000 | 0.891 | 0.947 | 0.861 | 0.889 | 0.875 | 0.833 | 0.917 | 0.717 | 0.938 |
| | MDA-MB-435 | 0.996 | 0.998 | 0.998 | 0.995 | 1.000 | 0.989 | 1.000 | 0.832 | 0.896 | 0.705 | 0.932 | 0.478 | 0.466 | 0.825 | 0.848 | 0.904 | 0.758 | 0.919 | 0.597 | 0.541 | 0.870 |
| | MDA-MB-453 | 0.994 | 0.995 | 0.994 | 1.000 | 0.988 | 0.989 | 0.994 | 0.750 | 0.766 | 0.750 | 0.818 | 0.682 | 0.505 | 0.750 | 0.795 | 0.809 | 0.795 | 0.864 | 0.727 | 0.596 | 0.795 |
| | MDA-MB-468 | 0.999 | 1.000 | 1.000 | 0.999 | 1.000 | 0.998 | 1.000 | 0.834 | 0.893 | 0.759 | 0.901 | 0.617 | 0.531 | 0.759 | 0.843 | 0.895 | 0.815 | 0.868 | 0.761 | 0.593 | 0.815 |
| | SK-BR-3 | 0.863 | 0.912 | 0.785 | 0.940 | 0.630 | 0.614 | 0.931 | 0.852 | 0.906 | 0.758 | 0.947 | 0.569 | 0.580 | 0.876 | 0.812 | 0.879 | 0.714 | 0.908 | 0.520 | 0.463 | 0.831 |
| | T-47D | 0.998 | 0.999 | 0.999 | 0.997 | 1.000 | 0.995 | 1.000 | 0.783 | 0.850 | 0.721 | 0.877 | 0.564 | 0.464 | 0.851 | 0.802 | 0.860 | 0.760 | 0.864 | 0.656 | 0.523 | 0.841 |
| NB | Bcap37 | 0.736 | 0.613 | 0.701 | 0.523 | 0.879 | 0.437 | 0.830 | 0.643 | 0.444 | 0.594 | 0.364 | 0.824 | 0.211 | 0.733 | 0.667 | 0.571 | 0.648 | 0.545 | 0.750 | 0.301 | 0.756 |
| | BT-20 | 0.778 | 0.780 | 0.800 | 0.672 | 0.928 | 0.598 | 0.877 | 0.724 | 0.692 | 0.765 | 0.529 | 1.000 | 0.564 | 0.824 | 0.724 | 0.750 | 0.728 | 0.706 | 0.750 | 0.449 | 0.789 |
| | BT-474 | 0.749 | 0.819 | 0.785 | 0.723 | 0.847 | 0.475 | 0.860 | 0.679 | 0.764 | 0.710 | 0.656 | 0.765 | 0.346 | 0.805 | 0.741 | 0.817 | 0.750 | 0.734 | 0.765 | 0.421 | 0.767 |
| | BT-549 | 0.654 | 0.696 | 0.691 | 0.586 | 0.795 | 0.359 | 0.765 | 0.619 | 0.676 | 0.636 | 0.588 | 0.684 | 0.254 | 0.621 | 0.542 | 0.585 | 0.580 | 0.475 | 0.684 | 0.151 | 0.623 |
| | HBL-100 | 0.775 | 0.767 | 0.801 | 0.639 | 0.962 | 0.609 | 0.917 | 0.531 | 0.571 | 0.528 | 0.556 | 0.500 | 0.055 | 0.671 | 0.645 | 0.645 | 0.662 | 0.556 | 0.769 | 0.325 | 0.761 |
| | HS-578T | 0.731 | 0.791 | 0.751 | 0.705 | 0.798 | 0.454 | 0.853 | 0.745 | 0.813 | 0.729 | 0.765 | 0.692 | 0.425 | 0.750 | 0.596 | 0.678 | 0.602 | 0.588 | 0.615 | 0.183 | 0.688 |
| | MCF-7 | 0.553 | 0.522 | 0.567 | 0.440 | 0.694 | 0.137 | 0.601 | 0.566 | 0.545 | 0.578 | 0.468 | 0.688 | 0.159 | 0.620 | 0.552 | 0.516 | 0.567 | 0.430 | 0.705 | 0.138 | 0.599 |
| | MDA-MB-231 | 0.549 | 0.559 | 0.586 | 0.450 | 0.722 | 0.170 | 0.635 | 0.566 | 0.579 | 0.603 | 0.468 | 0.739 | 0.203 | 0.639 | 0.545 | 0.553 | 0.584 | 0.441 | 0.727 | 0.166 | 0.627 |
| | MDA-MB-361 | 0.806 | 0.831 | 0.843 | 0.725 | 0.960 | 0.651 | 0.904 | 0.784 | 0.818 | 0.798 | 0.750 | 0.846 | 0.571 | 0.849 | 0.833 | 0.857 | 0.875 | 0.750 | 1.000 | 0.707 | 0.931 |
| | MDA-MB-435 | 0.581 | 0.662 | 0.652 | 0.526 | 0.778 | 0.252 | 0.735 | 0.554 | 0.640 | 0.612 | 0.508 | 0.716 | 0.187 | 0.670 | 0.587 | 0.675 | 0.634 | 0.551 | 0.716 | 0.222 | 0.720 |
| | MDA-MB-453 | 0.795 | 0.782 | 0.798 | 0.713 | 0.883 | 0.603 | 0.863 | 0.727 | 0.700 | 0.727 | 0.636 | 0.818 | 0.462 | 0.818 | 0.727 | 0.684 | 0.727 | 0.591 | 0.864 | 0.472 | 0.803 |
| | MDA-MB-468 | 0.717 | 0.791 | 0.736 | 0.700 | 0.771 | 0.405 | 0.808 | 0.678 | 0.763 | 0.679 | 0.678 | 0.681 | 0.310 | 0.734 | 0.652 | 0.738 | 0.667 | 0.638 | 0.696 | 0.284 | 0.759 |
| | SK-BR-3 | 0.608 | 0.672 | 0.681 | 0.535 | 0.826 | 0.314 | 0.739 | 0.567 | 0.645 | 0.606 | 0.526 | 0.686 | 0.185 | 0.668 | 0.614 | 0.678 | 0.690 | 0.539 | 0.840 | 0.330 | 0.724 |
| | T-47D | 0.725 | 0.795 | 0.703 | 0.758 | 0.649 | 0.387 | 0.777 | 0.723 | 0.795 | 0.693 | 0.768 | 0.617 | 0.371 | 0.749 | 0.668 | 0.748 | 0.646 | 0.700 | 0.591 | 0.273 | 0.696 |
| RF | Bcap37 | 0.995 | 0.994 | 0.996 | 1.000 | 0.992 | 0.991 | 1.000 | 0.786 | 0.750 | 0.791 | 0.818 | 0.765 | 0.571 | 0.869 | 0.741 | 0.667 | 0.724 | 0.636 | 0.813 | 0.457 | 0.776 |
| | BT-20 | 1.000 | 1.000 | 1.000 | 1.000 | 1.000 | 1.000 | 1.000 | 0.793 | 0.813 | 0.799 | 0.765 | 0.833 | 0.589 | 0.919 | 0.759 | 0.811 | 0.733 | 0.882 | 0.583 | 0.496 | 0.860 |
| | BT-474 | 1.000 | 1.000 | 1.000 | 1.000 | 1.000 | 1.000 | 1.000 | 0.852 | 0.912 | 0.690 | 0.969 | 0.412 | 0.493 | 0.918 | 0.827 | 0.897 | 0.653 | 0.953 | 0.353 | 0.397 | 0.846 |
| | BT-549 | 0.998 | 0.998 | 0.998 | 0.998 | 0.997 | 0.995 | 1.000 | 0.822 | 0.877 | 0.758 | 0.938 | 0.579 | 0.574 | 0.878 | 0.788 | 0.855 | 0.713 | 0.925 | 0.500 | 0.486 | 0.713 |
| | HBL-100 | 0.917 | 0.929 | 0.913 | 0.939 | 0.887 | 0.829 | 0.977 | 0.844 | 0.872 | 0.829 | 0.944 | 0.714 | 0.688 | 0.869 | 0.742 | 0.778 | 0.735 | 0.778 | 0.692 | 0.470 | 0.831 |
| | HS-578T | 1.000 | 1.000 | 1.000 | 1.000 | 1.000 | 1.000 | 1.000 | 0.787 | 0.868 | 0.639 | 0.971 | 0.308 | 0.404 | 0.863 | 0.830 | 0.886 | 0.764 | 0.912 | 0.615 | 0.557 | 0.838 |
| | MCF-7 | 0.995 | 0.995 | 0.995 | 0.995 | 0.994 | 0.989 | 1.000 | 0.804 | 0.826 | 0.799 | 0.837 | 0.762 | 0.601 | 0.885 | 0.804 | 0.826 | 0.800 | 0.835 | 0.766 | 0.603 | 0.880 |
| | MDA-MB-231 | 0.997 | 0.998 | 0.997 | 0.998 | 0.996 | 0.994 | 1.000 | 0.841 | 0.881 | 0.811 | 0.922 | 0.700 | 0.649 | 0.894 | 0.816 | 0.861 | 0.788 | 0.891 | 0.685 | 0.594 | 0.886 |
| | MDA-MB-361 | 0.901 | 0.924 | 0.897 | 0.912 | 0.881 | 0.784 | 0.962 | 0.919 | 0.939 | 0.902 | 0.958 | 0.846 | 0.820 | 0.946 | 0.861 | 0.894 | 0.854 | 0.875 | 0.833 | 0.695 | 0.955 |
| | MDA-MB-435 | 0.996 | 0.998 | 0.996 | 0.996 | 0.996 | 0.989 | 1.000 | 0.832 | 0.899 | 0.673 | 0.958 | 0.388 | 0.443 | 0.897 | 0.848 | 0.908 | 0.710 | 0.958 | 0.463 | 0.510 | 0.870 |
| | MDA-MB-453 | 0.994 | 0.994 | 0.994 | 0.994 | 0.994 | 0.989 | 1.000 | 0.864 | 0.857 | 0.864 | 0.818 | 0.909 | 0.730 | 0.928 | 0.841 | 0.844 | 0.841 | 0.864 | 0.818 | 0.683 | 0.888 |
| | MDA-MB-468 | 0.999 | 1.000 | 0.999 | 1.000 | 0.997 | 0.998 | 1.000 | 0.889 | 0.931 | 0.795 | 0.974 | 0.617 | 0.675 | 0.922 | 0.874 | 0.922 | 0.766 | 0.967 | 0.565 | 0.619 | 0.920 |
| | SK-BR-3 | 0.998 | 0.999 | 0.997 | 0.999 | 0.995 | 0.995 | 1.000 | 0.833 | 0.895 | 0.712 | 0.954 | 0.471 | 0.512 | 0.884 | 0.832 | 0.892 | 0.734 | 0.928 | 0.540 | 0.516 | 0.867 |
| | T-47D | 0.998 | 0.998 | 0.997 | 0.998 | 0.996 | 0.994 | 1.000 | 0.803 | 0.868 | 0.722 | 0.923 | 0.521 | 0.499 | 0.864 | 0.812 | 0.874 | 0.729 | 0.932 | 0.527 | 0.520 | 0.887 |
| SVM | Bcap37 | 0.995 | 0.994 | 0.994 | 0.989 | 1.000 | 0.991 | 1.000 | 0.786 | 0.727 | 0.775 | 0.727 | 0.824 | 0.551 | 0.770 | 0.778 | 0.700 | 0.756 | 0.636 | 0.875 | 0.533 | 0.790 |
| | BT-20 | 0.893 | 0.908 | 0.891 | 0.905 | 0.876 | 0.780 | 0.937 | 0.793 | 0.813 | 0.799 | 0.765 | 0.833 | 0.589 | 0.868 | 0.724 | 0.789 | 0.691 | 0.882 | 0.500 | 0.421 | 0.828 |
| | BT-474 | 0.915 | 0.947 | 0.850 | 0.963 | 0.737 | 0.736 | 0.967 | 0.778 | 0.864 | 0.622 | 0.891 | 0.353 | 0.270 | 0.884 | 0.864 | 0.917 | 0.741 | 0.953 | 0.529 | 0.553 | 0.852 |
| | BT-549 | 0.855 | 0.898 | 0.806 | 0.947 | 0.666 | 0.660 | 0.931 | 0.746 | 0.826 | 0.667 | 0.888 | 0.447 | 0.378 | 0.804 | 0.788 | 0.855 | 0.713 | 0.925 | 0.500 | 0.486 | 0.675 |
| | HBL-100 | 0.996 | 0.997 | 0.997 | 0.993 | 1.000 | 0.992 | 1.000 | 0.719 | 0.757 | 0.710 | 0.778 | 0.643 | 0.425 | 0.865 | 0.742 | 0.778 | 0.735 | 0.778 | 0.692 | 0.470 | 0.739 |
| | HS-578T | 0.997 | 0.998 | 0.995 | 1.000 | 0.990 | 0.993 | 1.000 | 0.809 | 0.880 | 0.678 | 0.971 | 0.385 | 0.476 | 0.876 | 0.766 | 0.857 | 0.601 | 0.971 | 0.231 | 0.323 | 0.803 |
| | MCF-7 | 0.921 | 0.928 | 0.920 | 0.926 | 0.914 | 0.839 | 0.977 | 0.785 | 0.809 | 0.781 | 0.819 | 0.744 | 0.564 | 0.857 | 0.787 | 0.810 | 0.784 | 0.814 | 0.754 | 0.569 | 0.860 |
| | MDA-MB-231 | 0.935 | 0.950 | 0.925 | 0.961 | 0.890 | 0.859 | 0.984 | 0.816 | 0.858 | 0.795 | 0.873 | 0.717 | 0.598 | 0.863 | 0.810 | 0.853 | 0.790 | 0.863 | 0.717 | 0.585 | 0.869 |
| | MDA-MB-361 | 0.905 | 0.926 | 0.904 | 0.907 | 0.901 | 0.794 | 0.965 | 0.811 | 0.863 | 0.766 | 0.917 | 0.615 | 0.572 | 0.929 | 0.861 | 0.894 | 0.854 | 0.875 | 0.833 | 0.695 | 0.958 |
| | MDA-MB-435 | 0.834 | 0.901 | 0.657 | 0.974 | 0.340 | 0.442 | 0.915 | 0.805 | 0.885 | 0.608 | 0.962 | 0.254 | 0.319 | 0.826 | 0.822 | 0.895 | 0.629 | 0.975 | 0.284 | 0.389 | 0.830 |
| | MDA-MB-453 | 0.878 | 0.877 | 0.879 | 0.851 | 0.906 | 0.757 | 0.956 | 0.795 | 0.800 | 0.795 | 0.818 | 0.773 | 0.592 | 0.884 | 0.795 | 0.816 | 0.795 | 0.909 | 0.682 | 0.607 | 0.818 |
| | MDA-MB-468 | 0.996 | 0.997 | 0.992 | 0.999 | 0.984 | 0.988 | 0.999 | 0.879 | 0.925 | 0.781 | 0.967 | 0.596 | 0.643 | 0.839 | 0.864 | 0.914 | 0.775 | 0.941 | 0.609 | 0.595 | 0.860 |
| | SK-BR-3 | 0.841 | 0.902 | 0.705 | 0.976 | 0.434 | 0.535 | 0.915 | 0.837 | 0.900 | 0.696 | 0.980 | 0.412 | 0.527 | 0.851 | 0.822 | 0.891 | 0.674 | 0.967 | 0.380 | 0.463 | 0.855 |
| | T-47D | 0.949 | 0.964 | 0.930 | 0.977 | 0.883 | 0.877 | 0.987 | 0.771 | 0.843 | 0.696 | 0.882 | 0.511 | 0.423 | 0.818 | 0.831 | 0.883 | 0.777 | 0.909 | 0.645 | 0.581 | 0.870 |
| XGBoost | Bcap37 | 0.977 | 0.972 | 0.977 | 0.977 | 0.977 | 0.953 | 0.997 | 0.786 | 0.727 | 0.775 | 0.727 | 0.824 | 0.551 | 0.797 | 0.704 | 0.636 | 0.693 | 0.636 | 0.750 | 0.386 | 0.699 |
| | BT-20 | 1.000 | 1.000 | 1.000 | 1.000 | 1.000 | 1.000 | 1.000 | 0.828 | 0.848 | 0.828 | 0.824 | 0.833 | 0.651 | 0.902 | 0.793 | 0.833 | 0.775 | 0.882 | 0.667 | 0.569 | 0.877 |
| | BT-474 | 0.969 | 0.981 | 0.948 | 0.984 | 0.912 | 0.907 | 0.995 | 0.877 | 0.923 | 0.792 | 0.938 | 0.647 | 0.613 | 0.897 | 0.827 | 0.891 | 0.739 | 0.891 | 0.588 | 0.479 | 0.872 |
| | BT-549 | 0.998 | 0.998 | 0.998 | 0.998 | 0.997 | 0.995 | 1.000 | 0.797 | 0.857 | 0.739 | 0.900 | 0.579 | 0.514 | 0.816 | 0.788 | 0.852 | 0.726 | 0.900 | 0.553 | 0.491 | 0.672 |
| | HBL-100 | 0.996 | 0.997 | 0.997 | 0.993 | 1.000 | 0.992 | 1.000 | 0.719 | 0.743 | 0.718 | 0.722 | 0.714 | 0.434 | 0.857 | 0.774 | 0.800 | 0.774 | 0.778 | 0.769 | 0.542 | 0.812 |
| | HS-578T | 0.989 | 0.993 | 0.984 | 0.996 | 0.971 | 0.973 | 1.000 | 0.809 | 0.877 | 0.701 | 0.941 | 0.462 | 0.479 | 0.821 | 0.787 | 0.857 | 0.710 | 0.882 | 0.538 | 0.445 | 0.787 |
| | MCF-7 | 0.846 | 0.862 | 0.844 | 0.865 | 0.822 | 0.688 | 0.929 | 0.749 | 0.777 | 0.744 | 0.787 | 0.702 | 0.490 | 0.830 | 0.750 | 0.777 | 0.745 | 0.786 | 0.705 | 0.492 | 0.828 |
| | MDA-MB-231 | 0.912 | 0.932 | 0.897 | 0.951 | 0.844 | 0.808 | 0.973 | 0.797 | 0.847 | 0.767 | 0.878 | 0.655 | 0.552 | 0.862 | 0.794 | 0.843 | 0.767 | 0.866 | 0.667 | 0.546 | 0.864 |
| | MDA-MB-361 | 0.990 | 0.992 | 0.992 | 0.984 | 1.000 | 0.978 | 1.000 | 0.838 | 0.875 | 0.822 | 0.875 | 0.769 | 0.644 | 0.917 | 0.861 | 0.894 | 0.854 | 0.875 | 0.833 | 0.695 | 0.951 |
| | MDA-MB-435 | 0.982 | 0.988 | 0.966 | 0.995 | 0.936 | 0.947 | 0.998 | 0.822 | 0.892 | 0.672 | 0.941 | 0.403 | 0.417 | 0.872 | 0.838 | 0.901 | 0.709 | 0.941 | 0.478 | 0.484 | 0.869 |
| | MDA-MB-453 | 0.994 | 0.994 | 0.994 | 0.994 | 0.994 | 0.989 | 1.000 | 0.682 | 0.696 | 0.682 | 0.727 | 0.636 | 0.365 | 0.843 | 0.795 | 0.780 | 0.795 | 0.727 | 0.864 | 0.596 | 0.872 |
| | MDA-MB-468 | 0.978 | 0.986 | 0.958 | 0.996 | 0.919 | 0.938 | 0.999 | 0.879 | 0.924 | 0.789 | 0.961 | 0.617 | 0.644 | 0.890 | 0.864 | 0.914 | 0.767 | 0.947 | 0.587 | 0.592 | 0.890 |
| | SK-BR-3 | 0.973 | 0.982 | 0.951 | 0.994 | 0.908 | 0.927 | 0.996 | 0.857 | 0.910 | 0.755 | 0.961 | 0.549 | 0.592 | 0.883 | 0.842 | 0.898 | 0.754 | 0.928 | 0.580 | 0.550 | 0.860 |
| | T-47D | 0.942 | 0.960 | 0.916 | 0.981 | 0.852 | 0.860 | 0.988 | 0.761 | 0.838 | 0.680 | 0.882 | 0.479 | 0.395 | 0.831 | 0.824 | 0.881 | 0.757 | 0.923 | 0.591 | 0.558 | 0.867 |

[a] ACC: Accuracy. [b] F1: F1-measure. [c] BA: Balanced accuracy. [d] SE:Sensitivity. [e] SP:Specificity. [f] MCC:Matthews correlation coefficient. [g] AUC:The area under receiver operating characteristic.

**Table S4.** The performance results of models based on MACCS keys.

| Methods | Cell lines | Training set | | | | | | | Validation set | | | | | | | Test set | | | | | | |
|---|---|---|---|---|---|---|---|---|---|---|---|---|---|---|---|---|---|---|---|---|---|---|
| | | ACC[a] | F1[b] | BA[c] | SE[d] | SP[e] | MCC[f] | AUC[g] | ACC | F1 | BA | SE | SP | MCC | AUC | ACC | F1 | BA | SE | SP | MCC | AUC |
| | Bcap37 | 0.736 | 0.651 | 0.716 | 0.614 | 0.818 | 0.442 | 0.810 | 0.607 | 0.421 | 0.564 | 0.364 | 0.765 | 0.139 | 0.781 | 0.741 | 0.696 | 0.739 | 0.727 | 0.750 | 0.472 | 0.824 |
| | BT-20 | 0.876 | 0.894 | 0.873 | 0.891 | 0.856 | 0.745 | 0.956 | 0.793 | 0.813 | 0.799 | 0.765 | 0.833 | 0.589 | 0.887 | 0.759 | 0.800 | 0.745 | 0.824 | 0.667 | 0.498 | 0.745 |
| | BT-474 | 0.884 | 0.929 | 0.780 | 0.961 | 0.599 | 0.627 | 0.916 | 0.852 | 0.913 | 0.669 | 0.984 | 0.353 | 0.489 | 0.864 | 0.815 | 0.884 | 0.710 | 0.891 | 0.529 | 0.430 | 0.866 |
| | BT-549 | 0.882 | 0.914 | 0.855 | 0.931 | 0.779 | 0.726 | 0.943 | 0.729 | 0.802 | 0.683 | 0.813 | 0.553 | 0.371 | 0.754 | 0.763 | 0.825 | 0.728 | 0.825 | 0.632 | 0.457 | 0.742 |
| | HBL-100 | 0.794 | 0.823 | 0.789 | 0.823 | 0.755 | 0.578 | 0.890 | 0.719 | 0.780 | 0.694 | 0.889 | 0.500 | 0.429 | 0.825 | 0.645 | 0.703 | 0.630 | 0.722 | 0.538 | 0.264 | 0.701 |
| | HS-578T | 0.867 | 0.913 | 0.783 | 0.970 | 0.596 | 0.651 | 0.935 | 0.766 | 0.857 | 0.601 | 0.971 | 0.231 | 0.323 | 0.781 | 0.851 | 0.904 | 0.755 | 0.971 | 0.538 | 0.606 | 0.758 |
| DNN | MCF-7 | 0.856 | 0.874 | 0.851 | 0.896 | 0.807 | 0.708 | 0.936 | 0.767 | 0.797 | 0.759 | 0.825 | 0.694 | 0.525 | 0.846 | 0.756 | 0.788 | 0.748 | 0.817 | 0.679 | 0.502 | 0.839 |
| | MDA-MB-231 | 0.898 | 0.922 | 0.882 | 0.941 | 0.822 | 0.778 | 0.962 | 0.795 | 0.845 | 0.764 | 0.875 | 0.653 | 0.546 | 0.856 | 0.804 | 0.851 | 0.778 | 0.874 | 0.682 | 0.570 | 0.864 |
| | MDA-MB-361 | 0.949 | 0.961 | 0.940 | 0.969 | 0.911 | 0.886 | 0.989 | 0.865 | 0.902 | 0.825 | 0.958 | 0.692 | 0.699 | 0.907 | 0.889 | 0.913 | 0.896 | 0.875 | 0.917 | 0.766 | 0.962 |
| | MDA-MB-435 | 0.896 | 0.935 | 0.819 | 0.957 | 0.680 | 0.683 | 0.946 | 0.835 | 0.898 | 0.718 | 0.928 | 0.507 | 0.483 | 0.843 | 0.802 | 0.878 | 0.664 | 0.911 | 0.418 | 0.371 | 0.833 |
| | MDA-MB-453 | 0.858 | 0.855 | 0.859 | 0.818 | 0.901 | 0.719 | 0.928 | 0.818 | 0.810 | 0.818 | 0.773 | 0.864 | 0.639 | 0.919 | 0.750 | 0.732 | 0.750 | 0.682 | 0.818 | 0.505 | 0.822 |
| | MDA-MB-468 | 0.909 | 0.942 | 0.843 | 0.966 | 0.720 | 0.734 | 0.950 | 0.879 | 0.924 | 0.789 | 0.961 | 0.617 | 0.644 | 0.870 | 0.843 | 0.901 | 0.746 | 0.928 | 0.565 | 0.534 | 0.873 |
| | SK-BR-3 | 0.858 | 0.911 | 0.747 | 0.968 | 0.526 | 0.590 | 0.925 | 0.813 | 0.883 | 0.686 | 0.941 | 0.431 | 0.449 | 0.830 | 0.817 | 0.885 | 0.690 | 0.941 | 0.440 | 0.456 | 0.834 |
| | T-47D | 0.863 | 0.907 | 0.802 | 0.954 | 0.650 | 0.661 | 0.940 | 0.796 | 0.864 | 0.708 | 0.927 | 0.489 | 0.479 | 0.810 | 0.818 | 0.877 | 0.743 | 0.927 | 0.559 | 0.539 | 0.852 |
| | Bcap37 | 0.977 | 0.971 | 0.973 | 0.955 | 0.992 | 0.953 | 0.999 | 0.679 | 0.609 | 0.671 | 0.636 | 0.706 | 0.338 | 0.794 | 0.778 | 0.750 | 0.784 | 0.818 | 0.750 | 0.559 | 0.801 |
| | BT-20 | 1.000 | 1.000 | 1.000 | 1.000 | 1.000 | 1.000 | 1.000 | 0.862 | 0.882 | 0.858 | 0.882 | 0.833 | 0.716 | 0.917 | 0.793 | 0.833 | 0.775 | 0.882 | 0.667 | 0.569 | 0.806 |
| | BT-474 | 0.995 | 0.997 | 0.992 | 0.998 | 0.985 | 0.986 | 0.992 | 0.914 | 0.947 | 0.816 | 0.984 | 0.647 | 0.724 | 0.816 | 0.790 | 0.868 | 0.673 | 0.875 | 0.471 | 0.353 | 0.673 |
| | BT-549 | 0.990 | 0.993 | 0.991 | 0.989 | 0.994 | 0.978 | 1.000 | 0.814 | 0.863 | 0.787 | 0.863 | 0.711 | 0.573 | 0.868 | 0.720 | 0.792 | 0.683 | 0.788 | 0.579 | 0.364 | 0.704 |
| | HBL-100 | 0.988 | 0.990 | 0.986 | 1.000 | 0.972 | 0.976 | 0.986 | 0.719 | 0.769 | 0.702 | 0.833 | 0.571 | 0.423 | 0.702 | 0.742 | 0.789 | 0.724 | 0.833 | 0.615 | 0.463 | 0.724 |
| | HS-578T | 0.987 | 0.991 | 0.991 | 0.982 | 1.000 | 0.968 | 1.000 | 0.809 | 0.873 | 0.725 | 0.912 | 0.538 | 0.492 | 0.805 | 0.830 | 0.886 | 0.764 | 0.912 | 0.615 | 0.557 | 0.768 |
| KNN | MCF-7 | 0.848 | 0.865 | 0.845 | 0.873 | 0.817 | 0.692 | 0.932 | 0.772 | 0.797 | 0.768 | 0.804 | 0.731 | 0.537 | 0.852 | 0.768 | 0.795 | 0.763 | 0.812 | 0.714 | 0.529 | 0.847 |
| | MDA-MB-231 | 0.849 | 0.883 | 0.829 | 0.900 | 0.759 | 0.669 | 0.931 | 0.801 | 0.849 | 0.773 | 0.875 | 0.670 | 0.561 | 0.854 | 0.813 | 0.856 | 0.790 | 0.873 | 0.707 | 0.589 | 0.872 |
| | MDA-MB-361 | 0.993 | 0.995 | 0.992 | 0.995 | 0.990 | 0.985 | 0.992 | 0.838 | 0.880 | 0.804 | 0.917 | 0.692 | 0.636 | 0.804 | 0.861 | 0.894 | 0.854 | 0.875 | 0.833 | 0.695 | 0.854 |
| | MDA-MB-435 | 0.871 | 0.920 | 0.767 | 0.954 | 0.579 | 0.598 | 0.925 | 0.835 | 0.900 | 0.691 | 0.949 | 0.433 | 0.463 | 0.851 | 0.822 | 0.891 | 0.683 | 0.932 | 0.433 | 0.426 | 0.853 |
| | MDA-MB-453 | 0.991 | 0.992 | 0.991 | 0.994 | 0.988 | 0.983 | 0.991 | 0.795 | 0.800 | 0.795 | 0.818 | 0.773 | 0.592 | 0.795 | 0.818 | 0.833 | 0.818 | 0.909 | 0.727 | 0.647 | 0.818 |
| | MDA-MB-468 | 0.987 | 0.992 | 0.985 | 0.989 | 0.981 | 0.965 | 0.998 | 0.879 | 0.922 | 0.818 | 0.934 | 0.702 | 0.657 | 0.891 | 0.889 | 0.928 | 0.837 | 0.934 | 0.739 | 0.684 | 0.903 |
| | SK-BR-3 | 0.973 | 0.982 | 0.974 | 0.973 | 0.975 | 0.931 | 0.998 | 0.823 | 0.882 | 0.758 | 0.888 | 0.627 | 0.523 | 0.841 | 0.797 | 0.864 | 0.738 | 0.855 | 0.620 | 0.466 | 0.802 |
| | T-47D | 0.974 | 0.982 | 0.973 | 0.976 | 0.971 | 0.940 | 0.973 | 0.739 | 0.817 | 0.677 | 0.832 | 0.521 | 0.363 | 0.677 | 0.802 | 0.862 | 0.747 | 0.882 | 0.613 | 0.512 | 0.747 |
| | Bcap37 | 0.682 | 0.635 | 0.684 | 0.693 | 0.674 | 0.361 | 0.767 | 0.714 | 0.667 | 0.717 | 0.727 | 0.706 | 0.424 | 0.722 | 0.630 | 0.615 | 0.645 | 0.727 | 0.563 | 0.287 | 0.795 |
| | BT-20 | 0.671 | 0.709 | 0.668 | 0.686 | 0.649 | 0.332 | 0.791 | 0.690 | 0.667 | 0.723 | 0.529 | 0.917 | 0.462 | 0.799 | 0.724 | 0.733 | 0.740 | 0.647 | 0.833 | 0.476 | 0.809 |
| | BT-474 | 0.744 | 0.822 | 0.736 | 0.750 | 0.723 | 0.404 | 0.810 | 0.704 | 0.797 | 0.661 | 0.734 | 0.588 | 0.279 | 0.744 | 0.728 | 0.817 | 0.677 | 0.766 | 0.588 | 0.312 | 0.767 |
| | BT-549 | 0.670 | 0.750 | 0.636 | 0.734 | 0.539 | 0.267 | 0.710 | 0.653 | 0.739 | 0.613 | 0.725 | 0.500 | 0.221 | 0.672 | 0.568 | 0.667 | 0.529 | 0.638 | 0.421 | 0.056 | 0.537 |
| | HBL-100 | 0.787 | 0.811 | 0.786 | 0.789 | 0.783 | 0.567 | 0.867 | 0.688 | 0.737 | 0.675 | 0.778 | 0.571 | 0.358 | 0.806 | 0.613 | 0.647 | 0.613 | 0.611 | 0.615 | 0.224 | 0.684 |
| | HS-578T | 0.699 | 0.781 | 0.661 | 0.745 | 0.577 | 0.304 | 0.731 | 0.617 | 0.719 | 0.569 | 0.676 | 0.462 | 0.128 | 0.654 | 0.681 | 0.769 | 0.637 | 0.735 | 0.538 | 0.258 | 0.751 |
| NB | MCF-7 | 0.580 | 0.606 | 0.579 | 0.581 | 0.577 | 0.158 | 0.610 | 0.589 | 0.612 | 0.589 | 0.584 | 0.595 | 0.178 | 0.625 | 0.575 | 0.605 | 0.573 | 0.586 | 0.561 | 0.146 | 0.606 |
| | MDA-MB-231 | 0.597 | 0.670 | 0.581 | 0.641 | 0.521 | 0.158 | 0.632 | 0.606 | 0.680 | 0.587 | 0.657 | 0.517 | 0.170 | 0.638 | 0.598 | 0.669 | 0.584 | 0.636 | 0.532 | 0.163 | 0.636 |
| | MDA-MB-361 | 0.789 | 0.826 | 0.802 | 0.762 | 0.842 | 0.576 | 0.894 | 0.784 | 0.833 | 0.763 | 0.833 | 0.692 | 0.526 | 0.856 | 0.778 | 0.810 | 0.813 | 0.708 | 0.917 | 0.589 | 0.938 |
| | MDA-MB-435 | 0.730 | 0.824 | 0.628 | 0.811 | 0.445 | 0.247 | 0.698 | 0.736 | 0.828 | 0.633 | 0.818 | 0.448 | 0.258 | 0.705 | 0.739 | 0.836 | 0.592 | 0.856 | 0.328 | 0.197 | 0.691 |
| | MDA-MB-453 | 0.756 | 0.768 | 0.755 | 0.785 | 0.725 | 0.511 | 0.804 | 0.682 | 0.667 | 0.682 | 0.636 | 0.727 | 0.365 | 0.748 | 0.682 | 0.682 | 0.682 | 0.682 | 0.682 | 0.364 | 0.824 |
| | MDA-MB-468 | 0.739 | 0.825 | 0.665 | 0.804 | 0.526 | 0.313 | 0.765 | 0.734 | 0.822 | 0.657 | 0.803 | 0.511 | 0.299 | 0.754 | 0.712 | 0.808 | 0.623 | 0.789 | 0.457 | 0.235 | 0.737 |
| | SK-BR-3 | 0.656 | 0.741 | 0.658 | 0.655 | 0.660 | 0.276 | 0.726 | 0.611 | 0.713 | 0.577 | 0.645 | 0.510 | 0.137 | 0.640 | 0.644 | 0.735 | 0.629 | 0.658 | 0.600 | 0.227 | 0.690 |
| | T-47D | 0.658 | 0.748 | 0.614 | 0.724 | 0.503 | 0.219 | 0.667 | 0.650 | 0.741 | 0.607 | 0.714 | 0.500 | 0.205 | 0.696 | 0.629 | 0.728 | 0.578 | 0.705 | 0.452 | 0.150 | 0.648 |
| | Bcap37 | 0.936 | 0.918 | 0.928 | 0.886 | 0.970 | 0.867 | 0.989 | 0.679 | 0.609 | 0.671 | 0.636 | 0.706 | 0.338 | 0.783 | 0.778 | 0.750 | 0.784 | 0.818 | 0.750 | 0.559 | 0.849 |
| | BT-20 | 1.000 | 1.000 | 1.000 | 1.000 | 1.000 | 1.000 | 1.000 | 0.862 | 0.882 | 0.858 | 0.882 | 0.833 | 0.716 | 0.902 | 0.759 | 0.811 | 0.733 | 0.882 | 0.583 | 0.496 | 0.801 |
| | BT-474 | 0.997 | 0.998 | 0.995 | 0.998 | 0.993 | 0.991 | 1.000 | 0.901 | 0.940 | 0.786 | 0.984 | 0.588 | 0.681 | 0.949 | 0.877 | 0.923 | 0.792 | 0.938 | 0.647 | 0.613 | 0.900 |
| | BT-549 | 0.990 | 0.993 | 0.987 | 0.997 | 0.977 | 0.978 | 0.999 | 0.797 | 0.857 | 0.739 | 0.900 | 0.579 | 0.514 | 0.858 | 0.737 | 0.814 | 0.675 | 0.850 | 0.500 | 0.372 | 0.707 |
| | HBL-100 | 0.988 | 0.990 | 0.988 | 0.986 | 0.991 | 0.976 | 1.000 | 0.719 | 0.769 | 0.702 | 0.833 | 0.571 | 0.423 | 0.859 | 0.710 | 0.769 | 0.686 | 0.833 | 0.538 | 0.392 | 0.801 |
| | HS-578T | 0.971 | 0.980 | 0.968 | 0.974 | 0.962 | 0.928 | 0.996 | 0.787 | 0.853 | 0.734 | 0.853 | 0.615 | 0.468 | 0.813 | 0.894 | 0.928 | 0.855 | 0.941 | 0.769 | 0.729 | 0.819 |
| RF | MCF-7 | 0.905 | 0.916 | 0.903 | 0.925 | 0.880 | 0.808 | 0.967 | 0.796 | 0.821 | 0.791 | 0.838 | 0.744 | 0.586 | 0.874 | 0.784 | 0.811 | 0.778 | 0.834 | 0.722 | 0.561 | 0.866 |
| | MDA-MB-231 | 0.971 | 0.977 | 0.967 | 0.982 | 0.952 | 0.937 | 0.998 | 0.799 | 0.847 | 0.772 | 0.871 | 0.672 | 0.557 | 0.874 | 0.826 | 0.868 | 0.799 | 0.896 | 0.702 | 0.616 | 0.885 |
| | MDA-MB-361 | 0.993 | 0.995 | 0.992 | 0.995 | 0.990 | 0.985 | 0.999 | 0.784 | 0.846 | 0.728 | 0.917 | 0.538 | 0.506 | 0.933 | 0.917 | 0.939 | 0.896 | 0.958 | 0.833 | 0.810 | 0.993 |
| | MDA-MB-435 | 0.979 | 0.987 | 0.960 | 0.994 | 0.927 | 0.938 | 0.998 | 0.822 | 0.892 | 0.666 | 0.945 | 0.388 | 0.413 | 0.898 | 0.832 | 0.898 | 0.684 | 0.949 | 0.418 | 0.450 | 0.873 |
| | MDA-MB-453 | 0.991 | 0.992 | 0.992 | 0.989 | 0.994 | 0.983 | 1.000 | 0.818 | 0.810 | 0.818 | 0.773 | 0.864 | 0.639 | 0.937 | 0.750 | 0.766 | 0.750 | 0.818 | 0.682 | 0.505 | 0.872 |
| | MDA-MB-468 | 0.987 | 0.992 | 0.980 | 0.994 | 0.965 | 0.965 | 0.999 | 0.879 | 0.924 | 0.789 | 0.961 | 0.617 | 0.644 | 0.923 | 0.879 | 0.925 | 0.777 | 0.967 | 0.587 | 0.636 | 0.935 |
| | SK-BR-3 | 0.973 | 0.982 | 0.961 | 0.986 | 0.935 | 0.929 | 0.997 | 0.833 | 0.892 | 0.738 | 0.928 | 0.549 | 0.525 | 0.877 | 0.802 | 0.871 | 0.714 | 0.888 | 0.540 | 0.448 | 0.850 |
| | T-47D | 0.980 | 0.986 | 0.974 | 0.990 | 0.959 | 0.953 | 0.998 | 0.755 | 0.834 | 0.670 | 0.882 | 0.457 | 0.375 | 0.827 | 0.805 | 0.869 | 0.725 | 0.923 | 0.527 | 0.504 | 0.886 |
| | Bcap37 | 0.977 | 0.971 | 0.975 | 0.966 | 0.985 | 0.953 | 0.993 | 0.714 | 0.667 | 0.717 | 0.727 | 0.706 | 0.424 | 0.791 | 0.815 | 0.762 | 0.801 | 0.727 | 0.875 | 0.613 | 0.784 |
| | BT-20 | 0.897 | 0.913 | 0.893 | 0.920 | 0.866 | 0.788 | 0.951 | 0.828 | 0.848 | 0.828 | 0.824 | 0.833 | 0.651 | 0.819 | 0.793 | 0.833 | 0.775 | 0.882 | 0.667 | 0.569 | 0.819 |
| | BT-474 | 0.914 | 0.947 | 0.820 | 0.982 | 0.657 | 0.726 | 0.959 | 0.877 | 0.926 | 0.727 | 0.984 | 0.471 | 0.590 | 0.901 | 0.827 | 0.894 | 0.696 | 0.922 | 0.471 | 0.435 | 0.872 |
| | BT-549 | 0.863 | 0.904 | 0.812 | 0.958 | 0.666 | 0.679 | 0.946 | 0.763 | 0.841 | 0.673 | 0.925 | 0.421 | 0.415 | 0.762 | 0.797 | 0.857 | 0.739 | 0.900 | 0.579 | 0.514 | 0.689 |
| | HBL-100 | 0.885 | 0.902 | 0.882 | 0.905 | 0.858 | 0.764 | 0.953 | 0.781 | 0.821 | 0.766 | 0.889 | 0.643 | 0.555 | 0.853 | 0.645 | 0.703 | 0.630 | 0.722 | 0.538 | 0.264 | 0.752 |
| | HS-578T | 0.869 | 0.915 | 0.788 | 0.970 | 0.606 | 0.658 | 0.949 | 0.766 | 0.853 | 0.624 | 0.941 | 0.308 | 0.334 | 0.713 | 0.851 | 0.901 | 0.778 | 0.941 | 0.615 | 0.608 | 0.794 |
| SVM | MCF-7 | 0.933 | 0.940 | 0.932 | 0.942 | 0.922 | 0.865 | 0.978 | 0.790 | 0.813 | 0.786 | 0.822 | 0.750 | 0.574 | 0.855 | 0.789 | 0.813 | 0.785 | 0.826 | 0.744 | 0.572 | 0.852 |
| | MDA-MB-231 | 0.909 | 0.930 | 0.893 | 0.953 | 0.833 | 0.802 | 0.968 | 0.807 | 0.855 | 0.775 | 0.891 | 0.660 | 0.573 | 0.872 | 0.813 | 0.857 | 0.788 | 0.878 | 0.697 | 0.588 | 0.876 |
| | MDA-MB-361 | 0.935 | 0.951 | 0.927 | 0.953 | 0.901 | 0.856 | 0.976 | 0.838 | 0.885 | 0.787 | 0.958 | 0.615 | 0.638 | 0.840 | 0.889 | 0.917 | 0.875 | 0.917 | 0.833 | 0.750 | 0.972 |
| | MDA-MB-435 | 0.979 | 0.987 | 0.958 | 0.995 | 0.921 | 0.938 | 0.981 | 0.828 | 0.896 | 0.671 | 0.953 | 0.388 | 0.433 | 0.838 | 0.835 | 0.899 | 0.696 | 0.945 | 0.448 | 0.467 | 0.800 |
| | MDA-MB-453 | 0.886 | 0.889 | 0.886 | 0.884 | 0.889 | 0.773 | 0.957 | 0.773 | 0.773 | 0.773 | 0.773 | 0.773 | 0.545 | 0.893 | 0.773 | 0.792 | 0.773 | 0.864 | 0.682 | 0.555 | 0.857 |
| | MDA-MB-468 | 0.950 | 0.968 | 0.903 | 0.992 | 0.814 | 0.858 | 0.986 | 0.894 | 0.934 | 0.806 | 0.974 | 0.638 | 0.691 | 0.893 | 0.869 | 0.918 | 0.770 | 0.954 | 0.587 | 0.606 | 0.876 |
| | SK-BR-3 | 0.859 | 0.912 | 0.743 | 0.975 | 0.511 | 0.594 | 0.931 | 0.842 | 0.902 | 0.719 | 0.967 | 0.471 | 0.543 | 0.849 | 0.837 | 0.898 | 0.710 | 0.961 | 0.460 | 0.518 | 0.817 |
| | T-47D | 0.921 | 0.945 | 0.885 | 0.974 | 0.796 | 0.807 | 0.969 | 0.777 | 0.850 | 0.692 | 0.905 | 0.479 | 0.431 | 0.789 | 0.808 | 0.867 | 0.752 | 0.891 | 0.613 | 0.526 | 0.866 |
| | Bcap37 | 0.977 | 0.972 | 0.977 | 0.977 | 0.977 | 0.953 | 0.997 | 0.750 | 0.720 | 0.762 | 0.818 | 0.706 | 0.512 | 0.701 | 0.741 | 0.696 | 0.739 | 0.727 | 0.750 | 0.472 | 0.767 |
| | BT-20 | 0.983 | 0.986 | 0.981 | 0.993 | 0.969 | 0.965 | 0.999 | 0.862 | 0.882 | 0.858 | 0.882 | 0.833 | 0.716 | 0.897 | 0.828 | 0.857 | 0.816 | 0.882 | 0.750 | 0.642 | 0.833 |
| | BT-474 | 0.995 | 0.997 | 0.992 | 0.998 | 0.985 | 0.986 | 1.000 | 0.889 | 0.931 | 0.800 | 0.953 | 0.647 | 0.646 | 0.928 | 0.815 | 0.882 | 0.732 | 0.875 | 0.588 | 0.454 | 0.866 |
| | BT-549 | 0.959 | 0.970 | 0.948 | 0.978 | 0.919 | 0.906 | 0.991 | 0.788 | 0.852 | 0.726 | 0.900 | 0.553 | 0.491 | 0.814 | 0.763 | 0.829 | 0.714 | 0.850 | 0.579 | 0.443 | 0.694 |
| | HBL-100 | 0.937 | 0.946 | 0.934 | 0.952 | 0.915 | 0.870 | 0.985 | 0.688 | 0.737 | 0.675 | 0.778 | 0.571 | 0.358 | 0.796 | 0.742 | 0.789 | 0.724 | 0.833 | 0.615 | 0.463 | 0.825 |
| | HS-578T | 0.915 | 0.943 | 0.870 | 0.970 | 0.769 | 0.781 | 0.981 | 0.787 | 0.868 | 0.639 | 0.971 | 0.308 | 0.404 | 0.751 | 0.809 | 0.870 | 0.749 | 0.882 | 0.615 | 0.511 | 0.796 |
| XGBoost | MCF-7 | 0.810 | 0.832 | 0.805 | 0.848 | 0.761 | 0.613 | 0.894 | 0.750 | 0.779 | 0.745 | 0.793 | 0.697 | 0.492 | 0.828 | 0.745 | 0.777 | 0.739 | 0.800 | 0.678 | 0.482 | 0.821 |
| | MDA-MB-231 | 0.873 | 0.904 | 0.849 | 0.938 | 0.760 | 0.722 | 0.948 | 0.796 | 0.847 | 0.760 | 0.888 | 0.633 | 0.546 | 0.867 | 0.791 | 0.841 | 0.763 | 0.866 | 0.660 | 0.540 | 0.865 |
| | MDA-MB-361 | 0.973 | 0.979 | 0.972 | 0.974 | 0.970 | 0.940 | 0.993 | 0.811 | 0.863 | 0.766 | 0.917 | 0.615 | 0.572 | 0.907 | 0.917 | 0.936 | 0.917 | 0.917 | 0.917 | 0.818 | 0.990 |
| | MDA-MB-435 | 0.946 | 0.966 | 0.903 | 0.980 | 0.826 | 0.839 | 0.987 | 0.848 | 0.907 | 0.716 | 0.953 | 0.478 | 0.513 | 0.882 | 0.845 | 0.905 | 0.719 | 0.945 | 0.493 | 0.506 | 0.858 |
| | MDA-MB-453 | 0.861 | 0.865 | 0.861 | 0.867 | 0.854 | 0.721 | 0.941 | 0.841 | 0.844 | 0.841 | 0.864 | 0.818 | 0.683 | 0.888 | 0.727 | 0.714 | 0.727 | 0.682 | 0.773 | 0.456 | 0.842 |
| | MDA-MB-468 | 0.977 | 0.985 | 0.964 | 0.988 | 0.941 | 0.935 | 0.997 | 0.874 | 0.921 | 0.778 | 0.961 | 0.596 | 0.628 | 0.901 | 0.899 | 0.937 | 0.813 | 0.974 | 0.652 | 0.701 | 0.909 |
| | SK-BR-3 | 0.932 | 0.955 | 0.885 | 0.978 | 0.792 | 0.812 | 0.975 | 0.833 | 0.892 | 0.738 | 0.928 | 0.549 | 0.525 | 0.858 | 0.812 | 0.879 | 0.714 | 0.908 | 0.520 | 0.463 | 0.845 |
| | T-47D | 0.955 | 0.969 | 0.938 | 0.981 | 0.895 | 0.892 | 0.991 | 0.761 | 0.836 | 0.689 | 0.868 | 0.511 | 0.403 | 0.803 | 0.827 | 0.880 | 0.778 | 0.900 | 0.656 | 0.576 | 0.876 |

[a] ACC: Accuracy. [b] F1: F1-measure. [c] BA: Balanced accuracy. [d] SE:Sensitivity. [e] SP:Specificity. [f] MCC:Matthews correlation coefficient. [g] AUC:The area under receiver operating characteristic.

**Table S5.** The performance results of models based on Morgan fingerprints.

| Methods | Cell lines | Training set | | | | | | | Validation set | | | | | | | Test set | | | | | | |
|---|---|---|---|---|---|---|---|---|---|---|---|---|---|---|---|---|---|---|---|---|---|---|
| | | ACC[a] | F1[b] | BA[c] | SE[d] | SP[e] | MCC[f] | AUC[g] | ACC | F1 | BA | SE | SP | MCC | AUC | ACC | F1 | BA | SE | SP | MCC | AUC |
| DNN | Bcap37 | 0.877 | 0.862 | 0.890 | 0.955 | 0.826 | 0.765 | 0.955 | 0.750 | 0.720 | 0.762 | 0.818 | 0.706 | 0.512 | 0.850 | 0.667 | 0.667 | 0.690 | 0.818 | 0.563 | 0.381 | 0.682 |
| | BT-20 | 0.996 | 0.996 | 0.996 | 0.993 | 1.000 | 0.991 | 1.000 | 0.793 | 0.813 | 0.799 | 0.765 | 0.833 | 0.589 | 0.912 | 0.724 | 0.778 | 0.703 | 0.824 | 0.583 | 0.422 | 0.824 |
| | BT-474 | 0.941 | 0.963 | 0.920 | 0.957 | 0.883 | 0.827 | 0.978 | 0.889 | 0.930 | 0.822 | 0.938 | 0.706 | 0.658 | 0.885 | 0.852 | 0.905 | 0.798 | 0.891 | 0.706 | 0.573 | 0.886 |
| | BT-549 | 0.920 | 0.943 | 0.883 | 0.987 | 0.779 | 0.817 | 0.985 | 0.822 | 0.877 | 0.758 | 0.938 | 0.579 | 0.574 | 0.849 | 0.754 | 0.832 | 0.674 | 0.900 | 0.447 | 0.397 | 0.696 |
| | HBL-100 | 0.945 | 0.953 | 0.942 | 0.959 | 0.925 | 0.886 | 0.990 | 0.719 | 0.757 | 0.710 | 0.778 | 0.643 | 0.425 | 0.845 | 0.677 | 0.722 | 0.669 | 0.722 | 0.615 | 0.338 | 0.778 |
| | HS-578T | 0.880 | 0.921 | 0.804 | 0.974 | 0.635 | 0.688 | 0.969 | 0.809 | 0.880 | 0.678 | 0.971 | 0.385 | 0.476 | 0.808 | 0.809 | 0.880 | 0.678 | 0.971 | 0.385 | 0.476 | 0.771 |
| | MCF-7 | 0.971 | 0.974 | 0.971 | 0.972 | 0.969 | 0.941 | 0.996 | 0.800 | 0.819 | 0.797 | 0.819 | 0.775 | 0.594 | 0.873 | 0.797 | 0.817 | 0.795 | 0.814 | 0.777 | 0.590 | 0.875 |
| | MDA-MB-231 | 0.974 | 0.980 | 0.969 | 0.988 | 0.950 | 0.944 | 0.997 | 0.809 | 0.855 | 0.781 | 0.882 | 0.680 | 0.579 | 0.873 | 0.815 | 0.858 | 0.793 | 0.874 | 0.712 | 0.595 | 0.880 |
| | MDA-MB-361 | 0.935 | 0.951 | 0.925 | 0.959 | 0.891 | 0.856 | 0.975 | 0.811 | 0.873 | 0.731 | 1.000 | 0.462 | 0.598 | 0.981 | 0.861 | 0.898 | 0.833 | 0.917 | 0.750 | 0.682 | 0.969 |
| | MDA-MB-435 | 0.904 | 0.941 | 0.809 | 0.980 | 0.637 | 0.705 | 0.966 | 0.828 | 0.896 | 0.671 | 0.953 | 0.388 | 0.433 | 0.873 | 0.832 | 0.897 | 0.689 | 0.945 | 0.433 | 0.454 | 0.815 |
| | MDA-MB-453 | 0.963 | 0.963 | 0.964 | 0.945 | 0.982 | 0.927 | 0.995 | 0.818 | 0.810 | 0.818 | 0.773 | 0.864 | 0.639 | 0.934 | 0.727 | 0.727 | 0.727 | 0.727 | 0.727 | 0.455 | 0.775 |
| | MDA-MB-468 | 0.988 | 0.992 | 0.983 | 0.993 | 0.973 | 0.967 | 0.999 | 0.874 | 0.919 | 0.808 | 0.934 | 0.681 | 0.640 | 0.897 | 0.869 | 0.916 | 0.801 | 0.928 | 0.674 | 0.621 | 0.893 |
| | SK-BR-3 | 0.922 | 0.950 | 0.870 | 0.974 | 0.767 | 0.785 | 0.976 | 0.862 | 0.913 | 0.765 | 0.961 | 0.569 | 0.608 | 0.900 | 0.842 | 0.899 | 0.747 | 0.934 | 0.560 | 0.546 | 0.847 |
| | T-47D | 0.883 | 0.921 | 0.824 | 0.972 | 0.676 | 0.713 | 0.957 | 0.764 | 0.845 | 0.661 | 0.918 | 0.404 | 0.386 | 0.836 | 0.783 | 0.855 | 0.696 | 0.909 | 0.484 | 0.443 | 0.813 |
| KNN | Bcap37 | 0.995 | 0.994 | 0.994 | 0.989 | 1.000 | 0.991 | 1.000 | 0.750 | 0.667 | 0.730 | 0.636 | 0.824 | 0.469 | 0.824 | 0.704 | 0.600 | 0.679 | 0.545 | 0.813 | 0.373 | 0.798 |
| | BT-20 | 1.000 | 1.000 | 1.000 | 1.000 | 1.000 | 1.000 | 1.000 | 0.759 | 0.774 | 0.770 | 0.706 | 0.833 | 0.531 | 0.770 | 0.759 | 0.811 | 0.733 | 0.882 | 0.583 | 0.496 | 0.733 |
| | BT-474 | 1.000 | 1.000 | 1.000 | 1.000 | 1.000 | 1.000 | 1.000 | 0.877 | 0.925 | 0.749 | 0.969 | 0.529 | 0.592 | 0.749 | 0.840 | 0.894 | 0.812 | 0.859 | 0.765 | 0.571 | 0.812 |
| | BT-549 | 0.998 | 0.998 | 0.998 | 0.997 | 1.000 | 0.995 | 1.000 | 0.822 | 0.870 | 0.793 | 0.875 | 0.711 | 0.590 | 0.895 | 0.703 | 0.780 | 0.664 | 0.775 | 0.553 | 0.325 | 0.691 |
| | HBL-100 | 0.996 | 0.997 | 0.997 | 0.993 | 1.000 | 0.992 | 1.000 | 0.750 | 0.778 | 0.746 | 0.778 | 0.714 | 0.492 | 0.839 | 0.774 | 0.811 | 0.763 | 0.833 | 0.692 | 0.533 | 0.784 |
| | HS-578T | 1.000 | 1.000 | 1.000 | 1.000 | 1.000 | 1.000 | 1.000 | 0.851 | 0.904 | 0.755 | 0.971 | 0.538 | 0.606 | 0.846 | 0.830 | 0.886 | 0.764 | 0.912 | 0.615 | 0.557 | 0.762 |
| | MCF-7 | 0.992 | 0.993 | 0.992 | 0.988 | 0.997 | 0.984 | 1.000 | 0.804 | 0.824 | 0.801 | 0.826 | 0.776 | 0.602 | 0.878 | 0.804 | 0.823 | 0.802 | 0.820 | 0.783 | 0.603 | 0.873 |
| | MDA-MB-231 | 0.995 | 0.996 | 0.996 | 0.994 | 0.998 | 0.989 | 1.000 | 0.829 | 0.869 | 0.808 | 0.885 | 0.732 | 0.627 | 0.883 | 0.837 | 0.873 | 0.820 | 0.881 | 0.759 | 0.644 | 0.894 |
| | MDA-MB-361 | 0.997 | 0.997 | 0.997 | 0.995 | 1.000 | 0.993 | 1.000 | 0.919 | 0.936 | 0.920 | 0.917 | 0.923 | 0.827 | 0.968 | 0.889 | 0.909 | 0.917 | 0.833 | 1.000 | 0.791 | 0.934 |
| | MDA-MB-435 | 0.995 | 0.997 | 0.996 | 0.994 | 0.998 | 0.986 | 1.000 | 0.848 | 0.904 | 0.764 | 0.915 | 0.612 | 0.546 | 0.887 | 0.818 | 0.885 | 0.718 | 0.898 | 0.537 | 0.454 | 0.833 |
| | MDA-MB-453 | 0.858 | 0.848 | 0.860 | 0.773 | 0.947 | 0.729 | 0.947 | 0.773 | 0.750 | 0.773 | 0.682 | 0.864 | 0.555 | 0.893 | 0.773 | 0.750 | 0.773 | 0.682 | 0.864 | 0.555 | 0.775 |
| | MDA-MB-468 | 0.999 | 1.000 | 1.000 | 0.999 | 1.000 | 0.998 | 1.000 | 0.889 | 0.930 | 0.810 | 0.961 | 0.660 | 0.677 | 0.934 | 0.854 | 0.906 | 0.783 | 0.914 | 0.652 | 0.580 | 0.898 |
| | SK-BR-3 | 0.997 | 0.998 | 0.997 | 0.997 | 0.998 | 0.992 | 1.000 | 0.828 | 0.886 | 0.761 | 0.895 | 0.627 | 0.533 | 0.870 | 0.837 | 0.893 | 0.771 | 0.901 | 0.640 | 0.553 | 0.842 |
| | T-47D | 0.998 | 0.998 | 0.998 | 0.997 | 1.000 | 0.994 | 1.000 | 0.787 | 0.850 | 0.735 | 0.864 | 0.606 | 0.481 | 0.832 | 0.831 | 0.880 | 0.796 | 0.882 | 0.710 | 0.593 | 0.865 |
| NB | Bcap37 | 0.777 | 0.688 | 0.750 | 0.614 | 0.886 | 0.528 | 0.872 | 0.714 | 0.600 | 0.684 | 0.545 | 0.824 | 0.386 | 0.775 | 0.667 | 0.571 | 0.648 | 0.545 | 0.750 | 0.301 | 0.719 |
| | BT-20 | 0.846 | 0.851 | 0.866 | 0.752 | 0.979 | 0.724 | 0.960 | 0.724 | 0.714 | 0.752 | 0.588 | 0.917 | 0.512 | 0.833 | 0.793 | 0.800 | 0.811 | 0.706 | 0.917 | 0.617 | 0.819 |
| | BT-474 | 0.846 | 0.894 | 0.873 | 0.826 | 0.920 | 0.647 | 0.938 | 0.778 | 0.847 | 0.773 | 0.781 | 0.765 | 0.472 | 0.896 | 0.815 | 0.872 | 0.840 | 0.797 | 0.882 | 0.582 | 0.872 |
| | BT-549 | 0.870 | 0.901 | 0.867 | 0.876 | 0.857 | 0.715 | 0.933 | 0.763 | 0.823 | 0.735 | 0.813 | 0.658 | 0.464 | 0.800 | 0.712 | 0.788 | 0.670 | 0.788 | 0.553 | 0.340 | 0.680 |
| | HBL-100 | 0.909 | 0.920 | 0.910 | 0.905 | 0.915 | 0.815 | 0.973 | 0.750 | 0.778 | 0.746 | 0.778 | 0.714 | 0.492 | 0.815 | 0.710 | 0.769 | 0.686 | 0.833 | 0.538 | 0.392 | 0.744 |
| | HS-578T | 0.901 | 0.931 | 0.887 | 0.919 | 0.856 | 0.760 | 0.947 | 0.809 | 0.877 | 0.701 | 0.941 | 0.462 | 0.479 | 0.785 | 0.702 | 0.788 | 0.652 | 0.765 | 0.538 | 0.291 | 0.701 |
| | MCF-7 | 0.645 | 0.652 | 0.650 | 0.599 | 0.702 | 0.300 | 0.705 | 0.655 | 0.664 | 0.659 | 0.616 | 0.703 | 0.317 | 0.713 | 0.609 | 0.617 | 0.614 | 0.568 | 0.660 | 0.227 | 0.682 |
| | MDA-MB-231 | 0.708 | 0.755 | 0.709 | 0.706 | 0.712 | 0.405 | 0.779 | 0.698 | 0.744 | 0.702 | 0.689 | 0.714 | 0.389 | 0.762 | 0.702 | 0.750 | 0.702 | 0.700 | 0.704 | 0.391 | 0.769 |
| | MDA-MB-361 | 0.871 | 0.892 | 0.897 | 0.813 | 0.980 | 0.756 | 0.970 | 0.865 | 0.889 | 0.843 | 0.917 | 0.769 | 0.699 | 0.974 | 0.889 | 0.913 | 0.896 | 0.875 | 0.917 | 0.766 | 0.958 |
| | MDA-MB-435 | 0.826 | 0.886 | 0.778 | 0.864 | 0.692 | 0.527 | 0.852 | 0.799 | 0.870 | 0.716 | 0.864 | 0.567 | 0.425 | 0.793 | 0.812 | 0.879 | 0.724 | 0.881 | 0.567 | 0.451 | 0.811 |
| | MDA-MB-453 | 0.892 | 0.890 | 0.893 | 0.851 | 0.936 | 0.788 | 0.941 | 0.795 | 0.791 | 0.795 | 0.773 | 0.818 | 0.592 | 0.876 | 0.727 | 0.727 | 0.727 | 0.727 | 0.727 | 0.455 | 0.789 |
| | MDA-MB-468 | 0.849 | 0.898 | 0.833 | 0.863 | 0.803 | 0.619 | 0.911 | 0.819 | 0.878 | 0.779 | 0.855 | 0.702 | 0.529 | 0.857 | 0.773 | 0.847 | 0.716 | 0.822 | 0.609 | 0.406 | 0.850 |
| | SK-BR-3 | 0.737 | 0.794 | 0.798 | 0.676 | 0.921 | 0.516 | 0.883 | 0.704 | 0.769 | 0.751 | 0.658 | 0.843 | 0.436 | 0.826 | 0.688 | 0.759 | 0.726 | 0.651 | 0.800 | 0.391 | 0.801 |
| | T-47D | 0.787 | 0.844 | 0.764 | 0.823 | 0.705 | 0.512 | 0.850 | 0.736 | 0.810 | 0.690 | 0.805 | 0.574 | 0.376 | 0.789 | 0.690 | 0.767 | 0.665 | 0.727 | 0.602 | 0.312 | 0.751 |
| RF | Bcap37 | 0.759 | 0.619 | 0.714 | 0.489 | 0.939 | 0.497 | 0.876 | 0.679 | 0.400 | 0.607 | 0.273 | 0.941 | 0.299 | 0.882 | 0.704 | 0.600 | 0.679 | 0.545 | 0.813 | 0.373 | 0.793 |
| | BT-20 | 0.944 | 0.952 | 0.947 | 0.934 | 0.959 | 0.887 | 0.983 | 0.828 | 0.865 | 0.804 | 0.941 | 0.667 | 0.647 | 0.931 | 0.759 | 0.800 | 0.745 | 0.824 | 0.667 | 0.498 | 0.838 |
| | BT-474 | 1.000 | 1.000 | 1.000 | 1.000 | 1.000 | 1.000 | 1.000 | 0.877 | 0.928 | 0.706 | 1.000 | 0.412 | 0.597 | 0.929 | 0.827 | 0.896 | 0.675 | 0.938 | 0.412 | 0.415 | 0.884 |
| | BT-549 | 0.998 | 0.998 | 0.998 | 0.998 | 0.997 | 0.995 | 1.000 | 0.847 | 0.894 | 0.791 | 0.950 | 0.632 | 0.639 | 0.887 | 0.754 | 0.830 | 0.681 | 0.888 | 0.474 | 0.402 | 0.698 |
| | HBL-100 | 0.996 | 0.997 | 0.995 | 1.000 | 0.991 | 0.992 | 1.000 | 0.781 | 0.821 | 0.766 | 0.889 | 0.643 | 0.555 | 0.857 | 0.839 | 0.872 | 0.818 | 0.944 | 0.692 | 0.672 | 0.823 |
| | HS-578T | 1.000 | 1.000 | 1.000 | 1.000 | 1.000 | 1.000 | 1.000 | 0.830 | 0.895 | 0.692 | 1.000 | 0.385 | 0.558 | 0.835 | 0.809 | 0.873 | 0.725 | 0.912 | 0.538 | 0.492 | 0.741 |
| | MCF-7 | 0.992 | 0.993 | 0.992 | 0.994 | 0.990 | 0.984 | 1.000 | 0.814 | 0.833 | 0.811 | 0.839 | 0.783 | 0.623 | 0.888 | 0.808 | 0.827 | 0.805 | 0.830 | 0.780 | 0.610 | 0.886 |
| | MDA-MB-231 | 0.995 | 0.996 | 0.995 | 0.996 | 0.993 | 0.989 | 1.000 | 0.836 | 0.876 | 0.807 | 0.910 | 0.704 | 0.638 | 0.894 | 0.837 | 0.876 | 0.811 | 0.905 | 0.717 | 0.640 | 0.904 |
| | MDA-MB-361 | 0.997 | 0.997 | 0.995 | 1.000 | 0.990 | 0.992 | 1.000 | 0.892 | 0.923 | 0.846 | 1.000 | 0.692 | 0.770 | 0.981 | 0.917 | 0.936 | 0.917 | 0.917 | 0.917 | 0.818 | 0.972 |
| | MDA-MB-435 | 0.995 | 0.997 | 0.991 | 0.998 | 0.985 | 0.986 | 1.000 | 0.835 | 0.900 | 0.680 | 0.958 | 0.403 | 0.457 | 0.907 | 0.825 | 0.893 | 0.679 | 0.941 | 0.418 | 0.431 | 0.866 |
| | MDA-MB-453 | 0.994 | 0.994 | 0.994 | 0.994 | 0.994 | 0.989 | 1.000 | 0.818 | 0.800 | 0.818 | 0.727 | 0.909 | 0.647 | 0.930 | 0.750 | 0.744 | 0.750 | 0.727 | 0.773 | 0.501 | 0.851 |
| | MDA-MB-468 | 0.999 | 1.000 | 0.999 | 1.000 | 0.997 | 0.998 | 1.000 | 0.894 | 0.934 | 0.806 | 0.974 | 0.638 | 0.691 | 0.935 | 0.869 | 0.918 | 0.763 | 0.961 | 0.565 | 0.603 | 0.923 |
| | SK-BR-3 | 0.997 | 0.998 | 0.996 | 0.998 | 0.995 | 0.992 | 1.000 | 0.833 | 0.892 | 0.738 | 0.928 | 0.549 | 0.525 | 0.886 | 0.822 | 0.884 | 0.741 | 0.901 | 0.580 | 0.503 | 0.869 |
| | T-47D | 0.998 | 0.998 | 0.997 | 0.998 | 0.996 | 0.994 | 1.000 | 0.790 | 0.860 | 0.701 | 0.923 | 0.479 | 0.462 | 0.862 | 0.840 | 0.892 | 0.772 | 0.941 | 0.602 | 0.599 | 0.885 |
| SVM | Bcap37 | 0.995 | 0.994 | 0.994 | 0.989 | 1.000 | 0.991 | 1.000 | 0.786 | 0.700 | 0.759 | 0.636 | 0.882 | 0.542 | 0.743 | 0.741 | 0.632 | 0.710 | 0.545 | 0.875 | 0.452 | 0.682 |
| | BT-20 | 0.949 | 0.956 | 0.949 | 0.949 | 0.948 | 0.895 | 0.989 | 0.828 | 0.848 | 0.828 | 0.824 | 0.833 | 0.651 | 0.838 | 0.690 | 0.757 | 0.662 | 0.824 | 0.500 | 0.344 | 0.824 |
| | BT-474 | 0.982 | 0.988 | 0.956 | 1.000 | 0.912 | 0.944 | 0.999 | 0.877 | 0.924 | 0.771 | 0.953 | 0.588 | 0.601 | 0.841 | 0.864 | 0.915 | 0.784 | 0.922 | 0.647 | 0.582 | 0.856 |
| | BT-549 | 0.940 | 0.956 | 0.922 | 0.973 | 0.870 | 0.861 | 0.986 | 0.856 | 0.899 | 0.804 | 0.950 | 0.658 | 0.660 | 0.851 | 0.754 | 0.828 | 0.688 | 0.875 | 0.500 | 0.407 | 0.697 |
| | HBL-100 | 0.996 | 0.997 | 0.995 | 1.000 | 0.991 | 0.992 | 1.000 | 0.781 | 0.821 | 0.766 | 0.889 | 0.643 | 0.555 | 0.845 | 0.742 | 0.800 | 0.714 | 0.889 | 0.538 | 0.465 | 0.799 |
| | HS-578T | 0.917 | 0.945 | 0.863 | 0.985 | 0.740 | 0.789 | 0.990 | 0.809 | 0.880 | 0.678 | 0.971 | 0.385 | 0.476 | 0.803 | 0.851 | 0.904 | 0.755 | 0.971 | 0.538 | 0.606 | 0.826 |
| | MCF-7 | 0.980 | 0.982 | 0.979 | 0.982 | 0.977 | 0.959 | 0.996 | 0.810 | 0.831 | 0.807 | 0.839 | 0.774 | 0.615 | 0.876 | 0.810 | 0.831 | 0.807 | 0.840 | 0.774 | 0.615 | 0.872 |
| | MDA-MB-231 | 0.995 | 0.996 | 0.994 | 0.997 | 0.991 | 0.989 | 0.998 | 0.835 | 0.875 | 0.807 | 0.909 | 0.704 | 0.636 | 0.876 | 0.829 | 0.871 | 0.799 | 0.906 | 0.692 | 0.621 | 0.887 |
| | MDA-MB-361 | 0.952 | 0.964 | 0.950 | 0.959 | 0.941 | 0.895 | 0.979 | 0.865 | 0.906 | 0.808 | 1.000 | 0.615 | 0.714 | 0.968 | 0.917 | 0.936 | 0.917 | 0.917 | 0.917 | 0.818 | 0.941 |
| | MDA-MB-435 | 0.910 | 0.944 | 0.817 | 0.983 | 0.652 | 0.723 | 0.984 | 0.832 | 0.899 | 0.662 | 0.966 | 0.358 | 0.438 | 0.884 | 0.825 | 0.895 | 0.658 | 0.958 | 0.358 | 0.415 | 0.863 |
| | MDA-MB-453 | 0.963 | 0.963 | 0.964 | 0.939 | 0.988 | 0.927 | 0.993 | 0.864 | 0.864 | 0.864 | 0.864 | 0.864 | 0.727 | 0.936 | 0.750 | 0.756 | 0.750 | 0.773 | 0.727 | 0.501 | 0.839 |
| | MDA-MB-468 | 0.941 | 0.962 | 0.886 | 0.989 | 0.784 | 0.830 | 0.990 | 0.874 | 0.923 | 0.756 | 0.980 | 0.532 | 0.626 | 0.897 | 0.859 | 0.913 | 0.734 | 0.967 | 0.500 | 0.566 | 0.902 |
| | SK-BR-3 | 0.893 | 0.932 | 0.813 | 0.973 | 0.653 | 0.700 | 0.977 | 0.837 | 0.898 | 0.722 | 0.954 | 0.490 | 0.529 | 0.875 | 0.827 | 0.889 | 0.731 | 0.921 | 0.540 | 0.504 | 0.868 |
| | T-47D | 0.928 | 0.950 | 0.899 | 0.972 | 0.825 | 0.826 | 0.983 | 0.803 | 0.869 | 0.716 | 0.932 | 0.500 | 0.497 | 0.857 | 0.815 | 0.874 | 0.747 | 0.914 | 0.581 | 0.534 | 0.872 |
| XGBoost | Bcap37 | 0.982 | 0.977 | 0.979 | 0.966 | 0.992 | 0.962 | 0.998 | 0.750 | 0.667 | 0.730 | 0.636 | 0.824 | 0.469 | 0.754 | 0.704 | 0.667 | 0.707 | 0.727 | 0.688 | 0.408 | 0.744 |
| | BT-20 | 0.889 | 0.902 | 0.893 | 0.869 | 0.918 | 0.777 | 0.966 | 0.793 | 0.824 | 0.787 | 0.824 | 0.750 | 0.574 | 0.868 | 0.759 | 0.811 | 0.733 | 0.882 | 0.583 | 0.496 | 0.740 |
| | BT-474 | 0.997 | 0.998 | 0.993 | 1.000 | 0.985 | 0.991 | 1.000 | 0.877 | 0.923 | 0.792 | 0.938 | 0.647 | 0.613 | 0.884 | 0.802 | 0.879 | 0.659 | 0.906 | 0.412 | 0.353 | 0.827 |
| | BT-549 | 0.944 | 0.959 | 0.927 | 0.975 | 0.880 | 0.871 | 0.991 | 0.814 | 0.872 | 0.745 | 0.938 | 0.553 | 0.553 | 0.808 | 0.695 | 0.783 | 0.630 | 0.813 | 0.447 | 0.273 | 0.651 |
| | HBL-100 | 0.941 | 0.947 | 0.946 | 0.912 | 0.981 | 0.883 | 0.987 | 0.719 | 0.769 | 0.702 | 0.833 | 0.571 | 0.423 | 0.812 | 0.774 | 0.800 | 0.774 | 0.778 | 0.769 | 0.542 | 0.776 |
| | HS-578T | 0.891 | 0.928 | 0.818 | 0.982 | 0.654 | 0.718 | 0.972 | 0.787 | 0.868 | 0.639 | 0.971 | 0.308 | 0.404 | 0.744 | 0.809 | 0.880 | 0.678 | 0.971 | 0.385 | 0.476 | 0.753 |
| | MCF-7 | 0.829 | 0.846 | 0.827 | 0.846 | 0.809 | 0.654 | 0.912 | 0.757 | 0.783 | 0.753 | 0.788 | 0.718 | 0.507 | 0.830 | 0.749 | 0.773 | 0.746 | 0.769 | 0.723 | 0.492 | 0.826 |
| | MDA-MB-231 | 0.880 | 0.910 | 0.857 | 0.943 | 0.771 | 0.738 | 0.951 | 0.800 | 0.850 | 0.766 | 0.889 | 0.643 | 0.556 | 0.863 | 0.785 | 0.839 | 0.750 | 0.877 | 0.623 | 0.522 | 0.865 |
| | MDA-MB-361 | 0.983 | 0.987 | 0.982 | 0.984 | 0.980 | 0.962 | 0.997 | 0.838 | 0.885 | 0.787 | 0.958 | 0.615 | 0.638 | 0.968 | 0.917 | 0.939 | 0.896 | 0.958 | 0.833 | 0.810 | 0.976 |
| | MDA-MB-435 | 0.930 | 0.957 | 0.859 | 0.987 | 0.731 | 0.789 | 0.983 | 0.832 | 0.899 | 0.667 | 0.962 | 0.373 | 0.440 | 0.882 | 0.838 | 0.901 | 0.698 | 0.949 | 0.448 | 0.477 | 0.853 |
| | MDA-MB-453 | 0.989 | 0.989 | 0.989 | 0.983 | 0.994 | 0.977 | 1.000 | 0.750 | 0.744 | 0.750 | 0.727 | 0.773 | 0.501 | 0.880 | 0.750 | 0.732 | 0.750 | 0.682 | 0.818 | 0.505 | 0.810 |
| | MDA-MB-468 | 0.957 | 0.973 | 0.921 | 0.989 | 0.854 | 0.878 | 0.993 | 0.884 | 0.928 | 0.777 | 0.980 | 0.574 | 0.658 | 0.877 | 0.859 | 0.912 | 0.749 | 0.954 | 0.543 | 0.571 | 0.896 |
| | SK-BR-3 | 0.927 | 0.953 | 0.869 | 0.984 | 0.754 | 0.798 | 0.978 | 0.828 | 0.891 | 0.715 | 0.941 | 0.490 | 0.501 | 0.865 | 0.812 | 0.879 | 0.714 | 0.908 | 0.520 | 0.463 | 0.848 |
| | T-47D | 0.927 | 0.950 | 0.893 | 0.978 | 0.809 | 0.824 | 0.983 | 0.803 | 0.868 | 0.719 | 0.927 | 0.511 | 0.498 | 0.856 | 0.792 | 0.860 | 0.713 | 0.909 | 0.516 | 0.471 | 0.821 |

[a] ACC: Accuracy. [b] F1: F1-measure. [c] BA: Balanced accuracy. [d] SE:Sensitivity. [e] SP:Specificity. [f] MCC:Matthews correlation coefficient. [g] AUC:The area under receiver operating characteristic.

**Table S6.** The performance results of models based on PharmacoPFP.

| Methods | Cell lines | Training set | | | | | | | Validation set | | | | | | | Test set | | | | | | |
|---|---|---|---|---|---|---|---|---|---|---|---|---|---|---|---|---|---|---|---|---|---|---|
| | | ACC[a] | F1[b] | BA[c] | SE[d] | SP[e] | MCC[f] | AUC[g] | ACC | F1 | BA | SE | SP | MCC | AUC | ACC | F1 | BA | SE | SP | MCC | AUC |
| DNN | Bcap37 | 0.850 | 0.825 | 0.856 | 0.886 | 0.826 | 0.700 | 0.930 | 0.786 | 0.750 | 0.791 | 0.818 | 0.765 | 0.571 | 0.821 | 0.778 | 0.750 | 0.784 | 0.818 | 0.750 | 0.559 | 0.781 |
| | BT-20 | 0.671 | 0.748 | 0.638 | 0.832 | 0.443 | 0.302 | 0.758 | 0.862 | 0.889 | 0.846 | 0.941 | 0.750 | 0.716 | 0.922 | 0.552 | 0.649 | 0.520 | 0.706 | 0.333 | 0.042 | 0.510 |
| | BT-474 | 0.932 | 0.959 | 0.842 | 0.998 | 0.686 | 0.790 | 0.984 | 0.901 | 0.941 | 0.765 | 1.000 | 0.529 | 0.686 | 0.936 | 0.802 | 0.884 | 0.594 | 0.953 | 0.235 | 0.273 | 0.809 |
| | BT-549 | 0.832 | 0.882 | 0.777 | 0.934 | 0.620 | 0.603 | 0.913 | 0.788 | 0.854 | 0.719 | 0.913 | 0.526 | 0.488 | 0.800 | 0.712 | 0.800 | 0.636 | 0.850 | 0.421 | 0.298 | 0.682 |
| | HBL-100 | 0.905 | 0.920 | 0.899 | 0.939 | 0.858 | 0.805 | 0.970 | 0.781 | 0.811 | 0.774 | 0.833 | 0.714 | 0.553 | 0.863 | 0.774 | 0.811 | 0.763 | 0.833 | 0.692 | 0.533 | 0.795 |
| | HS-578T | 0.845 | 0.901 | 0.739 | 0.978 | 0.500 | 0.592 | 0.905 | 0.851 | 0.907 | 0.731 | 1.000 | 0.462 | 0.619 | 0.837 | 0.809 | 0.877 | 0.701 | 0.941 | 0.462 | 0.479 | 0.767 |
| | MCF-7 | 0.900 | 0.910 | 0.899 | 0.911 | 0.887 | 0.798 | 0.967 | 0.767 | 0.790 | 0.763 | 0.792 | 0.735 | 0.527 | 0.847 | 0.771 | 0.795 | 0.767 | 0.801 | 0.733 | 0.535 | 0.851 |
| | MDA-MB-231 | 0.934 | 0.949 | 0.927 | 0.953 | 0.902 | 0.857 | 0.984 | 0.815 | 0.857 | 0.795 | 0.868 | 0.722 | 0.596 | 0.867 | 0.808 | 0.849 | 0.793 | 0.847 | 0.739 | 0.585 | 0.868 |
| | MDA-MB-361 | 0.929 | 0.945 | 0.927 | 0.933 | 0.921 | 0.844 | 0.975 | 0.892 | 0.920 | 0.864 | 0.958 | 0.769 | 0.760 | 0.965 | 0.833 | 0.870 | 0.833 | 0.833 | 0.833 | 0.645 | 0.948 |
| | MDA-MB-435 | 0.851 | 0.912 | 0.674 | 0.992 | 0.355 | 0.518 | 0.943 | 0.809 | 0.889 | 0.589 | 0.983 | 0.194 | 0.319 | 0.819 | 0.812 | 0.890 | 0.607 | 0.975 | 0.239 | 0.341 | 0.816 |
| | MDA-MB-453 | 0.906 | 0.907 | 0.907 | 0.890 | 0.924 | 0.813 | 0.965 | 0.818 | 0.810 | 0.818 | 0.773 | 0.864 | 0.639 | 0.857 | 0.727 | 0.739 | 0.727 | 0.773 | 0.682 | 0.456 | 0.858 |
| | MDA-MB-468 | 0.906 | 0.941 | 0.812 | 0.988 | 0.636 | 0.724 | 0.959 | 0.864 | 0.917 | 0.735 | 0.980 | 0.489 | 0.592 | 0.875 | 0.843 | 0.905 | 0.693 | 0.974 | 0.413 | 0.510 | 0.876 |
| | SK-BR-3 | 0.893 | 0.926 | 0.894 | 0.892 | 0.896 | 0.740 | 0.961 | 0.842 | 0.893 | 0.810 | 0.875 | 0.745 | 0.598 | 0.872 | 0.797 | 0.857 | 0.785 | 0.809 | 0.760 | 0.522 | 0.878 |
| | T-47D | 0.804 | 0.874 | 0.693 | 0.970 | 0.415 | 0.500 | 0.889 | 0.732 | 0.831 | 0.596 | 0.936 | 0.255 | 0.269 | 0.812 | 0.783 | 0.860 | 0.672 | 0.945 | 0.398 | 0.432 | 0.807 |
| KNN | Bcap37 | 0.895 | 0.867 | 0.888 | 0.852 | 0.924 | 0.781 | 0.949 | 0.750 | 0.667 | 0.730 | 0.636 | 0.824 | 0.469 | 0.832 | 0.778 | 0.700 | 0.756 | 0.636 | 0.875 | 0.533 | 0.795 |
| | BT-20 | 0.970 | 0.974 | 0.970 | 0.971 | 0.969 | 0.938 | 0.970 | 0.759 | 0.774 | 0.770 | 0.706 | 0.833 | 0.531 | 0.770 | 0.724 | 0.789 | 0.691 | 0.882 | 0.500 | 0.421 | 0.691 |
| | BT-474 | 0.989 | 0.993 | 0.982 | 0.994 | 0.971 | 0.968 | 1.000 | 0.852 | 0.912 | 0.690 | 0.969 | 0.412 | 0.493 | 0.891 | 0.790 | 0.876 | 0.586 | 0.938 | 0.235 | 0.236 | 0.682 |
| | BT-549 | 0.982 | 0.987 | 0.978 | 0.989 | 0.968 | 0.959 | 0.978 | 0.780 | 0.841 | 0.734 | 0.863 | 0.605 | 0.483 | 0.734 | 0.720 | 0.802 | 0.656 | 0.838 | 0.474 | 0.330 | 0.656 |
| | HBL-100 | 0.889 | 0.905 | 0.886 | 0.905 | 0.868 | 0.773 | 0.955 | 0.750 | 0.778 | 0.746 | 0.778 | 0.714 | 0.492 | 0.833 | 0.839 | 0.857 | 0.840 | 0.833 | 0.846 | 0.674 | 0.848 |
| | HS-578T | 0.981 | 0.987 | 0.984 | 0.978 | 0.990 | 0.955 | 0.999 | 0.872 | 0.917 | 0.793 | 0.971 | 0.615 | 0.666 | 0.813 | 0.872 | 0.912 | 0.840 | 0.912 | 0.769 | 0.681 | 0.817 |
| | MCF-7 | 0.950 | 0.954 | 0.952 | 0.935 | 0.968 | 0.900 | 0.994 | 0.773 | 0.795 | 0.771 | 0.792 | 0.750 | 0.541 | 0.836 | 0.784 | 0.803 | 0.783 | 0.794 | 0.771 | 0.564 | 0.841 |
| | MDA-MB-231 | 0.958 | 0.967 | 0.961 | 0.951 | 0.972 | 0.912 | 0.995 | 0.810 | 0.851 | 0.793 | 0.853 | 0.734 | 0.588 | 0.861 | 0.806 | 0.848 | 0.792 | 0.845 | 0.739 | 0.582 | 0.851 |
| | MDA-MB-361 | 0.983 | 0.987 | 0.982 | 0.984 | 0.980 | 0.962 | 0.982 | 0.892 | 0.920 | 0.864 | 0.958 | 0.769 | 0.760 | 0.864 | 0.833 | 0.875 | 0.813 | 0.875 | 0.750 | 0.625 | 0.813 |
| | MDA-MB-435 | 0.875 | 0.918 | 0.843 | 0.900 | 0.787 | 0.656 | 0.932 | 0.799 | 0.869 | 0.726 | 0.856 | 0.597 | 0.438 | 0.831 | 0.812 | 0.878 | 0.735 | 0.873 | 0.597 | 0.463 | 0.809 |
| | MDA-MB-453 | 0.986 | 0.986 | 0.986 | 0.989 | 0.982 | 0.972 | 0.986 | 0.773 | 0.762 | 0.773 | 0.727 | 0.818 | 0.548 | 0.773 | 0.750 | 0.776 | 0.750 | 0.864 | 0.636 | 0.513 | 0.750 |
| | MDA-MB-468 | 0.978 | 0.986 | 0.973 | 0.982 | 0.965 | 0.939 | 0.973 | 0.869 | 0.916 | 0.804 | 0.928 | 0.681 | 0.628 | 0.804 | 0.879 | 0.920 | 0.845 | 0.908 | 0.783 | 0.671 | 0.845 |
| | SK-BR-3 | 0.955 | 0.970 | 0.937 | 0.973 | 0.901 | 0.879 | 0.937 | 0.837 | 0.895 | 0.755 | 0.921 | 0.588 | 0.545 | 0.755 | 0.772 | 0.847 | 0.708 | 0.836 | 0.580 | 0.405 | 0.708 |
| | T-47D | 0.981 | 0.986 | 0.981 | 0.981 | 0.981 | 0.955 | 0.998 | 0.771 | 0.843 | 0.699 | 0.877 | 0.521 | 0.426 | 0.812 | 0.821 | 0.874 | 0.777 | 0.886 | 0.667 | 0.564 | 0.837 |
| NB | Bcap37 | 0.745 | 0.685 | 0.737 | 0.693 | 0.780 | 0.472 | 0.785 | 0.607 | 0.560 | 0.612 | 0.636 | 0.588 | 0.219 | 0.676 | 0.704 | 0.692 | 0.722 | 0.818 | 0.625 | 0.438 | 0.804 |
| | BT-20 | 0.705 | 0.716 | 0.720 | 0.635 | 0.804 | 0.435 | 0.805 | 0.690 | 0.690 | 0.711 | 0.588 | 0.833 | 0.422 | 0.882 | 0.483 | 0.545 | 0.473 | 0.529 | 0.417 | -0.053 | 0.529 |
| | BT-474 | 0.741 | 0.826 | 0.686 | 0.781 | 0.591 | 0.333 | 0.814 | 0.778 | 0.855 | 0.708 | 0.828 | 0.588 | 0.387 | 0.784 | 0.679 | 0.783 | 0.602 | 0.734 | 0.471 | 0.181 | 0.710 |
| | BT-549 | 0.655 | 0.727 | 0.643 | 0.679 | 0.607 | 0.272 | 0.700 | 0.542 | 0.614 | 0.545 | 0.538 | 0.553 | 0.084 | 0.566 | 0.729 | 0.792 | 0.710 | 0.763 | 0.658 | 0.406 | 0.668 |
| | HBL-100 | 0.672 | 0.698 | 0.676 | 0.653 | 0.698 | 0.347 | 0.763 | 0.688 | 0.722 | 0.683 | 0.722 | 0.643 | 0.365 | 0.736 | 0.548 | 0.611 | 0.536 | 0.611 | 0.462 | 0.073 | 0.650 |
| | HS-578T | 0.723 | 0.801 | 0.684 | 0.771 | 0.596 | 0.350 | 0.761 | 0.766 | 0.831 | 0.743 | 0.794 | 0.692 | 0.459 | 0.751 | 0.681 | 0.762 | 0.661 | 0.706 | 0.615 | 0.296 | 0.681 |
| | MCF-7 | 0.570 | 0.588 | 0.572 | 0.553 | 0.591 | 0.142 | 0.599 | 0.578 | 0.592 | 0.581 | 0.552 | 0.610 | 0.161 | 0.608 | 0.577 | 0.595 | 0.579 | 0.560 | 0.597 | 0.156 | 0.600 |
| | MDA-MB-231 | 0.587 | 0.640 | 0.590 | 0.577 | 0.603 | 0.174 | 0.625 | 0.585 | 0.644 | 0.584 | 0.588 | 0.579 | 0.161 | 0.626 | 0.579 | 0.629 | 0.587 | 0.559 | 0.616 | 0.168 | 0.637 |
| | MDA-MB-361 | 0.762 | 0.814 | 0.748 | 0.793 | 0.703 | 0.486 | 0.844 | 0.757 | 0.830 | 0.689 | 0.917 | 0.462 | 0.439 | 0.806 | 0.667 | 0.727 | 0.667 | 0.667 | 0.667 | 0.316 | 0.800 |
| | MDA-MB-435 | 0.760 | 0.849 | 0.622 | 0.869 | 0.374 | 0.260 | 0.669 | 0.723 | 0.826 | 0.571 | 0.843 | 0.299 | 0.151 | 0.606 | 0.756 | 0.847 | 0.613 | 0.869 | 0.358 | 0.244 | 0.643 |
| | MDA-MB-453 | 0.693 | 0.723 | 0.691 | 0.779 | 0.602 | 0.388 | 0.792 | 0.727 | 0.760 | 0.727 | 0.864 | 0.591 | 0.472 | 0.806 | 0.705 | 0.735 | 0.705 | 0.818 | 0.591 | 0.420 | 0.776 |
| | MDA-MB-468 | 0.662 | 0.746 | 0.678 | 0.648 | 0.709 | 0.304 | 0.750 | 0.678 | 0.765 | 0.672 | 0.684 | 0.660 | 0.298 | 0.743 | 0.667 | 0.750 | 0.684 | 0.651 | 0.717 | 0.314 | 0.744 |
| | SK-BR-3 | 0.685 | 0.776 | 0.647 | 0.723 | 0.571 | 0.266 | 0.730 | 0.611 | 0.719 | 0.558 | 0.664 | 0.451 | 0.104 | 0.631 | 0.658 | 0.761 | 0.592 | 0.724 | 0.460 | 0.170 | 0.672 |
| | T-47D | 0.616 | 0.702 | 0.595 | 0.646 | 0.545 | 0.177 | 0.652 | 0.640 | 0.717 | 0.634 | 0.650 | 0.617 | 0.247 | 0.691 | 0.553 | 0.643 | 0.539 | 0.573 | 0.505 | 0.072 | 0.607 |
| RF | Bcap37 | 0.914 | 0.891 | 0.909 | 0.886 | 0.932 | 0.820 | 0.981 | 0.750 | 0.667 | 0.730 | 0.636 | 0.824 | 0.469 | 0.805 | 0.778 | 0.700 | 0.756 | 0.636 | 0.875 | 0.533 | 0.813 |
| | BT-20 | 0.970 | 0.974 | 0.970 | 0.971 | 0.969 | 0.938 | 0.996 | 0.793 | 0.833 | 0.775 | 0.882 | 0.667 | 0.569 | 0.917 | 0.690 | 0.769 | 0.650 | 0.882 | 0.417 | 0.344 | 0.775 |
| | BT-474 | 0.938 | 0.962 | 0.857 | 0.998 | 0.715 | 0.810 | 0.991 | 0.852 | 0.913 | 0.669 | 0.984 | 0.353 | 0.489 | 0.949 | 0.790 | 0.878 | 0.565 | 0.953 | 0.176 | 0.202 | 0.846 |
| | BT-549 | 0.982 | 0.987 | 0.979 | 0.987 | 0.971 | 0.959 | 0.999 | 0.822 | 0.879 | 0.751 | 0.950 | 0.553 | 0.575 | 0.866 | 0.746 | 0.830 | 0.654 | 0.913 | 0.395 | 0.369 | 0.708 |
| | HBL-100 | 0.957 | 0.962 | 0.956 | 0.959 | 0.953 | 0.911 | 0.995 | 0.750 | 0.789 | 0.738 | 0.833 | 0.643 | 0.488 | 0.867 | 0.774 | 0.811 | 0.763 | 0.833 | 0.692 | 0.533 | 0.855 |
| | HS-578T | 0.981 | 0.987 | 0.969 | 0.996 | 0.942 | 0.953 | 0.999 | 0.872 | 0.919 | 0.769 | 1.000 | 0.538 | 0.677 | 0.891 | 0.851 | 0.901 | 0.778 | 0.941 | 0.615 | 0.608 | 0.774 |
| | MCF-7 | 0.950 | 0.955 | 0.949 | 0.957 | 0.942 | 0.899 | 0.993 | 0.778 | 0.800 | 0.775 | 0.798 | 0.752 | 0.550 | 0.859 | 0.790 | 0.811 | 0.787 | 0.812 | 0.761 | 0.574 | 0.861 |
| | MDA-MB-231 | 0.958 | 0.968 | 0.954 | 0.971 | 0.937 | 0.910 | 0.994 | 0.817 | 0.861 | 0.789 | 0.891 | 0.687 | 0.596 | 0.880 | 0.807 | 0.852 | 0.783 | 0.870 | 0.697 | 0.577 | 0.876 |
| | MDA-MB-361 | 0.983 | 0.987 | 0.985 | 0.979 | 0.990 | 0.963 | 0.998 | 0.838 | 0.885 | 0.787 | 0.958 | 0.615 | 0.638 | 0.978 | 0.833 | 0.875 | 0.813 | 0.875 | 0.750 | 0.625 | 0.938 |
| | MDA-MB-435 | 0.971 | 0.981 | 0.955 | 0.983 | 0.927 | 0.914 | 0.995 | 0.838 | 0.902 | 0.693 | 0.953 | 0.433 | 0.474 | 0.839 | 0.822 | 0.891 | 0.677 | 0.936 | 0.418 | 0.421 | 0.864 |
| | MDA-MB-453 | 0.986 | 0.986 | 0.986 | 0.983 | 0.988 | 0.972 | 0.999 | 0.818 | 0.810 | 0.818 | 0.773 | 0.864 | 0.639 | 0.877 | 0.727 | 0.760 | 0.727 | 0.864 | 0.591 | 0.472 | 0.826 |
| | MDA-MB-468 | 0.983 | 0.989 | 0.971 | 0.993 | 0.949 | 0.952 | 0.998 | 0.889 | 0.930 | 0.803 | 0.967 | 0.638 | 0.675 | 0.923 | 0.889 | 0.931 | 0.791 | 0.974 | 0.609 | 0.668 | 0.920 |
| | SK-BR-3 | 0.960 | 0.973 | 0.941 | 0.979 | 0.903 | 0.892 | 0.993 | 0.823 | 0.889 | 0.699 | 0.947 | 0.451 | 0.480 | 0.885 | 0.802 | 0.870 | 0.721 | 0.882 | 0.560 | 0.454 | 0.834 |
| | T-47D | 0.981 | 0.986 | 0.974 | 0.990 | 0.959 | 0.954 | 0.998 | 0.768 | 0.846 | 0.670 | 0.914 | 0.426 | 0.398 | 0.845 | 0.831 | 0.885 | 0.768 | 0.923 | 0.613 | 0.576 | 0.873 |
| SVM | Bcap37 | 0.914 | 0.888 | 0.903 | 0.852 | 0.955 | 0.820 | 0.957 | 0.786 | 0.700 | 0.759 | 0.636 | 0.882 | 0.542 | 0.805 | 0.815 | 0.706 | 0.773 | 0.545 | 1.000 | 0.645 | 0.884 |
| | BT-20 | 0.906 | 0.918 | 0.908 | 0.898 | 0.918 | 0.809 | 0.957 | 0.724 | 0.750 | 0.728 | 0.706 | 0.750 | 0.449 | 0.833 | 0.690 | 0.757 | 0.662 | 0.824 | 0.500 | 0.344 | 0.721 |
| | BT-474 | 0.920 | 0.951 | 0.829 | 0.986 | 0.672 | 0.747 | 0.963 | 0.864 | 0.920 | 0.698 | 0.984 | 0.412 | 0.541 | 0.907 | 0.827 | 0.897 | 0.653 | 0.953 | 0.353 | 0.397 | 0.795 |
| | BT-549 | 0.867 | 0.906 | 0.822 | 0.950 | 0.695 | 0.689 | 0.941 | 0.839 | 0.889 | 0.778 | 0.950 | 0.605 | 0.618 | 0.830 | 0.754 | 0.830 | 0.681 | 0.888 | 0.474 | 0.402 | 0.705 |
| | HBL-100 | 0.901 | 0.916 | 0.897 | 0.925 | 0.868 | 0.796 | 0.943 | 0.688 | 0.737 | 0.675 | 0.778 | 0.571 | 0.358 | 0.823 | 0.742 | 0.778 | 0.735 | 0.778 | 0.692 | 0.470 | 0.859 |
| | HS-578T | 0.848 | 0.902 | 0.753 | 0.967 | 0.538 | 0.598 | 0.959 | 0.809 | 0.880 | 0.678 | 0.971 | 0.385 | 0.476 | 0.910 | 0.830 | 0.889 | 0.740 | 0.941 | 0.538 | 0.545 | 0.792 |
| | MCF-7 | 0.917 | 0.925 | 0.915 | 0.927 | 0.903 | 0.831 | 0.965 | 0.779 | 0.805 | 0.774 | 0.822 | 0.726 | 0.551 | 0.844 | 0.786 | 0.812 | 0.779 | 0.836 | 0.722 | 0.564 | 0.853 |
| | MDA-MB-231 | 0.958 | 0.967 | 0.953 | 0.971 | 0.935 | 0.909 | 0.976 | 0.807 | 0.857 | 0.771 | 0.903 | 0.638 | 0.572 | 0.850 | 0.796 | 0.847 | 0.763 | 0.885 | 0.640 | 0.549 | 0.849 |
| | MDA-MB-361 | 0.932 | 0.949 | 0.920 | 0.959 | 0.881 | 0.848 | 0.971 | 0.892 | 0.920 | 0.864 | 0.958 | 0.769 | 0.760 | 0.968 | 0.861 | 0.894 | 0.854 | 0.875 | 0.833 | 0.695 | 0.920 |
| | MDA-MB-435 | 0.835 | 0.902 | 0.659 | 0.974 | 0.344 | 0.447 | 0.913 | 0.799 | 0.882 | 0.588 | 0.966 | 0.209 | 0.280 | 0.760 | 0.815 | 0.892 | 0.609 | 0.979 | 0.239 | 0.356 | 0.805 |
| | MDA-MB-453 | 0.986 | 0.986 | 0.986 | 0.983 | 0.988 | 0.972 | 0.994 | 0.818 | 0.789 | 0.818 | 0.682 | 0.955 | 0.661 | 0.860 | 0.727 | 0.727 | 0.727 | 0.727 | 0.727 | 0.455 | 0.809 |
| | MDA-MB-468 | 0.899 | 0.937 | 0.803 | 0.984 | 0.623 | 0.703 | 0.953 | 0.864 | 0.916 | 0.742 | 0.974 | 0.511 | 0.592 | 0.867 | 0.854 | 0.911 | 0.715 | 0.974 | 0.457 | 0.547 | 0.869 |
| | SK-BR-3 | 0.851 | 0.908 | 0.725 | 0.976 | 0.474 | 0.568 | 0.928 | 0.823 | 0.891 | 0.680 | 0.967 | 0.392 | 0.474 | 0.842 | 0.802 | 0.877 | 0.667 | 0.934 | 0.400 | 0.406 | 0.834 |
| | T-47D | 0.860 | 0.905 | 0.800 | 0.948 | 0.653 | 0.651 | 0.926 | 0.768 | 0.846 | 0.673 | 0.909 | 0.436 | 0.400 | 0.798 | 0.815 | 0.873 | 0.753 | 0.905 | 0.602 | 0.538 | 0.830 |
| XGBoost | Bcap37 | 0.914 | 0.890 | 0.907 | 0.875 | 0.939 | 0.819 | 0.980 | 0.786 | 0.700 | 0.759 | 0.636 | 0.882 | 0.542 | 0.799 | 0.815 | 0.783 | 0.815 | 0.818 | 0.813 | 0.624 | 0.787 |
| | BT-20 | 0.962 | 0.967 | 0.961 | 0.964 | 0.959 | 0.921 | 0.993 | 0.759 | 0.774 | 0.770 | 0.706 | 0.833 | 0.531 | 0.902 | 0.724 | 0.778 | 0.703 | 0.824 | 0.583 | 0.422 | 0.819 |
| | BT-474 | 0.978 | 0.986 | 0.952 | 0.998 | 0.905 | 0.935 | 0.998 | 0.864 | 0.917 | 0.741 | 0.953 | 0.529 | 0.553 | 0.902 | 0.790 | 0.874 | 0.608 | 0.922 | 0.294 | 0.267 | 0.832 |
| | BT-549 | 0.979 | 0.984 | 0.975 | 0.986 | 0.964 | 0.952 | 0.998 | 0.814 | 0.872 | 0.745 | 0.938 | 0.553 | 0.553 | 0.794 | 0.729 | 0.814 | 0.648 | 0.875 | 0.421 | 0.334 | 0.694 |
| | HBL-100 | 0.953 | 0.959 | 0.950 | 0.966 | 0.934 | 0.902 | 0.994 | 0.813 | 0.833 | 0.810 | 0.833 | 0.786 | 0.619 | 0.855 | 0.806 | 0.842 | 0.791 | 0.889 | 0.692 | 0.599 | 0.829 |
| | HS-578T | 0.816 | 0.886 | 0.677 | 0.989 | 0.365 | 0.508 | 0.878 | 0.766 | 0.861 | 0.577 | 1.000 | 0.154 | 0.341 | 0.805 | 0.787 | 0.868 | 0.639 | 0.971 | 0.308 | 0.404 | 0.750 |
| | MCF-7 | 0.804 | 0.824 | 0.801 | 0.828 | 0.774 | 0.603 | 0.889 | 0.738 | 0.765 | 0.733 | 0.771 | 0.696 | 0.468 | 0.815 | 0.744 | 0.769 | 0.742 | 0.766 | 0.717 | 0.483 | 0.813 |
| | MDA-MB-231 | 0.865 | 0.897 | 0.844 | 0.920 | 0.769 | 0.704 | 0.941 | 0.793 | 0.844 | 0.761 | 0.878 | 0.643 | 0.541 | 0.856 | 0.782 | 0.834 | 0.754 | 0.856 | 0.653 | 0.520 | 0.855 |
| | MDA-MB-361 | 0.949 | 0.961 | 0.949 | 0.948 | 0.950 | 0.889 | 0.981 | 0.865 | 0.894 | 0.861 | 0.875 | 0.846 | 0.710 | 0.958 | 0.833 | 0.864 | 0.854 | 0.792 | 0.917 | 0.672 | 0.924 |
| | MDA-MB-435 | 0.866 | 0.919 | 0.722 | 0.980 | 0.464 | 0.570 | 0.936 | 0.822 | 0.894 | 0.640 | 0.966 | 0.313 | 0.394 | 0.825 | 0.828 | 0.898 | 0.655 | 0.966 | 0.343 | 0.424 | 0.824 |
| | MDA-MB-453 | 0.986 | 0.986 | 0.986 | 0.983 | 0.988 | 0.972 | 0.999 | 0.773 | 0.762 | 0.773 | 0.727 | 0.818 | 0.548 | 0.835 | 0.750 | 0.766 | 0.750 | 0.818 | 0.682 | 0.505 | 0.811 |
| | MDA-MB-468 | 0.979 | 0.986 | 0.963 | 0.993 | 0.933 | 0.940 | 0.997 | 0.864 | 0.914 | 0.772 | 0.947 | 0.596 | 0.599 | 0.872 | 0.874 | 0.921 | 0.781 | 0.954 | 0.609 | 0.623 | 0.906 |
| | SK-BR-3 | 0.923 | 0.950 | 0.867 | 0.978 | 0.757 | 0.787 | 0.973 | 0.833 | 0.896 | 0.699 | 0.967 | 0.431 | 0.509 | 0.859 | 0.787 | 0.862 | 0.691 | 0.882 | 0.500 | 0.402 | 0.853 |
| | T-47D | 0.930 | 0.951 | 0.902 | 0.970 | 0.834 | 0.830 | 0.983 | 0.777 | 0.850 | 0.695 | 0.900 | 0.489 | 0.433 | 0.843 | 0.805 | 0.864 | 0.753 | 0.882 | 0.624 | 0.521 | 0.850 |

[a] ACC: Accuracy. [b] F1: F1-measure. [c] BA: Balanced accuracy. [d] SE:Sensitivity. [e] SP:Specificity. [f] MCC:Matthews correlation coefficient. [g] AUC:The area under receiver operating characteristic.

**Table S7.** The performance results of models based on molecular graph.

| Methods | Cell lines | Training set | | | | | | | Validation set | | | | | | | Test set | | | | | | |
|---|---|---|---|---|---|---|---|---|---|---|---|---|---|---|---|---|---|---|---|---|---|---|
| | | ACC[a] | F1[b] | BA[c] | SE[d] | SP[e] | MCC[f] | AUC[g] | ACC | F1 | BA | SE | SP | MCC | AUC | ACC | F1 | BA | SE | SP | MCC | AUC |
| Attentive FP | Bcap37 | 0.836 | 0.788 | 0.824 | 0.761 | 0.886 | 0.656 | 0.918 | 0.821 | 0.737 | 0.789 | 0.636 | 0.941 | 0.624 | 0.882 | 0.778 | 0.727 | 0.770 | 0.727 | 0.813 | 0.540 | 0.858 |
| | BT-20 | 0.868 | 0.877 | 0.879 | 0.810 | 0.948 | 0.748 | 0.953 | 0.793 | 0.813 | 0.799 | 0.765 | 0.833 | 0.589 | 0.887 | 0.690 | 0.743 | 0.674 | 0.765 | 0.583 | 0.353 | 0.735 |
| | BT-474 | 0.874 | 0.922 | 0.770 | 0.949 | 0.591 | 0.594 | 0.903 | 0.889 | 0.930 | 0.822 | 0.938 | 0.706 | 0.658 | 0.902 | 0.778 | 0.864 | 0.622 | 0.891 | 0.353 | 0.270 | 0.787 |
| | BT-549 | 0.720 | 0.826 | 0.576 | 0.989 | 0.162 | 0.298 | 0.741 | 0.712 | 0.817 | 0.580 | 0.950 | 0.211 | 0.248 | 0.662 | 0.729 | 0.832 | 0.586 | 0.988 | 0.184 | 0.319 | 0.630 |
| | HBL-100 | 0.704 | 0.780 | 0.665 | 0.905 | 0.425 | 0.384 | 0.757 | 0.688 | 0.750 | 0.667 | 0.833 | 0.500 | 0.357 | 0.718 | 0.613 | 0.700 | 0.581 | 0.778 | 0.385 | 0.177 | 0.645 |
| | HS-578T | 0.821 | 0.889 | 0.684 | 0.993 | 0.375 | 0.527 | 0.844 | 0.766 | 0.857 | 0.601 | 0.971 | 0.231 | 0.323 | 0.851 | 0.809 | 0.880 | 0.678 | 0.971 | 0.385 | 0.476 | 0.830 |
| | MCF-7 | 0.918 | 0.927 | 0.915 | 0.944 | 0.885 | 0.833 | 0.977 | 0.767 | 0.795 | 0.761 | 0.813 | 0.709 | 0.526 | 0.847 | 0.764 | 0.792 | 0.759 | 0.807 | 0.710 | 0.521 | 0.845 |
| | MDA-MB-231 | 0.947 | 0.958 | 0.946 | 0.950 | 0.942 | 0.887 | 0.990 | 0.808 | 0.849 | 0.794 | 0.846 | 0.741 | 0.586 | 0.868 | 0.805 | 0.845 | 0.796 | 0.831 | 0.761 | 0.585 | 0.870 |
| | MDA-MB-361 | 0.932 | 0.948 | 0.929 | 0.938 | 0.921 | 0.851 | 0.981 | 0.838 | 0.889 | 0.769 | 1.000 | 0.538 | 0.656 | 0.949 | 0.861 | 0.894 | 0.854 | 0.875 | 0.833 | 0.695 | 0.938 |
| | MDA-MB-435 | 0.923 | 0.951 | 0.870 | 0.966 | 0.774 | 0.770 | 0.975 | 0.835 | 0.896 | 0.734 | 0.915 | 0.552 | 0.496 | 0.853 | 0.835 | 0.897 | 0.723 | 0.924 | 0.522 | 0.487 | 0.824 |
| | MDA-MB-453 | 0.875 | 0.888 | 0.872 | 0.961 | 0.784 | 0.760 | 0.955 | 0.705 | 0.745 | 0.705 | 0.864 | 0.545 | 0.432 | 0.822 | 0.773 | 0.792 | 0.773 | 0.864 | 0.682 | 0.555 | 0.872 |
| | MDA-MB-468 | 0.941 | 0.961 | 0.936 | 0.947 | 0.925 | 0.844 | 0.989 | 0.839 | 0.897 | 0.755 | 0.914 | 0.596 | 0.536 | 0.874 | 0.884 | 0.924 | 0.841 | 0.921 | 0.761 | 0.677 | 0.875 |
| | SK-BR-3 | 0.823 | 0.892 | 0.675 | 0.970 | 0.380 | 0.469 | 0.838 | 0.818 | 0.889 | 0.663 | 0.974 | 0.353 | 0.456 | 0.835 | 0.822 | 0.888 | 0.700 | 0.941 | 0.460 | 0.474 | 0.805 |
| | T-47D | 0.947 | 0.963 | 0.930 | 0.972 | 0.889 | 0.872 | 0.987 | 0.787 | 0.853 | 0.720 | 0.886 | 0.553 | 0.468 | 0.820 | 0.789 | 0.852 | 0.738 | 0.864 | 0.613 | 0.486 | 0.812 |
| GAT | Bcap37 | 0.732 | 0.638 | 0.708 | 0.591 | 0.826 | 0.431 | 0.798 | 0.714 | 0.600 | 0.684 | 0.545 | 0.824 | 0.386 | 0.759 | 0.741 | 0.667 | 0.724 | 0.636 | 0.813 | 0.457 | 0.767 |
| | BT-20 | 0.786 | 0.830 | 0.765 | 0.891 | 0.639 | 0.555 | 0.838 | 0.793 | 0.813 | 0.799 | 0.765 | 0.833 | 0.589 | 0.863 | 0.690 | 0.757 | 0.662 | 0.824 | 0.500 | 0.344 | 0.721 |
| | BT-474 | 0.838 | 0.901 | 0.708 | 0.934 | 0.482 | 0.469 | 0.818 | 0.815 | 0.885 | 0.688 | 0.906 | 0.471 | 0.406 | 0.849 | 0.728 | 0.833 | 0.547 | 0.859 | 0.235 | 0.105 | 0.657 |
| | BT-549 | 0.793 | 0.863 | 0.698 | 0.970 | 0.425 | 0.507 | 0.894 | 0.703 | 0.804 | 0.595 | 0.900 | 0.289 | 0.241 | 0.712 | 0.746 | 0.831 | 0.647 | 0.925 | 0.368 | 0.365 | 0.710 |
| | HBL-100 | 0.739 | 0.805 | 0.703 | 0.925 | 0.481 | 0.466 | 0.821 | 0.781 | 0.837 | 0.750 | 1.000 | 0.500 | 0.600 | 0.833 | 0.581 | 0.683 | 0.543 | 0.778 | 0.308 | 0.096 | 0.641 |
| | HS-578T | 0.819 | 0.880 | 0.738 | 0.919 | 0.558 | 0.521 | 0.853 | 0.851 | 0.904 | 0.755 | 0.971 | 0.538 | 0.606 | 0.862 | 0.851 | 0.901 | 0.778 | 0.941 | 0.615 | 0.608 | 0.758 |
| | MCF-7 | 0.756 | 0.774 | 0.757 | 0.753 | 0.760 | 0.511 | 0.835 | 0.733 | 0.755 | 0.732 | 0.741 | 0.722 | 0.462 | 0.801 | 0.721 | 0.742 | 0.721 | 0.723 | 0.718 | 0.439 | 0.800 |
| | MDA-MB-231 | 0.743 | 0.823 | 0.672 | 0.934 | 0.410 | 0.420 | 0.815 | 0.723 | 0.809 | 0.648 | 0.922 | 0.374 | 0.366 | 0.801 | 0.724 | 0.807 | 0.656 | 0.905 | 0.406 | 0.369 | 0.770 |
| | MDA-MB-361 | 0.891 | 0.914 | 0.896 | 0.881 | 0.911 | 0.770 | 0.956 | 0.892 | 0.923 | 0.846 | 1.000 | 0.692 | 0.770 | 0.955 | 0.861 | 0.889 | 0.875 | 0.833 | 0.917 | 0.717 | 0.896 |
| | MDA-MB-435 | 0.842 | 0.904 | 0.695 | 0.958 | 0.432 | 0.483 | 0.892 | 0.802 | 0.882 | 0.611 | 0.953 | 0.269 | 0.313 | 0.826 | 0.838 | 0.903 | 0.672 | 0.970 | 0.373 | 0.464 | 0.830 |
| | MDA-MB-453 | 0.858 | 0.848 | 0.861 | 0.768 | 0.953 | 0.731 | 0.948 | 0.750 | 0.744 | 0.750 | 0.727 | 0.773 | 0.501 | 0.829 | 0.750 | 0.703 | 0.750 | 0.591 | 0.909 | 0.527 | 0.812 |
| | MDA-MB-468 | 0.894 | 0.934 | 0.792 | 0.983 | 0.601 | 0.685 | 0.947 | 0.869 | 0.920 | 0.745 | 0.980 | 0.511 | 0.609 | 0.877 | 0.869 | 0.919 | 0.748 | 0.974 | 0.522 | 0.600 | 0.875 |
| | SK-BR-3 | 0.869 | 0.914 | 0.808 | 0.929 | 0.687 | 0.638 | 0.913 | 0.862 | 0.911 | 0.784 | 0.941 | 0.627 | 0.614 | 0.863 | 0.812 | 0.875 | 0.748 | 0.875 | 0.620 | 0.495 | 0.840 |
| | T-47D | 0.864 | 0.908 | 0.798 | 0.962 | 0.634 | 0.663 | 0.929 | 0.752 | 0.839 | 0.637 | 0.923 | 0.351 | 0.343 | 0.755 | 0.748 | 0.830 | 0.659 | 0.877 | 0.441 | 0.353 | 0.763 |
| GCN | Bcap37 | 0.791 | 0.753 | 0.792 | 0.795 | 0.788 | 0.575 | 0.834 | 0.786 | 0.727 | 0.775 | 0.727 | 0.824 | 0.551 | 0.850 | 0.741 | 0.696 | 0.739 | 0.727 | 0.750 | 0.472 | 0.693 |
| | BT-20 | 0.829 | 0.832 | 0.851 | 0.723 | 0.979 | 0.698 | 0.965 | 0.724 | 0.714 | 0.752 | 0.588 | 0.917 | 0.512 | 0.868 | 0.655 | 0.688 | 0.657 | 0.647 | 0.667 | 0.309 | 0.740 |
| | BT-474 | 0.961 | 0.976 | 0.925 | 0.988 | 0.861 | 0.882 | 0.990 | 0.889 | 0.932 | 0.778 | 0.969 | 0.588 | 0.638 | 0.901 | 0.790 | 0.870 | 0.651 | 0.891 | 0.412 | 0.326 | 0.866 |
| | BT-549 | 0.873 | 0.914 | 0.809 | 0.994 | 0.623 | 0.713 | 0.975 | 0.814 | 0.876 | 0.724 | 0.975 | 0.474 | 0.559 | 0.890 | 0.763 | 0.839 | 0.680 | 0.913 | 0.447 | 0.418 | 0.669 |
| | HBL-100 | 0.889 | 0.908 | 0.878 | 0.946 | 0.811 | 0.773 | 0.975 | 0.813 | 0.842 | 0.802 | 0.889 | 0.714 | 0.618 | 0.841 | 0.677 | 0.762 | 0.637 | 0.889 | 0.385 | 0.323 | 0.658 |
| | HS-578T | 0.733 | 0.844 | 0.519 | 1.000 | 0.038 | 0.168 | 0.746 | 0.723 | 0.840 | 0.500 | 1.000 | 0.000 | nan | 0.796 | 0.723 | 0.840 | 0.500 | 1.000 | 0.000 | nan | 0.636 |
| | MCF-7 | 0.828 | 0.839 | 0.830 | 0.808 | 0.852 | 0.656 | 0.912 | 0.754 | 0.771 | 0.755 | 0.746 | 0.765 | 0.508 | 0.834 | 0.752 | 0.765 | 0.754 | 0.728 | 0.781 | 0.506 | 0.833 |
| | MDA-MB-231 | 0.903 | 0.925 | 0.888 | 0.944 | 0.831 | 0.788 | 0.968 | 0.801 | 0.848 | 0.775 | 0.868 | 0.682 | 0.562 | 0.870 | 0.797 | 0.845 | 0.771 | 0.866 | 0.677 | 0.554 | 0.859 |
| | MDA-MB-361 | 0.884 | 0.909 | 0.888 | 0.876 | 0.901 | 0.756 | 0.952 | 0.838 | 0.875 | 0.822 | 0.875 | 0.769 | 0.644 | 0.929 | 0.917 | 0.939 | 0.896 | 0.958 | 0.833 | 0.810 | 0.955 |
| | MDA-MB-435 | 0.944 | 0.964 | 0.926 | 0.958 | 0.893 | 0.840 | 0.981 | 0.815 | 0.882 | 0.726 | 0.886 | 0.567 | 0.458 | 0.855 | 0.828 | 0.892 | 0.730 | 0.907 | 0.552 | 0.481 | 0.858 |
| | MDA-MB-453 | 0.906 | 0.916 | 0.904 | 0.994 | 0.813 | 0.824 | 0.996 | 0.705 | 0.755 | 0.705 | 0.909 | 0.500 | 0.448 | 0.876 | 0.682 | 0.750 | 0.682 | 0.955 | 0.409 | 0.434 | 0.866 |
| | MDA-MB-468 | 0.955 | 0.971 | 0.929 | 0.977 | 0.881 | 0.872 | 0.987 | 0.899 | 0.937 | 0.809 | 0.980 | 0.638 | 0.706 | 0.917 | 0.848 | 0.904 | 0.757 | 0.928 | 0.587 | 0.552 | 0.887 |
| | SK-BR-3 | 0.878 | 0.913 | 0.905 | 0.851 | 0.958 | 0.733 | 0.974 | 0.808 | 0.860 | 0.826 | 0.789 | 0.863 | 0.585 | 0.898 | 0.757 | 0.819 | 0.785 | 0.730 | 0.840 | 0.500 | 0.839 |
| | T-47D | 0.895 | 0.929 | 0.835 | 0.984 | 0.686 | 0.744 | 0.968 | 0.774 | 0.850 | 0.680 | 0.914 | 0.447 | 0.417 | 0.832 | 0.780 | 0.852 | 0.697 | 0.900 | 0.495 | 0.437 | 0.819 |
| MPNN | Bcap37 | 0.800 | 0.728 | 0.778 | 0.670 | 0.886 | 0.577 | 0.882 | 0.714 | 0.556 | 0.668 | 0.455 | 0.882 | 0.380 | 0.866 | 0.778 | 0.700 | 0.756 | 0.636 | 0.875 | 0.533 | 0.807 |
| | BT-20 | 0.838 | 0.862 | 0.831 | 0.869 | 0.794 | 0.665 | 0.908 | 0.793 | 0.813 | 0.799 | 0.765 | 0.833 | 0.589 | 0.902 | 0.759 | 0.811 | 0.733 | 0.882 | 0.583 | 0.496 | 0.784 |
| | BT-474 | 0.938 | 0.962 | 0.878 | 0.982 | 0.774 | 0.808 | 0.973 | 0.852 | 0.910 | 0.712 | 0.953 | 0.471 | 0.504 | 0.919 | 0.802 | 0.881 | 0.637 | 0.922 | 0.353 | 0.327 | 0.847 |
| | BT-549 | 0.892 | 0.922 | 0.867 | 0.939 | 0.795 | 0.751 | 0.943 | 0.797 | 0.852 | 0.760 | 0.863 | 0.658 | 0.528 | 0.809 | 0.712 | 0.795 | 0.649 | 0.825 | 0.474 | 0.314 | 0.634 |
| | HBL-100 | 0.684 | 0.758 | 0.652 | 0.850 | 0.453 | 0.334 | 0.715 | 0.750 | 0.810 | 0.722 | 0.944 | 0.500 | 0.509 | 0.790 | 0.645 | 0.732 | 0.609 | 0.833 | 0.385 | 0.246 | 0.701 |
| | HS-578T | 0.723 | 0.839 | 0.500 | 1.000 | 0.000 | nan | 0.646 | 0.723 | 0.840 | 0.500 | 1.000 | 0.000 | nan | 0.681 | 0.723 | 0.840 | 0.500 | 1.000 | 0.000 | nan | 0.665 |
| | MCF-7 | 0.842 | 0.859 | 0.838 | 0.871 | 0.805 | 0.679 | 0.925 | 0.759 | 0.787 | 0.754 | 0.803 | 0.705 | 0.510 | 0.835 | 0.766 | 0.791 | 0.763 | 0.795 | 0.730 | 0.526 | 0.843 |
| | MDA-MB-231 | 0.879 | 0.908 | 0.859 | 0.932 | 0.786 | 0.736 | 0.953 | 0.796 | 0.846 | 0.767 | 0.874 | 0.660 | 0.551 | 0.859 | 0.794 | 0.842 | 0.768 | 0.863 | 0.672 | 0.547 | 0.850 |
| | MDA-MB-361 | 0.898 | 0.920 | 0.901 | 0.891 | 0.911 | 0.783 | 0.964 | 0.865 | 0.902 | 0.825 | 0.958 | 0.692 | 0.699 | 0.962 | 0.917 | 0.933 | 0.938 | 0.875 | 1.000 | 0.837 | 0.972 |
| | MDA-MB-435 | 0.847 | 0.906 | 0.711 | 0.953 | 0.469 | 0.505 | 0.895 | 0.799 | 0.874 | 0.678 | 0.894 | 0.463 | 0.381 | 0.795 | 0.809 | 0.884 | 0.642 | 0.941 | 0.343 | 0.360 | 0.749 |
| | MDA-MB-453 | 0.869 | 0.873 | 0.869 | 0.873 | 0.865 | 0.738 | 0.939 | 0.795 | 0.809 | 0.795 | 0.864 | 0.727 | 0.596 | 0.847 | 0.636 | 0.667 | 0.636 | 0.727 | 0.545 | 0.277 | 0.715 |
| | MDA-MB-468 | 0.936 | 0.959 | 0.899 | 0.968 | 0.830 | 0.817 | 0.978 | 0.910 | 0.942 | 0.853 | 0.961 | 0.745 | 0.741 | 0.906 | 0.879 | 0.921 | 0.830 | 0.921 | 0.739 | 0.660 | 0.858 |
| | SK-BR-3 | 0.809 | 0.883 | 0.653 | 0.963 | 0.342 | 0.417 | 0.790 | 0.803 | 0.881 | 0.634 | 0.974 | 0.294 | 0.399 | 0.759 | 0.807 | 0.883 | 0.644 | 0.967 | 0.320 | 0.406 | 0.760 |
| | T-47D | 0.815 | 0.874 | 0.752 | 0.909 | 0.595 | 0.539 | 0.867 | 0.768 | 0.844 | 0.682 | 0.895 | 0.468 | 0.406 | 0.800 | 0.722 | 0.811 | 0.635 | 0.850 | 0.419 | 0.292 | 0.751 |

[a] ACC: Accuracy. [b] F1: F1-measure. [c] BA: Balanced accuracy. [d] SE:Sensitivity. [e] SP:Specificity. [f] MCC:Matthews correlation coefficient. [g] AUC:The area under receiver operating characteristic.

**Table S8.** AUC results for Multi-task models.

| Datasets | Based on DNN::Morgan | Based on graph::GCN | Based on graph::Attentive FP |
|---|---|---|---|
| Bcap37 | 0.508 | 0.616 | 0.571 |
| BT-20 | 0.414 | 0.464 | 0.520 |
| BT-474 | 0.493 | 0.579 | 0.385 |
| BT-549 | 0.546 | 0.502 | 0.432 |
| HS-578T | 0.530 | 0.512 | 0.541 |
| MCF-7 | 0.519 | 0.536 | 0.472 |
| MDA-MB-231 | 0.460 | 0.553 | 0.440 |
| MDA-MB-361 | 0.359 | 0.600 | 0.654 |
| MDA-MB-435 | 0.581 | 0.483 | 0.555 |
| MDA-MB-453 | 0.408 | 0.620 | 0.425 |
| MDA-MB-468 | 0.495 | 0.429 | 0.536 |
| SK-BR-3 | 0.503 | 0.410 | 0.363 |
| T-47D | 0.565 | 0.563 | 0.447 |

**Table S9.** The optimal *in silico* predictive model for each breast cell line.

| Cell lines | The optional model | Training set | | | | | | | Validation set | | | | | | | Test set | | | | | | |
|---|---|---|---|---|---|---|---|---|---|---|---|---|---|---|---|---|---|---|---|---|---|---|
| | | ACC[a] | F1[b] | BA[c] | SE[d] | SP[e] | MCC[f] | AUC[g] | ACC | F1 | BA | SE | SP | MCC | AUC | ACC | F1 | BA | SE | SP | MCC | AUC |
| Bcap37 | KNN::AtomPairs | 0.995 | 0.994 | 0.994 | 0.989 | 1.000 | 0.991 | 1.000 | 0.750 | 0.696 | 0.746 | 0.727 | 0.765 | 0.486 | 0.837 | 0.852 | 0.800 | 0.832 | 0.727 | 0.938 | 0.693 | 0.869 |
| BT-20 | XGBoost::MACCS | 0.983 | 0.986 | 0.981 | 0.993 | 0.969 | 0.965 | 0.999 | 0.862 | 0.882 | 0.858 | 0.882 | 0.833 | 0.716 | 0.897 | 0.828 | 0.857 | 0.816 | 0.882 | 0.750 | 0.642 | 0.833 |
| BT-474 | RF::MACCS | 0.997 | 0.998 | 0.995 | 0.998 | 0.993 | 0.991 | 1.000 | 0.901 | 0.940 | 0.786 | 0.984 | 0.588 | 0.681 | 0.949 | 0.877 | 0.923 | 0.792 | 0.938 | 0.647 | 0.613 | 0.900 |
| BT-549 | SVM::MACCS | 0.863 | 0.904 | 0.812 | 0.958 | 0.666 | 0.679 | 0.946 | 0.763 | 0.841 | 0.673 | 0.925 | 0.421 | 0.415 | 0.762 | 0.797 | 0.857 | 0.739 | 0.900 | 0.579 | 0.514 | 0.689 |
| HBL-100 | RF::Morgan | 0.996 | 0.997 | 0.995 | 1.000 | 0.991 | 0.992 | 1.000 | 0.781 | 0.821 | 0.766 | 0.889 | 0.643 | 0.555 | 0.857 | 0.839 | 0.872 | 0.818 | 0.944 | 0.692 | 0.672 | 0.823 |
| HS-578T | RF::MACCS | 0.971 | 0.980 | 0.968 | 0.974 | 0.962 | 0.928 | 0.996 | 0.787 | 0.853 | 0.734 | 0.853 | 0.615 | 0.468 | 0.813 | 0.894 | 0.928 | 0.855 | 0.941 | 0.769 | 0.729 | 0.819 |
| MCF-7 | SVM::Morgan | 0.980 | 0.982 | 0.979 | 0.982 | 0.977 | 0.959 | 0.996 | 0.810 | 0.831 | 0.807 | 0.839 | 0.774 | 0.615 | 0.876 | 0.810 | 0.831 | 0.807 | 0.840 | 0.774 | 0.615 | 0.872 |
| MDA-MB-231 | RF::Morgan | 0.995 | 0.996 | 0.995 | 0.996 | 0.993 | 0.989 | 1.000 | 0.836 | 0.876 | 0.807 | 0.910 | 0.704 | 0.638 | 0.894 | 0.837 | 0.876 | 0.811 | 0.905 | 0.717 | 0.640 | 0.904 |
| MDA-MB-361 | RF::MACCS | 0.993 | 0.995 | 0.992 | 0.995 | 0.990 | 0.985 | 0.999 | 0.784 | 0.846 | 0.728 | 0.917 | 0.538 | 0.506 | 0.933 | 0.917 | 0.939 | 0.896 | 0.958 | 0.833 | 0.810 | 0.993 |
| MDA-MB-435 | RF::AtomPairs | 0.996 | 0.998 | 0.996 | 0.996 | 0.996 | 0.989 | 1.000 | 0.832 | 0.899 | 0.673 | 0.958 | 0.388 | 0.443 | 0.897 | 0.848 | 0.908 | 0.710 | 0.958 | 0.463 | 0.510 | 0.870 |
| MDA-MB-453 | RF::AtomPairs | 0.994 | 0.994 | 0.994 | 0.994 | 0.994 | 0.989 | 1.000 | 0.864 | 0.857 | 0.864 | 0.818 | 0.909 | 0.730 | 0.928 | 0.841 | 0.844 | 0.841 | 0.864 | 0.818 | 0.683 | 0.888 |
| MDA-MB-468 | XGBoost::MACCS | 0.977 | 0.985 | 0.964 | 0.988 | 0.941 | 0.935 | 0.997 | 0.874 | 0.921 | 0.778 | 0.961 | 0.596 | 0.628 | 0.901 | 0.899 | 0.937 | 0.813 | 0.974 | 0.652 | 0.701 | 0.909 |
| SK-BR-3 | DNN::AtomPairs | 0.948 | 0.967 | 0.896 | 1.000 | 0.792 | 0.861 | 0.994 | 0.852 | 0.909 | 0.725 | 0.980 | 0.471 | 0.576 | 0.896 | 0.842 | 0.901 | 0.727 | 0.954 | 0.500 | 0.537 | 0.860 |
| T-47D | RF::Morgan | 0.998 | 0.998 | 0.997 | 0.998 | 0.996 | 0.994 | 1.000 | 0.790 | 0.860 | 0.701 | 0.923 | 0.479 | 0.462 | 0.862 | 0.840 | 0.892 | 0.772 | 0.941 | 0.602 | 0.599 | 0.885 |

[a] ACC: Accuracy. [b] F1: F1-measure. [c] BA: Balanced accuracy. [d] SE:Sensitivity. [e] SP:Specificity. [f] MCC:Matthews correlation coefficient. [g] AUC:The area under receiver operating characteristic.

**Table S10.** The performance results of voting models based on Morgan fingerprints.

| Methods | Cell lines | Training set | | | | | | | Validation set | | | | | | | Test set | | | | | | |
|---|---|---|---|---|---|---|---|---|---|---|---|---|---|---|---|---|---|---|---|---|---|---|
| | | ACC[a] | F1[b] | BA[c] | SE[d] | SP[e] | MCC[f] | AUC[g] | ACC | F1 | BA | SE | SP | MCC | AUC | ACC | F1 | BA | SE | SP | MCC | AUC |
| RF+SVM | Bcap37 | 0.964 | 0.953 | 0.958 | 0.932 | 0.985 | 0.924 | 0.996 | 0.786 | 0.700 | 0.759 | 0.636 | 0.882 | 0.542 | 0.802 | 0.741 | 0.667 | 0.724 | 0.636 | 0.813 | 0.457 | 0.739 |
| | BT-20 | 0.962 | 0.967 | 0.960 | 0.971 | 0.948 | 0.921 | 0.998 | 0.793 | 0.813 | 0.799 | 0.765 | 0.833 | 0.589 | 0.892 | 0.724 | 0.778 | 0.703 | 0.824 | 0.583 | 0.422 | 0.843 |
| | BT-474 | 0.975 | 0.985 | 0.950 | 0.994 | 0.905 | 0.925 | 0.999 | 0.877 | 0.924 | 0.771 | 0.953 | 0.588 | 0.601 | 0.911 | 0.802 | 0.881 | 0.637 | 0.922 | 0.353 | 0.327 | 0.865 |
| | BT-549 | 0.977 | 0.983 | 0.969 | 0.991 | 0.948 | 0.947 | 0.999 | 0.831 | 0.884 | 0.764 | 0.950 | 0.579 | 0.596 | 0.873 | 0.763 | 0.835 | 0.694 | 0.888 | 0.500 | 0.426 | 0.708 |
| | HBL-100 | 0.968 | 0.973 | 0.969 | 0.966 | 0.972 | 0.935 | 0.998 | 0.750 | 0.800 | 0.730 | 0.889 | 0.571 | 0.493 | 0.833 | 0.742 | 0.789 | 0.724 | 0.833 | 0.615 | 0.463 | 0.782 |
| | HS-578T | 0.965 | 0.977 | 0.938 | 1.000 | 0.875 | 0.914 | 1.000 | 0.809 | 0.880 | 0.678 | 0.971 | 0.385 | 0.476 | 0.814 | 0.830 | 0.892 | 0.716 | 0.971 | 0.462 | 0.543 | 0.787 |
| | MCF-7 | 0.954 | 0.959 | 0.953 | 0.962 | 0.944 | 0.907 | 0.995 | 0.808 | 0.829 | 0.805 | 0.835 | 0.775 | 0.611 | 0.884 | 0.804 | 0.823 | 0.802 | 0.820 | 0.784 | 0.604 | 0.884 |
| | MDA-MB-231 | 0.960 | 0.969 | 0.954 | 0.977 | 0.931 | 0.914 | 0.996 | 0.825 | 0.867 | 0.798 | 0.896 | 0.700 | 0.614 | 0.890 | 0.841 | 0.878 | 0.819 | 0.899 | 0.739 | 0.651 | 0.904 |
| | MDA-MB-361 | 0.973 | 0.979 | 0.975 | 0.969 | 0.980 | 0.941 | 0.999 | 0.865 | 0.906 | 0.808 | 1.000 | 0.615 | 0.714 | 0.971 | 0.889 | 0.917 | 0.875 | 0.917 | 0.833 | 0.750 | 0.965 |
| | MDA-MB-435 | 0.958 | 0.974 | 0.913 | 0.993 | 0.834 | 0.875 | 0.997 | 0.835 | 0.902 | 0.659 | 0.975 | 0.343 | 0.448 | 0.903 | 0.835 | 0.901 | 0.675 | 0.962 | 0.388 | 0.454 | 0.870 |
| | MDA-MB-453 | 0.983 | 0.983 | 0.983 | 0.972 | 0.994 | 0.966 | 0.999 | 0.841 | 0.829 | 0.841 | 0.773 | 0.909 | 0.688 | 0.921 | 0.705 | 0.698 | 0.705 | 0.682 | 0.727 | 0.410 | 0.855 |
| | MDA-MB-468 | 0.965 | 0.978 | 0.936 | 0.991 | 0.881 | 0.902 | 0.998 | 0.879 | 0.925 | 0.767 | 0.980 | 0.553 | 0.642 | 0.906 | 0.859 | 0.913 | 0.741 | 0.961 | 0.522 | 0.568 | 0.928 |
| | SK-BR-3 | 0.952 | 0.969 | 0.915 | 0.989 | 0.841 | 0.871 | 0.996 | 0.837 | 0.898 | 0.722 | 0.954 | 0.490 | 0.529 | 0.878 | 0.832 | 0.890 | 0.754 | 0.908 | 0.600 | 0.531 | 0.865 |
| | T-47D | 0.967 | 0.977 | 0.955 | 0.986 | 0.924 | 0.921 | 0.998 | 0.803 | 0.870 | 0.710 | 0.941 | 0.479 | 0.495 | 0.861 | 0.821 | 0.880 | 0.745 | 0.932 | 0.559 | 0.547 | 0.889 |
| RF+XGBoost | Bcap37 | 0.991 | 0.989 | 0.991 | 0.989 | 0.992 | 0.981 | 1.000 | 0.750 | 0.667 | 0.730 | 0.636 | 0.824 | 0.469 | 0.807 | 0.741 | 0.696 | 0.739 | 0.727 | 0.750 | 0.472 | 0.744 |
| | BT-20 | 1.000 | 1.000 | 1.000 | 1.000 | 1.000 | 1.000 | 1.000 | 0.793 | 0.824 | 0.787 | 0.824 | 0.750 | 0.574 | 0.897 | 0.759 | 0.800 | 0.745 | 0.824 | 0.667 | 0.498 | 0.833 |
| | BT-474 | 1.000 | 1.000 | 1.000 | 1.000 | 1.000 | 1.000 | 1.000 | 0.889 | 0.931 | 0.800 | 0.953 | 0.647 | 0.646 | 0.917 | 0.790 | 0.872 | 0.630 | 0.906 | 0.353 | 0.297 | 0.840 |
| | BT-549 | 0.988 | 0.991 | 0.986 | 0.994 | 0.977 | 0.973 | 1.000 | 0.847 | 0.895 | 0.784 | 0.963 | 0.605 | 0.640 | 0.857 | 0.720 | 0.805 | 0.649 | 0.850 | 0.447 | 0.323 | 0.679 |
| | HBL-100 | 0.980 | 0.983 | 0.983 | 0.966 | 1.000 | 0.960 | 0.999 | 0.719 | 0.769 | 0.702 | 0.833 | 0.571 | 0.423 | 0.837 | 0.839 | 0.865 | 0.829 | 0.889 | 0.769 | 0.667 | 0.823 |
| | HS-578T | 0.968 | 0.978 | 0.942 | 1.000 | 0.885 | 0.920 | 1.000 | 0.809 | 0.880 | 0.678 | 0.971 | 0.385 | 0.476 | 0.769 | 0.830 | 0.892 | 0.716 | 0.971 | 0.462 | 0.543 | 0.794 |
| | MCF-7 | 0.986 | 0.987 | 0.986 | 0.988 | 0.983 | 0.971 | 0.999 | 0.810 | 0.830 | 0.807 | 0.838 | 0.775 | 0.615 | 0.886 | 0.805 | 0.825 | 0.802 | 0.831 | 0.773 | 0.605 | 0.885 |
| | MDA-MB-231 | 0.993 | 0.994 | 0.992 | 0.995 | 0.988 | 0.984 | 1.000 | 0.829 | 0.869 | 0.808 | 0.885 | 0.732 | 0.627 | 0.895 | 0.834 | 0.873 | 0.811 | 0.895 | 0.727 | 0.635 | 0.904 |
| | MDA-MB-361 | 0.990 | 0.992 | 0.990 | 0.990 | 0.990 | 0.977 | 1.000 | 0.838 | 0.885 | 0.787 | 0.958 | 0.615 | 0.638 | 0.971 | 0.917 | 0.936 | 0.917 | 0.917 | 0.917 | 0.818 | 0.979 |
| | MDA-MB-435 | 0.989 | 0.993 | 0.980 | 0.997 | 0.963 | 0.969 | 0.999 | 0.842 | 0.905 | 0.685 | 0.966 | 0.403 | 0.479 | 0.911 | 0.832 | 0.898 | 0.684 | 0.949 | 0.418 | 0.450 | 0.869 |
| | MDA-MB-453 | 0.991 | 0.992 | 0.992 | 0.989 | 0.994 | 0.983 | 1.000 | 0.750 | 0.744 | 0.750 | 0.727 | 0.773 | 0.501 | 0.893 | 0.773 | 0.750 | 0.773 | 0.682 | 0.864 | 0.555 | 0.826 |
| | MDA-MB-468 | 0.987 | 0.991 | 0.975 | 0.997 | 0.954 | 0.963 | 1.000 | 0.879 | 0.925 | 0.774 | 0.974 | 0.574 | 0.642 | 0.908 | 0.864 | 0.915 | 0.752 | 0.961 | 0.543 | 0.586 | 0.925 |
| | SK-BR-3 | 0.981 | 0.988 | 0.965 | 0.998 | 0.933 | 0.950 | 0.999 | 0.828 | 0.891 | 0.715 | 0.941 | 0.490 | 0.501 | 0.874 | 0.817 | 0.882 | 0.724 | 0.908 | 0.540 | 0.481 | 0.860 |
| | T-47D | 0.986 | 0.990 | 0.980 | 0.996 | 0.964 | 0.968 | 1.000 | 0.787 | 0.859 | 0.692 | 0.927 | 0.457 | 0.451 | 0.863 | 0.805 | 0.871 | 0.715 | 0.936 | 0.495 | 0.500 | 0.871 |
| SVM+XGBoost | Bcap37 | 0.932 | 0.910 | 0.920 | 0.864 | 0.977 | 0.859 | 0.990 | 0.750 | 0.667 | 0.730 | 0.636 | 0.824 | 0.469 | 0.786 | 0.704 | 0.636 | 0.693 | 0.636 | 0.750 | 0.386 | 0.722 |
| | BT-20 | 0.936 | 0.945 | 0.936 | 0.934 | 0.938 | 0.869 | 0.990 | 0.828 | 0.848 | 0.828 | 0.824 | 0.833 | 0.651 | 0.897 | 0.690 | 0.757 | 0.662 | 0.824 | 0.500 | 0.344 | 0.804 |
| | BT-474 | 0.975 | 0.984 | 0.952 | 0.992 | 0.912 | 0.925 | 0.998 | 0.901 | 0.938 | 0.830 | 0.953 | 0.706 | 0.691 | 0.915 | 0.827 | 0.891 | 0.739 | 0.891 | 0.588 | 0.479 | 0.859 |
| | BT-549 | 0.947 | 0.962 | 0.930 | 0.980 | 0.880 | 0.879 | 0.990 | 0.856 | 0.901 | 0.797 | 0.963 | 0.632 | 0.661 | 0.831 | 0.703 | 0.788 | 0.643 | 0.813 | 0.474 | 0.298 | 0.683 |
| | HBL-100 | 0.945 | 0.952 | 0.946 | 0.939 | 0.953 | 0.887 | 0.989 | 0.719 | 0.769 | 0.702 | 0.833 | 0.571 | 0.423 | 0.825 | 0.806 | 0.850 | 0.780 | 0.944 | 0.615 | 0.609 | 0.803 |
| | HS-578T | 0.909 | 0.941 | 0.842 | 0.993 | 0.692 | 0.770 | 0.981 | 0.809 | 0.880 | 0.678 | 0.971 | 0.385 | 0.476 | 0.749 | 0.809 | 0.880 | 0.678 | 0.971 | 0.385 | 0.476 | 0.826 |
| | MCF-7 | 0.918 | 0.926 | 0.917 | 0.926 | 0.908 | 0.834 | 0.981 | 0.803 | 0.823 | 0.800 | 0.826 | 0.773 | 0.600 | 0.880 | 0.805 | 0.824 | 0.802 | 0.823 | 0.782 | 0.605 | 0.880 |
| | MDA-MB-231 | 0.946 | 0.958 | 0.939 | 0.967 | 0.911 | 0.883 | 0.990 | 0.819 | 0.861 | 0.795 | 0.882 | 0.707 | 0.602 | 0.890 | 0.826 | 0.866 | 0.804 | 0.882 | 0.727 | 0.619 | 0.900 |
| | MDA-MB-361 | 0.973 | 0.979 | 0.975 | 0.969 | 0.980 | 0.941 | 0.995 | 0.838 | 0.885 | 0.787 | 0.958 | 0.615 | 0.638 | 0.971 | 0.889 | 0.917 | 0.875 | 0.917 | 0.833 | 0.750 | 0.979 |
| | MDA-MB-435 | 0.926 | 0.954 | 0.849 | 0.987 | 0.712 | 0.776 | 0.989 | 0.828 | 0.898 | 0.655 | 0.966 | 0.343 | 0.424 | 0.909 | 0.838 | 0.902 | 0.688 | 0.958 | 0.418 | 0.471 | 0.870 |
| | MDA-MB-453 | 0.980 | 0.981 | 0.980 | 0.972 | 0.988 | 0.960 | 0.998 | 0.750 | 0.744 | 0.750 | 0.727 | 0.773 | 0.501 | 0.911 | 0.750 | 0.732 | 0.750 | 0.682 | 0.818 | 0.505 | 0.837 |
| | MDA-MB-468 | 0.949 | 0.967 | 0.904 | 0.989 | 0.819 | 0.854 | 0.993 | 0.884 | 0.928 | 0.777 | 0.980 | 0.574 | 0.658 | 0.890 | 0.864 | 0.915 | 0.752 | 0.961 | 0.543 | 0.586 | 0.908 |
| | SK-BR-3 | 0.909 | 0.942 | 0.838 | 0.979 | 0.697 | 0.746 | 0.979 | 0.837 | 0.898 | 0.722 | 0.954 | 0.490 | 0.529 | 0.876 | 0.827 | 0.889 | 0.731 | 0.921 | 0.540 | 0.504 | 0.868 |
| | T-47D | 0.929 | 0.951 | 0.898 | 0.974 | 0.822 | 0.828 | 0.986 | 0.803 | 0.869 | 0.713 | 0.936 | 0.489 | 0.496 | 0.866 | 0.802 | 0.867 | 0.723 | 0.918 | 0.527 | 0.496 | 0.865 |
| RF+SVM+XGBoost | Bcap37 | 0.964 | 0.953 | 0.958 | 0.932 | 0.985 | 0.924 | 0.996 | 0.786 | 0.700 | 0.759 | 0.636 | 0.882 | 0.542 | 0.802 | 0.741 | 0.667 | 0.724 | 0.636 | 0.813 | 0.457 | 0.739 |
| | BT-20 | 0.962 | 0.967 | 0.960 | 0.971 | 0.948 | 0.921 | 0.998 | 0.793 | 0.813 | 0.799 | 0.765 | 0.833 | 0.589 | 0.892 | 0.724 | 0.778 | 0.703 | 0.824 | 0.583 | 0.422 | 0.843 |
| | BT-474 | 0.975 | 0.985 | 0.950 | 0.994 | 0.905 | 0.925 | 0.999 | 0.877 | 0.924 | 0.771 | 0.953 | 0.588 | 0.601 | 0.911 | 0.802 | 0.881 | 0.637 | 0.922 | 0.353 | 0.327 | 0.865 |
| | BT-549 | 0.977 | 0.983 | 0.969 | 0.991 | 0.948 | 0.947 | 0.999 | 0.831 | 0.884 | 0.764 | 0.950 | 0.579 | 0.596 | 0.873 | 0.763 | 0.835 | 0.694 | 0.888 | 0.500 | 0.426 | 0.708 |
| | HBL-100 | 0.968 | 0.973 | 0.969 | 0.966 | 0.972 | 0.935 | 0.998 | 0.750 | 0.800 | 0.730 | 0.889 | 0.571 | 0.493 | 0.833 | 0.742 | 0.789 | 0.724 | 0.833 | 0.615 | 0.463 | 0.782 |
| | HS-578T | 0.965 | 0.977 | 0.938 | 1.000 | 0.875 | 0.914 | 1.000 | 0.809 | 0.880 | 0.678 | 0.971 | 0.385 | 0.476 | 0.814 | 0.830 | 0.892 | 0.716 | 0.971 | 0.462 | 0.543 | 0.787 |
| | MCF-7 | 0.954 | 0.959 | 0.953 | 0.962 | 0.944 | 0.907 | 0.995 | 0.808 | 0.829 | 0.805 | 0.835 | 0.775 | 0.611 | 0.884 | 0.804 | 0.823 | 0.802 | 0.820 | 0.784 | 0.604 | 0.884 |
| | MDA-MB-231 | 0.960 | 0.969 | 0.954 | 0.977 | 0.931 | 0.914 | 0.996 | 0.825 | 0.867 | 0.798 | 0.896 | 0.700 | 0.614 | 0.890 | 0.841 | 0.878 | 0.819 | 0.899 | 0.739 | 0.651 | 0.904 |
| | MDA-MB-361 | 0.973 | 0.979 | 0.975 | 0.969 | 0.980 | 0.941 | 0.999 | 0.865 | 0.906 | 0.808 | 1.000 | 0.615 | 0.714 | 0.971 | 0.889 | 0.917 | 0.875 | 0.917 | 0.833 | 0.750 | 0.965 |
| | MDA-MB-435 | 0.958 | 0.974 | 0.913 | 0.993 | 0.834 | 0.875 | 0.997 | 0.835 | 0.902 | 0.659 | 0.975 | 0.343 | 0.448 | 0.903 | 0.835 | 0.901 | 0.675 | 0.962 | 0.388 | 0.454 | 0.870 |
| | MDA-MB-453 | 0.983 | 0.983 | 0.983 | 0.972 | 0.994 | 0.966 | 0.999 | 0.841 | 0.829 | 0.841 | 0.773 | 0.909 | 0.688 | 0.921 | 0.705 | 0.698 | 0.705 | 0.682 | 0.727 | 0.410 | 0.855 |
| | MDA-MB-468 | 0.965 | 0.978 | 0.936 | 0.991 | 0.881 | 0.902 | 0.998 | 0.879 | 0.925 | 0.767 | 0.980 | 0.553 | 0.642 | 0.906 | 0.859 | 0.913 | 0.741 | 0.961 | 0.522 | 0.568 | 0.928 |
| | SK-BR-3 | 0.952 | 0.969 | 0.915 | 0.989 | 0.841 | 0.871 | 0.996 | 0.837 | 0.898 | 0.722 | 0.954 | 0.490 | 0.529 | 0.878 | 0.832 | 0.890 | 0.754 | 0.908 | 0.600 | 0.531 | 0.865 |
| | T-47D | 0.967 | 0.977 | 0.955 | 0.986 | 0.924 | 0.921 | 0.998 | 0.803 | 0.870 | 0.710 | 0.941 | 0.479 | 0.495 | 0.861 | 0.821 | 0.880 | 0.745 | 0.932 | 0.559 | 0.547 | 0.889 |

[a] ACC: Accuracy. [b] F1: F1-measure. [c] BA: Balanced accuracy. [d] SE:Sensitivity. [e] SP:Specificity. [f] MCC:Matthews correlation coefficient. [g] AUC:The area under receiver operating characteristic.

**Table S11.** The performance results of stacking models based on Morgan fingerprints.

| Methods | Cell lines | Training set | | | | | | | Validation set | | | | | | | Test set | | | | | | |
|---|---|---|---|---|---|---|---|---|---|---|---|---|---|---|---|---|---|---|---|---|---|---|
| | | ACC[a] | F1[b] | BA[c] | SE[d] | SP[e] | MCC[f] | AUC[g] | ACC | F1 | BA | SE | SP | MCC | AUC | ACC | F1 | BA | SE | SP | MCC | AUC |
| RF+SVM | Bcap37 | 0.986 | 0.983 | 0.983 | 0.966 | 1.000 | 0.972 | 1.000 | 0.750 | 0.632 | 0.714 | 0.545 | 0.882 | 0.462 | 0.786 | 0.667 | 0.526 | 0.634 | 0.455 | 0.813 | 0.287 | 0.699 |
| | BT-20 | 0.996 | 0.996 | 0.995 | 1.000 | 0.990 | 0.991 | 1.000 | 0.759 | 0.788 | 0.757 | 0.765 | 0.750 | 0.510 | 0.868 | 0.690 | 0.757 | 0.662 | 0.824 | 0.500 | 0.344 | 0.843 |
| | BT-474 | 0.974 | 0.984 | 0.938 | 1.000 | 0.876 | 0.921 | 0.999 | 0.852 | 0.914 | 0.647 | 1.000 | 0.294 | 0.498 | 0.919 | 0.778 | 0.875 | 0.492 | 0.984 | 0.000 | -0.058 | 0.868 |
| | BT-549 | 0.962 | 0.972 | 0.952 | 0.980 | 0.925 | 0.913 | 0.996 | 0.839 | 0.890 | 0.771 | 0.963 | 0.579 | 0.619 | 0.870 | 0.746 | 0.821 | 0.681 | 0.863 | 0.500 | 0.389 | 0.709 |
| | HBL-100 | 0.972 | 0.976 | 0.970 | 0.986 | 0.953 | 0.943 | 0.999 | 0.750 | 0.800 | 0.730 | 0.889 | 0.571 | 0.493 | 0.825 | 0.710 | 0.780 | 0.675 | 0.889 | 0.462 | 0.395 | 0.821 |
| | HS-578T | 0.723 | 0.839 | 0.500 | 1.000 | 0.000 | nan | 1.000 | 0.723 | 0.840 | 0.500 | 1.000 | 0.000 | nan | 0.810 | 0.723 | 0.840 | 0.500 | 1.000 | 0.000 | nan | 0.769 |
| | MCF-7 | 0.978 | 0.981 | 0.977 | 0.987 | 0.968 | 0.956 | 0.998 | 0.805 | 0.827 | 0.801 | 0.839 | 0.763 | 0.604 | 0.886 | 0.806 | 0.827 | 0.802 | 0.836 | 0.767 | 0.606 | 0.886 |
| | MDA-MB-231 | 0.963 | 0.971 | 0.952 | 0.993 | 0.911 | 0.920 | 0.998 | 0.822 | 0.869 | 0.784 | 0.924 | 0.643 | 0.606 | 0.889 | 0.831 | 0.875 | 0.796 | 0.924 | 0.667 | 0.627 | 0.904 |
| | MDA-MB-361 | 0.983 | 0.987 | 0.982 | 0.984 | 0.980 | 0.962 | 0.999 | 0.838 | 0.889 | 0.769 | 1.000 | 0.538 | 0.656 | 0.974 | 0.861 | 0.898 | 0.833 | 0.917 | 0.750 | 0.682 | 0.969 |
| | MDA-MB-435 | 0.969 | 0.980 | 0.931 | 0.998 | 0.864 | 0.908 | 0.999 | 0.835 | 0.903 | 0.648 | 0.983 | 0.313 | 0.447 | 0.906 | 0.828 | 0.898 | 0.644 | 0.975 | 0.313 | 0.420 | 0.878 |
| | MDA-MB-453 | 0.983 | 0.983 | 0.983 | 0.972 | 0.994 | 0.966 | 0.999 | 0.818 | 0.810 | 0.818 | 0.773 | 0.864 | 0.639 | 0.913 | 0.773 | 0.773 | 0.773 | 0.773 | 0.773 | 0.545 | 0.857 |
| | MDA-MB-468 | 0.982 | 0.989 | 0.965 | 0.998 | 0.933 | 0.950 | 1.000 | 0.894 | 0.934 | 0.799 | 0.980 | 0.617 | 0.691 | 0.909 | 0.848 | 0.906 | 0.727 | 0.954 | 0.500 | 0.535 | 0.921 |
| | SK-BR-3 | 0.985 | 0.990 | 0.971 | 0.999 | 0.943 | 0.960 | 1.000 | 0.833 | 0.896 | 0.699 | 0.967 | 0.431 | 0.509 | 0.877 | 0.827 | 0.890 | 0.717 | 0.934 | 0.500 | 0.495 | 0.871 |
| | T-47D | 0.972 | 0.980 | 0.954 | 0.998 | 0.911 | 0.934 | 0.999 | 0.790 | 0.864 | 0.679 | 0.955 | 0.404 | 0.457 | 0.867 | 0.821 | 0.883 | 0.727 | 0.959 | 0.495 | 0.545 | 0.900 |
| RF+XGBoost | Bcap37 | 0.986 | 0.983 | 0.983 | 0.966 | 1.000 | 0.972 | 1.000 | 0.750 | 0.667 | 0.730 | 0.636 | 0.824 | 0.469 | 0.802 | 0.704 | 0.600 | 0.679 | 0.545 | 0.813 | 0.373 | 0.727 |
| | BT-20 | 1.000 | 1.000 | 1.000 | 1.000 | 1.000 | 1.000 | 1.000 | 0.793 | 0.824 | 0.787 | 0.824 | 0.750 | 0.574 | 0.902 | 0.690 | 0.757 | 0.662 | 0.824 | 0.500 | 0.344 | 0.843 |
| | BT-474 | 0.969 | 0.981 | 0.927 | 1.000 | 0.854 | 0.907 | 1.000 | 0.864 | 0.921 | 0.676 | 1.000 | 0.353 | 0.549 | 0.926 | 0.765 | 0.865 | 0.506 | 0.953 | 0.059 | 0.022 | 0.863 |
| | BT-549 | 0.995 | 0.996 | 0.993 | 0.998 | 0.987 | 0.988 | 1.000 | 0.847 | 0.898 | 0.770 | 0.988 | 0.553 | 0.648 | 0.869 | 0.771 | 0.846 | 0.686 | 0.925 | 0.447 | 0.439 | 0.677 |
| | HBL-100 | 0.992 | 0.993 | 0.991 | 1.000 | 0.981 | 0.984 | 1.000 | 0.750 | 0.789 | 0.738 | 0.833 | 0.643 | 0.488 | 0.857 | 0.806 | 0.842 | 0.791 | 0.889 | 0.692 | 0.599 | 0.838 |
| | HS-578T | 0.723 | 0.839 | 0.500 | 1.000 | 0.000 | nan | 1.000 | 0.723 | 0.840 | 0.500 | 1.000 | 0.000 | nan | 0.824 | 0.723 | 0.840 | 0.500 | 1.000 | 0.000 | nan | 0.708 |
| | MCF-7 | 0.992 | 0.993 | 0.992 | 0.994 | 0.989 | 0.984 | 1.000 | 0.802 | 0.825 | 0.797 | 0.842 | 0.752 | 0.598 | 0.881 | 0.805 | 0.827 | 0.801 | 0.839 | 0.762 | 0.604 | 0.879 |
| | MDA-MB-231 | 0.995 | 0.996 | 0.994 | 0.998 | 0.990 | 0.989 | 1.000 | 0.821 | 0.868 | 0.784 | 0.920 | 0.648 | 0.604 | 0.886 | 0.826 | 0.871 | 0.790 | 0.922 | 0.658 | 0.615 | 0.899 |
| | MDA-MB-361 | 0.993 | 0.995 | 0.992 | 0.995 | 0.990 | 0.985 | 1.000 | 0.838 | 0.889 | 0.769 | 1.000 | 0.538 | 0.656 | 0.974 | 0.917 | 0.939 | 0.896 | 0.958 | 0.833 | 0.810 | 0.972 |
| | MDA-MB-435 | 0.991 | 0.994 | 0.980 | 0.999 | 0.961 | 0.974 | 1.000 | 0.838 | 0.905 | 0.650 | 0.987 | 0.313 | 0.462 | 0.904 | 0.822 | 0.895 | 0.634 | 0.970 | 0.299 | 0.392 | 0.879 |
| | MDA-MB-453 | 0.994 | 0.995 | 0.994 | 1.000 | 0.988 | 0.989 | 1.000 | 0.841 | 0.837 | 0.841 | 0.818 | 0.864 | 0.683 | 0.924 | 0.727 | 0.727 | 0.727 | 0.727 | 0.727 | 0.455 | 0.835 |
| | MDA-MB-468 | 0.999 | 1.000 | 0.999 | 1.000 | 0.997 | 0.998 | 1.000 | 0.889 | 0.931 | 0.795 | 0.974 | 0.617 | 0.675 | 0.911 | 0.864 | 0.915 | 0.760 | 0.954 | 0.565 | 0.588 | 0.925 |
| | SK-BR-3 | 0.994 | 0.996 | 0.989 | 1.000 | 0.978 | 0.985 | 1.000 | 0.823 | 0.891 | 0.680 | 0.967 | 0.392 | 0.474 | 0.875 | 0.817 | 0.884 | 0.704 | 0.928 | 0.480 | 0.465 | 0.873 |
| | T-47D | 0.993 | 0.995 | 0.988 | 0.999 | 0.977 | 0.983 | 1.000 | 0.787 | 0.862 | 0.677 | 0.950 | 0.404 | 0.447 | 0.870 | 0.802 | 0.872 | 0.695 | 0.959 | 0.430 | 0.489 | 0.882 |
| SVM+XGBoost | Bcap37 | 0.959 | 0.947 | 0.951 | 0.909 | 0.992 | 0.916 | 0.997 | 0.750 | 0.667 | 0.730 | 0.636 | 0.824 | 0.469 | 0.775 | 0.704 | 0.600 | 0.679 | 0.545 | 0.813 | 0.373 | 0.693 |
| | BT-20 | 0.966 | 0.971 | 0.963 | 0.978 | 0.948 | 0.929 | 0.999 | 0.793 | 0.824 | 0.787 | 0.824 | 0.750 | 0.574 | 0.887 | 0.724 | 0.789 | 0.691 | 0.882 | 0.500 | 0.421 | 0.838 |
| | BT-474 | 0.972 | 0.983 | 0.934 | 1.000 | 0.869 | 0.916 | 0.999 | 0.877 | 0.926 | 0.727 | 0.984 | 0.471 | 0.590 | 0.925 | 0.765 | 0.867 | 0.484 | 0.969 | 0.000 | -0.082 | 0.864 |
| | BT-549 | 0.958 | 0.969 | 0.947 | 0.978 | 0.916 | 0.903 | 0.992 | 0.831 | 0.884 | 0.764 | 0.950 | 0.579 | 0.596 | 0.867 | 0.737 | 0.817 | 0.668 | 0.863 | 0.474 | 0.365 | 0.718 |
| | HBL-100 | 0.988 | 0.990 | 0.987 | 0.993 | 0.981 | 0.976 | 0.999 | 0.750 | 0.789 | 0.738 | 0.833 | 0.643 | 0.488 | 0.849 | 0.806 | 0.842 | 0.791 | 0.889 | 0.692 | 0.599 | 0.825 |
| | HS-578T | 0.723 | 0.839 | 0.500 | 1.000 | 0.000 | nan | 0.980 | 0.723 | 0.840 | 0.500 | 1.000 | 0.000 | nan | 0.771 | 0.723 | 0.840 | 0.500 | 1.000 | 0.000 | nan | 0.778 |
| | MCF-7 | 0.899 | 0.910 | 0.897 | 0.919 | 0.875 | 0.796 | 0.962 | 0.793 | 0.816 | 0.788 | 0.827 | 0.750 | 0.579 | 0.864 | 0.796 | 0.818 | 0.792 | 0.825 | 0.759 | 0.586 | 0.866 |
| | MDA-MB-231 | 0.896 | 0.922 | 0.872 | 0.961 | 0.782 | 0.774 | 0.958 | 0.809 | 0.858 | 0.771 | 0.909 | 0.633 | 0.576 | 0.851 | 0.817 | 0.863 | 0.783 | 0.906 | 0.660 | 0.595 | 0.863 |
| | MDA-MB-361 | 0.973 | 0.979 | 0.975 | 0.969 | 0.980 | 0.941 | 0.997 | 0.838 | 0.889 | 0.769 | 1.000 | 0.538 | 0.656 | 0.974 | 0.889 | 0.917 | 0.875 | 0.917 | 0.833 | 0.750 | 0.976 |
| | MDA-MB-435 | 0.898 | 0.938 | 0.775 | 0.995 | 0.555 | 0.686 | 0.989 | 0.818 | 0.894 | 0.611 | 0.983 | 0.239 | 0.371 | 0.898 | 0.828 | 0.899 | 0.639 | 0.979 | 0.299 | 0.418 | 0.857 |
| | MDA-MB-453 | 0.972 | 0.972 | 0.972 | 0.956 | 0.988 | 0.944 | 0.997 | 0.795 | 0.791 | 0.795 | 0.773 | 0.818 | 0.592 | 0.903 | 0.750 | 0.744 | 0.750 | 0.727 | 0.773 | 0.501 | 0.864 |
| | MDA-MB-468 | 0.945 | 0.965 | 0.893 | 0.990 | 0.795 | 0.841 | 0.993 | 0.869 | 0.920 | 0.745 | 0.980 | 0.511 | 0.609 | 0.894 | 0.848 | 0.907 | 0.719 | 0.961 | 0.478 | 0.532 | 0.913 |
| | SK-BR-3 | 0.901 | 0.937 | 0.810 | 0.990 | 0.630 | 0.724 | 0.987 | 0.818 | 0.889 | 0.663 | 0.974 | 0.353 | 0.456 | 0.876 | 0.832 | 0.894 | 0.720 | 0.941 | 0.500 | 0.508 | 0.862 |
| | T-47D | 0.931 | 0.952 | 0.898 | 0.981 | 0.814 | 0.833 | 0.990 | 0.796 | 0.867 | 0.693 | 0.950 | 0.436 | 0.476 | 0.857 | 0.824 | 0.885 | 0.732 | 0.959 | 0.505 | 0.554 | 0.887 |
| RF+SVM+XGBoost | Bcap37 | 0.986 | 0.983 | 0.983 | 0.966 | 1.000 | 0.972 | 1.000 | 0.750 | 0.667 | 0.730 | 0.636 | 0.824 | 0.469 | 0.807 | 0.704 | 0.600 | 0.679 | 0.545 | 0.813 | 0.373 | 0.727 |
| | BT-20 | 0.996 | 0.996 | 0.995 | 1.000 | 0.990 | 0.991 | 1.000 | 0.793 | 0.813 | 0.799 | 0.765 | 0.833 | 0.589 | 0.887 | 0.690 | 0.757 | 0.662 | 0.824 | 0.500 | 0.344 | 0.833 |
| | BT-474 | 0.972 | 0.983 | 0.934 | 1.000 | 0.869 | 0.916 | 1.000 | 0.877 | 0.928 | 0.706 | 1.000 | 0.412 | 0.597 | 0.923 | 0.765 | 0.867 | 0.484 | 0.969 | 0.000 | -0.082 | 0.862 |
| | BT-549 | 0.962 | 0.972 | 0.952 | 0.980 | 0.925 | 0.913 | 0.996 | 0.839 | 0.890 | 0.771 | 0.963 | 0.579 | 0.619 | 0.871 | 0.737 | 0.817 | 0.668 | 0.863 | 0.474 | 0.365 | 0.709 |
| | HBL-100 | 0.988 | 0.990 | 0.987 | 0.993 | 0.981 | 0.976 | 0.999 | 0.750 | 0.789 | 0.738 | 0.833 | 0.643 | 0.488 | 0.849 | 0.806 | 0.842 | 0.791 | 0.889 | 0.692 | 0.599 | 0.842 |
| | HS-578T | 0.723 | 0.839 | 0.500 | 1.000 | 0.000 | nan | 1.000 | 0.723 | 0.840 | 0.500 | 1.000 | 0.000 | nan | 0.819 | 0.723 | 0.840 | 0.500 | 1.000 | 0.000 | nan | 0.733 |
| | MCF-7 | 0.985 | 0.987 | 0.984 | 0.990 | 0.979 | 0.970 | 0.999 | 0.811 | 0.833 | 0.807 | 0.846 | 0.767 | 0.616 | 0.884 | 0.807 | 0.828 | 0.804 | 0.834 | 0.773 | 0.609 | 0.883 |
| | MDA-MB-231 | 0.975 | 0.981 | 0.967 | 0.995 | 0.939 | 0.946 | 0.998 | 0.821 | 0.867 | 0.786 | 0.916 | 0.655 | 0.604 | 0.877 | 0.835 | 0.876 | 0.804 | 0.916 | 0.692 | 0.635 | 0.895 |
| | MDA-MB-361 | 0.983 | 0.987 | 0.982 | 0.984 | 0.980 | 0.962 | 0.999 | 0.838 | 0.889 | 0.769 | 1.000 | 0.538 | 0.656 | 0.978 | 0.889 | 0.917 | 0.875 | 0.917 | 0.833 | 0.750 | 0.969 |
| | MDA-MB-435 | 0.968 | 0.980 | 0.929 | 0.998 | 0.860 | 0.905 | 0.999 | 0.835 | 0.903 | 0.648 | 0.983 | 0.313 | 0.447 | 0.906 | 0.828 | 0.898 | 0.644 | 0.975 | 0.313 | 0.420 | 0.878 |
| | MDA-MB-453 | 0.983 | 0.983 | 0.983 | 0.972 | 0.994 | 0.966 | 0.999 | 0.795 | 0.791 | 0.795 | 0.773 | 0.818 | 0.592 | 0.921 | 0.773 | 0.773 | 0.773 | 0.773 | 0.773 | 0.545 | 0.857 |
| | MDA-MB-468 | 0.982 | 0.989 | 0.965 | 0.998 | 0.933 | 0.950 | 1.000 | 0.894 | 0.934 | 0.799 | 0.980 | 0.617 | 0.691 | 0.909 | 0.848 | 0.906 | 0.727 | 0.954 | 0.500 | 0.535 | 0.921 |
| | SK-BR-3 | 0.986 | 0.991 | 0.972 | 0.999 | 0.945 | 0.962 | 1.000 | 0.833 | 0.896 | 0.699 | 0.967 | 0.431 | 0.509 | 0.876 | 0.827 | 0.890 | 0.717 | 0.934 | 0.500 | 0.495 | 0.870 |
| | T-47D | 0.967 | 0.977 | 0.947 | 0.996 | 0.899 | 0.921 | 0.999 | 0.796 | 0.869 | 0.684 | 0.964 | 0.404 | 0.476 | 0.869 | 0.818 | 0.881 | 0.721 | 0.959 | 0.484 | 0.536 | 0.895 |

[a] ACC: Accuracy. [b] F1: F1-measure. [c] BA: Balanced accuracy. [d] SE:Sensitivity. [e] SP:Specificity. [f] MCC:Matthews correlation coefficient. [g] AUC:The area under receiver operating characteristic.

**Table S12.** The performance results of 10-fold cross validation models based on Morgan fingerprints.

| Methods | Cell lines | Training set | | | | Validation set | | | | Test set | | | |
|---|---|---|---|---|---|---|---|---|---|---|---|---|---|
| | | ACC[a] | F1[b] | BA[c] | AUC[d] | ACC | F1 | BA | AUC | ACC | F1 | BA | AUC |
| RF::Morgan | Bcap37 | 0.779 ± 0.019 | 0.661 ± 0.039 | 0.739 ± 0.024 | 0.892 ± 0.010 | 0.746 ± 0.043 | 0.569 ± 0.118 | 0.693 ± 0.055 | 0.844 ± 0.031 | 0.685 ± 0.047 | 0.582 ± 0.047 | 0.662 ± 0.042 | 0.742 ± 0.048 |
| | BT-20 | 0.918 ± 0.017 | 0.931 ± 0.014 | 0.914 ± 0.019 | 0.980 ± 0.005 | 0.800 ± 0.032 | 0.829 ± 0.027 | 0.794 ± 0.038 | 0.900 ± 0.023 | 0.724 ± 0.054 | 0.781 ± 0.043 | 0.700 ± 0.056 | 0.810 ± 0.053 |
| | BT-474 | 1.000 ± 0.000 | 1.000 ± 0.000 | 1.000 ± 0.000 | 1.000 ± 0.000 | 0.874 ± 0.018 | 0.925 ± 0.010 | 0.717 ± 0.041 | 0.928 ± 0.009 | 0.815 ± 0.020 | 0.888 ± 0.011 | 0.652 ± 0.046 | 0.868 ± 0.012 |
| | BT-549 | 0.998 ± 0.001 | 0.999 ± 0.001 | 0.998 ± 0.001 | 1.000 ± 0.000 | 0.826 ± 0.014 | 0.880 ± 0.009 | 0.762 ± 0.024 | 0.877 ± 0.007 | 0.751 ± 0.018 | 0.828 ± 0.013 | 0.676 ± 0.021 | 0.704 ± 0.016 |
| | HBL-100 | 0.996 ± 0.002 | 0.997 ± 0.002 | 0.996 ± 0.002 | 1.000 ± 0.000 | 0.766 ± 0.027 | 0.808 ± 0.021 | 0.750 ± 0.029 | 0.861 ± 0.010 | 0.765 ± 0.068 | 0.804 ± 0.063 | 0.750 ± 0.066 | 0.809 ± 0.025 |
| | HS-578T | 1.000 ± 0.000 | 1.000 ± 0.000 | 1.000 ± 0.000 | 1.000 ± 0.000 | 0.815 ± 0.023 | 0.884 ± 0.014 | 0.682 ± 0.033 | 0.843 ± 0.022 | 0.809 ± 0.030 | 0.874 ± 0.020 | 0.720 ± 0.036 | 0.749 ± 0.019 |
| | MCF-7 | 0.993 ± 0.000 | 0.993 ± 0.000 | 0.992 ± 0.000 | 1.000 ± 0.000 | 0.808 ± 0.002 | 0.829 ± 0.002 | 0.805 ± 0.002 | 0.884 ± 0.001 | 0.806 ± 0.002 | 0.827 ± 0.002 | 0.803 ± 0.002 | 0.882 ± 0.001 |
| | MDA-MB-231 | 0.995 ± 0.000 | 0.996 ± 0.000 | 0.995 ± 0.000 | 1.000 ± 0.000 | 0.832 ± 0.006 | 0.873 ± 0.004 | 0.803 ± 0.007 | 0.891 ± 0.003 | 0.831 ± 0.003 | 0.873 ± 0.002 | 0.803 ± 0.004 | 0.900 ± 0.003 |
| | MDA-MB-361 | 0.997 ± 0.002 | 0.998 ± 0.001 | 0.997 ± 0.002 | 1.000 ± 0.000 | 0.854 ± 0.036 | 0.898 ± 0.023 | 0.798 ± 0.054 | 0.981 ± 0.007 | 0.878 ± 0.030 | 0.910 ± 0.022 | 0.856 ± 0.036 | 0.960 ± 0.013 |
| | MDA-MB-435 | 0.996 ± 0.001 | 0.997 ± 0.000 | 0.992 ± 0.001 | 1.000 ± 0.000 | 0.838 ± 0.010 | 0.903 ± 0.006 | 0.678 ± 0.019 | 0.902 ± 0.006 | 0.825 ± 0.008 | 0.894 ± 0.005 | 0.672 ± 0.015 | 0.861 ± 0.005 |
| | MDA-MB-453 | 0.995 ± 0.002 | 0.995 ± 0.002 | 0.995 ± 0.002 | 1.000 ± 0.000 | 0.843 ± 0.017 | 0.838 ± 0.019 | 0.843 ± 0.017 | 0.934 ± 0.011 | 0.741 ± 0.022 | 0.728 ± 0.025 | 0.741 ± 0.022 | 0.836 ± 0.010 |
| | MDA-MB-468 | 0.999 ± 0.000 | 1.000 ± 0.000 | 0.999 ± 0.001 | 1.000 ± 0.000 | 0.881 ± 0.010 | 0.926 ± 0.006 | 0.782 ± 0.012 | 0.932 ± 0.005 | 0.862 ± 0.005 | 0.914 ± 0.003 | 0.753 ± 0.009 | 0.917 ± 0.007 |
| | SK-BR-3 | 0.997 ± 0.001 | 0.998 ± 0.000 | 0.996 ± 0.001 | 1.000 ± 0.000 | 0.833 ± 0.011 | 0.894 ± 0.007 | 0.728 ± 0.013 | 0.880 ± 0.013 | 0.819 ± 0.010 | 0.883 ± 0.006 | 0.729 ± 0.017 | 0.860 ± 0.006 |
| | T-47D | 0.998 ± 0.001 | 0.998 ± 0.000 | 0.997 ± 0.001 | 1.000 ± 0.000 | 0.792 ± 0.008 | 0.863 ± 0.005 | 0.700 ± 0.010 | 0.860 ± 0.006 | 0.822 ± 0.012 | 0.881 ± 0.007 | 0.742 ± 0.017 | 0.879 ± 0.006 |
| XGBoost::Morgan | Bcap37 | 0.940 ± 0.054 | 0.923 ± 0.071 | 0.935 ± 0.059 | 0.979 ± 0.028 | 0.729 ± 0.086 | 0.637 ± 0.123 | 0.707 ± 0.091 | 0.744 ± 0.109 | 0.685 ± 0.053 | 0.621 ± 0.065 | 0.678 ± 0.055 | 0.721 ± 0.040 |
| | BT-20 | 0.951 ± 0.034 | 0.958 ± 0.030 | 0.951 ± 0.034 | 0.988 ± 0.013 | 0.762 ± 0.044 | 0.785 ± 0.047 | 0.765 ± 0.040 | 0.869 ± 0.027 | 0.710 ± 0.040 | 0.773 ± 0.030 | 0.683 ± 0.045 | 0.799 ± 0.040 |
| | BT-474 | 0.986 ± 0.011 | 0.991 ± 0.007 | 0.969 ± 0.021 | 0.999 ± 0.002 | 0.873 ± 0.018 | 0.922 ± 0.011 | 0.770 ± 0.040 | 0.882 ± 0.009 | 0.806 ± 0.030 | 0.882 ± 0.018 | 0.657 ± 0.057 | 0.838 ± 0.016 |
| | BT-549 | 0.934 ± 0.050 | 0.954 ± 0.032 | 0.909 ± 0.077 | 0.980 ± 0.035 | 0.797 ± 0.029 | 0.861 ± 0.017 | 0.726 ± 0.045 | 0.796 ± 0.026 | 0.697 ± 0.015 | 0.785 ± 0.013 | 0.629 ± 0.021 | 0.643 ± 0.039 |
| | HBL-100 | 0.962 ± 0.023 | 0.967 ± 0.020 | 0.962 ± 0.023 | 0.993 ± 0.010 | 0.738 ± 0.016 | 0.779 ± 0.012 | 0.725 ± 0.018 | 0.811 ± 0.024 | 0.700 ± 0.040 | 0.739 ± 0.037 | 0.694 ± 0.041 | 0.768 ± 0.023 |
| | HS-578T | 0.904 ± 0.030 | 0.937 ± 0.018 | 0.839 ± 0.054 | 0.974 ± 0.019 | 0.798 ± 0.031 | 0.875 ± 0.020 | 0.651 ± 0.037 | 0.746 ± 0.041 | 0.811 ± 0.029 | 0.879 ± 0.018 | 0.696 ± 0.043 | 0.768 ± 0.018 |
| | MCF-7 | 0.956 ± 0.005 | 0.960 ± 0.004 | 0.956 ± 0.005 | 0.993 ± 0.002 | 0.790 ± 0.006 | 0.812 ± 0.005 | 0.787 ± 0.006 | 0.870 ± 0.003 | 0.793 ± 0.004 | 0.814 ± 0.003 | 0.791 ± 0.004 | 0.871 ± 0.002 |
| | MDA-MB-231 | 0.953 ± 0.020 | 0.963 ± 0.015 | 0.946 ± 0.024 | 0.990 ± 0.008 | 0.805 ± 0.007 | 0.850 ± 0.006 | 0.781 ± 0.008 | 0.869 ± 0.005 | 0.812 ± 0.008 | 0.856 ± 0.006 | 0.789 ± 0.010 | 0.884 ± 0.005 |
| | MDA-MB-361 | 0.974 ± 0.015 | 0.980 ± 0.011 | 0.975 ± 0.014 | 0.993 ± 0.007 | 0.849 ± 0.023 | 0.890 ± 0.017 | 0.808 ± 0.024 | 0.949 ± 0.017 | 0.908 ± 0.023 | 0.930 ± 0.018 | 0.902 ± 0.030 | 0.976 ± 0.008 |
| | MDA-MB-435 | 0.936 ± 0.025 | 0.960 ± 0.015 | 0.872 ± 0.056 | 0.985 ± 0.010 | 0.827 ± 0.013 | 0.896 ± 0.007 | 0.666 ± 0.026 | 0.890 ± 0.014 | 0.821 ± 0.011 | 0.892 ± 0.007 | 0.664 ± 0.020 | 0.851 ± 0.010 |
| | MDA-MB-453 | 0.982 ± 0.011 | 0.982 ± 0.011 | 0.982 ± 0.010 | 0.998 ± 0.001 | 0.750 ± 0.021 | 0.745 ± 0.021 | 0.750 ± 0.021 | 0.862 ± 0.021 | 0.734 ± 0.034 | 0.719 ± 0.035 | 0.734 ± 0.034 | 0.802 ± 0.034 |
| | MDA-MB-468 | 0.946 ± 0.022 | 0.966 ± 0.013 | 0.898 ± 0.047 | 0.989 ± 0.008 | 0.872 ± 0.008 | 0.920 ± 0.004 | 0.768 ± 0.019 | 0.884 ± 0.014 | 0.856 ± 0.009 | 0.910 ± 0.005 | 0.744 ± 0.020 | 0.885 ± 0.013 |
| | SK-BR-3 | 0.962 ± 0.013 | 0.975 ± 0.008 | 0.936 ± 0.022 | 0.992 ± 0.004 | 0.836 ± 0.014 | 0.895 ± 0.010 | 0.746 ± 0.016 | 0.866 ± 0.008 | 0.811 ± 0.013 | 0.877 ± 0.009 | 0.726 ± 0.019 | 0.853 ± 0.007 |
| | T-47D | 0.941 ± 0.020 | 0.959 ± 0.014 | 0.914 ± 0.031 | 0.987 ± 0.007 | 0.787 ± 0.008 | 0.857 ± 0.005 | 0.704 ± 0.012 | 0.841 ± 0.010 | 0.788 ± 0.017 | 0.856 ± 0.011 | 0.710 ± 0.023 | 0.821 ± 0.013 |

[a] ACC: Accuracy. [b] F1: F1-measure. [c] BA: Balanced accuracy. [d] AUC:The area under receiver operating characteristic.
